# Precise exogenous insertion and sequence replacements in poplar by simultaneous HDR overexpression and NHEJ suppression using CRISPR-Cas9

**DOI:** 10.1101/2020.07.04.188219

**Authors:** Ali Movahedi, Hui Wei, Xiaohong Zhou, Jake C. Fountain, Zhong-Hua Chen, Zhiying Mu, Weibo Sun, Jiaxin Zhang, Dawei Li, Baozhu Guo, Rajeev K. Varshney, Liming Yang, Qiang Zhuge

## Abstract

CRISPR-mediated genome editing has become a powerful tool for genetic modification of biological traits. However, developing an efficient, site-specific, gene knock-in system based on homology-directed DNA repair (HDR) remains a significant challenge in plants, especially in woody species like poplar. Here, we show that simultaneous inhibition of non-homologous end joining (NHEJ) recombination cofactor XRCC4 and overexpression of HDR enhancer factors *CtIP and MRE11* can improve the HDR efficiency for gene knock-in. Using this approach, the *BleoR* gene was integrated onto the 3′ end of the *MKK2* MAP Kinase gene to generate a *BleoR-MKK2* fusion protein. Based on exogenous *BleoR* expression, the HDR-mediated knock-in efficiency was up to ∼40-fold greater when using a *XRCC4* silencing incorporated with a combination of *CtIP and MRE11* overexpression compared to no HDR enhancement or NHEJ silencing. Furthermore, this corporation of HDR enhancer overexpression and NHEJ repression also resulted in 7-fold fewer CRISPR-induced Insertions and Deletions (InDels), resulting in no functional effects on *MKK2*-based salt stress responses in poplar. Therefore, this approach may be useful not only in poplar and plants or crops but also in mammalians for improving CRISPR-mediated gene knock-in efficiency.

---

Several studies have been carried out on methods to improve crop genetic modification by CRISPR-mediated donor-dependent HDR^1^, such as increasing *ARGOS8* expression by replacing *GOS2* promoter with HDR and enhancing the efficiency of 35S promoter insertion upstream of *ANT1* gene in tomato ^2^. Several publications have reported the successful generation of null mutations in woody plants using NHEJ pathway since being implemented in poplar ^3,4^. However, precise gene targeting and replacement have only been reported in model plants such as *Arabidopsis* ^5^ and rice ^6^. No report has yet shown efficient HDR for gene replacement in woody perennials.

One of the main limitations of HDR efficiency is inadequate delivery of donor DNA patterns (DDPs) into nuclei. Previous studies have indicated that it is necessary to increase the number of cells containing DDPs at S/G2 cell division phases to increase HDR efficiency ^7^. Several traditional strategies have been applied to enrich DDP availability and introduction into cells, including particle bombardment ^8^, protoplast ^9^, geminiviral-based replication ^10^, and RNA transcription ^1^, but it remains a significant problem for woody plants. Although genes introduced by *Agrobacterium* are stable and the method is widely used to transduce genes into woody plant cells ^11,12^, there were few reports on enhancing the efficiency of transferring DDPs and, consequently, the recovery of DSBs by HDR ^13,14^.

Efforts to increase HDR efficiency through the introduction of HDR-promoting effectors have been attempted. Expression of Cas9 fusion proteins with HDR-enhancers *MRE11, CtIP*, or *Rad52* has been shown to enhance HDR efficiency in human cells while significantly decreasing NHEJ with at least a 2-fold increase HDR and a 6-fold increase in HDR/NHEJ ratio ^15^. However, little attention has been paid on the role of these proteins in plants to increase the HDR efficiency ^16^. In contrast, the inhibition of DNA ligase IV (LIG4), Ku 70, and Ku 80, which are outwardly involved only in NHEJ, has been shown to increase HDR efficiency up to 16-fold in *Arabidopsis* ^17^. The X-ray repair cross-complementing protein 4 (XRCC4) is another critical NHEJ factor that has not yet been extensively considered for its interfering effect on HDR efficiency in plants. XRCC4 is one cofactor of LIG4 to interact with KU 70 and KU 80 and ligate the DSB ^18,19^.

To date, there was no report combining HDR factor overexpression (i.e., CtIP and MRE11) and NHEJ factor suppression (i.e., XRCC4) to promote the HDR pathway in plants. In contrast to gene mutation via CRISPR-Cas9 systems, precise gene targeting and knock-in by homologous recombination are more challenging but necessary as a versatile tool for research and breeding in crops and woody plants. Therefore, our objective was to examine the effects of HDR cofactor overexpression (*CtIP* and *MRE11* ^15^) and simultaneous disruption of NHEJ promoter *XRCC4* ^20^ on knock-in efficiency using the *MKK2* gene as a case study. Mitogen-activated protein kinases (MAPKs or MPKs) like *MKK2* are involved in several key pathways responding to stresses, including disease, drought, cold, heat shock, osmotic, and salt stresses^21,22^.

Regarding highly efficient CRISPR-mediated homozygous mutations in poplars^23^, the *MKK2* gene was targeted to integrate the Zeocin resistance gene *BleoR* onto its 3′ end in the poplar genome (Figure 1a). The *MKK2* gene was targeted to integrate the Zeocin resistance gene *BleoR* onto its 3′ end in the poplar genome (Figure 1a). The guide RNA (gRNA) was designed near the 3′ UTR with the highest activity score and no off-target effects on CDS to avoid effects on *MKK2* expression and function and less chance of off-target CRISPR activity ^24,25^ (Figure 1a and Supplementary Table 1). According to Song et al. ^26^, lengths of homologous arms were optimized for use in homologous recombination and *BleoR* integration to 400 bp upstream and downstream of the PAM site designated as the 5′ and 3′ homology arms, respectively (Figure 1b). The DDP cassette was ligated into the pRGEB31 vector containing the Cas9 expression cassette to construct the pDDP vector (Supplementary Figure 1). Multiple fusion vectors were also constructed to manipulate HDR and NEHJ cofactors, including the Cas9 expression cassette, *CtIP* overexpression cassette, *MRE11* overexpression cassette, and XRCC4 mutative cassette (Supplementary Figure 2a-d). We used the pathogenic suspension of *Agrobacterium tumefaciens* with an OD_600_ = 2.5 (∼2×10^9^ cell ml^-1^) and the ratio of 4:1 pDDP/pgRNA to provide an excess of DDP fragments during S/G2 cell division ^15^ and to avoid off-target editing caused by the extra accumulation of pgRNA ^27^ (Figure 1c). Actively growing buds on Zeocin-containing selection medium were assumed to be positive transformants and were selected and rooted. The recovered events were then used for further analyses (Figure 1c).

**Figure 1:**
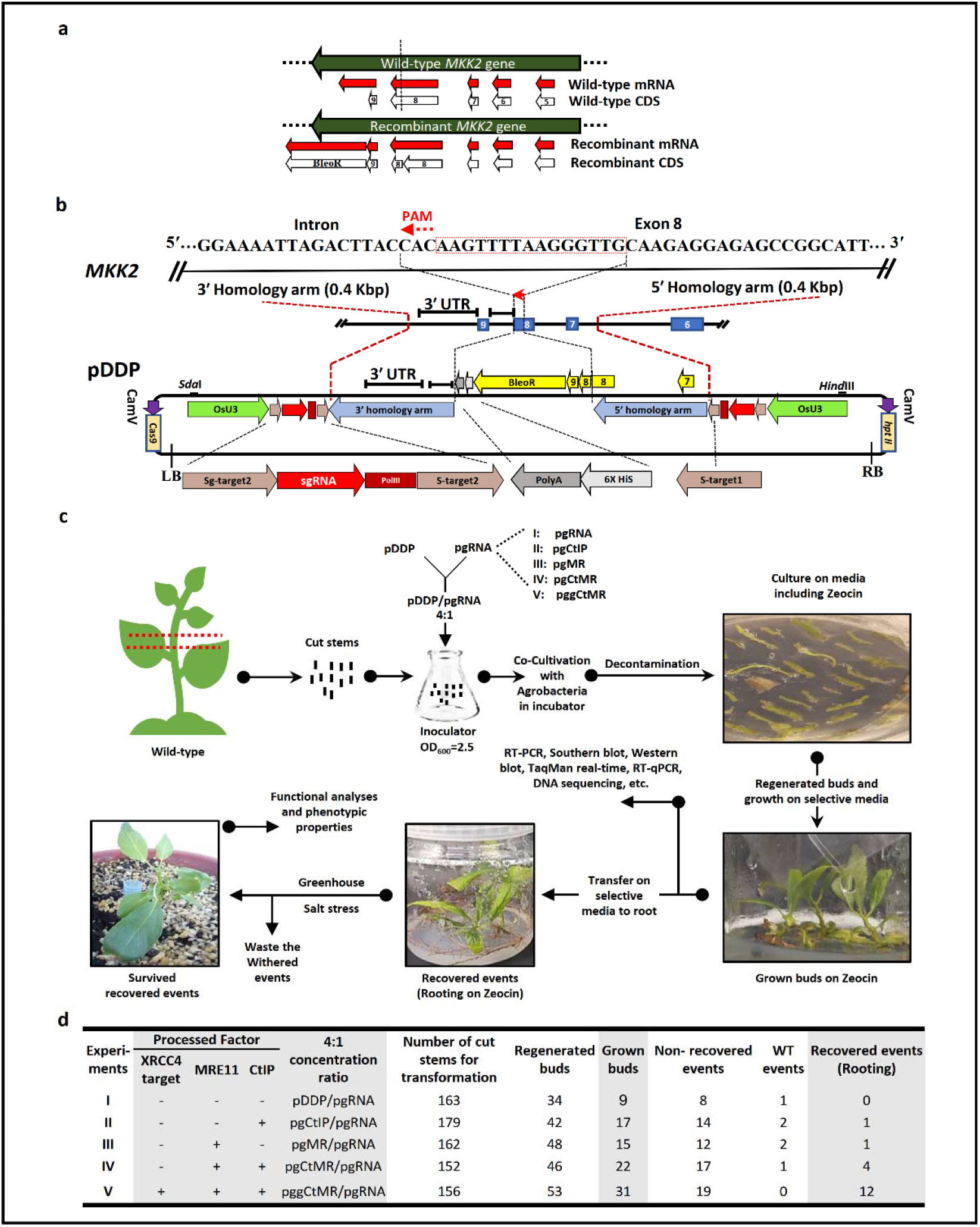
Exogenous *BleoR* CDS is integrated into the poplar genome. **a**, The purpose of this study to generate Recombinant mRNA, including *MKK2* and *BleoR*. Dash line reveals the target site. **b**, Protospacer Adjacent Motif (PAM) was detected at exon 8 to lead Cas9. 400 bp sequences from the up-and downstream of the CRISPR target were picked for HDR in this study. The 5′ homology arm included part sequences of the intron between exon 6 and -7, exon 7, intron sequences between exon 7 and -8, and a part of exon 8. The 3′ homology arm included intron sequences between exon 8 and -9 and 3′ UTR of the *MKK2* locus up to 400 bp. Designed DDP included remained sequences of exon 8, exon 9, *BleoR* CDS, 6xHis, and PolyA sequences flanked by the 3′- and 5′ homology arms. In addition, two special targets (S-target1 and -2) (No on- and -off-targets through the whole poplar) have been designed to attach besides DDP. The DDP was then ligated into the pRGEB31 vector to form pDDP. c, pDDP, and pgRNA were mixed 4:1 and introduced to the Agrobacterium tumefaciens to form inoculator suspension and condensed up to OD_600_=2.5. The putatively edited events were regenerated on Zeocin and were allowed to bud. The grown buds were then transferred on selective rooting media and allowed to be recovered. Recovered events were then planted on soil following by salt stress. **d**, The overview of designed experiments including (I) No HDR factors, (II) overexpressed CtIP, (III) overexpressed MRE11, (IV) overexpressed CtIP+MRE11, and (V) overexpressed CtIP+MRE11 with XRCC4 deficiency.

To examine the effect of increasing HDR effector expression and disruption of NHEJ component on knock-in efficiency, the pDDP cassette was co-transformed with one of five additional constructs in a 4:1 concentration ratio: pgRNA (Experiment I; ExI), pgCtIP overexpressing HDR effector CtIP (Experiment II; ExII), pgMR overexpressing HDR effector MRE11 (Experiment III; ExIII), pgCtMR overexpressing both *CtIP and MRE11* (Experiment IV; ExIV), and pggCtMR overexpressing both *CtIP and MRE11* along with CRISPR-based disruption of NHEJ effector XRCC4 (Experiment V; ExV) (Figure 1d; Supplementary Figure 2). While ExI resulted in no successfully recovered lines being generated from nine grown buds, ExII and ExIII resulted in one recovered event from 17 and 15 grown buds, respectively (Figure 1d). Combining *CtIP and MRE11* overexpression, ExIV resulted in four recovered events from 22 grown buds (Figure 1d). Seeing the positive influence of HDR effectors overexpression but still being unable to overcome the influence of NHEJ, pDDP was co-transformed with pggCtMR in ExV, resulting in a significant increase in recovered events to twelve from 31 grown buds (Figure 1d), which suggested that simultaneous overexpression of *CtIP and MRE11* and disruption of *XRCC4* resulted in a 3.4-fold increase in knock-in efficiency of the *BleoR* gene indicated by bud recovery compared to ExI.

In light of these results, Western blotting, RT-PCR, and Southern blotting were used to verify that HDR occurred in recovered transformants and to confirm the proper integration of *BleoR* to the 3′ end of the *MKK2* gene. A 6X His tag was fused with the *BleoR* C-terminal (Figure 1b) followed by Poly-A tail to show the integration of *BleoR* into target genomes using Western blotting. While screening the transformants grown on Zeocin using Western blotting, no edited events were detected in ExI, but one event showed a band of 54 KDa in ExII (Figure 2a; Uncropped documents in Supplementary Figure 16), which represents the successful integration of BleoR (∼13.7 KDa) fused with MKK2 (∼40.5 KDa) (Figure 2b). In screening events in ExIII, only one with a band of 13.7 KDa (Figure 2a) was identified, suggesting that the *BleoR* CDS were integrated but did not successfully form a fusion protein with MKK2 possibly due to mutation or knock out *MKK2* exons 7, 8, or 9 (Figure 2b). Screening events in ExIV showed three events with about 54 KDa bands and one event with about 14 KDa (Figure 2a). However, simultaneous HDR effector overexpression and NHEJ suppression in ExV resulted in ten events with about 54 KDa bands and two events with about 14 kDa bands (Figure 2a).

**Figure 2:**
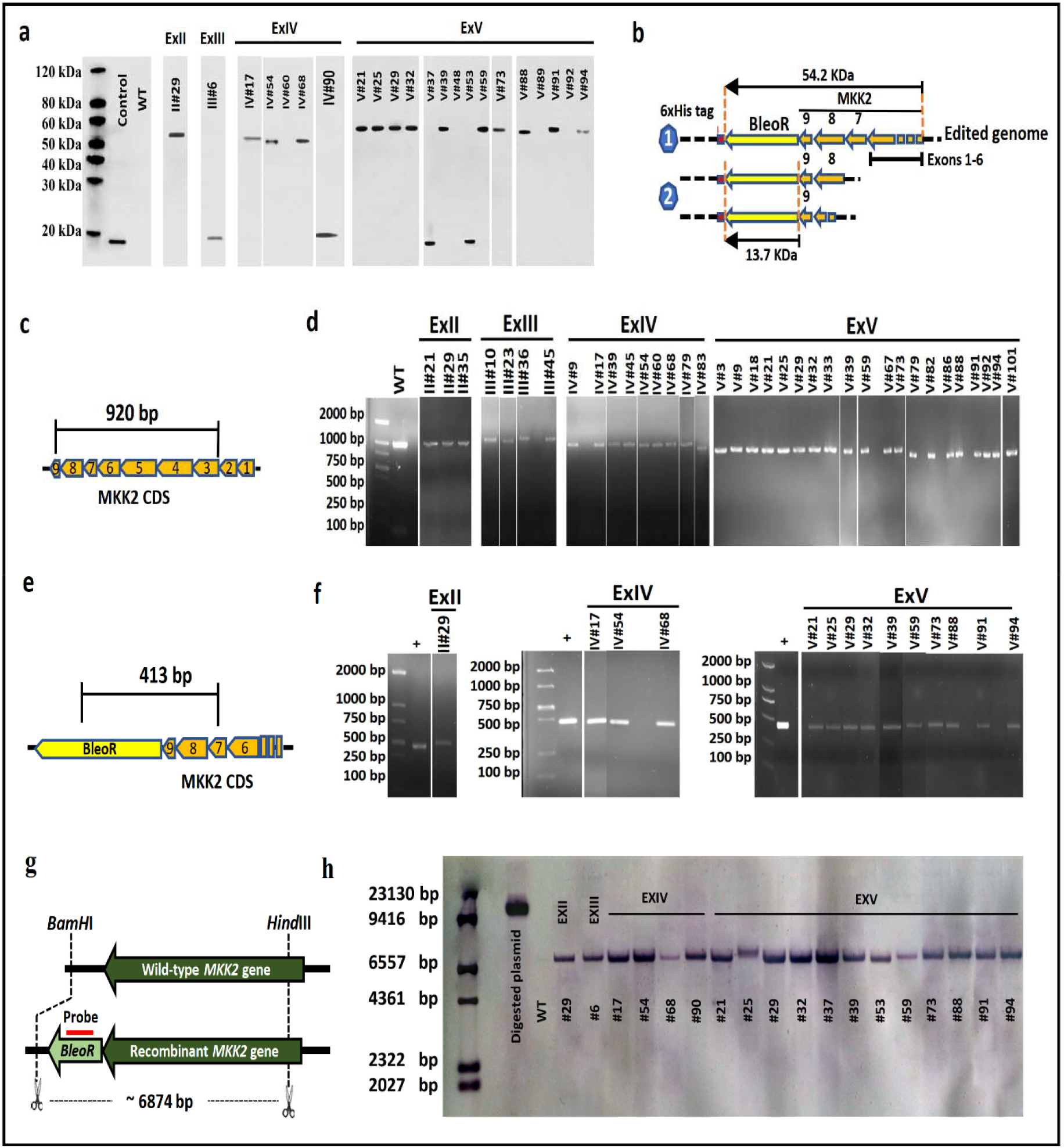
Western blotting, RT-PCR, and Southern blotting exhibited validation of *BleoR* integration in recovered events. **a**, Recovered events exhibited *BleoR* expression by Western blotting. **b**, Schematic of fusion 6xHis tag with edited poplar genome triggered by different constructions. Shape 1 reveals successful fusion of *BleoR* and *MKK2* with about 54.2 kDa. Shape 2 reveals only *BleoR* translation and unsuccessful fusion of *MKK2* and *BleoR* proteins because of *MKK2* disruption with about 13.7 kDa. **c**, Schematic of proper HDR happening caused to attach exon 8 and 9 resulting in *MKK2* transcription in the edited genome. **d**, RT-PCR exhibited the actual engineered events resulting in amplifying 920 bp of *MKK2* CDS. The β-actin was used as the control in all RT-PCR assays to normalize; WT was used as the positive control. **e**, Schematic of proper integration in edited genom caused to connect the *BleoR* to the C-terminal of *MKK2*. **f**, RT-PCR detected the recovered events with proper integration of *BleoR* and *MKK2* with amplifying 413 bp of their transcription. The β-actin was used in all RT-PCR assays to normalize; pDD plasmid was used as the positive control. WT was used as the negative control. **g**, Schematic of probing *BleoR* in edited events and WT as the control using Southern blotting. **h**, Southern blot proved the integrated *BleoR* into poplar genome resulted in Western blotting. Linearized pDDP plasmid was used as the positive control; WT was used as the negative control.

RT-PCR assays were performed to evaluate the accuracy of *MKK2* editing and to ensure the proper orientation of the insertion of *BleoR* at the 3′ end of *MKK2* exon 8 in the recovered transformants (Figure 1d). The first RT-PCR experiment was designed to determine whether HDR was successful and proper *MKK2* gene transcription occurred in the transformants (Figure 2c and d). A 920 bp region spanning exons 3 – 9 was amplified, and, as expected, there were no bands observed in ExI events. Three ExII events and four ExIII events showed WT with 920 bp bands, respectively. Also, nine ExIV events showed WT bands. Concurrent with increased transformation efficiency, more WT amplicons were observed in ExV than the other experiments with twenty events exhibiting WT bands (Figure 2d). The proper orientation of inserted *BleoR* at the 3′ end of *MKK2* was evaluated by the second RT-PCR experiment in which pDDP was designed as the positive control (Figure 2e and f). There was no 413 bp amplicon spanning exon 7 of *MKK2* to a point within *BleoR* (indicative of correct *BleoR* orientation) shown in ExI events. ExII events showed only one 413 bp amplification, but none of the ExIII events showed the desired band. Three positive events were observed in ExIV. However, in ExV, ten events showed the desired bands (Figure2f).

Relying on predicting the lack of CRISPR off-target and verifying the absence of *BleoR* integration elsewhere in the poplar genome (Supplementary Table 1), Sothern blotting was performed using a *BleoR*-specific probe for all recovered events (Figure 2g and h). In addition to validating the *BleoR* knock-in in the recovered events, a single band for each event also proved the precise knock-in in desired positions with one or more copy numbers, indicating the lack of *BleoR* in the off-target sites.

Overall, these results further support the concept that the overexpression of *CtIP* and *MER11* and the disruption of *XRCC4* resulted in increased *BleoR* knock-in efficiency when comparing ExV results to the others. To further validate HDR in the events, TaqMan real-time PCR was utilized with two probes, FAM1 and FAM2, for the 5′ and 3′-ends of the *BleoR* CDS (Figure 3a). Transformants exhibiting both FAM1 and FAM2 fluorescent signals were assumed to be fully edited, those exhibiting only FAM1 or FAM2 were considered to be partially edited, and those with no FAM1 or FAM2 signals were assumed to be either mutants or WT (Figure 3b). In ExI, the averages of fluorescent signal numbers of FAM1 and FAM2 ΔΔCt were between 4 to 8 (Supplementary Figure 3a), and most events exhibited as the mutant or WT, with a few partial FAM1 or FAM2 fluorescence (Figure 3c). The ExII and ExIII events showed enhanced fully and partially edited FAM signal numbers (Figures 3d, and e; Supplementary Figures 3b and c). In ExII, there were four fully edited events, four FAM1 partial edited events, and four FAM2 partial edited events (Supplementary Figure 9a). In ExIII, three fully edited events, five FAM1 partial edited events, and three FAM2 partial edited events were observed (Supplementary Figure 9b). In ExIV, the signal density of edited events increased significantly (Figure 3f). The mean fluorescent FAM1 and FAM2 signal numbers showed an increase of about 20 and 15, respectively (Supplementary Figure 3d). In total, nine fully edited events, seven FAM1 partial edited events, and four FAM2 partial edited events (Supplementary Figure 9c) were detected. Finally, the FAM signal densities of fully edited transformants in ExV were increased (Figure 3g) with means about 21 and 19, respectively (Supplementary Figure 3e), and fifteen fully edited events were discovered (Supplementary Figures 9d and 10). Furthermore, total FAM fluorescent signals (FAM1, FAM2, and FAM1&2) indicated an increasing trend of HDR occurrence through these experiments (ExI to ExV) (Figure 3h). These results were further validated by Sanger sequencing of the amplified regions followed by multisequence alignment (Supplementary Figures 4 – 8).

**Figure 3:**
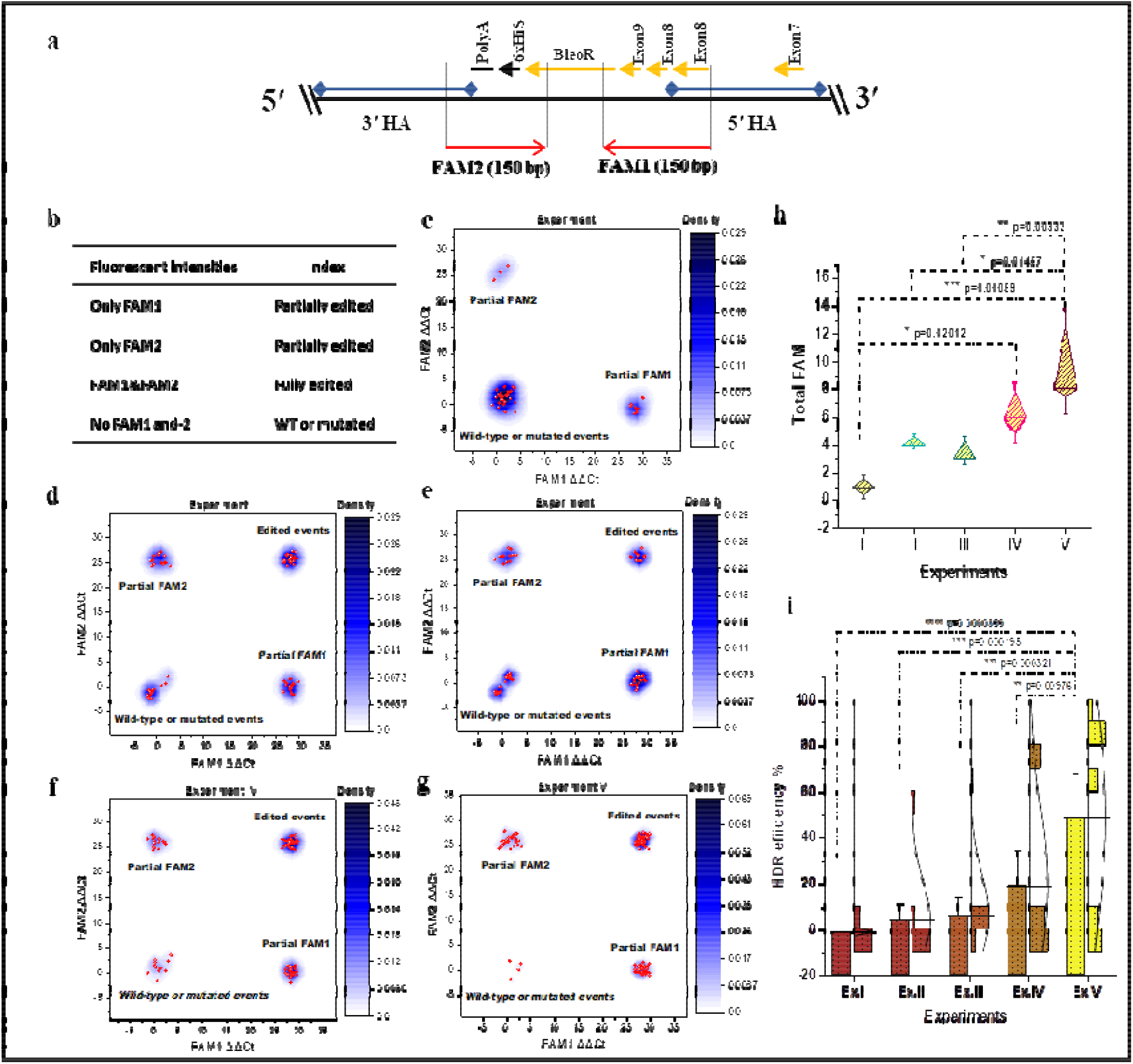
TaqMan real-time PCR and real-time PCR assays to validate and evaluate HDR occurrence and efficiency. **a**, Designing TaqMan real-time PCR assay to detect and evaluate HDR efficiency, included FAM1 and FAM2 DNA binding probes. **b**, Strategy to classify edited events. **c**, Experiment I revealed no edited events. **d**, The density plot of FAM1 and -2 intensities resulted from experiment II revealed an expansion in edited events against partial, mutant, and wild-types. **e**, The density plot of FAM1 and -2 signals resulted from experiment III revealed an increased intensity of partial FAM1 events. **f**, Experiment IV revealed a remarkable increase of edited events signals in confronting with three earlier experiments. **g**, The Density plot of experiment V revealed a significant increase of FAM1 and -2 intensities in edited events compared to the earlier experiments and a significant decrease in intensities in WT and mutated events. All samples were analyzed in quadruplicate. **h**, Diamond box, and whisker plot compared FAM signals (Partial FAM1 and -2 and FAM1&-2) detected in all experiments and showed more signals remarkably measured in ExV than ExI, II, and-III; Error bars represent SE; Asterisks represent p-value as *≤0.05, **≤0.01, and ***≤0.001. **i**, The exogenous *BleoR* expressions in the poplar genome resulting from HDR were evaluated (ΔΔCt) and compared to exhibit HDR efficiency %. The overlap data are shown as bin bars, and the standard distribution curves are added to confirm bars. HDR efficiency plot resulted from ExV events revealed significantly more *BleoR* expression than the other experiment events. Also, ExIV meaningfully revealed more HDR happening than ExII and -III.; Error bars represent SE; Asterisks represent p-value as **≤0.01, ***≤0.001, and ****≤0.0001; Triplicate technical repeats were considered for each sample.

Previously, HDR efficiency was improved ten-fold in tomatoes, and 35S promoter was integrated upstream of the *ANT1* gene ^2^. While some reports on *Populus* genome editing are limited only to knock-out genes and mutations that happened by Cas9 and Cas12a ^13,23,28^, the present study reports improving HDR efficiency for gene knock-ins in poplar. Estimating HDR efficiency was achieved by measuring the exogenous *BleoR* expression using real-time PCR and calculating the ratio of 100-fold of *BleoR* expression ΔΔCt mean:31, determining the percentage of *BleoR* expression ΔΔCt mean (HDR%) for each event (Supplementary Table 2a). The resulted HDR% from all events were then analyzed by One way-ANOVA, analyzing HDR% efficiency (Figure 3i; Supplementary Table 2b). Regarding improving the expression of HDR factors leading to increases in HDR efficiency ^29^, previous studies have shown that HDR efficiency could be improved up to 19-fold by inhibiting NHEJ factors in mammalian systems ^30^. While Tran et al.^15^ showed at least a 2-fold increase in HDR resulted from CtIP-Cas9 and MRE11-Cas9 fusions in human cells, here, the mean expression of *BleoR* increased from -1.23% in ExI to 4.40% (∼4-fold) in ExII and 6.11% (∼5-fold) in ExIII. When fusing both *CtIP and MRE11* with Cas9, the mean expression in ExIV raised to 19.07% (∼16-fold compared to ExI) in this study. Finally, while Qi et al. ^17^ reported 5∼16-fold or 3∼4-fold enhancement in HDR in *Arabidopsis* by knocking-out the *Ku*70 or *Lig*4, this study demonstrated that *XRCC4* deficiency in conjunction with *CtIP and MRE11* overexpression (ExV) raised the HDR by 48.90% (∼40-fold than ExI). Overall, XRCC4 deficiency, together with the overexpression of *CtIP* and *MRE11*, was the most efficient system for HDR-based integration resulting in more HDR occurrence than HDR expression effectors *CtIP and MRE11* alone or together (Figure 3i).

To evaluate the effect of *CtIP and MRE11* overexpression and simultaneous *XRCC4* suppression on CRISPR-induced polymorphisms, we analyzed the variant genotypes and protein effects within the 5′ and 3′ homologous arms and also the knocked-in fragments from the grown buds through all experiments (Supplementary Table 3). The increased HDR efficiency observed over the subsequent experiments from ExI to ExV suggested a shift from higher Insertion/Deletion (InDel) polymorphisms occurrence in the 5′ region of knocked-in fragments (*BleoR*) to the 3′ region (6xHis and PolyA) (Figure 4a). The result of this shift was less functional disruption of *BleoR* resulting from InDels localized more 3′ rather than 5′, resulting in more *BleoR* expression numbers in ExV events than the other experiments. Furthermore, the InDels number mean comparisons revealed that the promoted HDR by *XRCC4* deficiency caused a 7-fold decrease in InDels in ExV events compared to considerably ExI events and also the other experiments (Figure 4b). We then calculated the total number of polymorphisms throughout the homologous arms in each experiment. ExIV and ExV events resulted in the least numbers of polymorphisms, significantly 2.3-fold and 3.8-fold less than ExI events, respectively (Figure 4c). More polymorphisms also occurred within ExII and -III than ExIV and -V (Figure 4c). Examining all polymorphism classes observed in the data (Supplementary Table 4), the highest frequency of polymorphisms occurred in ExI events and the least within ExV events, with the majority being InDels (Figure 4d). Moreover, the whisker plot of total polymorphisms presented the maximum of polymorphisms through ExI events and the minimum of those in ExV events (Figure 4d).

**Figure 4:**
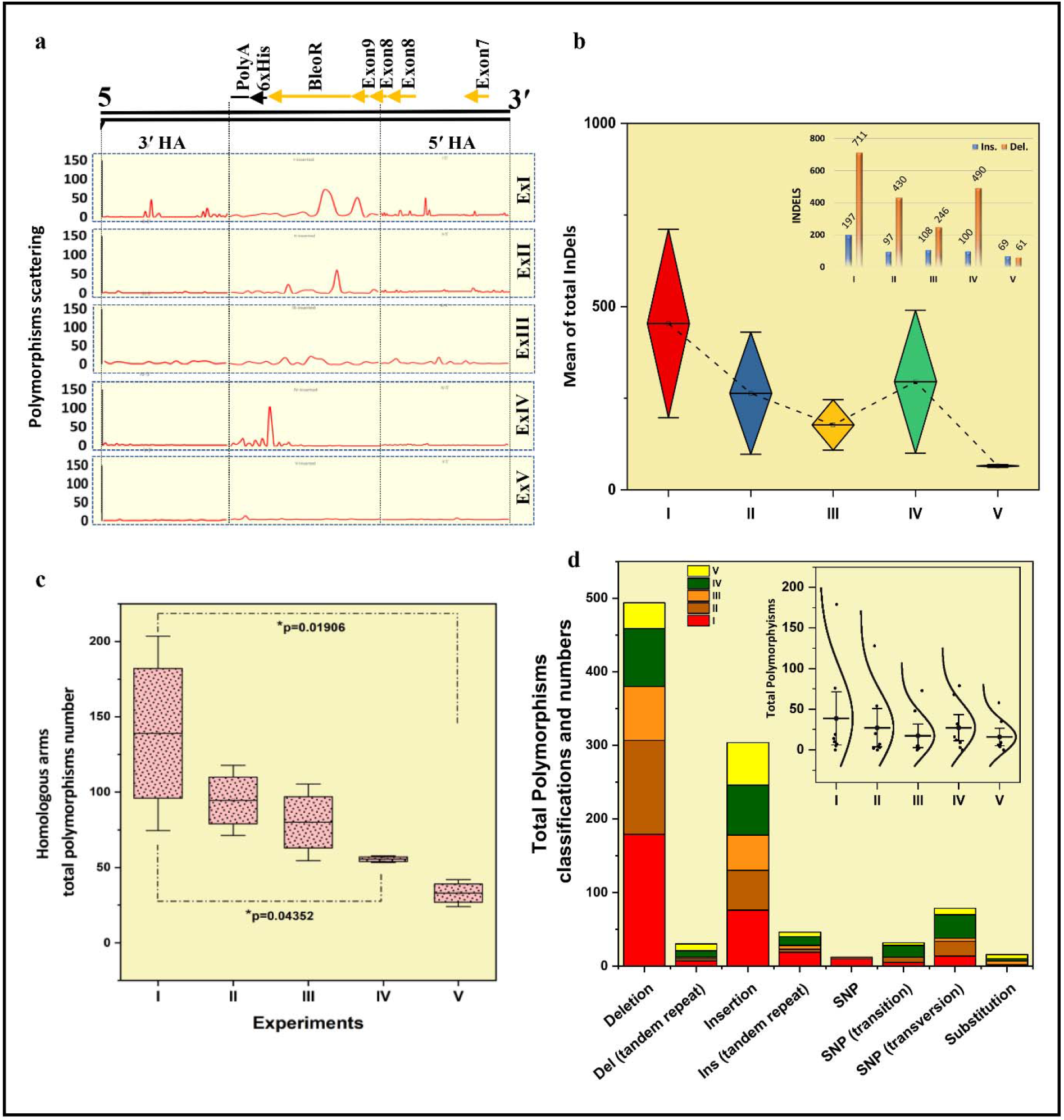
HDR promotion by *XRCC4* deficiency besides *CtIP and MRE11* overexpression caused a decrease in InDels considerably via ExV events compared to the other experiments. **a**, Analyses of distributed InDels happened on 5′ and 3′ homology arms (HA) and knocked in fragments throughout experiment events. **b**, Diamond box, and whiskers for the mean comparisons of total InDels through experiment events. The exact numbers of InDels (Excluding SNPs and substitutions) are presented via the punching column bars on the top-right corner. **c**, Identification of the happened polymorphisms through homology arms among all the experiments. Box and Whisker plot revealed that most polymorphisms happened in homology arms via ExI, and it was significantly more than those in ExV and –IV; Asterisks represent p-value as *≤0.05; Error bars represent SE. **d**, Stacked column plot of total polymorphisms classification and numbers in DDP integration among all the experiments. Insertions and deletions occurred much more than the other types. SNP and substitutions occurred less than the other types. Whisker and standard curves exposed the least total polymorphisms that happened through ExV than the other experiments.

It has been shown that *MAPKs* genes direct cellular responses against abiotic stresses such as salinity^22,31^., The *MKK2* functional analysis and comparing them with WT poplars revealed a regular expression (∼95%-100%) before stress in successfully transformed events from ExII, -IV, and -V, and stable overexpression induced by salt stress application (168%-173%). In addition, no loss of salt stress tolerance was observed in the successfully transformed events confirming that *MKK2* remained functional and no deleterious mutations occurred following HDR across exons 7, 8, and 9 (Supplementary Figure 17). Concerning unidentified bands in Figures 2d and f, the events II#6, IV#90, V#37, and V#53 could not resist salinity due to *MKK2* mutations and withered. In addition, Sun et al. ^32^ reported that the *MKK2* family genes play vital roles in maize development. Thus, a lack of *MKK2* expression may result in reduced plant stem length and diameter. Our results showed no significant differences between surviving recovered events before and after salt stress means and WT poplars in stem lengths and diameters, which validated the precise editing of the *MKK2* locus by efficient HDR within exons 7, 8, and 9 (Supplementary Figure 17 and Table 5).

In summary, this study demonstrated that NHEJ factor deficiency, together with HDR factor overexpression, results in enhanced HDR efficiency up to a 40-fold increase and dramatically expands our capacity for trait improvement in poplar. We also proved that HDR promotion by the overexpressing HDR factors (ExII, ExIII, and ExIV) drives the resected DNA repair through homologous arms with 1.7-, 2.5-, and 1.5-fold respectively fewer InDels than ExI, but not as much as the HDR factors overexpressing together with NHEJ factors deficiency (ExV) with 7-fold fewer. This breakthrough technology will prove helpful in biotechnological research, forest conservation of tree species, crop improvement and development, and mammalians.

## Acknowledgments

This project was funded by the National Key Program on Transgenic Research (2018ZX08020002), the National Natural Science Foundation of China (No. 31971682, 31570650 and 2045210646).

## Author contribution

A.M. conceived, planned, and coordinated the project, performed data analysis, and wrote the draft, and finalized the manuscript. H.W. carried out the experiments and contributed to data analysis and curation. X.Z., J.C.F., and Z.H.C. validated and contributed to data analysis and curation, revised and finalized the manuscript. Z.M., W.S., J.Z., D.L. reviewed and edited the manuscript. B.G. validated and contributed to data curation, review, and editing. R.K.V., L.Y., and Q.Z. planned, coordinated, contributed to data curation, and revised and finalized the manuscript. A.M., H.W., X.Z., and J.C.F. contributed equally as the first author.

## Supplementary Information

**Supplementary Figure 1:** Schematic DDP and pDDP. pDDP included DDP ligated into pRGEB31 by restriction enzyme cloning method.

**Supplementary Figure 2:** Schematics of constructed cassettes and plasmids. a, pgRNA included the *MKK2* target seed and Cas9. b, pgCtIP plasmid including CtIP cassette. c, pgMR plasmid, including MR cassette. d, pgCtMR plasmid including CtMR cassette. e, pggCtMR plasmid, including XRCC4 cassette.

**Supplementary Figure 3:** Box-and-whisker (Min-Max) plots of one-dimensional FAM delta-delta Ct signals in designed experiments. All signals were calculated as quadruplicates.

**Supplementary Figure 4:** Alignment of events involved in experiment I.

**Supplementary Figure 5:** Alignment of events involved in experiment II.

**Supplementary Figure 6:** Alignment of events involved in experiment III.

**Supplementary Figure 7:** Alignment of events involved in experiment IV.

**Supplementary Figure 8:** Alignment of events involved in experiment V.

**Supplementary Figure 9:** Schematics of sequence analyzing of triggered events from different experiments. **a**, Sequence analysis of triggered events included in EXII reveals one recovered event. **b**, Sequence analysis of triggered events included in EXIII reveals one recovered event. **c**, Sequence analysis of triggered events included in EXIV reveals four recovered events. **d**, Sequence analysis of triggered events included in EXV reveals 12 recovered events.

**Supplementary Figure 10:** Chromatogram alignments of events included in experiment V.

**Supplementary Figure 11:** Schematic constructions of DDP and pDDP fragments.

**Supplementary Figure 12:** Constructions of CtIP, MRE11, CtIP+MRE11, and XRCC4 cassettes with their primers and oligos.

**Supplementary Figure 13:** Constructions of designed plasmids pgCtIP and pgMR with their primers.

**Supplementary Figure 14:** Constructions of designed plasmids pgCtMR and pggCtMR with their primers.

**Supplementary Figure 15:** Schematic of TaqMan real-time PCR FAM and VIC target assays in this study. Yellow rectangles exhibited CDS.

**Supplementary Figure 16:** Uncropped agarose gel images used in figure 2a, d, f, and h. **a**, Western blotting. **b** and **c**, RT-PCR. **d**, Southern blotting; Red dashed boxes indicate the cropped sections handled in figures.

**Supplementary Figure 17:** Radar diagrams of *MKK2* expressions, stem lengths, and - diameters from WT and survived recovered events after NaCl treatment. No significant differences in *MKK2* expressions and phenotypic changes before and after salt stress between WT and survived recovered events confirmed the exact genetic engineering induced by efficient HDR to keep *MKK2* exons arrangement.

**Supplementary Table 1:** CRISPR sites located on 3′ region of *MKK2*. The yellow highlight reveals the selected CRISPR target in this study.

**Supplementary Table 2:** The HDR efficiency calculated in this study. **a**, The percentage of *BleoR* expression ΔΔCt mean (HDR%) achieved from all grown buds on Zeocin. Each event was investigated with three technical repeats; To balance the variation of each experiment event number, we considered zero for all the *BleoR* expressions up to 31 (Equals to ExV events). **b**, One way-ANOVA descriptive statistics computed from all events HDR% has been used for analyzing each experiments HDR% efficiency in Figure 3i.

**Supplementary Table 3:** Polymorphisms detected in homologous arms happened by HDR through all experiments grown buds.

**Supplementary Table 4:** Total variant genotyping happened by CRISPR-induced polymorphisms through all experiments grown buds.

**Supplementary Table 5:** The raw data of RT-qPCR of *MKK2* expressions and phenotypic analyzes in WT and survived recovered events before and after salt stress. **a**, *MKK2* expressions (%) Rt-qPCR; All events were analyzed in triplicates. **b** and **c**, Stem lengths and diameters before and after stress. **d, e**, and **f**, Duncan test tables for *MKK2* expression, stem length, and stem diameters, respectively. The mean of before and after treatments from each event was compared with WT as the control.

**Supplementary Table 6:** Oligos and primers used in this study.

## Methods

### Design of Experiments and Construct Transformation

#### Design of Experiments

This study was based on promoting HDR efficiency in poplar using designed plasmid harboring gRNA (pgRNA) (Supplementary Figure 2a), plasmids (pgCtIP, pgMR, pggCtMR) harboring gRNA, and HDR factors CtIP and MRE11 (Supplementary Figures 2b, c, and d), and plasmid (pggCtMR) harboring gRNA to target NHEJ factor XRCC4 (Supplementary Figure 2e). All plasmids included gRNA to target *MKK2* locus on *Populus trichocarpa*. The designed plasmids were then classified into five experiments: ExI including pgRNA, ExII including pgCtIP, ExIII including pgMR, ExIV including pgCtMR, and ExV including pggCtMR.

#### Construction of DDP and pDDP

To produce DDP (Supplementary Figure 1), five fragments were designed, constructed, and ligated (Supplementary Figure 11a). To construct fragment one, the OsU3 promoter and gRNA scaffold were isolated from pRGEB31 (Supplementary Table 6, OS1-F and -R) flanked by *HindIII* and *BamHI* sites. To increase the amount of DDP in the cell nucleus and improve HDR efficiency, the cleavage property of Cas9 was harnessed by designing two special gRNA targets 1 and -2 (No on- and -off-targets through the whole poplar genome and only detect special targets besides of DDP) in addition to the DDP ^33^ (Supplementary Figure 1). Special gRNA oligos (Sgo1-F and -R) (Supplementary Figure 11a; Supplementary Table 6, special gRNA oligo1-F and - R) were then designed as previously described ^34^ to form special gRNA target1 (Sgt1), which was then ligated into the fragment one. To construct fragment two, we decided to isolate 400 bp nucleotides of upstream of the target from the poplar genome (5′ homology arm) (Supplementary Table 6, 5′ Ho-F-1 and -R-1). Afterward, regular PCR was carried out using primers with the extensions of *BamHI*-special target1 (St1) and 39 bp from complemented 5′ of fragment 3 (Supplementary Table 6, 5′ Ho-F-2 and -R-2) (Supplementary Figure 11a). To construct fragment three, we isolated the *BleoR* CDS (Zeocin resistance gene) from the PCR®-XL-Topo® vector (Supplementary Table 6, *BleoR*-1092F and -2276R). Then, overlap-PCR was performed (Supplementary Table 6, BP1,2,3-F and -R) using the isolated *BleoR* CDS as a template to include sequences rather than remaining nucleotides from exon 8 (Leu-Ala-Thr-Leu-Lys-Thr-Cys) and exon 9 (Val-Leu-Val-Lys-Met) for adding to the 5′ *BleoR* CDS region and also 18 bp 6xHis tag and 30 bp Poly A tail for adding to the 3′ *BleoR* CDS area (Supplementary Figures 1 and 11a). We decided to isolate 400 bp nucleotides of downstream of the target from the poplar genome (3′ homology arm) (Supplementary Table 6, 3′ Ho-F-1 and -R-1) to assemble fragment four. Then, PCR was performed to extend 3′ homology arm with 30 bp Poly-T and NcoI-special target2 (St2) sequences (Supplementary Table 6, Ho-F-2 and -R-2) (Supplementary Figure 11a). Finally, standard PCR was used to isolate the OsU3 promoter and gRNA scaffold from pRGEB31 (Supplementary Table 6, Os2-F and Os2-R). Moreover, special gRNA oligos were designed (Sgo2-F and -R) (Supplementary Figure 11a; Supplementary Table 6, special gRNA oligo2-F and -R) again as previously described ^34^ to form special gRNA target2 (Sgt2) and ligated it into fragment five.

To construct the final pDDP construct, we ligated fragments two and three using PCR (Supplementary Figure 11b). For this, we designed a 39 bp overhang on fragment two that was complementary to the end of fragment three to form preliminary DDP (Supplementary Figure 11b). For this reaction, PCR was prepared with 500 ng of each component. Initially, all components were used in the PCR reaction except primers, and then the fragments were denatured at 95°C for 5 minutes, followed by two annealing and extension cycles. Next, the PCR products were allowed to anneal at 68°C to avoid nonspecific hybridization amongst the long PCR products for 30 seconds, followed by extension for one minute at 74°C resulting in a double-stranded template. The primers were then added for the distal ends of fragments two and three, and PCR proceeded normally. The PCR products were then purified and ligated into the pEASY vector for sequencing and confirmation. The preliminary DDP product was then ligated into fragment four as previously described and formed secondary DDP products (Supplementary Figure 11b). After sequencing and confirmation, restriction cloning was used to ligate secondary DDP products to fragments one and four (Supplementary Figure 11b). Briefly, we incubated a reaction including 50 ng of each digested fragments, 10x T4 DNA ligase buffer 0.5 µl, T4 DNA ligase (NEB) 1 µl, and H_2_O to 5 µl at 25°C for 4 hours and transferred into E. Coli DH5α competent cells for sequencing and confirmation. Subsequently, the restriction cloning technique was used to merge the DDP product and pRGEB31 vector to form the pDDP vector (Supplementary Figure 11b).

#### Synthesis of pgCtIP and pgMR

To design a fused CtIP and Cas9 cassette, the CaMV35S promoter, 3xFLAG, and Cas9 CDS were isolated from pRGEB31 (Supplementary Figure 12a) using designed primers (Supplementary Table 6). CtIP CDS were then obtained using RT-PCR from the *Populus trichocarpa* genome (Supplementary Figure 12a; Supplementary Table 6, CtIP-F and -R). Next, the 3′UTR and PolyA fragments were isolated from the pCAG-T3-hCAS-pA plasmid (Supplementary Figure 12a; Supplementary Table 6, PolyA-F and -R). To complete pgCtIP, CaMV35S and 3xFLAG fragments were ligated using restriction cloning to form backbone 1 (Supplementary Figure 13a). Next, the isolated Cas9 and the obtained CtIP CDS were also ligated, applying restriction cloning to form backbone 2 (Supplementary Figure 13a). Backbones 1 and 2 were then ligated using *HindIII* restriction cloning to form backbone 3 (Supplementary Figure 13a). In the next step, the resulted backbone 3 was ligated to the assembled 3′UTR-PolyA using StuI restriction cloning to form the CtIP cassette (Supplementary Figures 13a and 12a). SdaI and PmeI restriction enzymes were then used to restrict the cloning of the CtIP cassette and pRGEB31 and assemble the pgCtIP plasmid (Supplementary Figures 13a and 12a).

To construct a fusion of MRE11 and Cas9, the CaMV35 promoter, 3xFLAG, Cas9, 3′UTR, and PolyA were isolated as previously described (Supplementary Figure 12b; Supplementary Table 6). The MRE11 CDS were obtained from *P. trichocarpa* total RNA, and RT-PCR was carried out as mentioned above (Supplementary Figure 12b; Supplementary Table 6, MRE-F and R). To complete pgMR, we ligated the isolated CaMV35S and 3xFLAG fragments concerning *XhoI* endonuclease to form backbone 1 (Supplementary Figure 13b). Backbone 2 was then constructed using the isolated Cas9 and 3′UTR-PolyA fragments (Supplementary Figure 13b). Backbone 1, backbone 2, and MRE11 CDS product were then merged using NotI and NdeI (NEB) restriction cloning to form the MR cassette (Supplementary Figures 13b and 12b). Then restriction cloning with SdaI and PmeI was used to construct the pgMR plasmid (Supplementary Figures 13b and 12b).

#### Synthesis of pgCtMR and pggCtMR

To construct the CtMR cassette, we prepared all the required fragments, as described above (Supplementary Figure 12c). Afterward, CaMV35S and 3xFLAG components were merged using *XhoI* restriction cloning to form backbone 1 (Supplementary Figure 14a). Backbone 1 and the already obtained MRE11 CDS product (Supplementary Table 6, MRE-F and -R) were then ligated using *NotI* restriction cloning to form backbone 2 (Supplementary Figure 14a). The isolated Cas9 and the obtained RT-PCR product CtIP CDS were ligated using *BamHI* restriction cloning to form backbone 3 (Supplementary Figure 14a). Backbone 3 and isolated 3′UTR-PolyA fragments were then used to form backbone 4 (Supplementary Figure 14a). Backbones 2 and 4 were then used to construct the CtMR cassette (Supplementary Figures 14a and 12c), followed by *SdaI* and *PmeI* restriction cloning to ligate the CtMR cassettes into pRGEB31 forming the pgCtMR plasmid (Supplementary Figures 14a and 12c). To target the XRCC4 gene and *MKK2* simultaneously, we designed one cassette, including both *XRCC4* [by adding one CRISPR site (Located on 5′ region of target CDS) to mutate XRCC4 (Non-off-target site on whole poplar genome; Activity score: 0.415; Specificity score: 100%) ^24,25^] and *MKK2* gRNAs. PCR was then used (Supplementary Table 6, XR-Cass1-F and -R) to isolate the OsU3 promoter and gRNA scaffold from the pRGEB31 vector, and *MKK2* designed oligos (Supplementary Table 6, *MKK2* Oligo-F and -R) were then used to ligate the *MKK2* target duplex (Supplementary Figure 12d). In addition, PCR was used (Supplementary Table 6; XR-Cass2-F and -R) to isolate the OsU3 promoter and gRNA scaffold again. In this process, we applied *XRCC4* designed oligos (Supplementary Table 6; XRCC4-Oligo1 and -2) to ligate the *XRCC4* target duplex (Supplementary figure 12d). The resulting fragments were then cloned using *K*asI restriction cloning to form *XRCC4*-Cassette (Backbone 1) (Supplementary Figures 14b and 12d). The *XRCC4*-Cassette was then cloned into pRGEB31 using *Hind*III and S*daI* restriction cloning to form backbone 2 (Supplementary Figure 14b). Finally, *S*daI and *P*meI restriction cloning was used to clone the CtMR cassette into backbone 2, forming the pggCtMR plasmid (Supplementary Figures 14b and 12d). Validation of construct assembly was performed using PCR, cloning into pEASY T3 vector, and DNA sequencing throughout construction.

### Transformation and targets detection

#### Plant transformation

For transformation, poplar (*P. trichocarpa*) seedlings were cultivated in a Phytotron at 23 ± 2°C under a 16/8 light/dark time ^11^. To generate transgenic lines, stems from four weeks old clones were dipped in an optimized *Agrobacterium tumefaciens* suspension (OD_600_: 2.5, 120 min, pH ∼5, Acetosyringone (As): 200 µM) ^12^ for 5 min with gentle shaking. The transformed stems were then transferred to a semi-solid woody plant medium (WPM) containing 0.05 mg/L Indole-3-butyric acid (IBA), 0.006 mg/L thidiazuron (TDZ), 200 µM As, and 0.5% (w/v) agar. Afterward, the stimulated stems were incubated in the dark at 23°C for two days. The assumed transformants were then co-cultivated in selection media enriched with 0.1 mg/L IBA, 0.006 mg/L TDZ, 100 mg/L cefotaxime, 8 mg/L hygromycin, 50 mg/L Zeocin, and 0.8% (w/v) agar. Two weeks later, buds were regenerated and then sub-cultured independently in media containing 0.1 mg/L IBA, 0.001 mg/L TDZ, 100 mg/L cefotaxime, 8 mg/L hygromycin, 50 mg/L Zeocin, and 0.8% (w/v) agar. After six weeks, buds with four to six small leaves were transferred to MS media containing 0.1 mg/L IBA, 200 mg/L cefotaxime, 70 mg/L Zeocin, and 0.8% (w/v) agar to root. Five independent transgenic lines were used for each experiment, and each line included about ten individuals.

#### Targets and protein detection

The *MKK2* gene from P. trichocarpa (POPTR_0018s05420g; Chromosome 18) was selected as a target for editing due to its vital role in transcriptional regulation against environmental stresses. Uniprot database (https://www.uniprot.org/) was used to download the *MKK2* protein sequence and then used the BLAST database of the National Center for Biotechnology Information (NCBI) (https://blast.ncbi.nlm.nih.gov/) to download full DNA sequences and CDS. To detect targets, Geneious Prime® 2020.1.1 was used to analyze the *MKK2* locus and detect targets relative to the whole genome of *P. trichocarpa* downloaded from NCBI (Supplementary Table 1) ^24,25^. Geneious Prime was also used to analyze the XRCC4 (POPTR_0010s08650g, Chromosome 10) gene for knocking out. The PAM motif target sequences were associated with exon 8 from *MKK2* and exon 1 from XRCC4. Furthermore, to evaluate the effect of HDR proteins and also proper function of the edited *MKK2* gene in transformants, relevant protein sequences CtIP (POPTR_001G269700v3), MRE11 (POPTR_0001s41800g), BRCA1 (POPTR_0005s26150g), Rad50 (POPTR_0001s32760g), Rad51 (POPTR_0014s06360g), Lig4 (POPTR_0018s13870g) were downloaded from Uniprot and used to identify and isolate the proper CDS sequences from the wild type *P. trichocarpa* genome.

#### *MKK2* locus target oligo synthesis

For vector construction, a pair of oligos (Supplementary Table 6; *MKK2* Oligo-F and -R) were designed flanked by *Bs*aI adaptors. Synthesized oligos were then ligated into pRGEB31 vectors following BsaI digestion ^34^ to construct pgRNA (Supplementary Figure 2a). Afterward, all vectors were transferred into *E. coli* (DH5α) and propagated under 37°C for 8 hours (Normal conditions). Vectors were then extracted using a plasmid midi kit (Qiagen, USA) and confirmed by Sanger sequencing (GeneScript, Nanjing).

### Transformation detection and confirmation

#### Western blotting

We used Western blotting to validate the successful integration of exogenous *BleoR* in the edited events genome. For extraction of proteins, 150 mg fresh leaves of five-week-old buds were milled in 500 μl extraction buffer (125 mM Tris, pH 6.8, 4 M Urea, 5% β - mercaptoethanol, 4% w/v SDS). Centrifugation was then performed at 13,000 rpm for 10 min, and the supernatant was collected for gel analysis. The extracted protein was then boiled in loading buffer (24% (w/v) glycerol, 100 mM Tris, 0.05% (w/v) Bromophenol Blue, 4% v/v β - mercaptoethanol, 8% (w/v) SDS) for 10 min. The extracted protein was analyzed by SDS-PAGE and visualized using Coomassie brilliant blue R-250 staining. Western blotting was then performed as described by ^35^, using a rabbit anti-His polyclonal antibody developed in our laboratory as the primary antibody and peroxidase-conjugated goat antirabbit IgG (Zhongshan Biotechnique, Beijing, China) as the secondary antibody.

#### RT-PCR

We performed RT-PCR to verify the whole and precise integration of exogenous *BleoR* regarding designed primers and complete transcription of *BleoR* and *MKK2* resulting from efficient HDR. Total RNA (100 ng/ml) was extracted by TRIzol from young leaves of five-week-old buds grown on Zeocin-containing medium. Reverse transcription was then carried out using total RNA and oligo-dT primers to synthesize the first strand cDNA using the PrimeScript One-Step RT-PCR Kit (Ver.2, Takara Biotechnology, Dalian, China) according to the manufacturer’s instructions. Afterward, two RT-PCR experiments were designed for examination of *MKK2* transcription and proper HDR. The first RT-PCR was intended to isolate a 920 bp fragment of *MKK2* CDS (Supplementary Table 6, RT-F and R) with the primers designed to amplify from the 5′ region of exon 9 (15 bp) and 3′ region of exon 8 (15 bp). The purpose was to show the precise attaching of exon 8 and 9 to direct the transcription of *MKK2* correctly. A second RT-PCR was performed to isolate a 413 bp fragment of recombinant CDS (Supplementary Table 6, RT-F-107 and RT-R-519). The forward primer was designed from *BleoR*, and the reverse primer was designed from exon 7 of *MKK2* with the purposes of showing the occurrence of HDR via transcription of a single mRNA from *MKK2* and *BleoR*.

#### DNA sequencing

Genomic DNA was extracted from leaves of five-week-old buds grown on Zeocin-containing medium using a DNeasy Plant Mini Kit (Qiagen, USA). The quality of the extracted genomic DNA (250–350 ng/μl) was determined by a BioDrop spectrophotometer (UK).

DNA sequencing was performed to evaluate and confirm the western blotting, RT-PCR, and southern blotting results. In addition, DNA sequencing was applied to assess the happened kind of mutations during genome editing. For DNA sequencing, we carried out PCR using designed primers (Supplementary Table 6, *MKK2*-S-7F and *MKK2*-S-1139R), Easy Taq polymerase (TransGene Biotech), and 50 ng of extracted genomic DNA as a template. Desired amplicons were then cloned into pEASY T3 vector (TransGen Biotech Co, Beijing, China) and used for Sanger sequencing (GeneScript, Nanjing, China), followed by alignment and data analysis (Supplementary Figures 4-10).

#### Southern blotting

Southern blotting was designed and performed to confirm Western blotting results and investigate the potential off-targets through genome editing. First, genomic DNA (500 ng) was cleaved with *BamH*I and *Hind*III at 37 °C for 4 h. The digested DNA was then used as a PCR template to label a 160 bp probe from integrated *BleoR* CDS into the genomic DNA (Supplementary Table 6; S-F and -R). DIG (digoxigenin) reagent was used for this procedure according to the manufacturer’s instructions (catalog number 11745832910; Roche, Basel, Switzerland). The PCR product was then electrophoresed on a 0.8% agarose gel. Finally, the separated fragments were shifted on a Hybond N+ nylon membrane (Amersham Biosciences BV, Eindhoven, The Netherlands).

### Evaluation of HDR validity and efficiency

#### HDR validity by TaqMan real-time PCR

TaqMan real-time PCR was performed to confirm the HDR validity with the proper integration of exogenous *BleoR* in both homology arms. For this purpose, TaqMan assay applying dye labels such as FAM and VIC was performed using an Applied Biosystems real-time PCR (Applied Biosystems, Thermo Scientific, USA). High-quality grown buds genomic DNA (refer to the southern blotting) was used as the template for running TaqMan real-time PCR. In this assay, two fluorescent markers, FAM and VIC, will attach to the 5′ region of the probe, while a non-fluorescent quencher (NFQ) binds to the 3′ region. Therefore, we designed primers to probe two 150 bp fragments FAM1 (Supplementary Table 6, FAM1-F and -R) and FAM2 (Supplementary Table 6, FAM2-F and -R). In detail, FAM1 could probe 114 bp nucleotides from the 5′ homology arm and 36 bp from *BleoR*. Also, FAM2 could probe 105 bp nucleotides from the 3′ homology arm and 45 bp from the *BleoR* (Supplementary Figure 15). In addition, primers (Supplementary Table 6, VIC-F, and -R) were also designed to probe one 106 bp fragment VIC on the actin gene as housekeeping (Supplementary Figure 15). Thus, all samples were analyzed in quadruplicate.

#### HDR efficiency by real-time PCR

Given the purpose of this study to express an external *BleoR* gene into the desired position from the *P. trichocarpa* genome, we evaluated HDR efficiency by calculating the mean ΔΔCt of *BleoR* expression integrated into the poplar genome from all grown buds across five designed experiments separately. In this step, we used the synthesized cDNA (Point to the RT-PCR section) and designed primers (Supplementary Table 6, *BleoR*-52F and -151R) to carry out real-time PCR. The Fast Start Universal SYBR Green Master mix (Rox; No. 04913914001: Roche, USA) was used with three technical repeats for each event. First, we used the achieved *BleoR* expressions from all grown buds on Zeocin (*BleoR* Ct mean) and desired reference (DDP Ct mean; plasmid extracted from *E. coli*) to evaluate the main ΔCt Mean. Then, the control group ΔCt was measured from a wild-type poplar for each experiment separately. Subsequently, the *BleoR* expression ΔΔCt mean was calculated by subtracting the control group ΔCt of the main ΔCt Mean. Finally, to balance the variation of each experiment event number, we considered zero for all the *BleoR* expressions up to 31 (Equals to ExV events), calculating the ratio of 100-fold of *BleoR* expression ΔΔCt mean:31, determining HDR% for each event (Supplementary Table 2a). Thus, One way-ANOVA descriptive statistics computed from all events HDR% has been used for analyzing HDR% efficiency in Figure 3i (Supplementary Table 2b).

### Salt Stress Phenotypic Evaluation

Given the roles of *MKK2* in plant protection against environmental stresses ^22,31^, and to confirm the exact HDR occurred through *MKK2* exons 7, 8, and 9 in the recovered transgenic lines, *MKK2* expression and phenotypic evaluations, specifically salt stress tolerance, were performed relative to WT, non-transformed poplar. Recovered events were planted into soil and transferred to the greenhouse. After two weeks of acclimation in a greenhouse, total RNA was isolated from WT leaves as a control and all transferred recovered events to evaluate the *MKK2* expression with RT-qPCR. For salt stress response evaluation, all recovered events were irrigated daily with 25 mM NaCl for one week following acclimation to the greenhouse. Total RNA was extracted from leaves of surviving events to perform RT-qPCR. Each reaction was performed in triplicate. Stem lengths (mm) and -diameters (mm) were also measured before and after salt stress.

### Statistical analysis

All data were analyzed using One-Way ANOVA with Turkey post-hoc comparisons calculated by OriginPro 2018 software (Northampton, USA). Differences were analyzed when the confidence intervals presented no overlap of the mean values with an error value of 0.05.

## References

1. Li, S. et al. Precise gene replacement in rice by RNA transcript-templated homologous recombination. Nat Biotechnol 37, 445–450; 10.1038/s41587-019-0065-7 (2019).

2. Cermak, T., Baltes, N. J., Cegan, R., Zhang, Y. & Voytas, D. F. High-frequency, precise modification of the tomato genome. Genome Biol 16, 232; 10.1186/s13059-015-0796-9 (2015).

3. Bewg, W. P., Ci, D. & Tsai, C.-J. Genome Editing in Trees. From Multiple Repair Pathways to Long-Term Stability. Frontiers in Plant Science 9; 10.3389/fpls.2018.01732 (2018).

4. Zhou, X., Jacobs, T. B., Xue, L.-J., Harding, S. A. & Tsai, C.-J. Exploiting SNP s for biallelic CRISPR mutations in the outcrossing woody perennial <em>Populus</em> reveals 4-coumarate. CoA ligase specificity and redundancy. New Phytologist 208, 298–301 (2015).

5. Schiml, S., Fauser, F. & Puchta, H. The CRISPR/Cas system can be used as nuclease for in planta gene targeting and as paired nickases for directed mutagenesis in Arabidopsis resulting in heritable progeny. Plant J 80, 1139–1150; 10.1111/tpj.12704 (2014).

6. Li, J. et al. Gene replacements and insertions in rice by intron targeting using CRISPR-Cas9. Nat Plants 2, 16139; 10.1038/nplants.2016.139 (2016).

7. Yang, D. et al. Enrichment of G2/M cell cycle phase in human pluripotent stem cells enhances HDR-mediated gene repair with customizable endonucleases. Sci Rep 6, 21264; 10.1038/srep21264 (2016).

8. Gil-Humanes, J. et al. High-efficiency gene targeting in hexaploid wheat using DNA replicons and CRISPR/Cas9. Plant J 89, 1251–1262; 10.1111/tpj.13446 (2017).

9. Svitashev, S., Schwartz, C., Lenderts, B., Young, J. K. & Cigan, A. M. Genome editing in maize directed by CRISPR-Cas9 ribonucleoprotein complexes. Nat Commun 7; 10.1038/ncomms13274 (2016).

10. van Vu, T. et al. Highly efficient homology-directed repair using CRISPR/Cpf1-geminiviral replicon in tomato. Plant Biotechnol J; 10.1111/pbi.13373 (2020).

11. Movahedi, A. et al. Expression of the chickpea CarNAC3 gene enhances salinity and drought tolerance in transgenic poplars. Plant Cell Tiss Org 120, 141–154; 10.1007/s11240-014-0588-z (2015).

12. Movahedi, A., Zhang, J. X., Amirian, R. & Zhuge, Q. An efficient Agrobacterium-mediated transformation system for poplar. Int J Mol Sci 15, 10780–10793; 10.3390/ijms150610780 (2014).

13. An, Y. et al. Efficient genome editing in populus using CRISPR/Cas12a. Front Plant Sci 11, 593938; 10.3389/fpls.2020.593938 (2020).

14. Ali, Z. et al. Fusion of the Cas9 endonuclease and the VirD2 relaxase facilitates homology-directed repair for precise genome engineering in rice. Commun Biol 3; 10.1038/s42003-020-0768-9 (2020).

15. Tran, N. T. et al. Enhancement of precise gene editing by the association of Cas9 with homologous recombination factors. Front Genet 10; 10.3389/fgene.2019.00365 (2019).

16. Hartun, F. & Puchta, H. Isolation of the complete cDNA of the Mre11 homologue of Arabidopsis (Accession No. AJ243822) indicates conservation of DNA recombination mechanisms between plants and other eucaryotes. (PGR99-132). Plant Physiol 121, 312 (1999).

17. Qi, Y. et al. Increasing frequencies of site-specific mutagenesis and gene targeting in Arabidopsis by manipulating DNA repair pathways. Genome Res 23, 547–554; 10.1101/gr.145557.112 (2013).

18. Grawunder, U., Zimmer, D., Kulesza, P. & Lieber, M. R. Requirement for an interaction of XRCC4 with DNA ligase IV for wild-type V(D)J recombination and DNA double-strand break repair in vivo. J Biol Chem 273, 24708–24714; 10.1074/jbc.273.38.24708 (1998).

19. West, C. E., Waterworth, W. M., Jiang, Q. & Bray, C. M. Arabidopsis DNA ligase IV is induced by gamma-irradiation and interacts with an Arabidopsis homologue of the double strand break repair protein XRCC4. Plant J 24, 67–78; 10.1046/j.1365-313x.2000.00856.x (2000).

20. Pierce, A. J., Hu, P., Han, M., Ellis, N. & Jasin, M. Ku DNA end-binding protein modulates homologous repair of double-strand breaks in mammalian cells. Genes Dev 15, 3237–3242; 10.1101/gad.946401 (2001).

21. Chen, X. Y. et al. The Liriodendron chinense MKK2 Gene Enhances Arabidopsis thaliana Salt Resistance. Forests 11; 10.3390/f11111160 (2020).

22. Mohanta, T. K., Arora, P. K., Mohanta, N., Parida, P. & Bae, H. Identification of new members of the MAPK gene family in plants shows diverse conserved domains and novel activation loop variants. BMC genomics 16, 58; 10.1186/s12864-015-1244-7 (2015).

23. Di Fan et al. Efficient CRISPR/Cas9-mediated Targeted Mutagenesis in Populus in the First Generation. Sci Rep-Uk 5, 12217; 10.1038/srep12217 (2015).

24. Hsu, P. D. et al. DNA targeting specificity of RNA-guided Cas9 nucleases. Nat Biotechnol 31, 827–832; 10.1038/nbt.2647 (2013).

25. Doench, J. G. et al. Rational design of highly active sgRNAs for CRISPR-Cas9-mediated gene inactivation. Nat Biotechnol 32, 1262–1267; 10.1038/nbt.3026 (2014).

26. Song, F. & Stieger, K. Optimizing the DNA Donor Template for Homology-Directed Repair of Double-Strand Breaks. Mol Ther Nucleic Acids 7, 53–60; 10.1016/j.omtn.2017.02.006 (2017).

27. Hajiahmadi, Z. et al. Strategies to Increase On-Target and Reduce Off-Target Effects of the CRISPR/Cas9 System in Plants. Int J Mol Sci 20; 10.3390/ijms20153719 (2019).

28. Liu, T. et al. Highly efficient CRISPR/Cas9-mediated targeted mutagenesis of multiple genes in Populus. Yi chuan = Hereditas 37, 1044–1052; 10.16288/j.yczz.15-303 (2015).

29. Shao, S. M. et al. Enhancing CRISPR/Cas9-mediated homology-directed repair in mammalian cells by expressing Saccharomyces cerevisiae Rad52. Int J Biochem Cell B 92, 43–52; 10.1016/j.biocel.2017.09.012 (2017).

30. Maruyama, T. et al. Increasing the efficiency of precise genome editing with CRISPR-Cas9 by inhibition of nonhomologous end joining. Nat Biotechnol 33, 538–542; 10.1038/nbt.3190 (2015).

31. Teige, M. et al. The MKK2 pathway mediates cold and salt stress signaling in Arabidopsis. Mol Cell 15, 141–152; 10.1016/j.molcel.2004.06.023 (2004).

32. Sun, W. et al. Expression analysis of genes encoding mitogen-activated protein kinases in maize provides a key link between abiotic stress signaling and plant reproduction. Funct Integr Genomic 15, 107–120; 10.1007/s10142-014-0410-3 (2015).

33. Zhang, J. P. et al. Efficient precise knockin with a double cut HDR donor after CRISPR/Cas9-mediated double-stranded DNA cleavage. Genome Biol 18, 35; 10.1186/s13059-017-1164-8 (2017).

34. Xie, K. B. & Yang, Y. N. RNA-Guided Genome Editing in Plants Using a CRISPRCas System. Mol Plant 6, 1975–1983; 10.1093/mp/sst119 (2013).

35. Sambrook, J., Fritsch, E. F. & Maniatis, T. Molecular cloning. a laboratory manual. 2nd ed. (Cold Spring Harbor Laboratory Press, Cold Spring Harbor, NY, 1989).

