## Supplementary figures and images for "Precise exogenous insertion and sequence replacements in poplar by simultaneous HDR overexpression and NHEJ suppression using CRISPR-Cas9" (from another version of this preprint: version 6, 2021-02-03)

### Suppl. 2

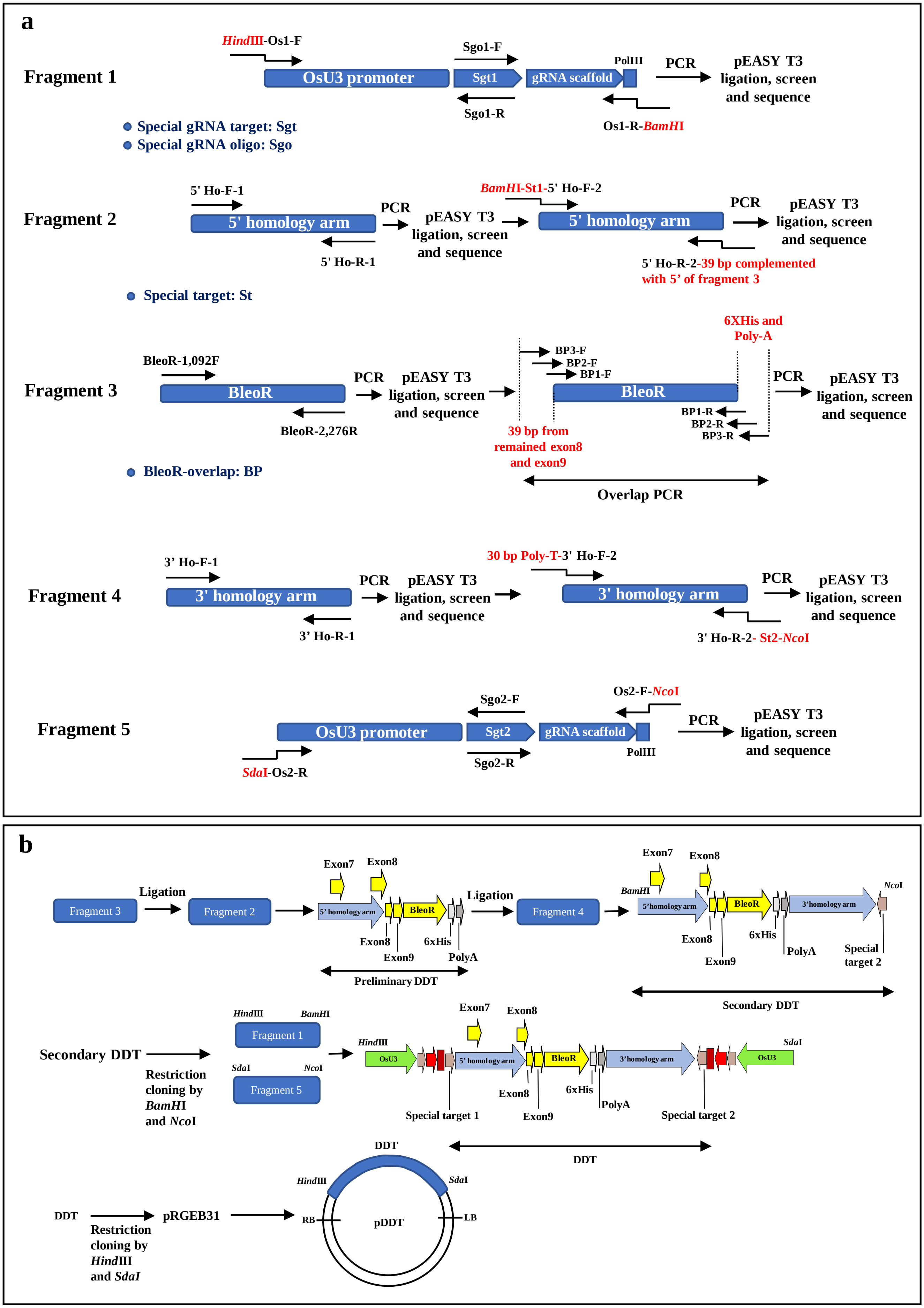


**Supplementary 2:** Schematic construction of DDT and pDDT fragments, primers and oligos.

### Suppl. 7

#### Supplementary 7: Alignment of events involved in experiment I.

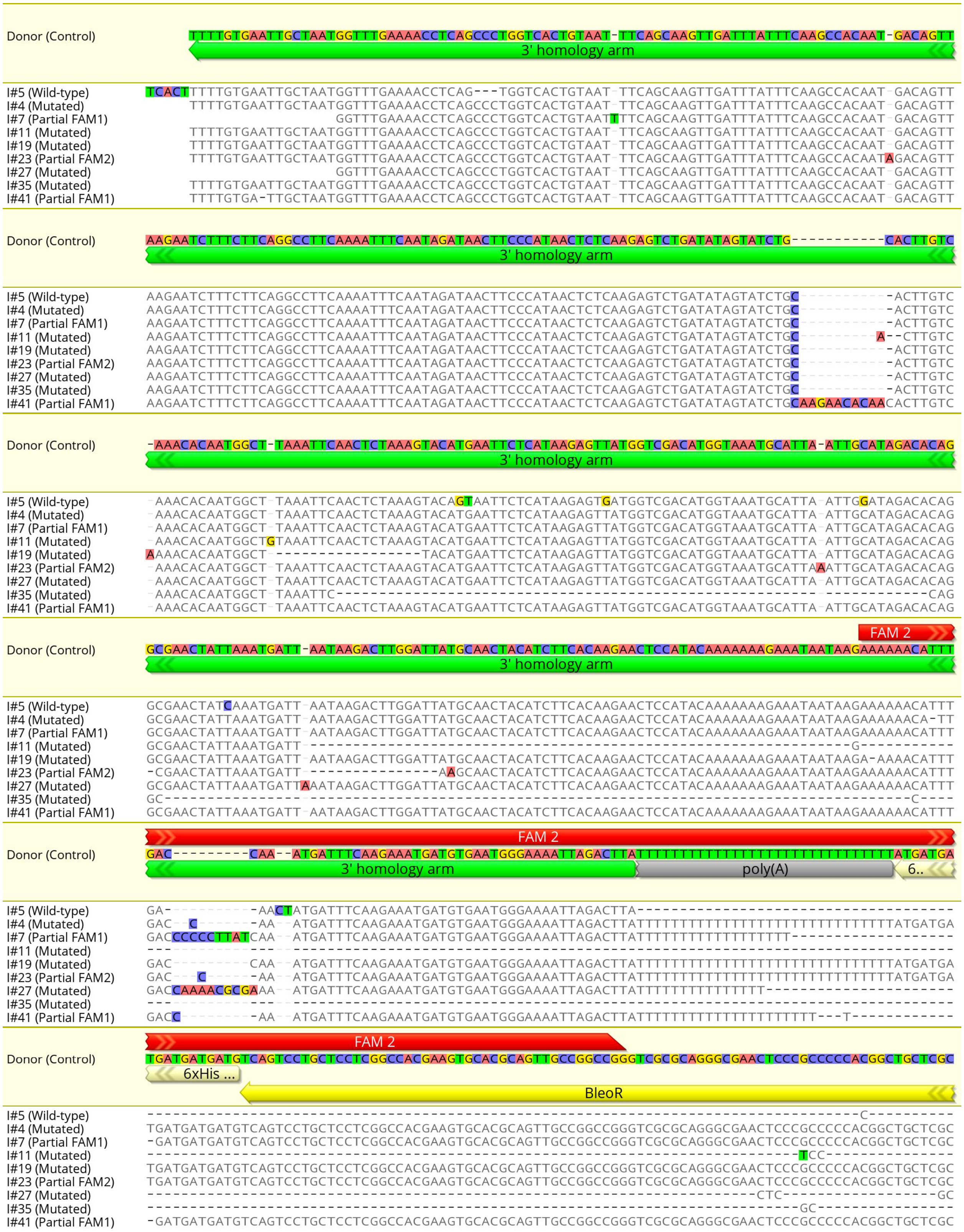

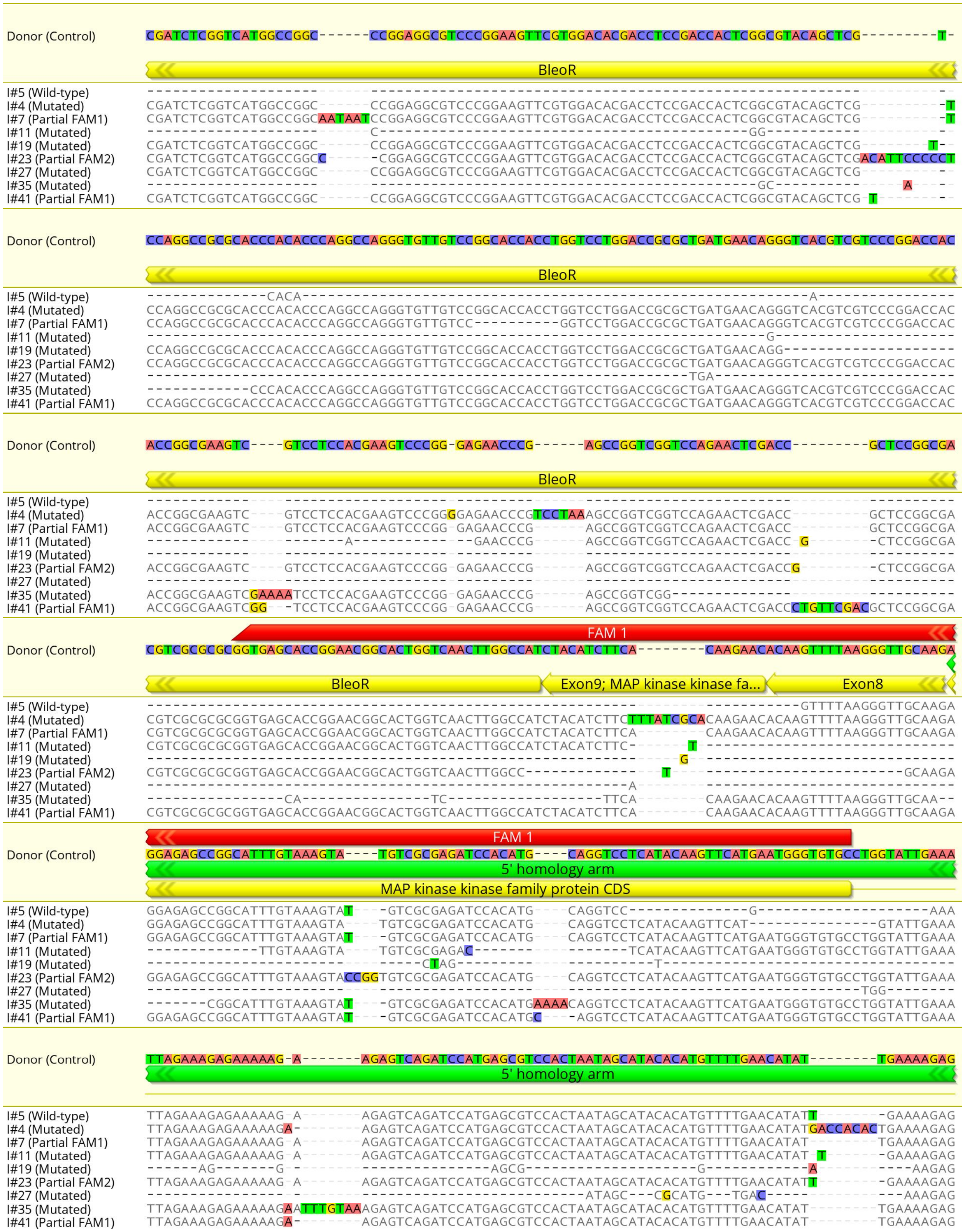

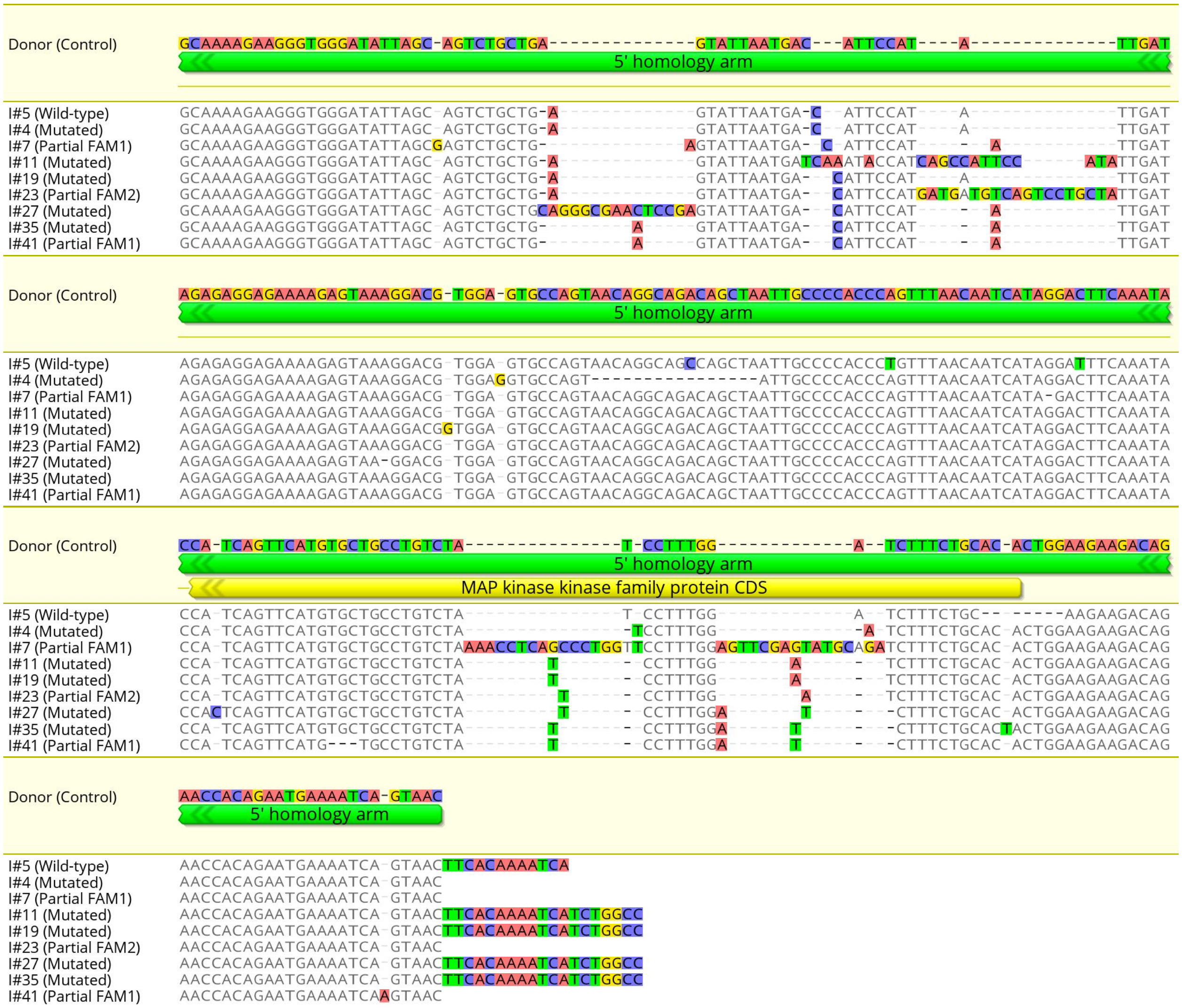

### Suppl. 10

#### Supplementary 10: Alignment of events involved in experiment IV.

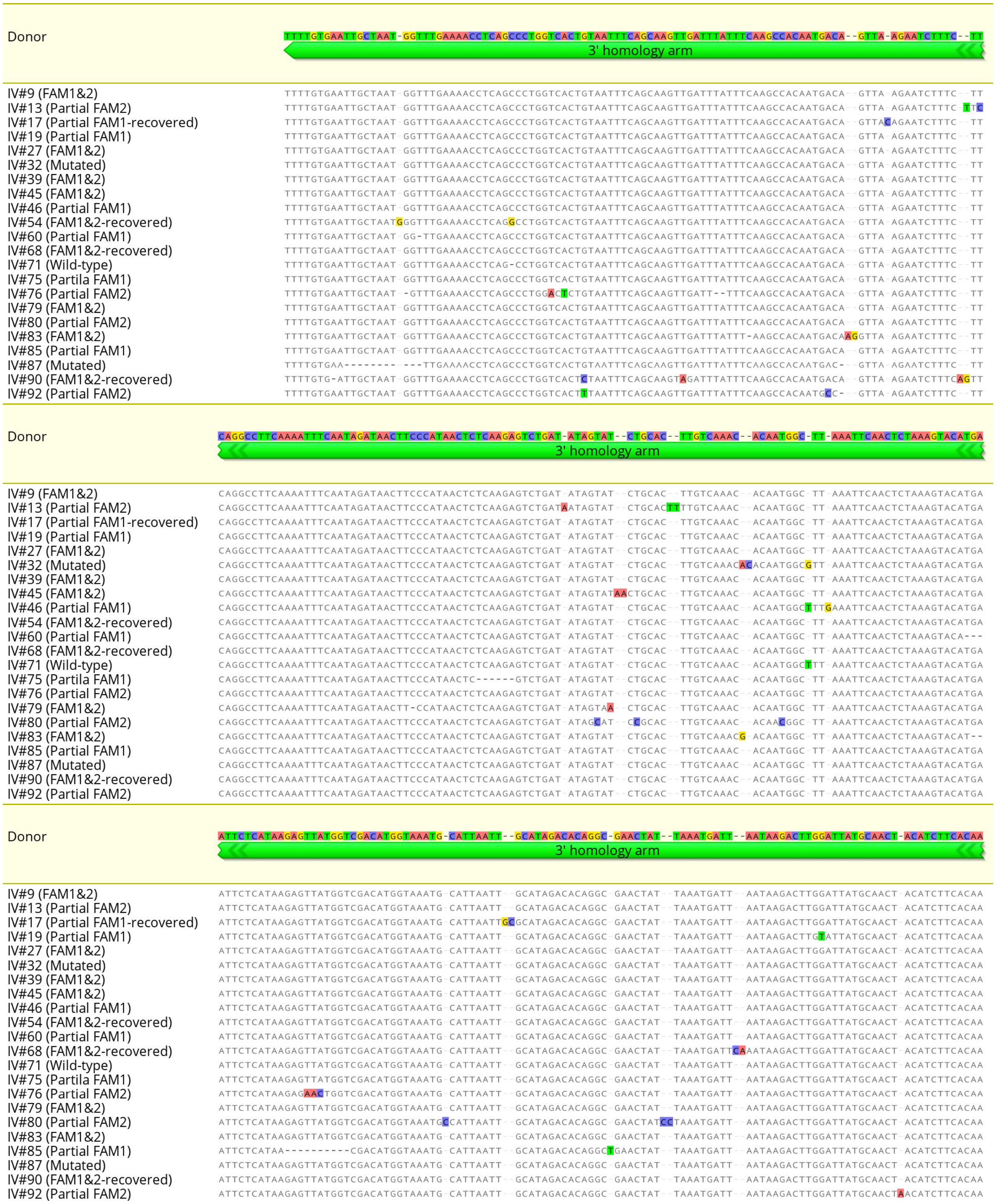

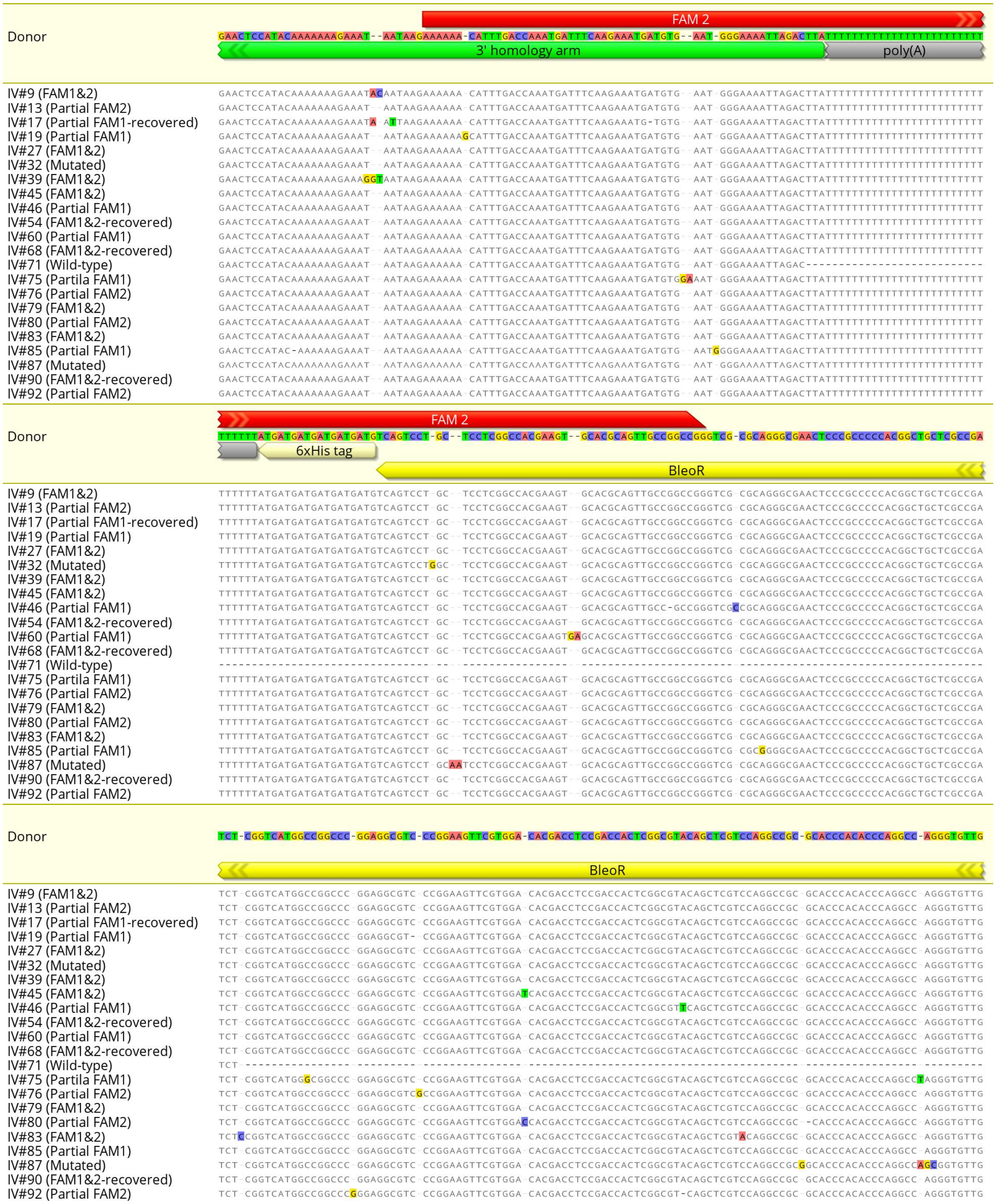

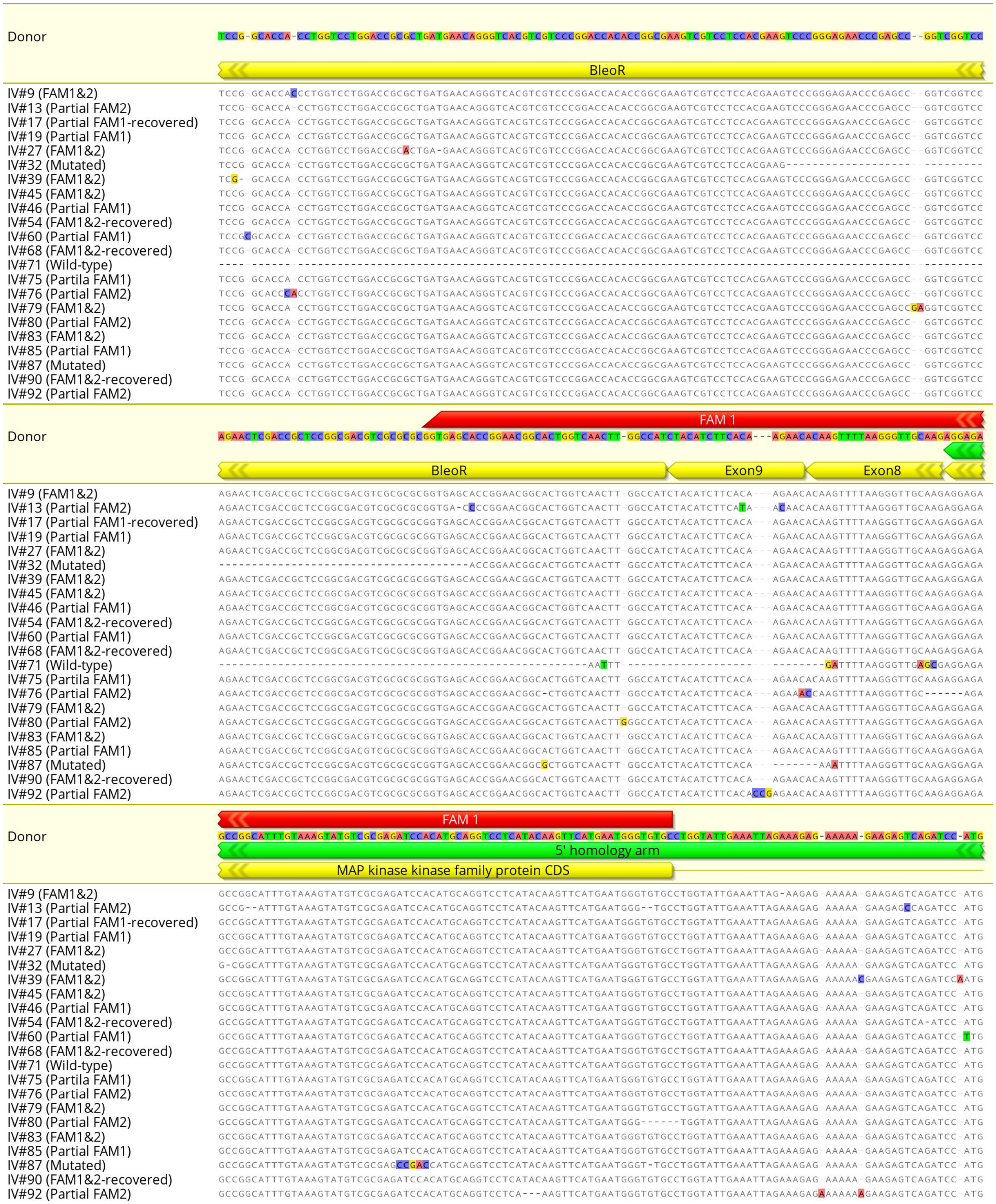

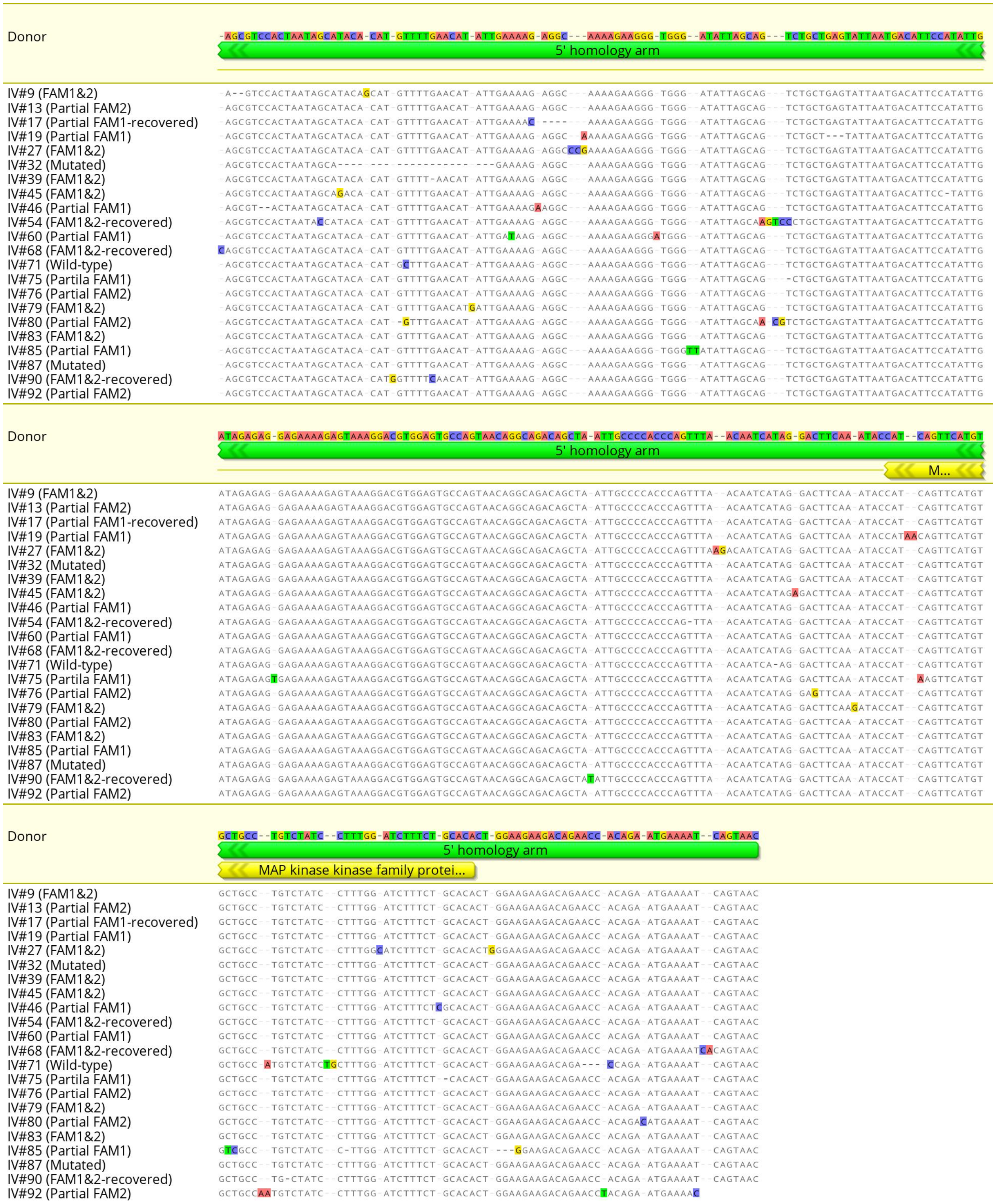

### Suppl. 11

#### Supplementary 11: Alignment of events involved in experiment V.

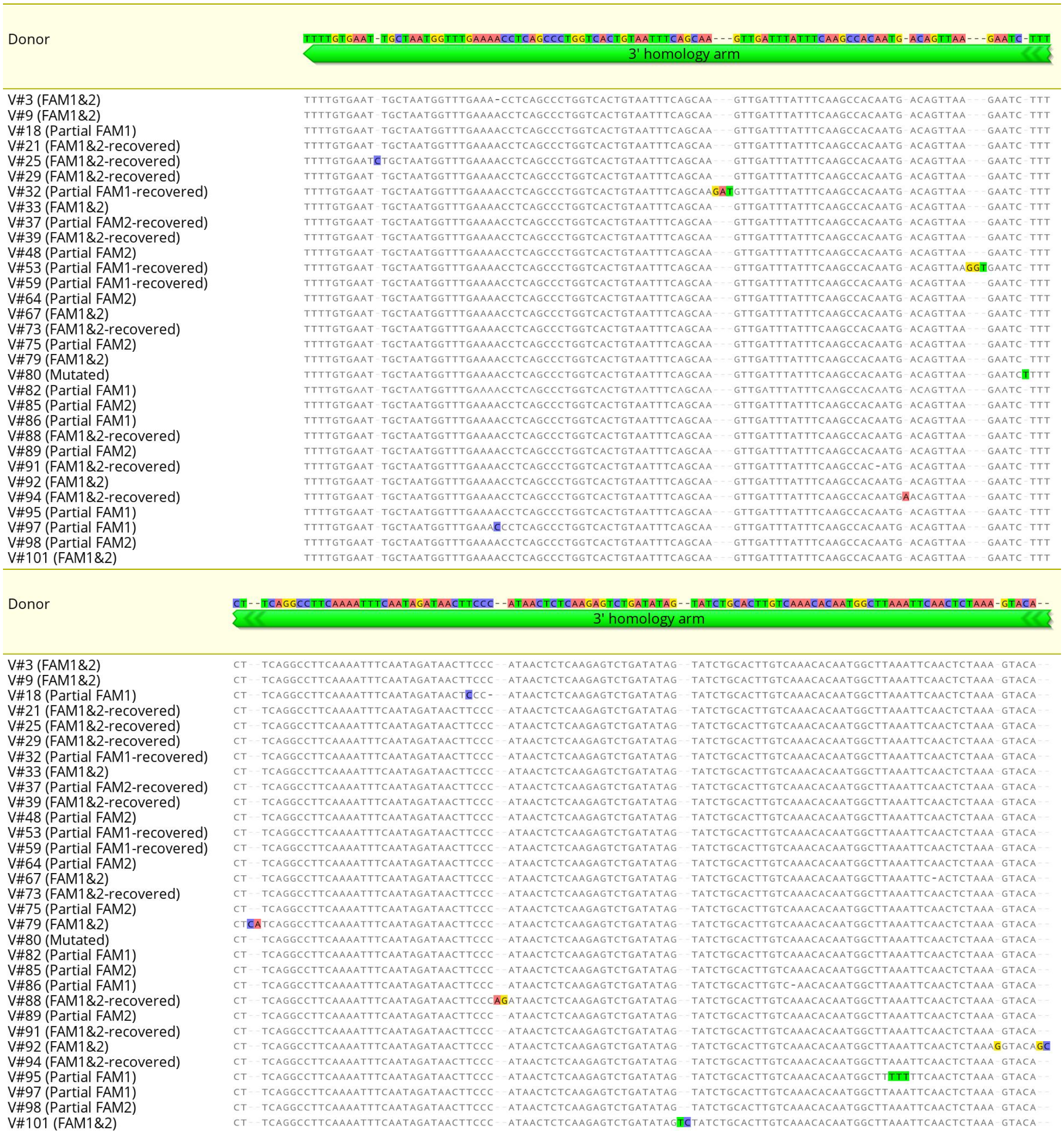

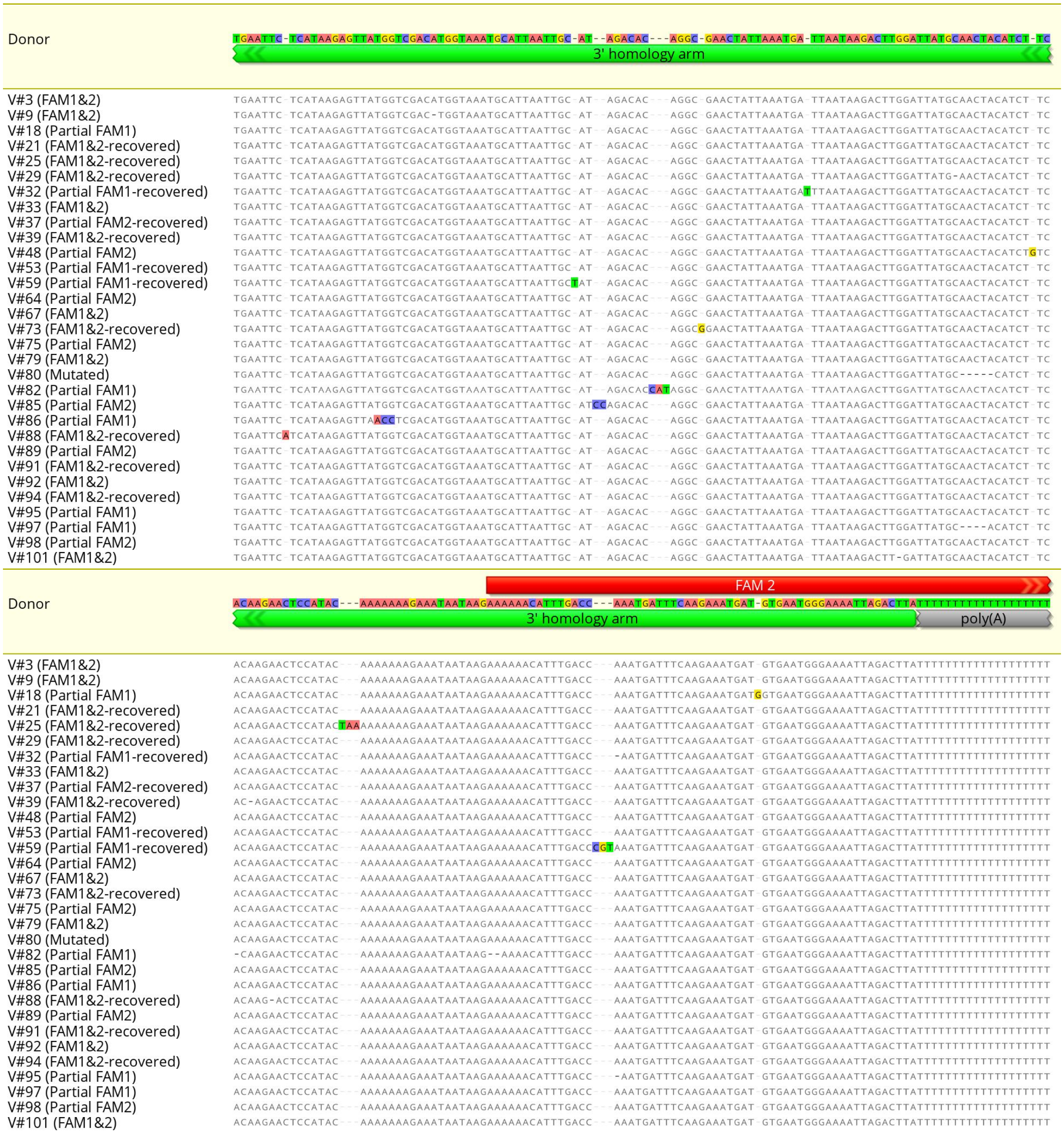

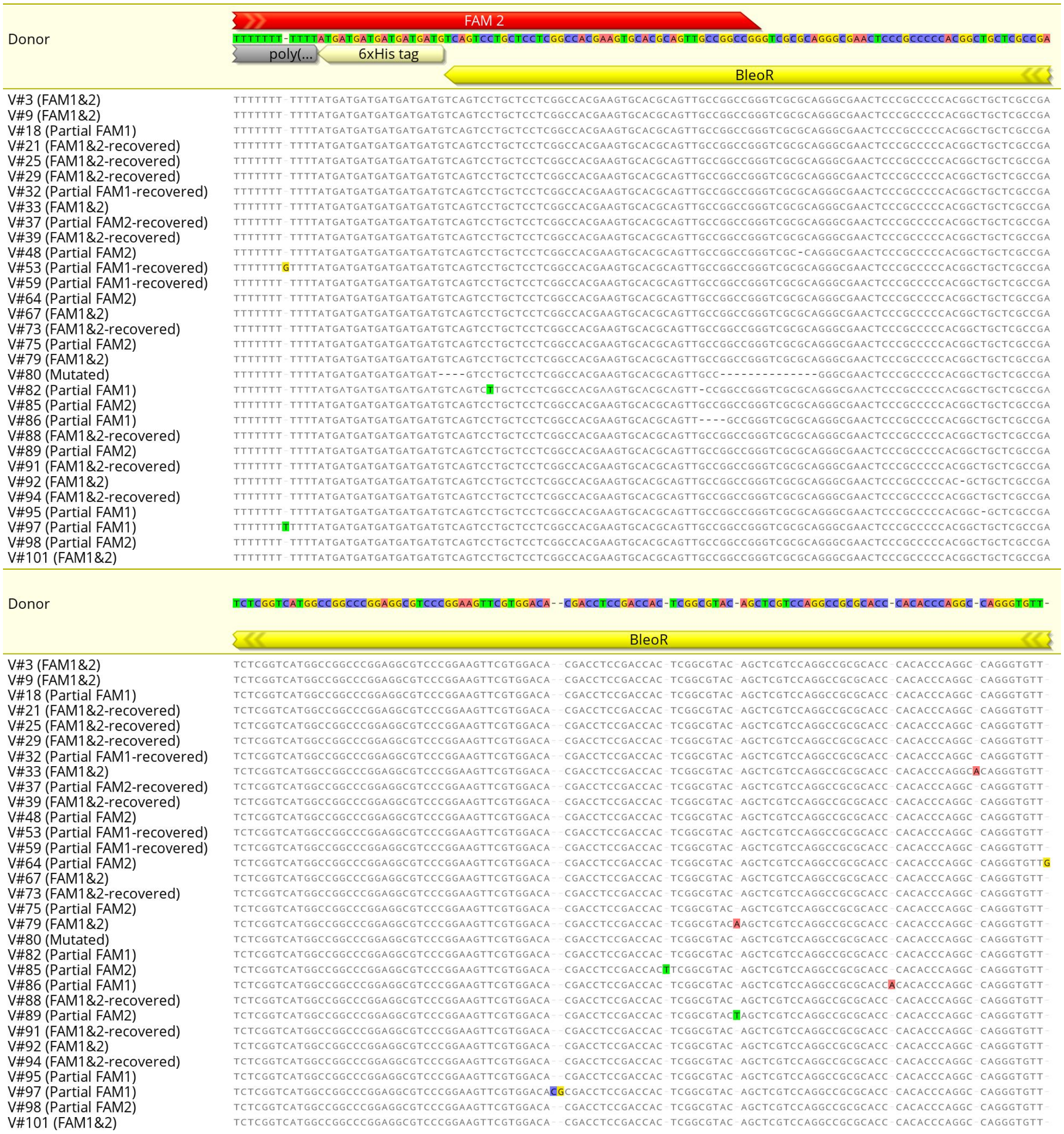

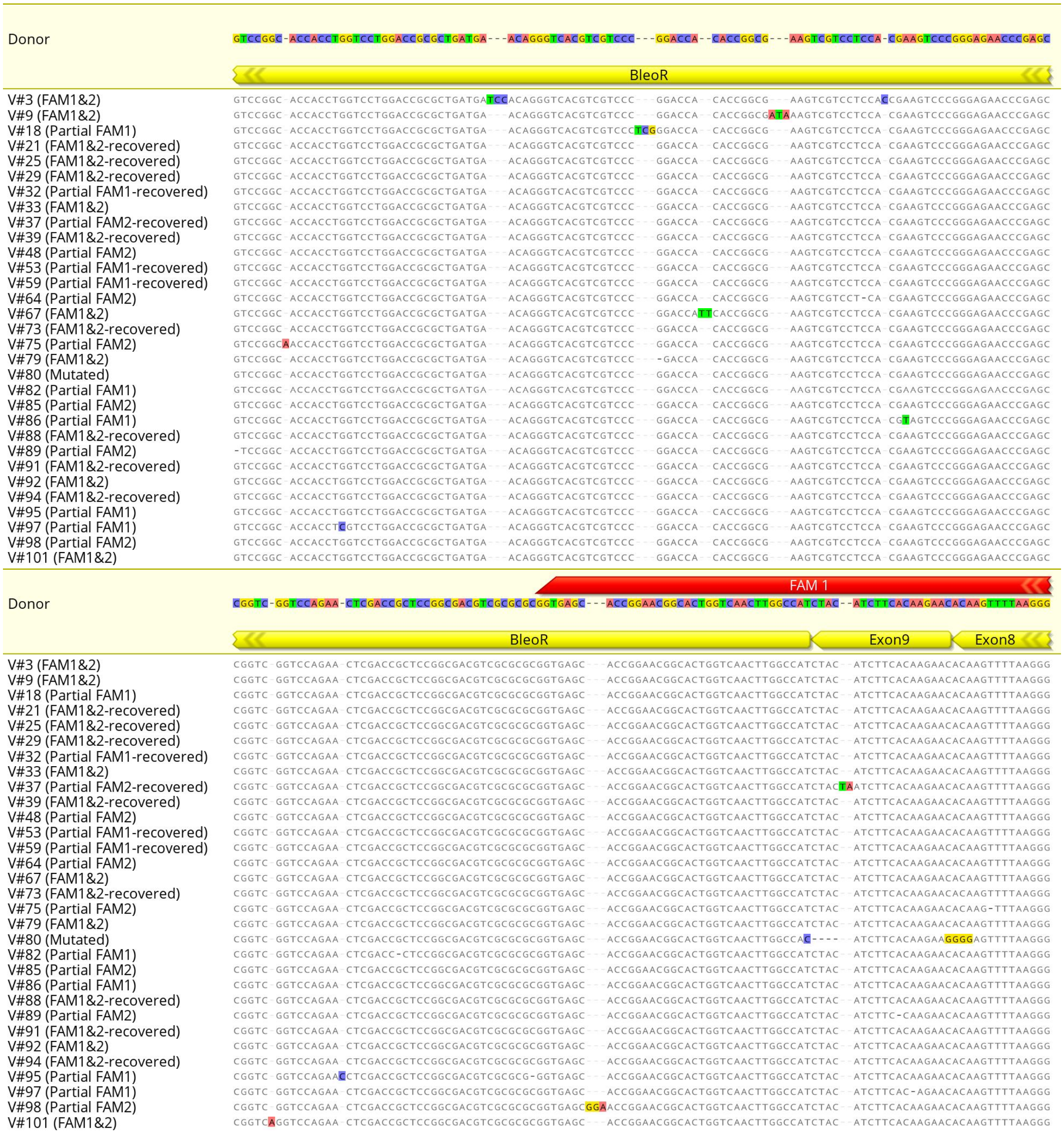

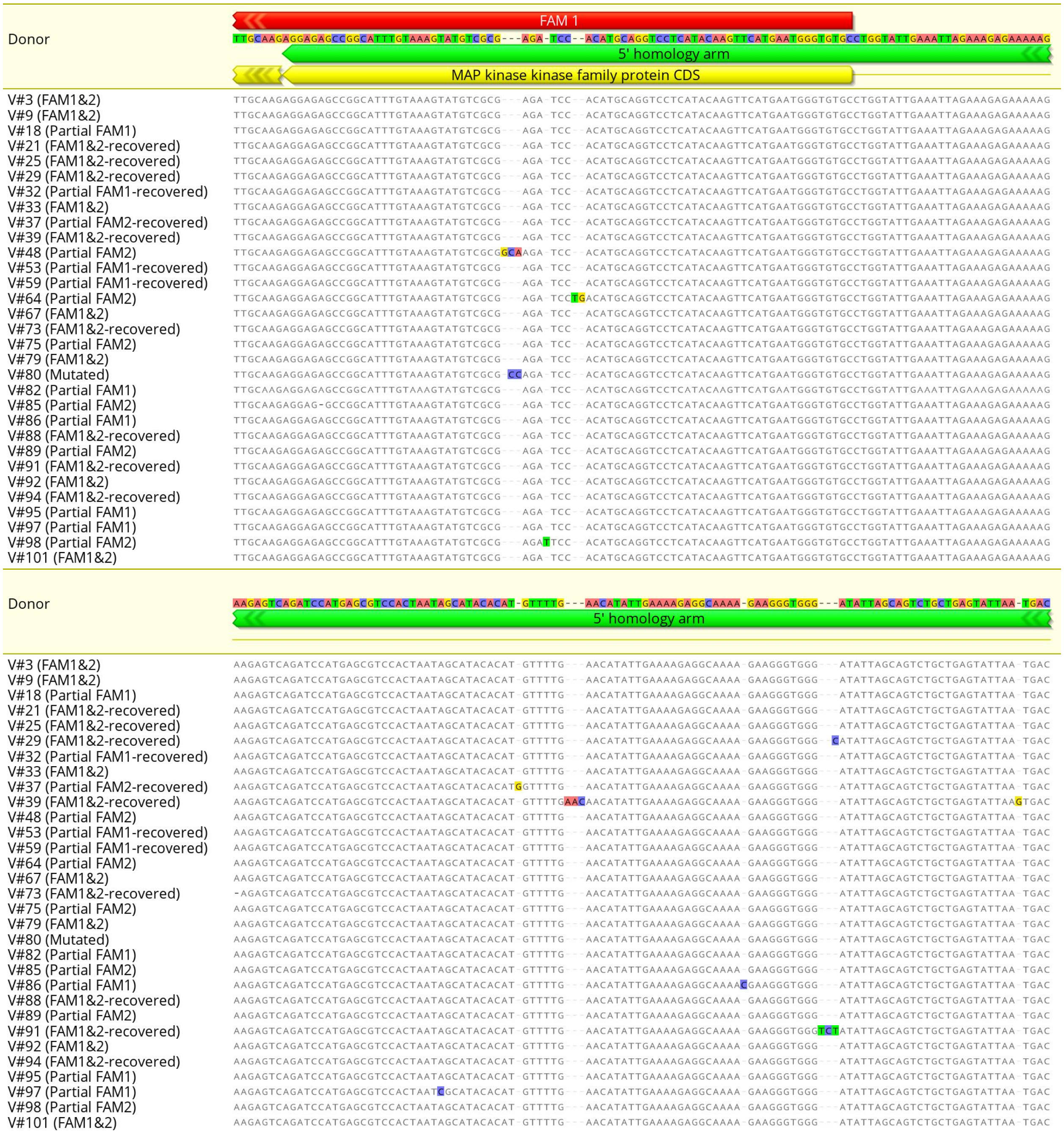

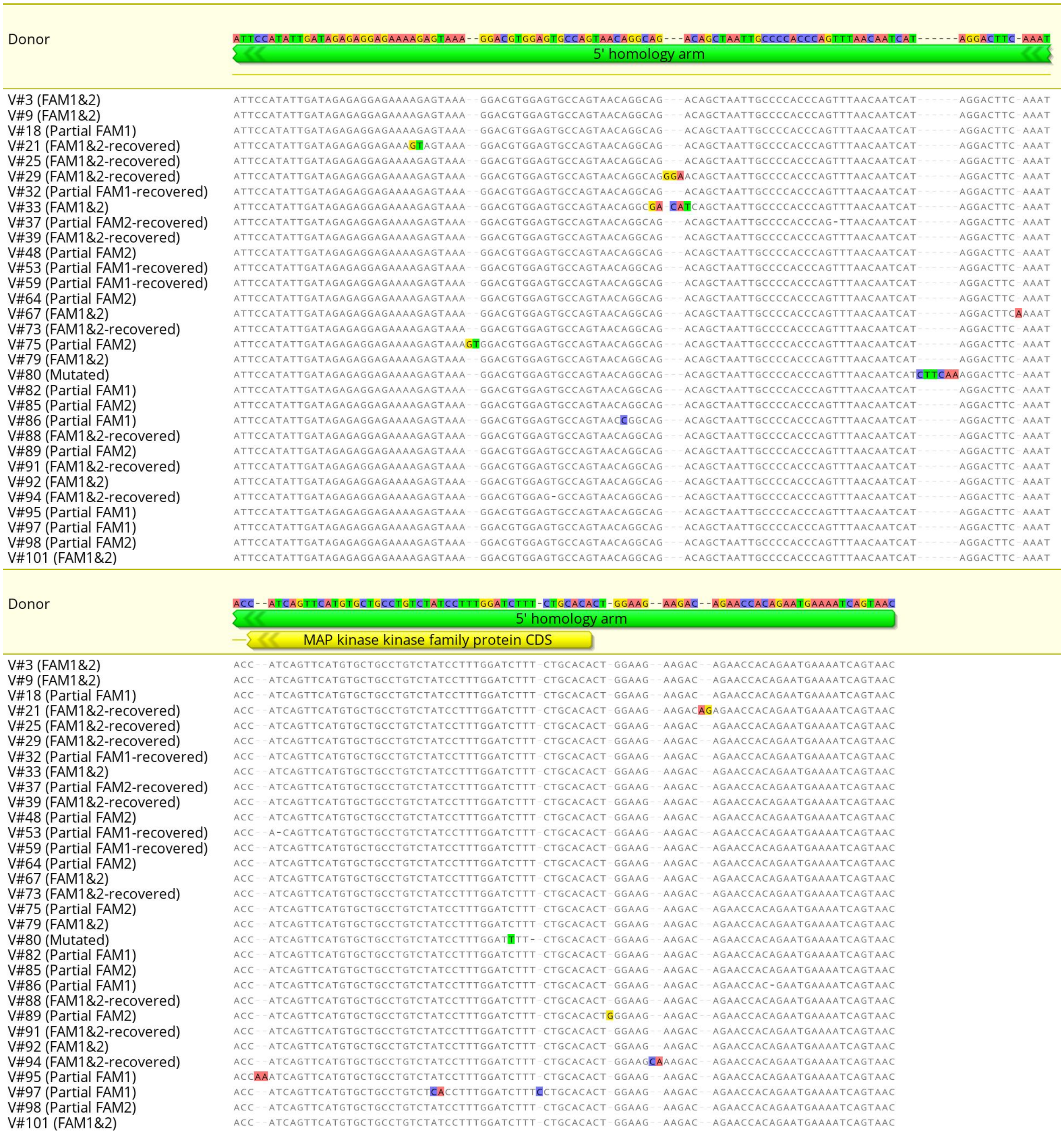
