## Supplementary material for "Precise exogenous insertion and sequence replacements in poplar by simultaneous HDR overexpression and NHEJ suppression using CRISPR-Cas9": Suppl. 3 (from another version of this preprint: version 6, 2021-02-03)

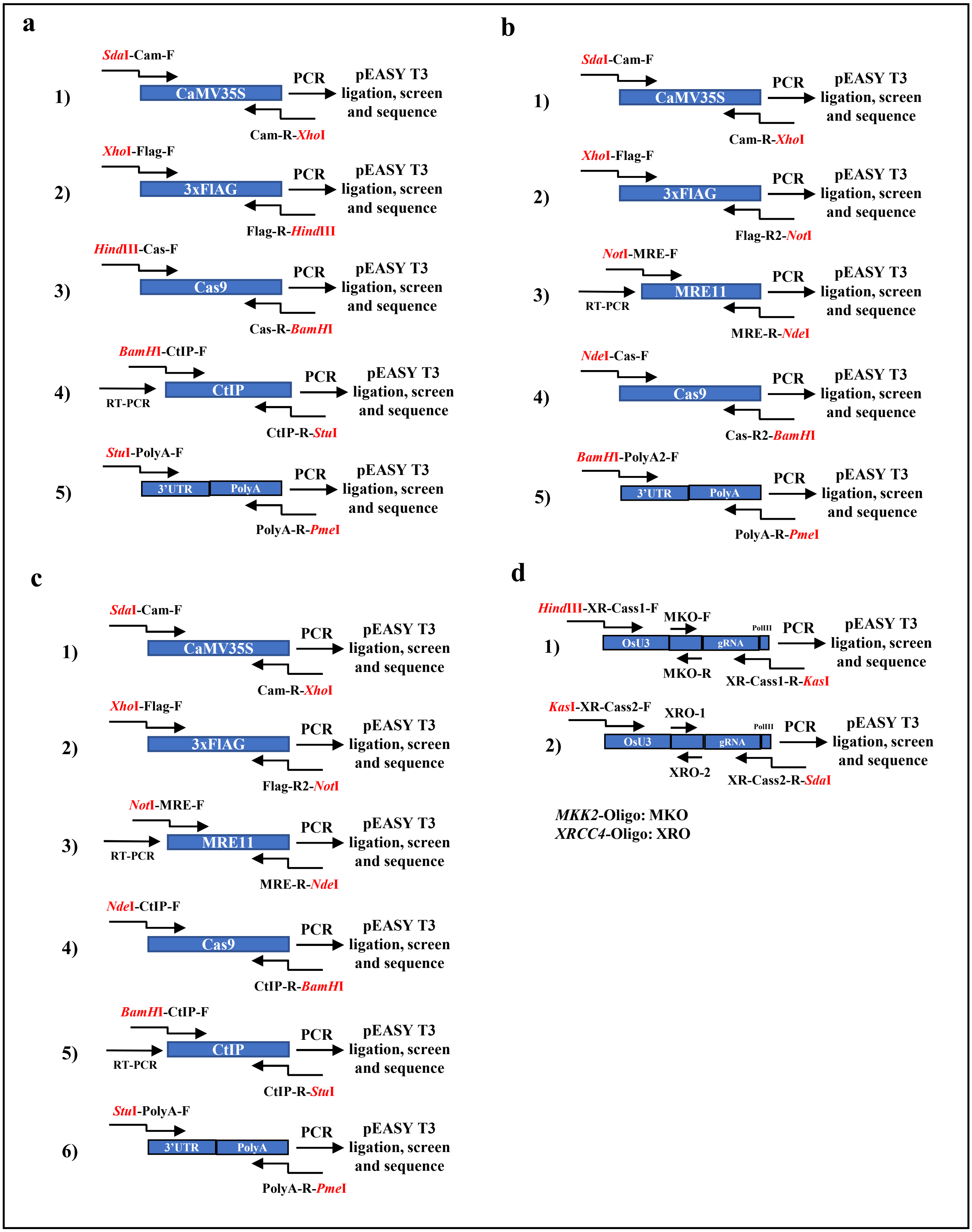


**Supplementary 3:** Schematic construction of CtIP, MRE11, CtIP+MRE11 and XRCC4 cassette primers and oligos.
