## Supplementary material for "Precise exogenous insertion and sequence replacements in poplar by simultaneous HDR overexpression and NHEJ suppression using CRISPR-Cas9": Suppl. 4 (from another version of this preprint: version 6, 2021-02-03)

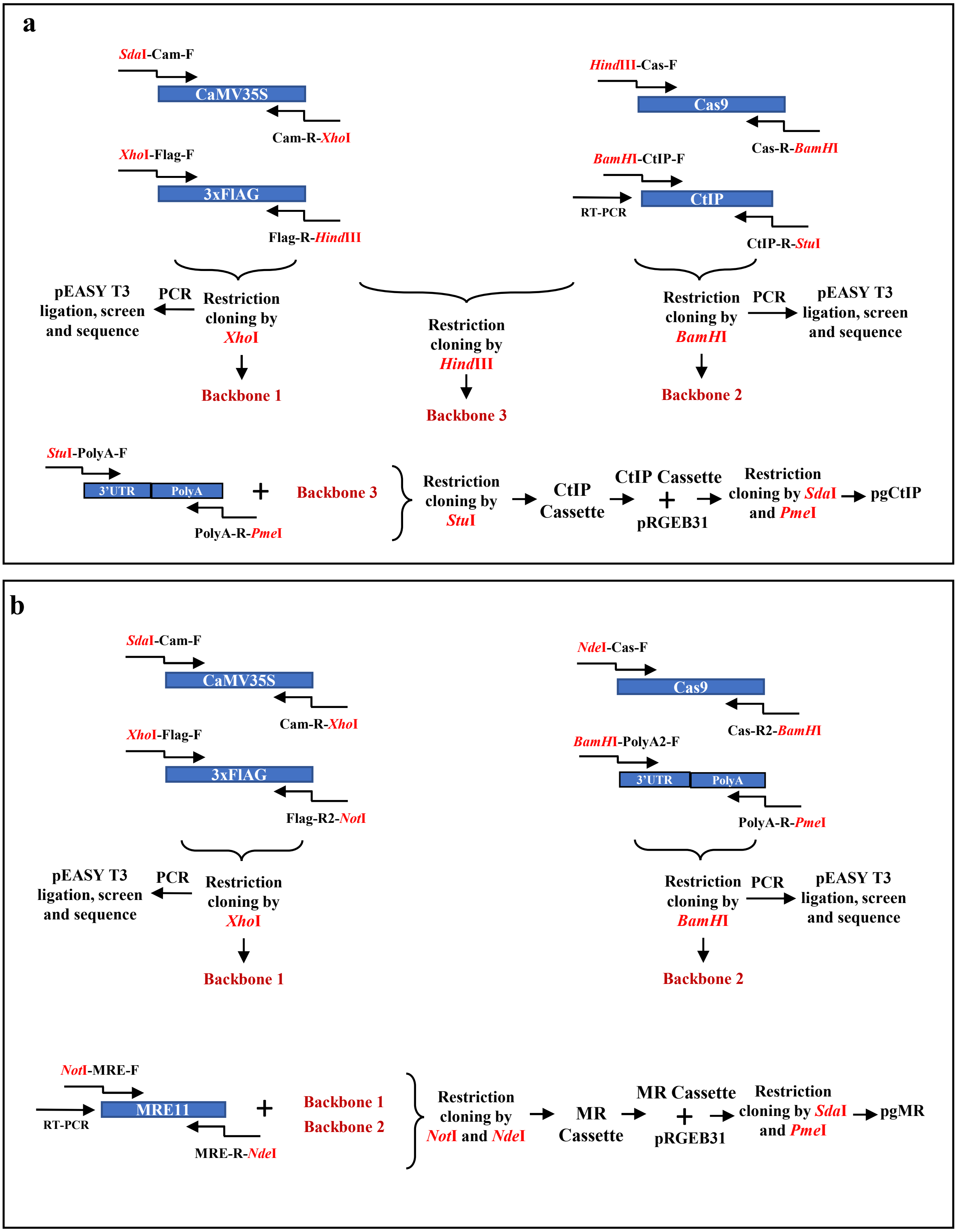


**Supplementary 4:** Schematic construction of cassettes (CtIP and MR) and vectors (pgCtIP and pgMR) and their primers.
