## Supplementary material for "Precise exogenous insertion and sequence replacements in poplar by simultaneous HDR overexpression and NHEJ suppression using CRISPR-Cas9": Suppl. 5 (from another version of this preprint: version 6, 2021-02-03)

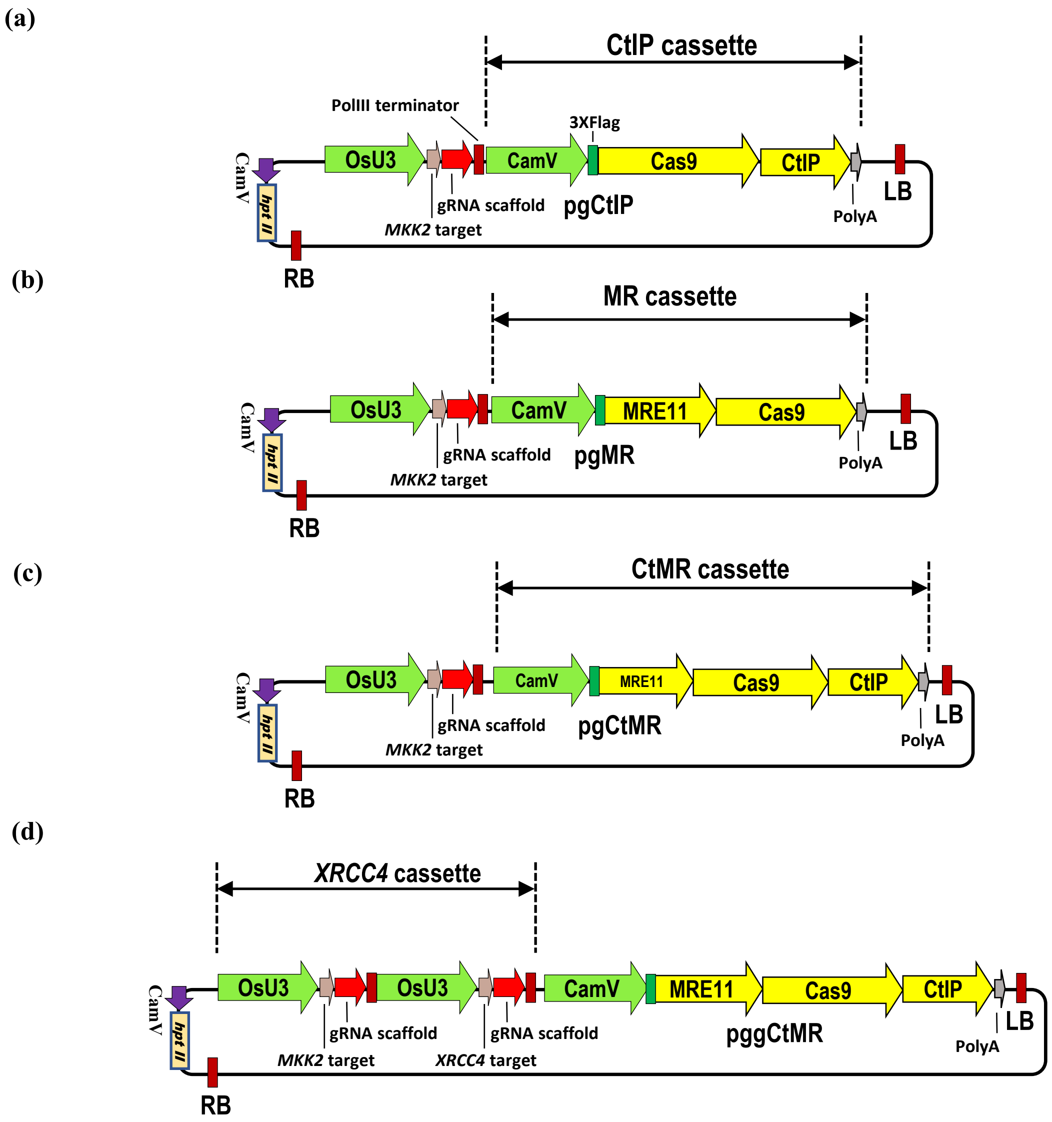


**Supplementary 5**: Schematics of constructed cassettes and plasmids. **(a)** pgCtIP plasmid including CtIP cassette. **(b)** pgMR plasmid including MR cassette. **(c)** pgCtMR plasmid including CtMR cassette. **(d)** pggCtMR plasmid including *XRCC4* cassette.
