## Supplementary material for "Precise exogenous insertion and sequence replacements in poplar by simultaneous HDR overexpression and NHEJ suppression using CRISPR-Cas9": Suppl. 6 (from another version of this preprint: version 6, 2021-02-03)

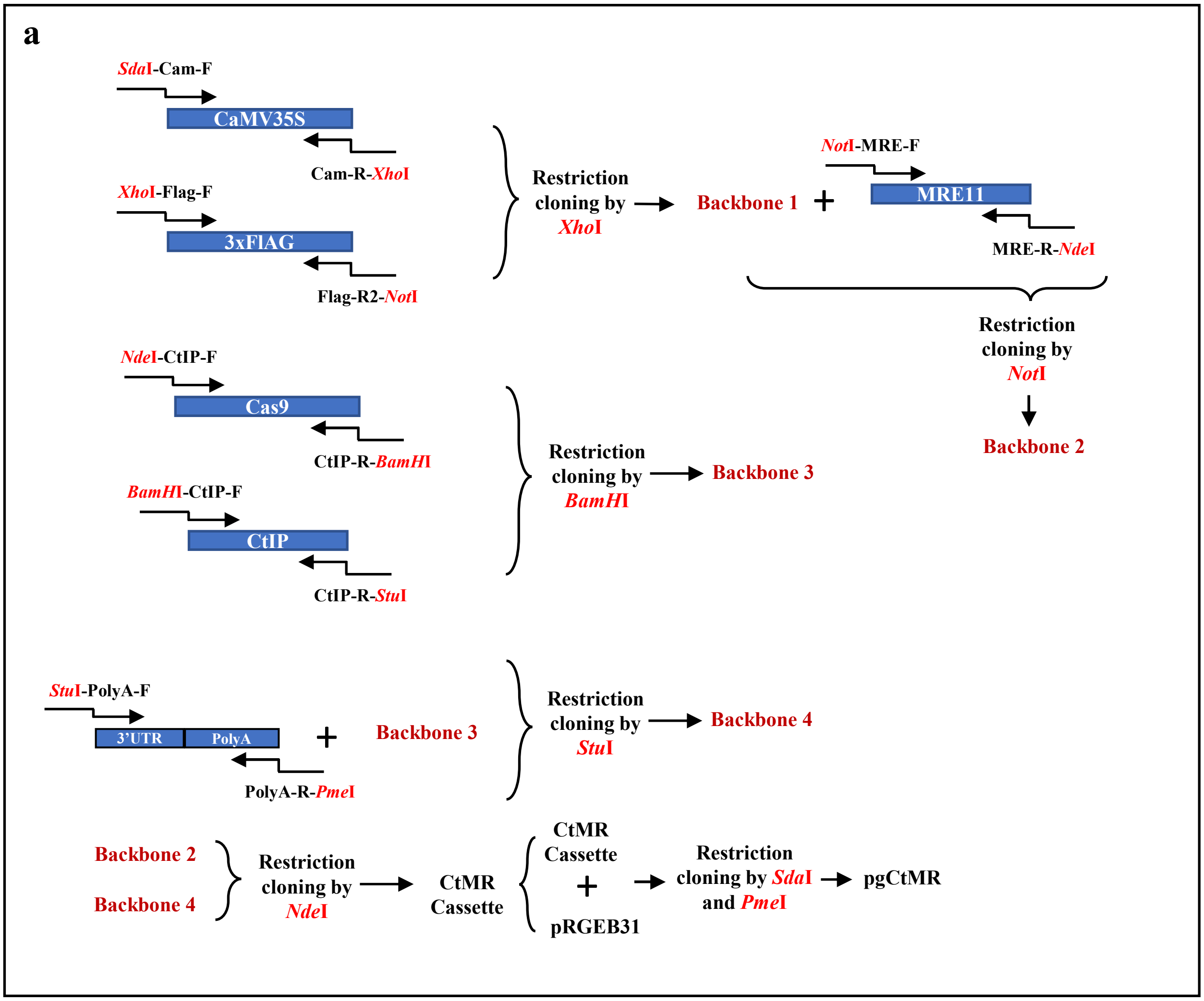


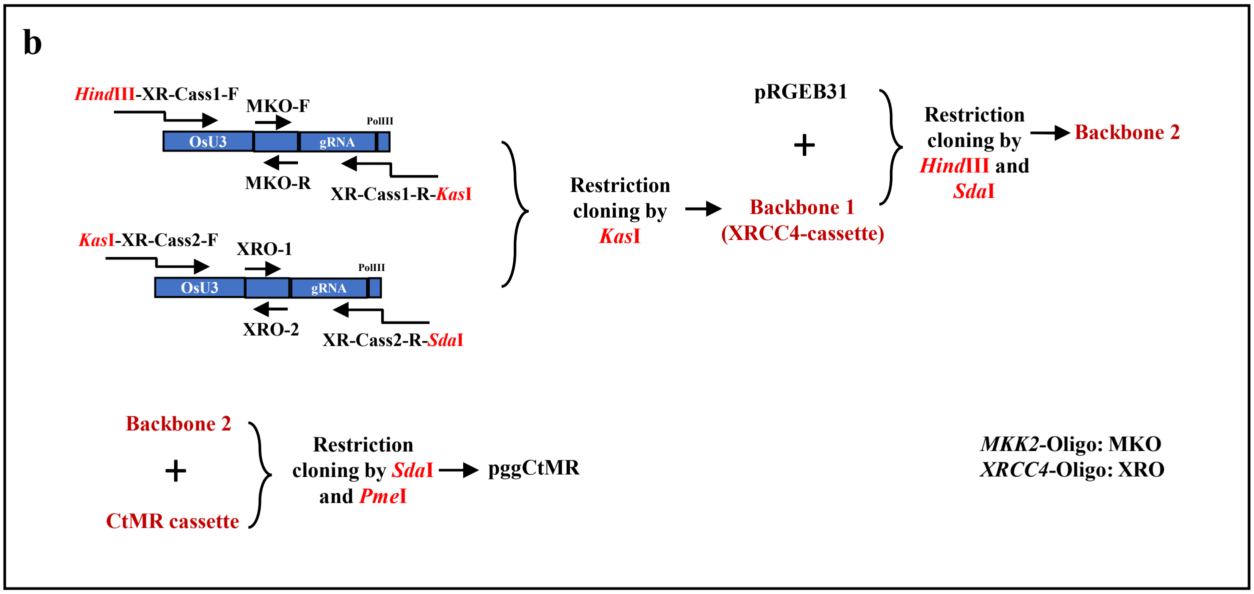


**Supplementary 6:** Schematic construction of cassettes (CtMR, XRCC4) and vectors (pgCtMR and pggCtMR) and their primers.
