## Supplementary material for "Precise exogenous insertion and sequence replacements in poplar by simultaneous HDR overexpression and NHEJ suppression using CRISPR-Cas9": Suppl. 8 (from another version of this preprint: version 6, 2021-02-03)

Supplementary 8: Alignment of events involved in experiment II.

```
Donor (Control)              TTTTGTGAATTGCTAATGGTTTGAAAACCTCAGCCCTGGTCACTGTAATTTCAGCAAGTTGATTTATTTCAAGCCACAATGACA
                             3' homology arm

II#3 (Partial FAM2)          TTTTGTGAATTGCTAATGGTTTGAAAACCTCAGCCCTGGTCACTGTAATTTCAGCAAGTTGATTTATTTCAAGCCACAATGACA
II#7 (FAM1&2)                TTTTGTGAATTGCTAATGGTTTGAAAACCTCAGCCCTGGTCACTGTAATTTCAGCAAGTTGATTTATTTCAAGCCACAATGACA
II#11 (Partial FAM2)         TTTTGTGAATTGCTAATGGTTTGAAAACCTCAGCCCTGGTCACTGTAATTTCAGCAAGTTGATTTATTTCAAGCCACAATGACA
II#13 (Partial FAM1)         TTTTGTGAATTGCTAATGGTTTGAAAACCTCAGCCCTGGTCACTGTAATTTCAGCAAGTTGATTTATTTCAAGCCACAATGACA
II#14 (Partial FAM2)         TTTTGTGAATTGCTAATGGTTTGAAAACCTCAGCCCTGGTCACTGTAATTTCAGCAAGTTGATTTATTTCAAGCCACAATGACA
II#19 (FAM1&2)               TTTTGTGAATTGCTAATGGTTTGAAAACCTCAGCCCTGGTCACTGTAATTTCAGCAAGTTGATTTATTTCAAGCCACAATGACA
II#21 (Partial FAM1)         TTTTGTGAATTGCTAATGGTTTGAAAACCTCAGCCCTGGTCACTGTAATTTCAGCAAGTTGATTTATTTCAAGCCACAATGACA
II#22 (Mutated)              TTTTGTGAATTGCTAATGGTTTGAAAACCTCAGCCCTGGTCACTGTAATTTCAGCAAGTTGATTTATTTCAAGCCACAATGACA
II#23 (Partial FAM2)         TTTTGTGAATTGCTAATGGTTTGAAAACCTCAGCCCTGGTCACTGTAATTTCAGCAAGTTGATTTATTTCAAGCCACAATGACA
II#29 (FAM1&2-Recovered)     TTTTGTGAATTGCT-ATGGTTTGAAAACCTCAGCCCTGGTCACTGTAATTTCAGCAAGTTGATTTATTTCAAGCCACAATGACA
II#35 (Partial FAM1)         TTTTGTGAATTGCTAATGGTTTGAAAACCTCAGCCCTGGTCACTGTAATTTCAGCAAGTTGATTTATTTCAAGCCACAATGACA
II#38 (Mutated)              TTTTGTGAATTGCTAATGGTTTGAAAACCTCAGCCCTGGTCACTGTAATTT-----AGTTGATTTATTTCAAGCCACAATGACA
II#40 (Mutated)              TTTTGTGAATTGCTAATGGTTTGAAAACCTCAGCCCTGGTCACTGTAATTTCAGCAAGTTGATTTATTTCAAGCCACAATGACA
II#41 (Partial FAM1)         TTTTGTGAATTGCTAATGGTTTGAAAACCTCAGCCCTGGTCACTGTAATTTCAGCAAGTTGATTTATTTCAAGCCACAATGACA
II#48 (Wild-type)            TTTTGTGAATTGCTAATGGTTTGAAAACCTCAGCCCTGGTCACTGTAATTTCAGCAAGTTGATTTATTTCAAGCCACAATGACA
II#53 (FAM1&2)               TTTTGTGAATTGCTAATGGTTTGAAAACCTCAGCCCTGGTCACTGTAATTTCAGCAAGTTGATTTATTTCAAGCCACAATGACA
II#59 (FAM1&2)               TTTTGTGAATTGCTAATGGTTTGAAAACCCCAGCCCTGGTCACTGTAATTTCAGCAAGTTGATTTATTTCAAGCCACAATGACA
```

```
Donor (Control)              GTTAAGAATCTTTCTTCAGGCCTT---CAAAATTTC-AATAGAT-AACTTCCCATAACTCTCAAGAGTCTGATATAGTAT---C
                             3' homology arm

II#3 (Partial FAM2)          GTT-AGAATCTTTCTTCAGGCCTT---CAAAATTTC-AATAGAT-AACTTCCCATAACTCTCAAGAGTCTGATATAGTAT---C
II#7 (FAM1&2)                GTTAAGAATCTTTCTTCAGGCCTT---CAAAATTTC-AATAGATCAACTTCCCATAACTCTCAAGAGTCTGATATAGTAT---C
II#11 (Partial FAM2)         GTTAAGAATCTTTCTTCAGGCCTT---CAAAATTTC-AATAGAT-AACTTCCCATAACTCTCAAGAGTCTGATATAGTAT---C
II#13 (Partial FAM1)         GTTAAGAATCTTTCTTCAGGCCTT---CAAAATTTC-AATAGAT-AACTTCCCATAACTCTCAAGAGTCTGATATAGTAT---C
II#14 (Partial FAM2)         GTTAAGAATCTTTCTTCAGGCCTTTTACAAAATTTC-AATAGAT-AACTTCCCATAACTCTCAAGAGTCTGATATAGTAT---C
II#19 (FAM1&2)               GTTAAGAATCTTTCTTCAGGCCTT---CAAAATTTC-AATAGAT-AACTTCCCATAACTCTCAAGAGTCTGATATAGTAT---C
II#21 (Partial FAM1)         GTTAAGAATCTTTCTTCAGGCCTT---CAAAATTTC-AATAGAT-AACTTCCCATAACTCTCAAGAGTCTGATATAGTAT---C
II#22 (Mutated)              GTTAAGAATCTTTCTTCAGGCCTT---CAAAATTTC-AATAGAT-AACTTCCCATAACTCTCAAGAGTCTGATATAGTAT---C
II#23 (Partial FAM2)         GTTAAGAATCTTTCTTCAGGCCTT---CAAAATTTC-AATAGAT-AACTTCCCATAACTCTCAAGAGTCTGATATAGTAT---C
II#29 (FAM1&2-Recovered)     GTTAAGAATCTTTCTTCAGGCCTT---CAAAATTTC-AATAGAT-AACTTCCCATAACTCTCAAGAGTCTGATATAGTAT---C
II#35 (Partial FAM1)         GTTAAGAATCTTTCTTCAGGCCTT---CAAAATTTC-AATAGAT-AACTTCCCATAACTCTCAAGAGTCTGATATAGTAT---C
II#38 (Mutated)              GTTAAGAATCTTTCTTCAGGCCTT---CAAAATTTC-AATAGAT-AACTTCCCATAACTCTCAAGAGTCTGATATAGTATTAAC
II#40 (Mutated)              GTTAAGAATCTTTCTTCAGGCCTT---CAAAATTTC-AATAGAT-AACTTCCCATAACTCTCAAGAGTCTGATATAGTAT---C
II#41 (Partial FAM1)         GTTAAGAATCTTTCTTCAGGCCTT---CAAAATTTC-AATAGAT-AACTTCCCATAACTCTCAAGAGTCTGATATAGTAT---C
II#48 (Wild-type)            GTTAAGAATCTTTCTTCAGGCCTT---CAAAATTTC-AATAGAT-AACTTCCCATAACTCTCAAGAGTCTGATATAGTAT---C
II#53 (FAM1&2)               GTTAAGAATCTTTCTTCAGGCCTT---CAAAATTTCTAATAGAT-AACTTCCCATAACTCTCAAGAGTCTGATATAGTAT---C
II#59 (FAM1&2)               GTTAAGAATCTTTCTTCAGGCCTT---CAAAATTTC-AATAGAT-AACTTCCCATAACTCTCAAGAGTCTGATATAGTAT---C
```

```
Donor (Control)              TGCACTTGTCAAACA-CAAT-GGCTT-AAATTCAACTCTAAAGTA--------C-ATGAATTCTCATAAGAGTTATGGTCGACA
                             3' homology arm

II#3 (Partial FAM2)          TG---TTGTCAAACA-CAAT-GGCTT-AAATTCAACTCTAAAGTA--------C-ATGAATTCTCAAAAGAGTTATGGTCGACA
II#7 (FAM1&2)                TGCACTTGTCAAACA-CAAT-GGCTT-AAATTCAACTCTAAAGTA--------C-ATGAATTCTCATAAGA-TTATGGTCGACA
II#11 (Partial FAM2)         TGCACTTGTCAAACA-CAAT-GGCTT-AAATTCAACTCTAAAGTA--------C-ATGAATTCTCATAAGAGTTATGGTCGACA
II#13 (Partial FAM1)         TGCACTTGTCAAACA-CAAT-GGCTT-AAATTCAACTCTAAAGTA--------C-ATGAATTCTCATAAGAGTTATGGTCGACA
II#14 (Partial FAM2)         TGCACTTGTCAAACA-CAAT-GGCTT--AATTCAACTCTAAAGTA--------C-ATGAATTCTCATAAGAGTTATGGTCGACA
II#19 (FAM1&2)               TGCACTTGTCAAACA-CAAT-GGCTT-AAATTCAACTCTAAAGTA--------C-ATGAATTCTCATAAGAGTTATGGTCGACA
II#21 (Partial FAM1)         TGCACTTGTCAAACA-CAAT-GGCTT-AAATTCAACTCTAAAGTA--------C-ATGAATTCTCATAAGAGTTATGGTCGACA
II#22 (Mutated)              TGCACTTGTCAAACA-CAAT-GGCTT-AAATTCAACTCTAAAGTA--------C-ATGAATTCTCATAAGAGTTATGGTCGACA
II#23 (Partial FAM2)         TGCACTTGTCAAACA-CAAT-GGCTT-AAATTCAACTCTAAAGTA--------C-ATGAATTCTCATAAGAGTTATGGTCGACA
II#29 (FAM1&2-Recovered)     TGCACTTGTCAAACA-CAATCGGCTT-AAATTCAACTCTAAAGTAATTAATCCC-ATGAATTCTCATAAGAGTTATGGTCGACA
II#35 (Partial FAM1)         TGCACTTGTCAAACA-CAAT-GGCTTAAAATTCAACTCTAAAGTA--------C-ATGAATTCTCATAAGAGTTATGGTCGACA
II#38 (Mutated)              TGCACTTGTCAAACA-CAAT-GGCTT-AAATTCAACTCTAAAGTA--------C-ATGAATTCTCATAAGAGTTATGGTCGACA
II#40 (Mutated)              TGCACTTGTCAAACA-CAAT-GGCTT-AAATTCAACTCTAAAGTA--------C-ATGAATTCTCATAAGAGTTATGGTCGACA
II#41 (Partial FAM1)         TGCACTTGTCAAACA-CAAT-GGCTT-AAATTCAACTCTAAAGTA--------C-ATGAATTCTCATAAGAGTTATGGTCGACA
II#48 (Wild-type)            TGCACTTGTCAAACACCAAT-GGCTT-AAATTCAACTCTAAAGTA--------C-ATGAATTCTCATAAGAGTTATCGTCGACA
II#53 (FAM1&2)               TGCACTTGTCAAACA-CAAT-GGCTT-AAATTCAACTCTAAAGTA--------C-ATGAATTCTCATAAGAGTTATGGTCGACA
II#59 (FAM1&2)               TGCACTTGTCAAACA-CAAT-GGCTT-AAATTCAACTCTAAAGTACCCCATAGCGATGAATTCTCATAAGAGTTATGGTCGACA
```

```
Donor (Control)              TGGTAAATGCATTAATTGCATAGACACAGGCGAACTAT----TAAATGATTAATAAGACTTGGATTA-TGCAACTACATCTTCA
                             3' homology arm

II#3 (Partial FAM2)          TGGTAAATGCATTAATTGCATAGACACAGGCGAACTAT----TAAATGATTAATAAGACTTGGATTA-TGCAACTACATCTTCA
II#7 (FAM1&2)                TGGTAAATGCATTAATTGCATAGACACAGGCGAACTAT----TAAATGATTAATAAGACTTGGATTA-TGCAACTACATCTTCA
II#11 (Partial FAM2)         TGGTAAATGCATTAATTGCATAGACACAGGCGAACTAT----TAAATGATTAATAAGACTTGGATTA-TGCAACTACATCTTCA
II#13 (Partial FAM1)         TGGTAAATGCATTAATTGCATAGACACAGGCGAACTAT----TAAATGATTAATAAGACTTGGATT--------TACATCTTCA
II#14 (Partial FAM2)         TGGTAAATGCATTAATTGCATAGACACAGGCGAACTAT----TAAATGATTAATAAGACTTGGATTA-TGCAACTACATCTTCA
II#19 (FAM1&2)               TGGTAAATG-ATTAATTGCATAGACACAGGCGAACTAT----TAAATGATTAATAAGACTTGGATTA-TGCAACTACATCTTCA
II#21 (Partial FAM1)         TGGTAAATGCATTAATTGCATAGACACAGGCGAACTAT----TAAATGATTAATAAGACTTGGATTA-TGCAACTACATCTTCA
II#22 (Mutated)              TGGTAAATGCATTAATTGCATAGACACAGGCGAACTAT----TAAATGATTAATAAGACTTGGATTA-TGCAACTACATCTTCA
II#23 (Partial FAM2)         TGGTAAATGCATTAATTGCATAGACACAGGCGAACTAT----TAAATGATTAATAAGACTTGGATTA-TGCAACTACATCTTCA
II#29 (FAM1&2-Recovered)     TGGTAAATGCATTAATTGCATAGACACAGGCGAACTAT----TAAATGATTAATAAGACTTGGATTATTGCAACTACATCTTCA
II#35 (Partial FAM1)         TGGTAAATGCATTAATTGCATAGACACAGGCGAACTAT----TAAATGATTAATAAGACTTGGATTA-TGCAACTACATCTTCA
II#38 (Mutated)              TGGTAAATGCATTAATTGCATAGACACAGGCGAACTAT----TAAATGATTAATAAGACTTGGATTA-TGCAACTACATCTTCA
II#40 (Mutated)              TGGTAAATGCATTAATTGCATAGAC-CAGGCGAACTAT----TAAATGATTAATAAGACTTGGATTA-TGCAACTACATCTTCA
II#41 (Partial FAM1)         TGGTAAATGCATTAATTGCATAGACACAGGCGAACTAT----TAAATGATTAATAAGACTTGGATTA-TGCAACTACATCTTCA
II#48 (Wild-type)            TGGTAAATGCATTAATTGCATAGACACAGGCGAACTATCCGCTAAATGATTAATAAGACTTGGATTA-TGCAACTACATCTTCA
II#53 (FAM1&2)               TGGTAAATGCATTAATTGCATAGACACAGGCGAACTAT----TAAATGATTAATAAGACTTGGATTA-TGCAACTACATCTTCA
II#59 (FAM1&2)               TGGTAAATGCATTAATTGCATAGACACAGGCGAACTAT----TAAATGATTAATAAGACTTGGATTA-TGCAACTACATCTTCA
```

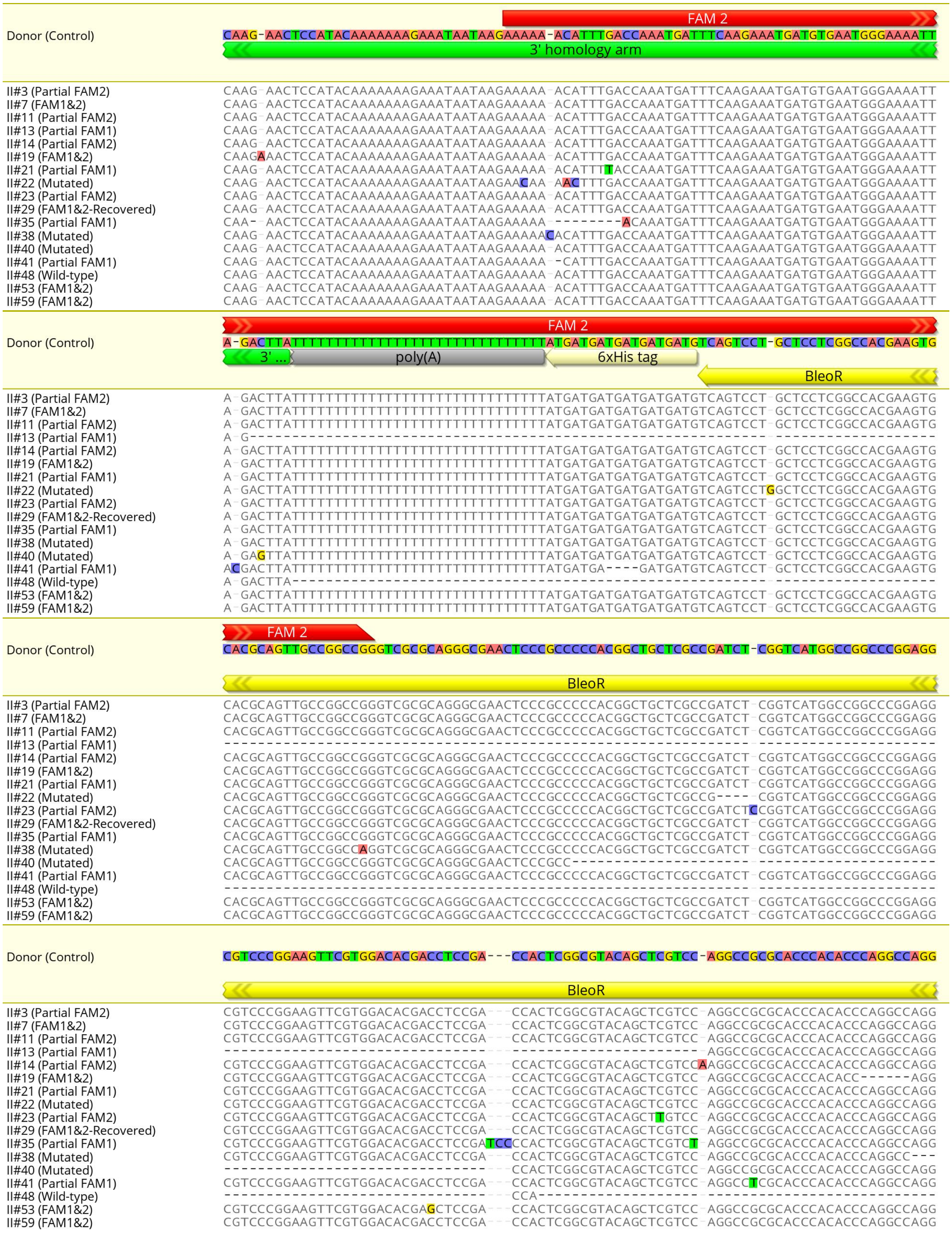
FAM 2
3' homology arm
3' ...
poly(A)
6xHis tag
BleoR
Donor (Control)
II#3 (Partial FAM2)
II#7 (FAM1&2)
II#11 (Partial FAM2)
II#13 (Partial FAM1)
II#14 (Partial FAM2)
II#19 (FAM1&2)
II#21 (Partial FAM1)
II#22 (Mutated)
II#23 (Partial FAM2)
II#29 (FAM1&2-Recovered)
II#35 (Partial FAM1)
II#38 (Mutated)
II#40 (Mutated)
II#41 (Partial FAM1)
II#48 (Wild-type)
II#53 (FAM1&2)
II#59 (FAM1&2)

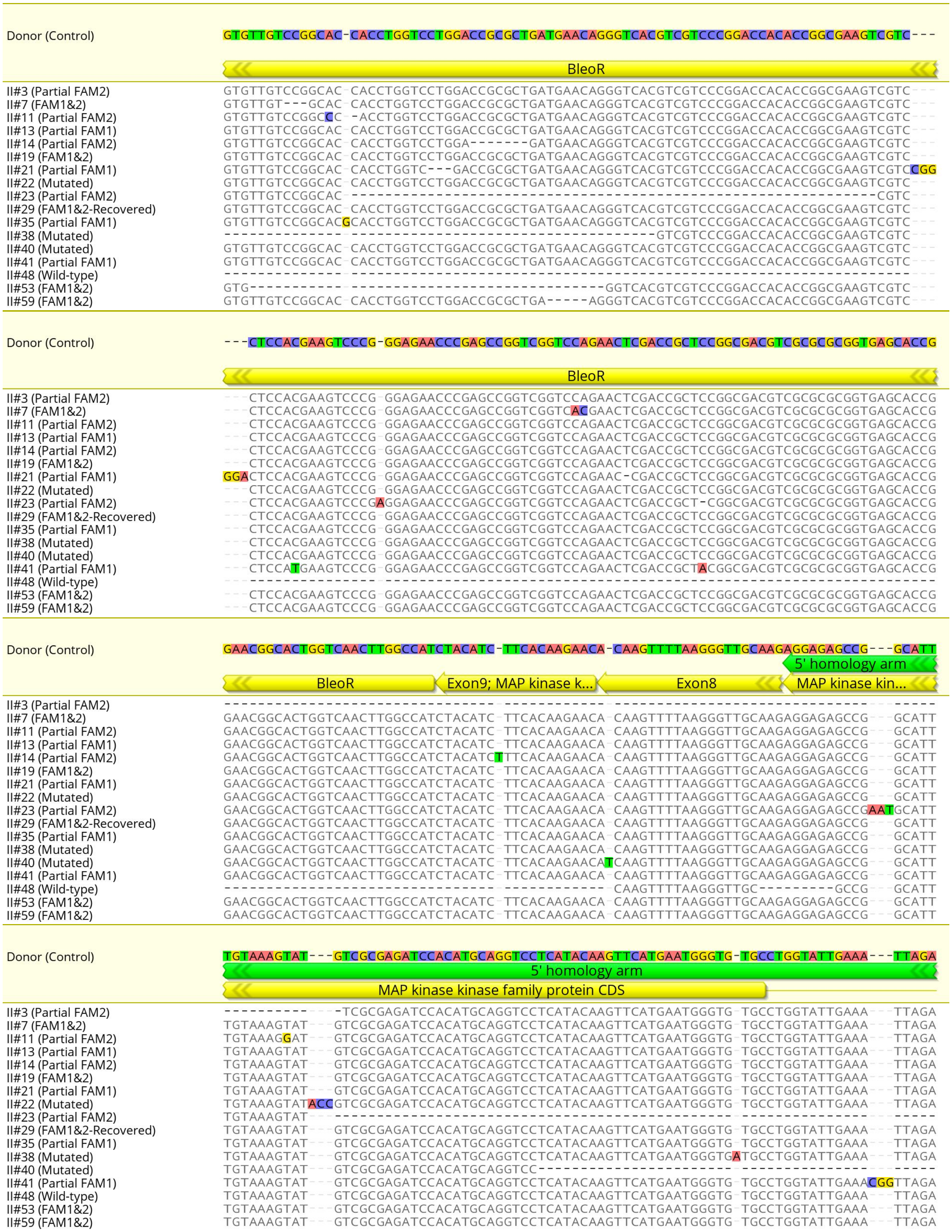
Donor (Control)
BleoR
Exon9; MAP kinase k...
Exon8
MAP kinase kin...
5' homology arm
MAP kinase kinase family protein CDS
II#3 (Partial FAM2)
II#7 (FAM1&2)
II#11 (Partial FAM2)
II#13 (Partial FAM1)
II#14 (Partial FAM2)
II#19 (FAM1&2)
II#21 (Partial FAM1)
II#22 (Mutated)
II#23 (Partial FAM2)
II#29 (FAM1&2-Recovered)
II#35 (Partial FAM1)
II#38 (Mutated)
II#40 (Mutated)
II#41 (Partial FAM1)
II#48 (Wild-type)
II#53 (FAM1&2)
II#59 (FAM1&2)

```
Donor (Control)               AAGAGAAAAA-GAAGAGTCAGA-----TCCATGAGCG-TCCACTAATAGCATACACATGT-TTTGAA-CATATTGAAAAGAGGC
                              5' homology arm

II#3 (Partial FAM2)           AAGAGAAAAA-GAAGAGTCAGA-----TCCATGAGCG-TCCACTAATAGCATACACATGT-TTTGAA-CATATTGAAAAGAGGC
II#7 (FAM1&2)                 AAGAGAAAAA-GAAGAGTCAGA-----TCCATGAGCG-TCCACTAATAGCATACACATGTCTTTGAA-CATATTGAAAAG----
II#11 (Partial FAM2)          AAGAGAAAAACGAAGAGTCAGA-----TCCATGAGCG-TCCACTAATAGC-------TGT-TTTGAA-CATATTGAAAAGAGGC
II#13 (Partial FAM1)          AAGAGAAAAA-GAAGAGTCAGA-----TCCATGAGCG-TCCACTAATAGCATACACATGT-TTTGAA-CATATTGAAAAGAGGC
II#14 (Partial FAM2)          AAGAGAAAAA-GAAGAGTCAGA-----TCCATGAGCG-TCCACTAATAGCATACACATGT-TTTGAA-CATATTGAAAAGAGGC
II#19 (FAM1&2)                AAGAGAAAAA-GAAGAGTCAGAAAGACTCCATGAGCG-TCCACTAATAGCATACACATGT-TTTGAA-CATATTGAAAAGAGGC
II#21 (Partial FAM1)          AAGAGAAAAA-GAAGAGTCAGA-----TCCATGAGCGTTCCACTAATAGCATACACATGT-TTTGAA-CATATTGAAAAGAGGC
II#22 (Mutated)               AAGAGAAAAA-GAAGAGTCAGA-----TCCATGAGCG-TCCACTAATAGCATACACATGT-TTTGAA-CATATTGAAAAGAGGC
II#23 (Partial FAM2)          --------------------------------------T----------------------TTTGAA-CATATTGAAAAGAGGC
II#29 (FAM1&2-Recovered)      AAGAGAAAAA-GAAGAGTCAGA-----TCCATGAGCG-TCCACTAATAGCATACACATGT-TTTGAA-CATATTGAAAAGAGGC
II#35 (Partial FAM1)          AAGAGAAAAA-GAAGAGTCAGA-----TCCATGAGCG-TCCACTAATAGCATACACATGT-TTTGAA-CATATTGAAAAGAGGC
II#38 (Mutated)               AAGAGAAAAA-GAAGAGTCAGA-----TCCATGAGCG-TCCACTAATAGCATACACATGT-TTTGAA-CATATTGAAAAGAGGC
II#40 (Mutated)               ------AAAA-GAAGAGTCAGA-----TCCATGAGCG-TCCACTAATAGCATACACATGT-TTTGAA-CATATTGAAAAGAGGC
II#41 (Partial FAM1)          AAGAGAAAAA-GAAGAGTCAGA-----TCCATGAGCG-TCCACTAATAGCATACACATGT-TTTGAA-CATATTGAAAAGAGGC
II#48 (Wild-type)             AAGAGAAAAA-GAAGAGTCAGA-----TCCATGAGCG-TCC----ATAGCATACACATGT-TTTGAA-CATATTGAAAAGAGGC
II#53 (FAM1&2)                AAGAGAAAAA-GAAGA-------------CATGAGCG-TCCACTAATAGCATACACATGT-TTTGAACCATATTGAAAAGAGGC
II#59 (FAM1&2)                AAGAGAAAAA-GAAGAGTCAGA-----TCCATGAGCG-TCCACTAATAGCATACACATGT-TTTGAA-CATATTGAAAAGAGGC

Donor (Control)               AAAAGAAGGGTGGGA-TATTAGCAGTCTGCTGAG-TATTAATGACATTCCATA-TTGATAGAGAGGAGAAAAGAGTAAAGGACG
                              5' homology arm

II#3 (Partial FAM2)           AAAAGAAGGGTGGGA-TATTAGCAGTCTGCTGAGTTATTAATGACATTCCATA-TTGATAGAGAGGAGAAAAGAGTAAAGGACG
II#7 (FAM1&2)                 --CACAAGGGTGGGA-TATTAGCAGTCTGCTGAG-TATTAATGACATTCCATA-TTGATAGAGAGGAGAAAAGAGTAAAGGACG
II#11 (Partial FAM2)          AAAAGAAGGGTGGGA-TATTAGCAGTCTGCTGAG-TATTAATGACATTCCATA-TTGATAGAGAGGAGAAAAGAGTAAAGGACG
II#13 (Partial FAM1)          AAAAGAAGGGTGGGA-TATTAGCAGTCTGCTGAG-TATTAATGACATTCCATA-TTGATAGAGAGGAGAAAAGAGTAAAGGACG
II#14 (Partial FAM2)          AAAAGAAGGGTGGGA-TATTAGCAGTCTGCTGAG-TATTAATGACATTCCATA-TTGATAGAGAGGAGAAAAGAGTAAAGGACG
II#19 (FAM1&2)                AAAAGAAGGGTGGGA-TATTAGCAGTCTGCTGAG-TATTAATGACATTCCATA-TTGATAGAGAGGAGAAAAGAGTAAAGGACG
II#21 (Partial FAM1)          AAAAGAAGGGTGGGAGTATTAGCAGTCTGCTGAG-TATTAATGACATTCCATA-TTGATAGAGAGGAGAAAAGAGTAAAGGACG
II#22 (Mutated)               AAAAGAAGGGTGGGA-TATTAGCAGTCTGCTGAG-TATTAATGACATTCCATA-TTGATAGAGAGGAGAAAAGGGTAAAGGACG
II#23 (Partial FAM2)          AAAAGAAGGGTGGGA-TATTAGCAGTCTGCTGAG-TATTAATGACATTCCATA-TTGATAGAGAGGAGAAAAGAGTAAAGGACG
II#29 (FAM1&2-Recovered)      AAAAGAAGGGTGGGA-TATTAGCA----GCTGAG-TATTAATGACATTCCATA-TTGATAGAGAGGAGAAAAGAGTAAAGGACG
II#35 (Partial FAM1)          AAAAGAAGGGTGGGA-TATTAGCAGTCTGCTGAG-TATTAATGACATTCCATA-TTGATAGAGAGGAGAAAAGAGTAAAGGACG
II#38 (Mutated)               AAAAGAAGGGTGGGA-TATTAGCAGTCTGCTGAG-TATT-ATGACATTCCATA-TTGATAGAGAGGAGAAAAGAGTAAAGGACG
II#40 (Mutated)               AAAAGAAGGGTGGGA-TATTAGCAGTCTGCTGAG-TATTAATGACATTCCATA-TTGATAGAGAGGAGAAAAGAGTAAAGGACG
II#41 (Partial FAM1)          AAAAGAAGGGTGGGA-TATTAGCAGTCTGCTGAG-TATTAATGACATTCCATA-TTGATAGAGAGGAGAAAAGAGTAAAGGACG
II#48 (Wild-type)             AAAAGAAGGGTGGGA-TATT-GCAGTCTGCTGAG-TATTAATGACATTCCATA-TTGATAGAGAGGAGAAAAGAGTAAAGGACG
II#53 (FAM1&2)                AAAAGAAGGGTGGGA-TATTAGCAGTCTGCTGAG-TATTAATG-CATTCCATA-TTGATAGAGAGGAGAAAAGAGTAAAGGACG
II#59 (FAM1&2)                AAAAGAAGGGTGGGA-TATTAGCAGTCTGCTGAG-TATTAATGACATTCCATATTTGATAGAGAGGAGAAAAGAGTAAAGGACG

Donor (Control)               TG-GAGTGCCAG----TAACAGGCAGACAGCTAATTGCCCCACCCAG--TTTAACAATCATAGGACTTCAAATACCATCAGTTC
                              5' homology arm

II#3 (Partial FAM2)           TG-GAGTGCCAGTGAATAACAGGCAGACAGCTAATTGCCCCACCCAG--TTTAACAATCATAGGACTTCAAATACCATCAGTTC
II#7 (FAM1&2)                 TG-GAGTGCCAG----TAACAGGCAGACAGCTAATTGCCCCACCCAG--TTTAACAATCATAGGACTTCAAATACCATCAGTTC
II#11 (Partial FAM2)          TG-GAGTGCCAG----TAACAGGCAGACAGCTAATTGCCCCACCCAGACTTTAACAATCATAGGACTTCAAATACCATCAGTTC
II#13 (Partial FAM1)          TG-GAGTGCCAG----TAACAGGCAGACAGCTAATTGCCCCACCCAG--TTTAACAATCATAGGACTTCAAATACCATCAGTTC
II#14 (Partial FAM2)          TG-GAGTGCCAG----TAACAGGCAGACAGCTAATTGCCCCACCCAG--TTTAACAATCATAGGACTTCAAATACCATCAGTTC
II#19 (FAM1&2)                TGTGAGTGCCAG----TAACAGGCAGACAGCTAATTGCCCCACCCAG--TTTAACAATCATAGGACTTCAAATACCATCAGTTC
II#21 (Partial FAM1)          TG-GAGTGCCAG----TAACAGGCAGACAGCTAATTGCCCCACCCAG--TTTAACAATCATAGGACTTCAAATACCATCAGTTC
II#22 (Mutated)               TG-GAGTGCCAG----TAACAGGCAGACAGCTAATTGCCCCACCCAG--TTTAACAATCATAGGACTTCAAATACCATCAGTTC
II#23 (Partial FAM2)          TG-GAGTGCCAG----TAACAGGCAGACAGCTAATTGCCCCACCCAG--TTTAACAATCATAGGACTTCAAATACCATCAGTTC
II#29 (FAM1&2-Recovered)      TG-GAGTGCCAG----TAACAGGCAGACAGCTAATTGCCCCACCCAG--TTTAACAATCATAGGACTTCAAATACCATCAGTTC
II#35 (Partial FAM1)          TG-GAGTGCCAG----TAACAGGCAGACAGCTAATTGCCCCACCCAG--TTTAACAATCATAGGACTTCAAATACCATCAGTTC
II#38 (Mutated)               TG-GAGTGCCAG----TAACAGGCAGACAGCTAATTGCCCCACCCAG--TTTAACAATCATAGGACTTCAAATACCATCAGTTC
II#40 (Mutated)               TG-GAGTGCCAG----TAACAGGCAGACAGCTAATTGCCCCACCCAG--TTTAACAATCATAGGACTTCAAATACCATCAGTTC
II#41 (Partial FAM1)          TG-GAGTGCCAG----TAACAGGCAGACAGCTAATTGCCCCACCCAG--TTTAACAATCATAGGACTTCAAATACCATCAGTTC
II#48 (Wild-type)             TG-GAGTGCCAG----TAACAGGCAGACAGCTAATTGCCCCACCCAG--TTTAACAATCATAGGACTTCAAATACCATCAGTTC
II#53 (FAM1&2)                TG-GAGTGCCAG----TAACAGGCAGACAGCTAATTGCCCCACCCAG--TTTAACAATCATAGGACTTCAAATACCATCAGTTC
II#59 (FAM1&2)                TG-GAGTGCCAG----TAACAGGCAGACAGCT-ATTGCCCCACCCAG--TTTAACAATCATAGGACTTCAAATACCATCAGTTC

Donor (Control)               ATG-TGCTGCCTGTCTATCCT-TTGGATCTTT------CTGCAC-ACTGGAAGAAGACAGAACCACAGAATGA--AAATCAGTA
                              5' homology arm
                              MAP kinase kinase family protein CDS

II#3 (Partial FAM2)           ATG-TGCTGCCTGTCTATCCT-TTGGATCTTT------CTGCAC-ACTGGAAGAAGACAGAACCACAGAATGA--AAATCAGTA
II#7 (FAM1&2)                 CTG-TGCTGCCTGTCTATCCT-TTGGATCTTT------CTGCAC-ACTGGAAGAAGACAGAACCACAGAATGA--AAATCAGTA
II#11 (Partial FAM2)          ATG-TGCTGCCTGTCTATCCTATTGGATCTTT------CTGCAC-ACTGGAAGAAGACAGAACCACAGAATGA--AAATCAGTA
II#13 (Partial FAM1)          ATG-TGCTGCCTGTCTATCCT-TTGGATCTT-------CTGCACTACTGGAAGAAGACAGAACCACAGAATGA--AAATCAGTA
II#14 (Partial FAM2)          ATG-TGCTGCCTGTCTATCCT-TTGGATCTTT------CTGCAC-ACTGGAAGAAGACAGAACCACAGAATGA--AAATCAGTA
II#19 (FAM1&2)                ATG-TGCTGCATGTCTATCCT-TTGGATCTCT------CTGCAC-ACTGGAAGAAGACAGAACCACAGAATGA--AAATCAGTA
II#21 (Partial FAM1)          ATG-TGCTGCCTGTCTATCCT-TTGGATCTTT------CTGCAC-ACTGGAAGAAGACAGAACCACAGAATGA--AAATCAGTA
II#22 (Mutated)               ATG-TGCTGCCTGTCTATCCT-TTGGATCTTT------CTGCAC-ACTGGAAGAAGACAGAACCACAGAATGA--AAATCAGTA
II#23 (Partial FAM2)          ATG-TGCTGCCTGTCTATCCT-TTGGATCTTT------CTGCAC-ACTGGAAGAAGACAGAACCACAGAATGA--AAATCAGTA
II#29 (FAM1&2-Recovered)      ATG-TGCTGCCTGTCTATCCT-TTGGATCTTT------CTGCAC-ACTGGAAGAAGACAGAACCACAGAATGA--AAATCAGTA
II#35 (Partial FAM1)          ATG-TGCTGCCTGTCTATCCT-TTGGATCTTT------CTGCAC-ACTGGAAGAAGACAGAACCACAGAATGA--AAATCAGTA
II#38 (Mutated)               ATG-TGCTGCCTGTCTATCCT-TTGGATCTTT------CTGCAC-ACTGGAAGAAGACAGAACCACAGAATGA--AAATCAGTA
II#40 (Mutated)               ATG-TGCTGCCTGTCTATCCT-TTGGATCTTT------CTGCAC-ACTGGAAGAAGACAGAACCACAGAATGA--AAATCAGTA
II#41 (Partial FAM1)          ATG-TGCTGCCTGTCTATCCT-TTGGATCTTT------CTGCAC-ACTGGAAGAAGACAGAACCACAGAATGA--AAATCAGTA
II#48 (Wild-type)             ATGTTGCTGCCTGTCTATCCT-TTGGATCT--------------------AAGAAGACAGAACCACAGAATGAATAAATCAGTA
II#53 (FAM1&2)                ATG-TGCTGCCTGTCTATCCT-TTGGATCTTTCCCACGCTGCAC-ACTGGAAGAAGACAGAACCACAGAATGA--AAATCAGTA
II#59 (FAM1&2)                ATG-TGCTGCCTGAATATCCT-TTGCATCTTT------CTGCAC-ACT----GAAGACAGAACCACAGAATGA--AAATCAGTA
```

| | |
|---|---|
| Donor (Control) | AC |
| II#3 (Partial FAM2) | AC |
| II#7 (FAM1&2) | AC |
| II#11 (Partial FAM2) | AC |
| II#13 (Partial FAM1) | AC |
| II#14 (Partial FAM2) | AC |
| II#19 (FAM1&2) | AC |
| II#21 (Partial FAM1) | AC |
| II#22 (Mutated) | AC |
| II#23 (Partial FAM2) | AC |
| II#29 (FAM1&2-Recovered) | AC |
| II#35 (Partial FAM1) | AC |
| II#38 (Mutated) | AC |
| II#40 (Mutated) | AC |
| II#41 (Partial FAM1) | AC |
| II#48 (Wild-type) | AC |
| II#53 (FAM1&2) | AC |
| II#59 (FAM1&2) | AC |
