## Supplementary material for "Precise exogenous insertion and sequence replacements in poplar by simultaneous HDR overexpression and NHEJ suppression using CRISPR-Cas9": Suppl. 9 (from another version of this preprint: version 6, 2021-02-03)

```
1. Donor     AATATTCACTTTTTGTGAATTGCTA-----ATGGTT-TGAAAACCTCAGCCCTGGTCACTGTAATTTCAGCAAGT
                       3' homology arm

2. III#6     AATATTCACTTTTTGTGAATTGCTAAGCTCATGGTT-TGAAAACCTCAGCCCTGGTCACTGTAATTTCAGCAAGT
3. III#8     AATATTCACTTTTTGTGAATTGCTA-----ATGGTT-TGAAAACCTCAGCCCTGGTCACTGTAATTTCAGCAAGT
4. III#10    AATATTCACTTTTTGTGAATTGCTA-----ATGGTT-TGAAAACCTCAGCCCTGGTCACTGTAATTTCAGCAAGT
5. III#14              TTTTGTGAATTGCTA-----ATGGTT-TGAAAACCTCAGCCCTGGTCACTGTAATTTCAGCAAGT
6. III#17    AATATTCACTTTTTGTGAATTGCTA-----ATGGTTCTGAAAACCTCAGCCCTGGTCACTGTAATTTCAGCAAGT
7. III#21    AATATTCACTTTTTGTGAATTGCTA-----ATGGTT-TGAAAACCTCAGCCCTGGTCACTGTAATTTCAGCAAGT
8. III#23    AATATTCA--------GAATTGCTA-----ATGGTT-TGAAAACCTCAGCCCTGGTCACTGTAATTTCAGCAAGT
9. III#24    AATATTCACTTTTTGTGAATTGCTA-----ATGGTT-TGAAAACCTCAGCCCTGGTCACTGTAATTTCAGCAAGT
10. III#27   AATATTCACTTTTTGTGAATTGCTA-----ATGGTT-TGAAAACCTCAGCCCTGGTCACTGTAATTTCAGCAAGT
11. III#32   AATATTCACTTTTTGTGAATTGCTA-----ATGGTT-TGAAAACCTCAGCCCTGGTCACTGTAATTTCAGCAAGT
12. III#36   AATATTCACTTTTTGTGAATTGCTA-----ATGGTT-TGAAAACCTCAGCCCTGGTCACTGTAATTTCAGCAAGT
13. III#43             TTTTGTGAATTGCTA-----ATGGTT-TGAAAACCTCAGCCCTGGTCACTGTAATTTCAGCAAGT
14. III#45     TATTCACTTTTTGTGAATTGCTA-----ATGGTT-TGAAAACCTCAGCCCTGGTCACTGTAATTTCAGCAAGT
15. III#53   AATATTCACTTTTTGTGAATTGCTA-----ATGGTT-TGAAAACCTCAGCCCTGGTCACTGTAATTTCAGCAAGT
16. III#61   AATATTCACTTTTTGTGAATTGCTA-----ATGGTT-TGAAAACCTCAGCCCTGGTCACTGTAATTTCAGCAAGT

1. Donor     TGATTTATTTCAAGCCACAATGACAGTTAAGAATCTTTCTTCAGGCCTTCAAAATT---------TCAATAGATA
                       3' homology arm

2. III#6     TGATTTATTTCAAGCCACAATGACAGTTAAGAATCTTTCTTCAGGCCTTCAAAATT---------TCAATAGATA
3. III#8     TGATTTATTTCAAGCCACAATGACAGTTAAGAATCTTTCTTCTGGCCTTCAAAATT---------TCAATAGATA
4. III#10    TGATTTATTTCAAGCCACAATGACAGTTAAGAATCTTTCTTCAGGCCTTCAAAATT---------TCAATAGATA
5. III#14    TGATTTATTTCAAGCCACAATGACAGTTAAGAATCTTTCTTCAGGCCTTCAAAATT---------TCAATAGATA
6. III#17    TGATTTATTTCAAGCCACAATGACAGTTAAGAATCTTTCTTCAGGCCTTCAAAATT---------TCAATAGATA
7. III#21    TGATTTATTTCAAGCCACAATGACAGTTAAGAATCTTTCTTCAGGCCTTCAAAATT---------TCAATAGATA
8. III#23    TGATTTATTTCAAGCCACAATGACAGTTAAGAATCTTTCTTCAGGCCTTCAAAATT---------TCAATAGATA
9. III#24    TGATTTATTTCAAGCCACAATGACAGTTAAGAATCTTTCTTCAGGCCTTCAAAATT---------TCAATAGATA
10. III#27   TGATTTATTTCAAGCCACAATGACAGTTAAGAATCTTTCTTCAGGCCTTCAAAATT---------TCAATAGATA
11. III#32   TGATTTATTTCAAGCCACAATGACAGTTAAGAATCTTTCTTCAGGCCTTCAAAATT---------TCAATAGATA
12. III#36   TGATTTATTTCAAGCCACAATGACAGTTAAGAATCTTTCTTCAGGCCTTCAAAATT---------TCAATAGATA
13. III#43   TGATTTATTTCAAGCCACAATGACAGTTAAGAATCTTTCTTCAGGCCTTCAAAATTTCCGTAAGGTCAATAGATA
14. III#45   TGATTTATTTCAAGCCACAATGACAGTTAAGAATCTTTCTTCAGGCCTTCAAAATT---------TCAATAGATA
15. III#53   TGATTTATTTCAAGCCACAATGACAGTTAAGAATCTTTCTTCAGGCCTTCAAAATT---------TCAATAGATA
16. III#61   TGATTTATTTCAAGCCACAATGACAGTTAAGAATCTTTCTTCAGGCCTTCAAAATT---------TCAATAGATA

1. Donor     ACTTCCCATAACTCTCAAGAGTCTGATATAGTATCTGCACTTGTCAAAC-ACAATGGCTTAAATTCAACTCTAAA
                       3' homology arm

2. III#6     ACTTCCCATAACTCTCAAGAGTCTGATATAGTATCTGCACTTGTCAAAC-ACAATGGCTTAAATTCAACTCTAAA
3. III#8     ACTTCCCATAACTCTCAAGAGTCTAGTATAGTATCTGCACTTGTCAAAC-ACAATGGCTTAAATTCAACTCTAAA
4. III#10    ACTTCCCATAACTCTCAAGAGTCTGATATAGTATCTGCACTTGTCAAAC-ACAATGGCTTAAATTCAACTCTAAA
5. III#14    ACTTCCCATAACTCTCAAGAGTCTGATATAGTATCTGCACTTGTCAAAC-ACAATGGCTTAAATTCAACTCTAAA
6. III#17    ACTTCCCATAACTCTCAAGAGTCTGATATAGTATCTGCACTTGTCAAAC-ACAATGGCTTAAATTCAACTCTAAA
7. III#21    ACTTCCCATAACTCTCAAGAGTCTGATATAGTATCTGCACTTGTCAAAC-ACAATGGCTTAAATTCAACTCTAAA
8. III#23    ACTTCCCATAACTCTCAAGAGTCTGATATAGTATCTGCACTTGTCAAAC-ACAATGGCTTAAATTCAACTCTAAA
9. III#24    ACTTCCCATAACTCTCAAGAGTCTGATATAGTATCTGCACTTGTCAAAC-ACAATGGCTTAAATTCAACTCTAAA
10. III#27   ACTTCCCATAACTCTCAAGAGTCTGATATAGTATCTGCACTTGTCAAAC-ACAATGGCTTAAATTCAACTCTAAA
11. III#32   ACTTCCCATAACTCTCAAGAGTCTGATATAGTATCTGCACTTGTCAAAC-ACAATGGCTTAAATTCAACTCTAAA
12. III#36   ACTTCCCATAACTCTCAAGAGTCTGATATAGTATCTGCACTTGTCAAACAACAATGGCTTAAATTCAACTCTAAA
13. III#43   ACTTCCCATAACTCTCAAGAGTCTGATATAGTATCTGCACTTGTCAAAC-ACAATGGCTTAAATTCAACTCTAAA
14. III#45   ACTTCCCATAACTCTCAAGAGTCTGATATAG-ATCTGCACTTGTCAAAC-ACAATGGCTTAAATTCAATCATAAA
15. III#53   ACTTCCCATAACTCTCAAGAGTCTGATATAGTATCTGCACTTGTCAAAC-ACAATGGCTTAAATTCAACTCTAAA
16. III#61   ACTTCCCATAACTCTCAAGAGTCTGATATAGTATCTGCACTTGTCAAAC-ACAATGGC------TCAACTCTAAA

1. Donor     GTACATGAATTCTCA-TAAGAGTTATGGTCGACATG---------GTAAATGCATTAATTGCATAGACACAGGCG
                       3' homology arm

2. III#6     GTACATGAATTCTCA-TAAGAGTTATGGTCGACATG---------GTAAATGCATTAATTGCATAGACACAGGCG
3. III#8     GTACATGAATTCTCA-TAAGAGTTATGGTCGACATG---------GTAAATGCATTAATTGCATAGACACAGGCG
4. III#10    GTACATGAATTCTCA-TAAGAGTTATGGTCGACATG---------GTAAATGCATTAATTGCATAGACACAGGCG
5. III#14    GTACATGAATTCTCA-TAAGAGTTATGGTCGACATG---------GTAAATGCATTAATTGCATAGACACAGGCG
6. III#17    GTACATGAATTCTCA-TAAGAGTTATGGTCGACATG---------GTAAATGCATTAATTGCATAGACACAGGCG
7. III#21    GTACATGAATTCTCACTAAGAGTTATGGTCGACATG---------GTAAATGCATTAATTGCATAGACACAGGCG
8. III#23    GTACATGAATTCTCA-TAAGAGTTATGGTCGACATG---------GTAAATGCATTAATTGCATAGACACAGGCG
9. III#24    GTACATGAATTCTCA-TAAGAGTTATGGTCGACATG---------GTAAATGCATTAATTGCATAGACACAGGCG
10. III#27   GTACATGAATTCTCA-TAAGAGTTATGGTCGACATG---------GTAAATGCATTAATTGCATAGACACAGGCG
11. III#32   GTACATGAATTCTCA-TAAGAGTTATGGTCGACATG---------GTAAATGCATTAATTGCATAGACACAGGCG
12. III#36   GTACATGAATTCTCA-TAAGAGTTATGGTCGACATG---------GTAAATGCATTAATTGCATAGACACAGGCG
13. III#43   GTACATGAATTCTCA-TAAGAGTTATGGTCGACATG---------GTAAATGCATTAATTGCATAGACACAGGCG
14. III#45   GTACATGAATTCTCA-TAAGAGTTATGGTCGACATG---------GTAAATGCATTAATTGCATAGACACAGGCG
15. III#53   GTACATGAATTCTCA-TAAGAGTTATGGTCGACATG---------GTAAATGCATTAATTGCATAGACACAGGCG
16. III#61   GTACATGAATTCTCA-TAAGAGTTATGGTCGACATGGGCGCTTTAGTAAATGCATTAATTGCATAGACACAGGCG
```

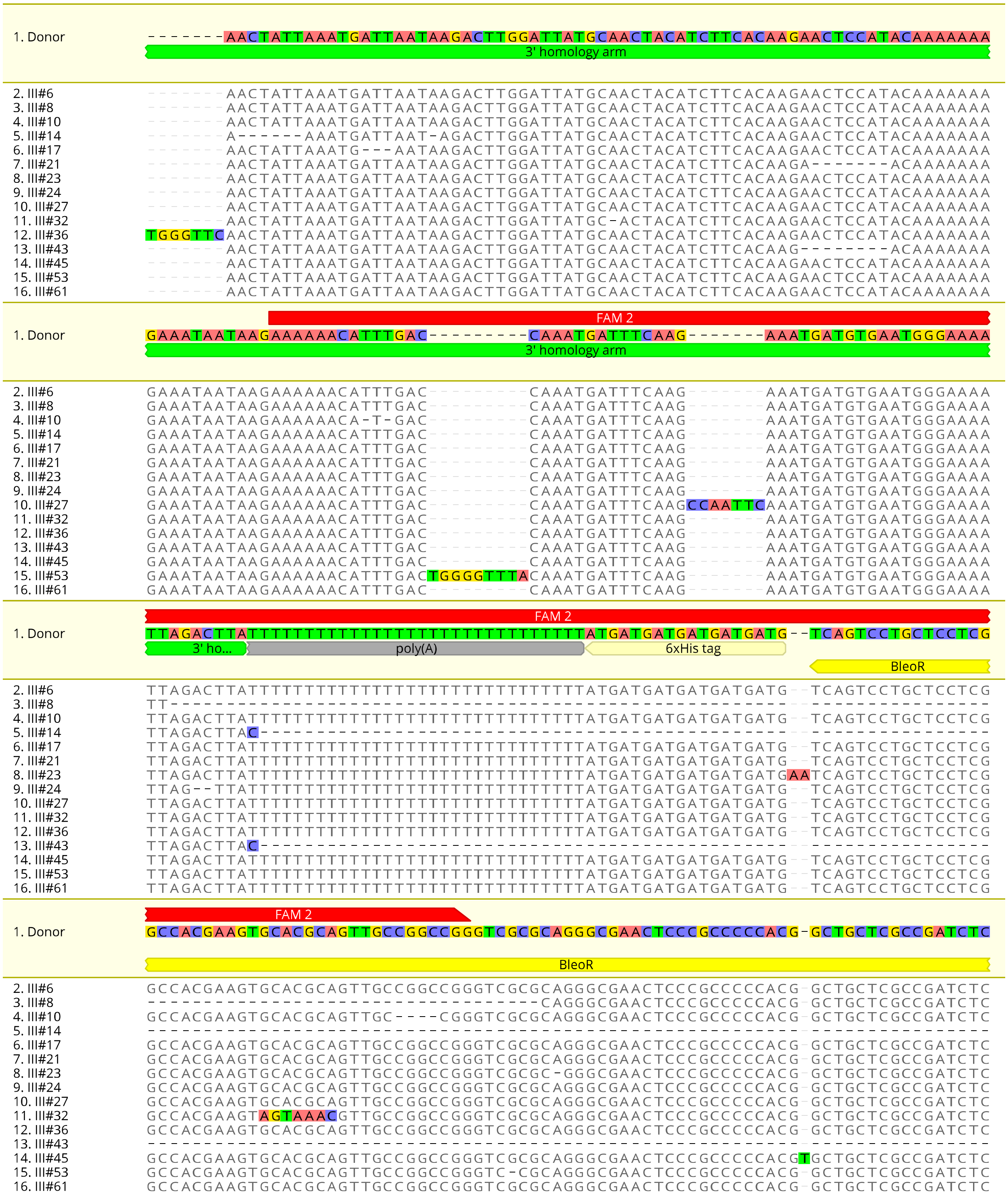
1. Donor
-------AACTATTAAATGATTAATAAGACTTGGATTATGCAACTACATCTTCACAAGAACTCCATACAAAAAAA
3' homology arm
2. III#6 -------AACTATTAAATGATTAATAAGACTTGGATTATGCAACTACATCTTCACAAGAACTCCATACAAAAAAA
3. III#8 -------AACTATTAAATGATTAATAAGACTTGGATTATGCAACTACATCTTCACAAGAACTCCATACAAAAAAA
4. III#10 -------AACTATTAAATGATTAATAAGACTTGGATTATGCAACTACATCTTCACAAGAACTCCATACAAAAAAA
5. III#14 -------A------AAATGATTAAT-AGACTTGGATTATGCAACTACATCTTCACAAGAACTCCATACAAAAAAA
6. III#17 -------AACTATTAAATG---AATAAGACTTGGATTATGCAACTACATCTTCACAAGAACTCCATACAAAAAAA
7. III#21 -------AACTATTAAATGATTAATAAGACTTGGATTATGCAACTACATCTTCACAAGA-------ACAAAAAAA
8. III#23 -------AACTATTAAATGATTAATAAGACTTGGATTATGCAACTACATCTTCACAAGAACTCCATACAAAAAAA
9. III#24 -------AACTATTAAATGATTAATAAGACTTGGATTATGCAACTACATCTTCACAAGAACTCCATACAAAAAAA
10. III#27 -------AACTATTAAATGATTAATAAGACTTGGATTATGCAACTACATCTTCACAAGAACTCCATACAAAAAAA
11. III#32 -------AACTATTAAATGATTAATAAGACTTGGATTATGC-ACTACATCTTCACAAGAACTCCATACAAAAAAA
12. III#36 TGGGTTCAACTATTAAATGATTAATAAGACTTGGATTATGCAACTACATCTTCACAAGAACTCCATACAAAAAAA
13. III#43 -------AACTATTAAATGATTAATAAGACTTGGATTATGCAACTACATCTTCACAAG--------ACAAAAAAA
14. III#45 -------AACTATTAAATGATTAATAAGACTTGGATTATGCAACTACATCTTCACAAGAACTCCATACAAAAAAA
15. III#53 -------AACTATTAAATGATTAATAAGACTTGGATTATGCAACTACATCTTCACAAGAACTCCATACAAAAAAA
16. III#61 -------AACTATTAAATGATTAATAAGACTTGGATTATGCAACTACATCTTCACAAGAACTCCATACAAAAAAA
FAM 2
1. Donor
GAAATAATAAGAAAAAACATTTGAC---------CAAATGATTTCAAG-------AAATGATGTGAATGGAAAA
3' homology arm
2. III#6 GAAATAATAAGAAAAAACATTTGAC---------CAAATGATTTCAAG-------AAATGATGTGAATGGAAAA
3. III#8 GAAATAATAAGAAAAAACATTTGAC---------CAAATGATTTCAAG-------AAATGATGTGAATGGAAAA
4. III#10 GAAATAATAAGAAAAAACA-T-GAC---------CAAATGATTTCAAG-------AAATGATGTGAATGGAAAA
5. III#14 GAAATAATAAGAAAAAACATTTGAC---------CAAATGATTTCAAG-------AAATGATGTGAATGGAAAA
6. III#17 GAAATAATAAGAAAAAACATTTGAC---------CAAATGATTTCAAG-------AAATGATGTGAATGGAAAA
7. III#21 GAAATAATAAGAAAAAACATTTGAC---------CAAATGATTTCAAG-------AAATGATGTGAATGGAAAA
8. III#23 GAAATAATAAGAAAAAACATTTGAC---------CAAATGATTTCAAG-------AAATGATGTGAATGGAAAA
9. III#24 GAAATAATAAGAAAAAACATTTGAC---------CAAATGATTTCAAG-------AAATGATGTGAATGGAAAA
10. III#27 GAAATAATAAGAAAAAACATTTGAC---------CAAATGATTTCAAGCCAATTCAAATGATGTGAATGGAAAA
11. III#32 GAAATAATAAGAAAAAACATTTGAC---------CAAATGATTTCAAG-------AAATGATGTGAATGGAAAA
12. III#36 GAAATAATAAGAAAAAACATTTGAC---------CAAATGATTTCAAG-------AAATGATGTGAATGGAAAA
13. III#43 GAAATAATAAGAAAAAACATTTGAC---------CAAATGATTTCAAG-------AAATGATGTGAATGGAAAA
14. III#45 GAAATAATAAGAAAAAACATTTGAC---------CAAATGATTTCAAG-------AAATGATGTGAATGGAAAA
15. III#53 GAAATAATAAGAAAAAACATTTGACTGGGGTTTACAAATGATTTCAAG-------AAATGATGTGAATGGAAAA
16. III#61 GAAATAATAAGAAAAAACATTTGAC---------CAAATGATTTCAAG-------AAATGATGTGAATGGAAAA
FAM 2
1. Donor
TTAGACTTATTTTTTTTTTTTTTTTTTTTTTTTTTTTTTATGATGATGATGATGATG--TCAGTCCTGCTCCTCG
3' ho...
poly(A)
6xHis tag
BleoR
2. III#6 TTAGACTTATTTTTTTTTTTTTTTTTTTTTTTTTTTTTTATGATGATGATGATGATG--TCAGTCCTGCTCCTCG
3. III#8 TT------------------------------------------------------------------------
4. III#10 TTAGACTTATTTTTTTTTTTTTTTTTTTTTTTTTTTTTTATGATGATGATGATGATG------------------
5. III#14 TTAGACTTAC-----------------------------------------------------------------
6. III#17 TTAGACTTATTTTTTTTTTTTTTTTTTTTTTTTTTTTTTATGATGATGATGATGATG--TCAGTCCTGCTCCTCG
7. III#21 TTAGACTTATTTTTTTTTTTTTTTTTTTTTTTTTTTTTTATGATGATGATGATGATG--TCAGTCCTGCTCCTCG
8. III#23 TTAGACTTATTTTTTTTTTTTTTTTTTTTTTTTTTTTTTATGATGATGATGATGATGAATCAGTCCTGCTCCTCG
9. III#24 TTAG--TTATTTTTTTTTTTTTTTTTTTTTTTTTTTTTTATGATGATGATGATGATG--TCAGTCCTGCTCCTCG
10. III#27 TTAGACTTATTTTTTTTTTTTTTTTTTTTTTTTTTTTTTATGATGATGATGATGATG--TCAGTCCTGCTCCTCG
11. III#32 TTAGACTTATTTTTTTTTTTTTTTTTTTTTTTTTTTTTTATGATGATGATGATGATG--TCAGTCCTGCTCCTCG
12. III#36 TTAGACTTATTTTTTTTTTTTTTTTTTTTTTTTTTTTTTATGATGATGATGATGATG--TCAGTCCTGCTCCTCG
13. III#43 TTAGACTTAC-----------------------------------------------------------------
14. III#45 TTAGACTTATTTTTTTTTTTTTTTTTTTTTTTTTTTTTTATGATGATGATGATGATG--TCAGTCCTGCTCCTCG
15. III#53 TTAGACTTATTTTTTTTTTTTTTTTTTTTTTTTTTTTTTATGATGATGATGATGATG--TCAGTCCTGCTCCTCG
16. III#61 TTAGACTTATTTTTTTTTTTTTTTTTTTTTTTTTTTTTTATGATGATGATGATGATG--TCAGTCCTGCTCCTCG
FAM 2
1. Donor
GCCACGAAGTGCACGCAGTTGCCGGCCGGGTCGCGCAGGGCGAACTCCCGCCCCCACG-GCTGCTCGCCGATCTC
BleoR
2. III#6 GCCACGAAGTGCACGCAGTTGCCGGCCGGGTCGCGCAGGGCGAACTCCCGCCCCCACG-GCTGCTCGCCGATCTC
3. III#8 ------------------------------------CAGGGCGAACTCCCGCCCCCACG-GCTGCTCGCCGATCTC
4. III#10 GCCACGAAGTGCACGCAGTTGC----CGGGTCGCGCAGGGCGAACTCCCGCCCCCACG-GCTGCTCGCCGATCTC
5. III#14 ---------------------------------------------------------------------------
6. III#17 GCCACGAAGTGCACGCAGTTGCCGGCCGGGTCGCGCAGGGCGAACTCCCGCCCCCACG-GCTGCTCGCCGATCTC
7. III#21 GCCACGAAGTGCACGCAGTTGCCGGCCGGGTCGCGCAGGGCGAACTCCCGCCCCCACG-GCTGCTCGCCGATCTC
8. III#23 GCCACGAAGTGCACGCAGTTGCCGGCCGGGTCGCGC-GGGCGAACTCCCGCCCCCACG-GCTGCTCGCCGATCTC
9. III#24 GCCACGAAGTGCACGCAGTTGCCGGCCGGGTCGCGCAGGGCGAACTCCCGCCCCCACG-GCTGCTCGCCGATCTC
10. III#27 GCCACGAAGTGCACGCAGTTGCCGGCCGGGTCGCGCAGGGCGAACTCCCGCCCCCACG-GCTGCTCGCCGATCTC
11. III#32 GCCACGAAGTAGTAAACGTTGCCGGCCGGGTCGCGCAGGGCGAACTCCCGCCCCCACG-GCTGCTCGCCGATCTC
12. III#36 GCCACGAAGTGCACGCAGTTGCCGGCCGGGTCGCGCAGGGCGAACTCCCGCCCCCACG-GCTGCTCGCCGATCTC
13. III#43 ---------------------------------------------------------------------------
14. III#45 GCCACGAAGTGCACGCAGTTGCCGGCCGGGTCGCGCAGGGCGAACTCCCGCCCCCACGTGCTGCTCGCCGATCTC
15. III#53 GCCACGAAGTGCACGCAGTTGCCGGCCGGGTC-CGCAGGGCGAACTCCCGCCCCCACG-GCTGCTCGCCGATCTC
16. III#61 GCCACGAAGTGCACGCAGTTGCCGGCCGGGTCGCGCAGGGCGAACTCCCGCCCCCACG-GCTGCTCGCCGATCTC

```
1. Donor     GGTCATGGCCGGCCCGGAGGCG-TCCCGGAAGTTCGTGGACACGACCTCCGACCACTCGGCGTACAGCTCGTCCA
             BleoR
2. III#6     GGTCATGGCCGGCCCGGAGGCG-TCCCGGAAGTTCGTGGACACGACCTCCGACCACTCGGCGTACAGCTCGTCCA
3. III#8     GGTCATGGCCGGCCCGGAGGCG-TCCCGGAAGTTCGTGGACACGACCTCCGACCACTCGGCGTACAGCTCGTCCA
4. III#10    GGTCATGGCCGGCCCGGAGGCG-TCCCGGAAGTTCGTGGACACGACCTCCGACCACTCGGCGTACAGCTCGTCCA
5. III#14    ---------------------------------------------------------------------------
6. III#17    GGTCATGGCCGGCCCGGAGGCG-TCCCGGAAGTTCGTGGACACGACCTCCGACCACTCGGCGTACAGCTCGTCCA
7. III#21    GGTCATGGCCGGCCCGGAGGCG-TCCCGGAAGTTCGTGGACACGACCTCCGACCACTCGGCGTACAGCTCGTCCA
8. III#23    GGTCATGGCCGGCCCGGAGGCG-TCCCGGAAGTTCGTGGACACGACCTCCGACCACTCGGCGTACAGCTCGTCCA
9. III#24    GGTCATGGCCGGCCCGGAGGCGTTCCCGGAAGTTCGTGGACACGACCTCCGACCACTCGGCGTACAGCTCGTCCA
10. III#27   GGTCATGGCCGGCCCGGAGGCG-TCCCGGAAGTTCGTGGACACGACCTCCGACCACTCGGCGTACAGCTCGTCCA
11. III#32   GGTCATGGCCGGCCCGGAGGCG-TCCCGGAAGTTCGTGGACACGACCTCCGACCACTCGGCGTACAGCTCGTCCA
12. III#36   GGTCATGGCCGGCCCGGAGGCG-TCCCGGAAGTTCGTGGACACGACCTCCGACCACTCGGCGTACAGCTCGTCCA
13. III#43   ---------------------------------------------------------------------------
14. III#45   GGTCATGGCCGGCCCGGAGGCG-TCCCGGAAGTTCGTGGACACGACCTCCGACCACTCGGCGTACAGCTCGTCCA
15. III#53   GGTCATGGCCGGCCCGGAGGCG-TCCCGGAAGTTCGTGGACACGACCTCCGACCACTCGGCGTACAGCTCGTCCA
16. III#61   GGTCATGGCCGGCCCGGAGGCG-TCCCGGAAGTTCGTGGACACGACCTCCGACCACTCGGCGTACAGCTCGTCCA
```

```
1. Donor     GGCCGCGCACCC--------ACACCCAGGCCAGGGTGTTGTC-CGGCACCACCTGGTCCTGGACCGCGC-TGATG
             BleoR
2. III#6     GGCCGCGCACCC--------ACACCCAGGCCAGGGTGTTGTC-CGGCACCACCTGGTCCTGGACCGCGC-TGATG
3. III#8     GGCCGCGCACCC--------ACACCCAGGCCAGGGTGTTGTC-CGGCACCACCTGGTCCTGGACCGCGC-TGATG
4. III#10    GGCCGCGCACCC--------ACACCCAGGCCAGGGTG----------ACCACCTGGTCCTGGACCGCGC-TGATG
5. III#14    ---------------------------------------------------------------------------
6. III#17    GGCCGCGCACCC--------ACACCCAGGCCAGGGTGTTGTC-CGGCACCACCTGGTCCTGGACCGCGC-TGATG
7. III#21    GGCCGCGCACCC--------ACACCCAGGCCAGGGTGTTGTC-CGGCACCACCTGGTCCTGGACCGCGC-TGATG
8. III#23    GGCCGCGCACCC--------ACACCCAGGCCAGGGTGTTGTC-CGGCACCACCTGGTCCTGGACCGCGC-TGATG
9. III#24    GGCCGCGCACCCATGGTACTACACCCAGGCCAGGGTGTTGTC-CGGCACCACCTGGTCCTGGACCGCGC-TGATG
10. III#27   GGCCGCGCACCC--------ACACCCAGGCCAGGGTGTTGTC-CGGCACCACCTGGTCCTGGACCGCGC-TGATG
11. III#32   GGCCGCGCACCC--------ACACCCAGGCCAGGGTGTTGTC-------CGCCTGGTCCTGGACCGCGC-TGATG
12. III#36   GGCCGCGCACCC--------ACACCCAGGCCAGGGTGTTGTC-CGGCACCACCTGGTCCTGGACCGCGC-TGATG
13. III#43   ---------------------------------------------------------------------------
14. III#45   GGCCGCGCACCC--------ACACCCAGGCCAGGGTGTTGTC-CGGCACCACCTGGTCCTGGACCGCGC-TGATG
15. III#53   GGCCGCGCACCC--------ACACCCAGGCCAGGGTGTTGTC-CGGCACCACCTGGTCCTGGACCGCGC-TGATG
16. III#61   GGCCGCGCACCC--------ACACCCAGGCCAGGGTGTTGTC-CGGCACCACCTGGTCCTGGACCGCGCTTGATG
```

```
1. Donor     AACAGGGTCACGTCGTCCCGGACCACACCGG-CGAAGTCGTCCTCCACGAAGTCCCGGGA---------GAACCC
             BleoR
2. III#6     AACAGGGTCACGTCGTCCCGGACCACACCGG-CGAAGTCGTCCTCCACGAAGTCCCGGGA---------GAACCC
3. III#8     AACAGGGTCACGTCGTCCCGGACCACACCGGCCGAAGTCGTCCTCCACGAAGTCCCGGGA---------GAACCC
4. III#10    AACAGGGTCACGTCGTCCCGGACCACACCGG-CGAAGTCGTCCTCCACGAAGTCCCGGGA---------GAACCC
5. III#14    ---------------------------------------------------------------------------
6. III#17    AACAGGGTCACGTCGTCCCGGACCACACCGG-CGAAGTCGTCCTCCACGAAGTCCCGGGA---------GAACCC
7. III#21    AACAGGGTCACGTCGTCCCGGACCACACCGG-CGAAGTCGTCCTCCACGAAGTCCCGGGA---------GAACCC
8. III#23    AACAGGGTCACGTCGTCCCGGACCACACCGG-CGAAGTCGTCCTCCACGAAGTCCCGGGA---------GAACCC
9. III#24    AACAGGGTCACGTCGTCCCGGACCACACCGG-CGAAGTCGTCCTCCACGAAGTCCCGGGA---------GAACCC
10. III#27   AACAGGGTCACGTCGTCCCGGACCACACCGG-CGAAGTCGTCCTCCACGAAGTCCCGGGA---------GAAC--
11. III#32   AACAGGGTCACGTCGTCCCGGACCACACCGG-CGAAGTCGTCCTCCACGAAGTCCCGGGA---------GAACCC
12. III#36   AACAGGGTCACGTCGTCCCGGACCACACCGG-CGAAGTCGTCCTCCACGAAGTCCCGGGA---------GAACCC
13. III#43   ---------------------------------------------------------------------------
14. III#45   AACAGGGTCACGTCGTCCCGGACCACACCGG-CGAAGTCGTCCTCCACGAAGTCCCGGGATTGGAGCTTGAACCC
15. III#53   AACAGGGTCACGTCGTCCCGGACCACACCGG-CGAAGTCGTCCTCCACGAAGTCCCGGGA---------GAACCC
16. III#61   AACAGGGTCACGTCGTCCCGGACCACACCGG-CGAAGTCGTCCTCCACGAAGTCCCGGGA---------GAACCC
```

```
                                                                            FAM 1
1. Donor     GAGCCGGTCGGTCC-----AGAACTCGACCGCTCCGGCGACGTCGCGCGCGGTGAGCACCGGAACGGCACTGGTC
             BleoR
2. III#6     GAGCCGGTCGGTCC-----AGAACTCGACCGCTCCGGCGACGTCGCGCGCGGTGAGCACCGGAACGGCACTGGTC
3. III#8     GAGCCGGTCGGTCC-----AGAACTCGA-CGCTCCGGCGACGTCGCGCGCGGTGAGCACCGGAACGGCACTGGTC
4. III#10    GAGCCGGTCGGTCC-----AGAACTCGACCGCTCCGGCGACGTCGCGCGCGGTGAGCACCGGAACGGCACTGGTC
5. III#14    ---------------------------------------------------------------------------
6. III#17    GAGCCGGTCGGTCC-----AGAACTCGACCGCTCC----------------------------------------
7. III#21    GAGCCGGTCGGTCCACAATAGAACTCGACCGCTCCGGCGACGTCGCGCGCGGTGAGCACCGGAACGGCACTGGTC
8. III#23    GAGCCGGTCGGTCC-----AGAACTCGACCGCTCCGGCGACGTCGCGCGCGGTGAGCACCGGAACGGCACTGGTC
9. III#24    GAGCCGGTCGGTCC-----AGAACTCGACCGCTCCGGCGACGTCGCGCGCGGTGAGCACCGG-ACGGCACTGGTC
10. III#27   --GCCGGTCGGTCC-----AGAACTCGACCGCTCCGGCGACGTCGCGCGCGGTGAGCACCGGAACGGCACTGGTC
11. III#32   GAGCCGGTCGGTCC-----AGAACTCGACCGCTCCGGCGACGTCGCGCGCGGTGAGCACCGGAACGGCACTGGTC
12. III#36   GAGCCGGTCGGTCC-----AGAACTCGACCGCTCCGGCGACGTCGCGCGCGGTGAG-ACCGGAACGGCACTGGTC
13. III#43   ---------------------------------------------------------------------------
14. III#45   GAGCCGGTCGGTCC-----AGAACTCGACCGCTCCGGCGACGTCGCGCGCGGTGAGCACCGGAACGGCACTGGTC
15. III#53   GAGCCGGTCGGTCC-----AGAACTCGACCGCTCCGGCGACGTCGCGCGCGGTGAGCACCGGAACGGCACTGGTC
16. III#61   GAGCCGGTCGGTCC-----AGAACTCGACCGCTCCGGCGACGTCGCGCGCGGTGAGCACCGGAACGGCACTGGTC
```

```
                FAM 1
1. Donor        AACTTGGCCATCTACATCTTCACAAGAACACAAGTTTTAAGGGTTGCAAGAGG-AGAGC-------CGGCATTTG
                5' homology arm
                BleoR   Exon9   Exon8   MAP kinase kinase family protein CDS

2. III#6        AACTTGGCCATCTACATCTTCACAAGAACACAAGTTTTAAGGGTTGCAAGAGG-AGAGC-------CGGCATTTG
3. III#8        AACTTGGCCATCTACATCTTCACAAGAACACAAGTTTTAAGGGTTGCAAGAGG-AGAGC-------CGGCATTTG
4. III#10       AACTTGGCCATCTACATCTTCACAAGAACACAAGTTTTAAGGGTTGCAAGAGG-AGAGC-------CGGCATTTG
5. III#14       ----------------------------CACAAGTTTTAAGGGTTGCAAGAGGCAGAGCTTAGGTACGGCATTTG
6. III#17       ----------------------------------------------GCAAGAGG-AG-GC-------CGGCATTTG
7. III#21       AACTTGGCCATCTACATCTTCACAAGAACACAAGTTTTAAGGGTTGCAAGAGG-AGAGC-------CGGCATTTG
8. III#23       AACTTGGCCATCTACATCTTCACAAGAACACAAGTTTTAAGGGTTGCAAGAGG-AGAGC-------CGGCATTTG
9. III#24       AACTTGGCCATCTACATCTTCACAAGAACACAAGTTTTAAGGGTTGCAAGAGG-AGAGC-------CGGCATTTG
10. III#27      AACTTGGCCATCTACATCTTCACAAGAACACAAGTTTTAAGGGTTGCAAGAGG-AGAGC-------CGGCATTTG
11. III#32      AACTTGGCCATCTACATCTTCACAAGAACACAAGTTTTAAGGGTTGCAAGAGG-AGAGC-------CGGCATTTG
12. III#36      AACTTGGCCATCTACATCTTCACAAGAACACAAGTTTTAAGGGTTGCAAGAGG-AGAGC-------CGGCATTTG
13. III#43      ----------------------------CACAAGTTTTAAGGGTTGCAAGAGG-AGAGC-------CGGCATTTG
14. III#45      AACTTGGCCATCTACATCTTCACAAGAACACAAGTTTTAAGGGTTGCAAGAGG-AGAGC-------CGGCATTTG
15. III#53      AACTTGGCCATCTACATCTTCACAAGAACACAAGTTTTAAGGGTTGCAAGAGG-AGAGC-------CGGCATTTG
16. III#61      AACTTGGCCATCTACATCTTCACAAGAACACAAGTTTTAAGGGTTGCAAGAGG-AGAGC-------CGGCATTTG
```

```
                FAM 1
1. Donor        TAAAGTATGTCGCGAGATCCACATGCAGGTCCTCATACAAGTTCATGAATGGGTGTGCCTGGTATTGAAATTAGA
                5' homology arm
                MAP kinase kinase family protein CDS

2. III#6        TAAAGTATGTCGCGAGATCCACATGCAGG---TCATACAAGTTCCTGAATGGGTGTGCCTGGTATTG---TTAGA
3. III#8        TAAAGTATGTCGCGAGATCCACATGCAGGTCC------------------------------TATTGAAATTAGA
4. III#10       TAAAGTATGTCGCGAGATCCACATGCAGGTCCTCATACAAGTTCATGAATGGGTGTGCCTGGTATTGAAATTAGA
5. III#14       TAAAGTATGTCGCGAGATCCACATGCAGGTCCTCATACAAGTTCATGAATGGGTGTGCCTGGTATTGAAATTAGA
6. III#17       TAAAGTATGTCGCGAGATCCACATGCAGGTCCTCATACAAGTTCATGAATGGGTGTGCCTGGTATTGAAATTAGA
7. III#21       TAAAGTATGTCGCGAGATCCACATGCAGGTCCTCATACAAGTTCATGAATGGGTGTGCCTGGTATTGAAATTAGA
8. III#23       TAAAGTATGTCGCGAGATCCACATGCAGGTCCTCATACAAGTTCATGAATGGGTGTGCCTGGTATTGAAATTAGA
9. III#24       TAAAGTATGTCGCGAGATCCACATGCAGGGCCTCATACAAGACCATGAATGGGTGTGCCTGGTATTGAAATTAGA
10. III#27      TAAAGTATGTCGCGAGATCCACATGCAGGTCCTCATACAAGTTCATGAATGGGTGTGCCTGGTATTGAAATTAGA
11. III#32      TAAAGTATGTCGCGAGATCCACATGCAGGTCCTCATACAAGTTCATGAATGGGTGTGCCTGGTATTGAAATTAGA
12. III#36      TAAAGTATGTCGCGAGATCCACATGCAGGTCCTCATACAAGTTCATGAATGGGTGTGCCTGGTATTGAAATTAGA
13. III#43      TAAAGTATGTCGCGAGATCCACATGCAGGTCCTCATACAAGTTCATGAATGGGTGTGCCTGGTATTGAAATTAGA
14. III#45      TAAAGTATGTCGCGAGATCCACATGCAGGTCCTCATACAAGTTCATGAATGGGTGTGCCTGGTATTGAAATTAGA
15. III#53      TAAAGTATGTCGCGAGATCCACATGCAGGTCCTCATACAAGTTCATGAATGGGTGTGCCTGGTATTGAAATTAGA
16. III#61      TAAAGTATGTCGCGAGATCCACATGCAGGTCCTCATACAAGTTCATGAATGGGTGTGCCTGGTATTGAAATTAGA
```

```
1. Donor        AAGAGAAAAA-GAAGAGTCAGATCCATGAGCGTCCA-CTAATAGCATACACATGTTTTGAACATATT-GAAAAGA
                5' homology arm

2. III#6        AAGAGA-AAA-GAAGAGTCAGATCCATGAGCGTCCA-CTAATAGCATACACATGTTTTGAACATATT-GAAAAGA
3. III#8        AAGAGAAAAA-GAAGAGTCAGATCCATGAGCGTCCA-CTAATAGCATACACATGTTTTGAACATATT-GAAAAGA
4. III#10       AAGAGAAAAA-GAAGAGTCAGATCCATGAGCGTCCA-CTAATAGCATACACATGTTTTGAACATATTCGAAAAGA
5. III#14       AAGAGAAAAA-GAAGAGTCAGATCCATGAGCGTCCA-CTAATAGCATACACATGTTTTGAACATATT-GAAAAGA
6. III#17       AAGAGAAAAA-GAAGAGTCAGATCCATGAGCGTCCA-CTAATAGCATACACATGTTTTGAACATATT-GAAAAGA
7. III#21       AAGAGAAAAA-GAAGAGTCAGATCCATGAGCGTCCA-CTAATAGCATACACATGTTTTGAACATATT-GAAAAGA
8. III#23       AAGAGAAAAA-GAAGAGTCAGATCCATGAGCGTCCA-CTAATAGCATACACATGTTTTGAACATATT-GAAAAGA
9. III#24       AAGAGAAAAA-GAAGAGTCAGATCCATGAGCGTCCA-CTAATAGCATACACATGTTTTGAACATATT-GAAAAGA
10. III#27      AAGAGAAAAA-GAAGAGTCAGATCCATGAGCGTCCA-CTAATAGCATACACATGTTTTGAACATATT-GAAAAGA
11. III#32      AAGAGAAAAA-GAAGAGTCAGATCCATGAGCGTCCA-CTAATAGCATACACATGTTTTGAACATATT-GAAAAGA
12. III#36      AAGAGAAAAA-GAAGAGTCAGATCCATGAGCGTCCA-CTAATAGCATACACATGTTTT----ATATT-GAAAAGA
13. III#43      AAGAGAAAAAGGAAGAGTCAGATCCATGAGCGTCCA-CTAATAGCATACACATGTTTTGAACATATT-GAAAAGA
14. III#45      AAGAGAAAAA-GAAGAGTCAGATCCATGAGC-TCCA-CTAATAGCATACACATGTTTTGAACATATT-GAAAAGA
15. III#53      AAGAGAAAAA-GAAGAGTCAGATCCATGAGCGTCCACCTAATAGCATACACATGTTTTGAACATATT-GAAAAGA
16. III#61      AAGAGAAAAA-GAAGAGTCAGATCCATGAGCGTCCA-CTAATAGCATACACATGTTTTGAACATATT-GAAAAGA
```

```
1. Donor        GGCAAAAGAAGGGTGGGATATTAGCAGTCTGCTGAGTATTAATGACATTCCATATTGATAGAGAGGAGAAAAGAG
                5' homology arm

2. III#6        GGCAAAAGAAGGGTGGGATATTAGCAGTCTGCTGAGTATTAATGACATTCCATATTGATAGAGAGGAGAAAAGAG
3. III#8        GGCAAAAGAAGGGTGGGATATTAGCAGTCTGCTGAGTATTAATGACATTCCATATTGATAGAGAGGAGAAAAGAG
4. III#10       GGCAAAAGAAGGGTGGGATATTAGCAGTCTGCTGAGTATTAATGACATTCCATATTGATAGAGAGGAGAAAAGAG
5. III#14       GGCAAAAGAAGGGTGGGATATTAGCAGTCTGCTGAGTATTAATGACATTCCATATTGATAGAGAGGAGAAAAGAG
6. III#17       GGCAAAAGAAGGGTGGGATATTAGCAGTCTGCTGAGTATTAATGACATTCCATATTGATAGAGAGGAGAAAAGAG
7. III#21       GGCAAAAGAAGGGTGGGATATTAGCAGTCTGCTGAGTATTAATGACATTCCATATTGATAGAGAGGAGAAAAGAG
8. III#23       GGCAAAAGAAGGGTGGGATATTAGCAGTCTGCTGAGTATTAATGACATTCCATATTGATAGAGAGGAGAAAAGAG
9. III#24       GGCAAAAGAAGGGTGGGATATTAGCAGTCTGCTGAGTATTAATGACATTCCATATTGATAGAGAGGAGAAAAGAG
10. III#27      GGCAAAAGAAGGGTGGGATATTAGCAGTCTGCTGAGTATTAATGACATTCCATATTGATAGAGAGGAGAAAAGAG
11. III#32      GGCAAAAGAAGGGTGGGATATTAGCAGTCTGCTGAGTATTAATGACATTCCATATTGATAGAGAGGAGAAAAGAG
12. III#36      GGCAAAAGAAGGGTGGGATATTAGCAGTCTGCTGAGTATTAATGACATTCCATATTGATAGAGAGGAGAAAAGAG
13. III#43      GGCAAAAGAAGGGTGGGATATTAGCAGTCTGCTGAGTATTAATGACATTCCATATTGATAGAGAGGAGAAAAGAG
14. III#45      GGCAAAAGAAGGGTGGGATATTAGCAGTCTGCTGAGTATTAATGACATTCCATATTGATAGAGAGGAGAAAAGAG
15. III#53      GGCAAAAGAAGGGTGGGATATTAGCAGTCTGCTGAGTATTAATGACATTCCATATTGATAGAGAGGAGAAAAGAG
16. III#61      GGCAAAAGAAGGGTGGGATATTAGCAGTCTGCTGAGTATTAATGACATTCCATATTGATAGAGAGGAGAAAAGAG
```

```
1. Donor     TAAAGGA---------CGTGGAGTGCCAGTAACAGGCAGACAGCTAATTGCCCCACCCAGTTTAACAATCATAGG
             5' homology arm

2. III#6     TAAAGGA---------CGTGGAGTGCCAGTAACAGGCAGACAGCTAATTGCCCCACCCAGTTTAACAATCATAGG
3. III#8     TAAAGGA-----------CGGAGTGCCAGTAACAGGCAGACAGCTAATTGCCCCACCCAGTTTAACAATCATAGG
4. III#10    TAAAGGA---------CGTGGAGTGCCAGTAACAGGCAGACAGCTAATTGCCCCACCCAGTTTAACAATCATAGG
5. III#14    TAAAGGA---------CGTGGAGTGCCAGTAACAGGCAGACAGCTAATTGCCCCACCCAGTTTAACAATCATAGG
6. III#17    TAAAGGACGCAGCCAGCGTGGAGTGCCAGTAACAGGCAGACAGCTAATTGCCCCACCCAGTTTAACAATCATAGG
7. III#21    TAAAGGA---------CGT-GAGTGCCAGTAACAGGCAGACAGCTAATTGCCCCACCCAGTTTAACAATCATAGG
8. III#23    TAAAGGA---------CGTGGAGTGCCAGTAACAGGCAGACAGCTAATTGCCCCACCCAGTTTAACAATCATAGG
9. III#24    TAAAGGA---------CGTGGAGTGCCAGTAACAGGCAGACAGCTAATTGCCCCACCCAGTTTAACAATCATAGG
10. III#27   TAAAGGA---------CGTGGAGTGCCAGTAACAGGCAGA-AGCTAATTGCCCCACCCAGTTTAACAATCATAGG
11. III#32   TAAAGGA---------CGTGGAGTGCCAGTAACAGGCAGACAGCTAATTGCCCCACCCAGTTTAACAATCATAGG
12. III#36   TAAAGGA---------CGTGGAGTGCCAGTAACAGGCAGACAGCTAATTGCCCCACCCAGTTTAACAATCATAGG
13. III#43   TAAAGGA---------CGTGGAGTGCCAGTAACAGGCAGACAGCTAATTGCCCCACCCAGTTTAACAATCATAGG
14. III#45   TAAAGGA---------CGTGGAGTGCCAGTAACAGGCAGACAGCTAATTGCCCCACCCAGTTTAACAATCATAGG
15. III#53   TAAAGGA---------CGTGGAGTGCCAGTAACAGGCAGACAGCTAATTGCCCCACCCAGTTTAACAATCATAGG
16. III#61   TAAAGGA---------CGTGGAGTGCCAGTAACAGGCAGACAGCTAATTGCCCCACCCAGTTTAACAATCATAGG
```

```
1. Donor     ACTTCAAATACCATCAGTTCATGTGCTGCCTGTC-T----ATCC--TTTGGATCTTTCTGC-ACACTGGAAGAAG
             5' homology arm
             MAP kinase kinase family protein CDS

2. III#6     ACTTCAAATACCATCAGTTCATGTGCTGCCTGTC-T----ATCC--TTTGGATCTTTCTGC-ACACTGGAAGAAG
3. III#8     ACTTCAAATACCATCAGTTCATGTGCTGCCTGTC-T----ATCCTATTTGGATCTTTCTGC-ACACTGGAAGAAG
4. III#10    ACTTCAAATACCATCAGTTCATGTGCTGCCTGTC-T----ATCC--TTTGGATCTTTCTGC-ACACTGGAAGAAG
5. III#14    ACTTCAAATACCATCAGTTCATGTGCTGCCTGTC-T----ATCC--TTTGGATCTTTCTGC-ACACTGGAAGAAG
6. III#17    ACTTCAAATACCATCAGTTCATGTGCTGCCTGTC-T----ATCC--TTTGGATCTTTCTGC-ACACTGGAAGAAG
7. III#21    ACTTCAAATACCATCAGTTCATGTCGTGCCTGTC-T----ATCC--TTTGGATCTTTCTGC-ACACTGGAAGAAG
8. III#23    ACTTCAAATACCATCAGTTCATGTGCTGCCTGTC-T----ATCC--TTTGGATCTTTCTGC-ACACTGGAAGAAG
9. III#24    ACTTCAAATACCATCAGTTCATGTGCTGCCTGTC-T----ATCC--TTTGGATCTTTCTGC-ACACTGGAAGAAG
10. III#27   ACTTCAAATACCATCAGTTCATGTGCTGCCTGTCAT----ATCC--TTTGGATCTTTCTGC-ACACTGGAAGAAG
11. III#32   ACTTCAAATACCATCAGTTCATGTGCTGCCTGTC-TCTCGATCC--TTTGGATCTTTCTGCTACACTGGAAGAAG
12. III#36   ACTTCAAATACCATCAGTTCATGTGCTGCCTGTC-T----ATCC--TTTGGATCTTTCTGC-ACACTGGAAGAAG
13. III#43   ACTTCAAATACCATCAGTTCATGTGCTGCCTGTC-T----ATCC--TTTGGATCTTTCTGC-ACACTGGAAGAAG
14. III#45   ACTTCAAATACCATCAGTTCATGTGCTGCCTGTC-T----ATCC--TTTGGATCTTTCTGC-ACACTGGAAGAAG
15. III#53   ACTTCAAATACCATCAGTTCATGTGCTGCCTGTC-T----ATCC--TTTGGATCTTTC--------TGGAAGAAG
16. III#61   ACTTCAAATACCATCA----ATGTGCTGCCTGTC-T----ATCC--TTTGGATCTTTCTGC-ACACTGGAAGAAG
```

```
1. Donor     ACAGAACCACAGAATG-AAAATCAGTA--ACTTCACAAAATCATCTGGCC
             5' homology arm

2. III#6     ACAGAACCACAGAATG-AAAATCAGTA--ACTTCACAAAATCATCTGGCC
3. III#8     ACAGAACCACAGAATG-AAAATCAGTA--ACTTCACAAAATCATCTGGCC
4. III#10    ACAGAACCACAGAATG-AAAATCAGTATTACTTCACAAAATCATCTGGCC
5. III#14    ACAGAACCACAGAATG-AAAATCAGTA--AC
6. III#17    ACAGAACCACAGAATG-AAAATCAGTA--ACTTCACAAAATCATCTGGCC
7. III#21    ACAGAACCACAGAATG-AAAATCAGTA--ACTTCACAAAATCATCTGGCC
8. III#23    ACAGAACCACAGAATGTAAAATCAGTA--ACTTCACAAAATCATCTGGCC
9. III#24    ACAGAACCACAGAATG-AAAATCAGTA--ACTTCACAAAATCATCTGGCC
10. III#27   ACAGAACCACAGAATG-AAAATCAGTA--ACTTCACAAAATCATCTGGCC
11. III#32   ACAGAACCACAGAATG-AAAATCAGTA--ACTTCACAAAATCATCTGGCC
12. III#36   ACAGAACCACAGAATG-AAAATCAGTA--ACTTCACAAAATCATCTGGCC
13. III#43   ACAGAACCACAGAATG-AAAATCAGAA----------------------C
14. III#45   ACAGAACCACAGAATG-AAAATCAGTA--ACTTCACAAAATCATCTGGCC
15. III#53   ACAGAACCACAGAATG-AAAATCAGTA--ACTTCACAAAATCATCTGGCC
16. III#61   ACAGAACCACAGAATG-AAAATCAGTA---CTTCACAAAATCATCTGGCC
```
