## Supplementary material for "Precise exogenous insertion and sequence replacements in poplar by simultaneous HDR overexpression and NHEJ suppression using CRISPR-Cas9": Suppl. 14 (from another version of this preprint: version 6, 2021-02-03)

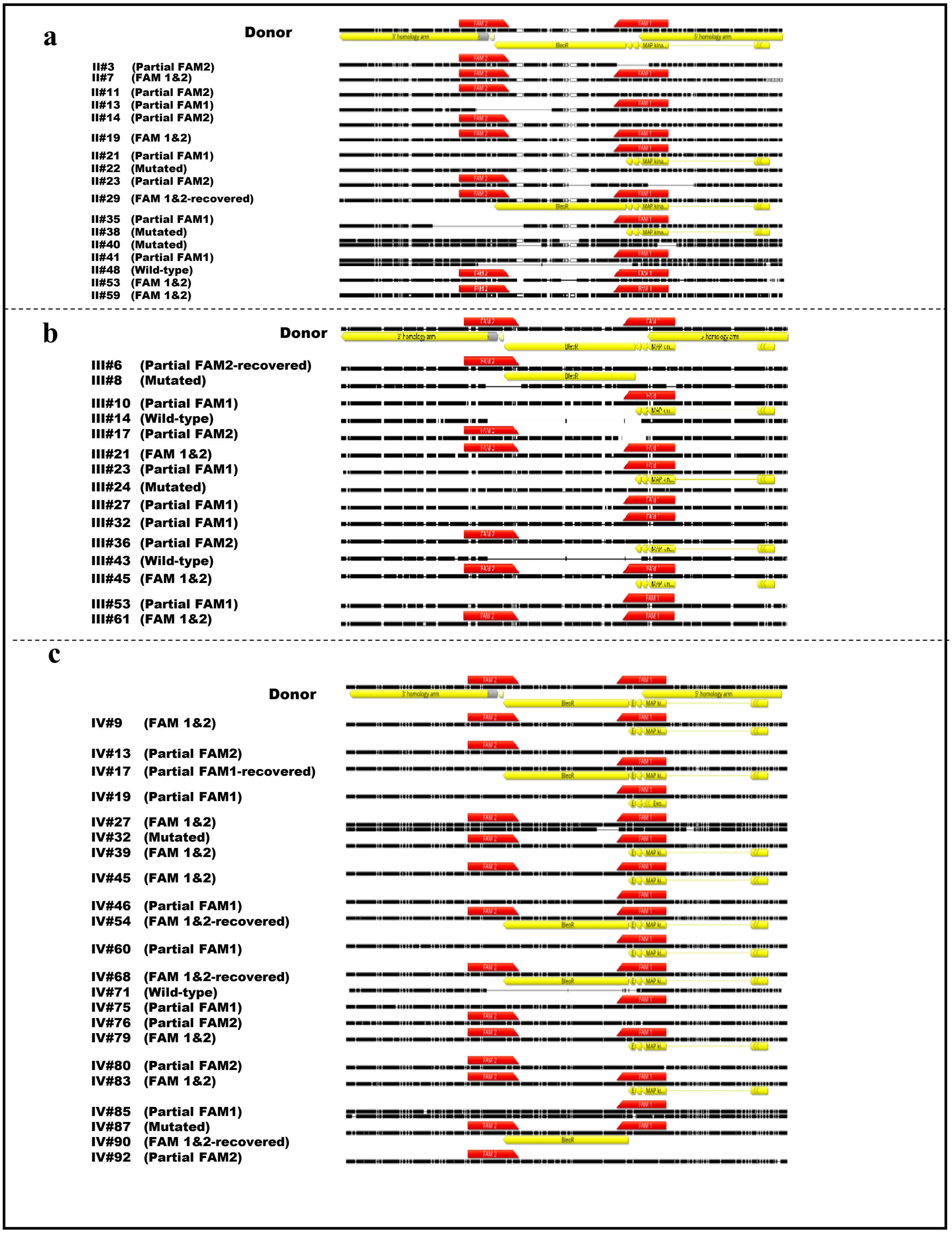
a
Donor
II#3 (Partial FAM2)
II#7 (FAM 1&2)
II#11 (Partial FAM2)
II#13 (Partial FAM1)
II#14 (Partial FAM2)
II#19 (FAM 1&2)
II#21 (Partial FAM1)
II#22 (Mutated)
II#23 (Partial FAM2)
II#29 (FAM 1&2-recovered)
II#35 (Partial FAM1)
II#38 (Mutated)
II#40 (Mutated)
II#41 (Partial FAM1)
II#48 (Wild-type)
II#53 (FAM 1&2)
II#59 (FAM 1&2)
b
Donor
III#6 (Partial FAM2-recovered)
III#8 (Mutated)
III#10 (Partial FAM1)
III#14 (Wild-type)
III#17 (Partial FAM2)
III#21 (FAM 1&2)
III#23 (Partial FAM1)
III#24 (Mutated)
III#27 (Partial FAM1)
III#32 (Partial FAM1)
III#36 (Partial FAM2)
III#43 (Wild-type)
III#45 (FAM 1&2)
III#53 (Partial FAM1)
III#61 (FAM 1&2)
c
Donor
IV#9 (FAM 1&2)
IV#13 (Partial FAM2)
IV#17 (Partial FAM1-recovered)
IV#19 (Partial FAM1)
IV#27 (FAM 1&2)
IV#32 (Mutated)
IV#39 (FAM 1&2)
IV#45 (FAM 1&2)
IV#46 (Partial FAM1)
IV#54 (FAM 1&2-recovered)
IV#60 (Partial FAM1)
IV#68 (FAM 1&2-recovered)
IV#71 (Wild-type)
IV#75 (Partial FAM1)
IV#76 (Partial FAM2)
IV#79 (FAM 1&2)
IV#80 (Partial FAM2)
IV#83 (FAM 1&2)
IV#85 (Partial FAM1)
IV#87 (Mutated)
IV#90 (FAM 1&2-recovered)
IV#92 (Partial FAM2)

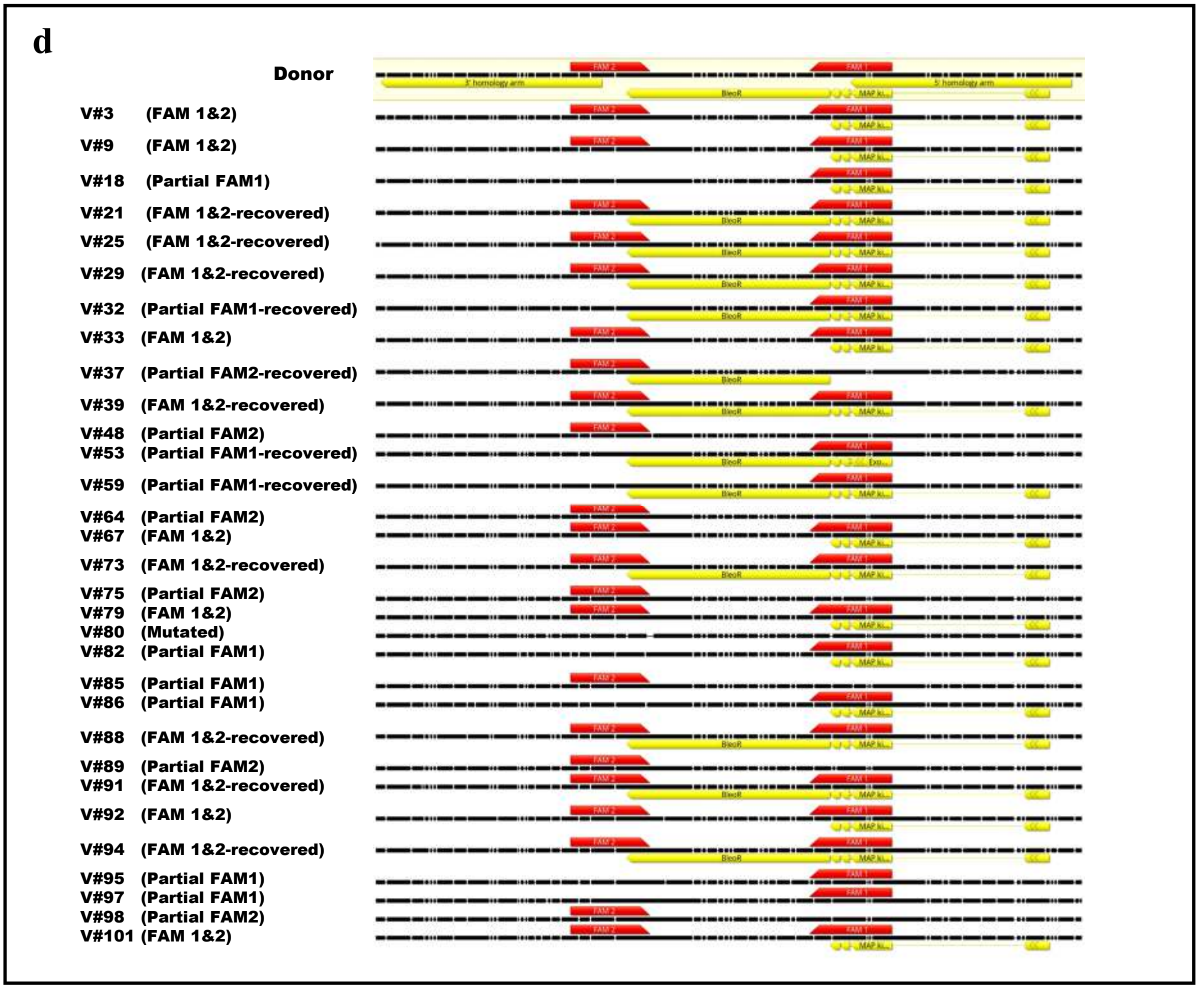


**Supplementary 14:** Schematics of sequence analyzing of triggered events from different experiments. **(a)** Sequence analysis of triggered events included in EXII reveals one recovered event. **(b)** Sequence analysis of triggered events included in EXIII reveals one recovered event. **(c)** Sequence analysis of triggered events included in EXIV reveals four recovered events. **(d)** Sequence analysis of triggered events included in EXV reveals 12 recovered events.
