## Supplementary material for "Precise exogenous insertion and sequence replacements in poplar by simultaneous HDR overexpression and NHEJ suppression using CRISPR-Cas9": Suppl. 15 (from another version of this preprint: version 6, 2021-02-03)

**a**

| | Ex.I | Ex.II | Ex.III | Ex.IV | Ex.V |
|---|---|---|---|---|---|
| **Event-1** | -1.444594 | -0.330585 | 64.255961 | 0.734755 | -0.239312 |
| **Event-2** | -2.643005 | -0.401498 | 1.453354 | 0.702302 | 1.236346 |
| **Event-3** | -0.653519 | -0.991272 | 1.128913 | 77.512145 | 0.998929 |
| **Event-4** | -2.047565 | -0.142852 | -0.355211 | -1.417974 | 91.135438 |
| **Event-5** | -2.257371 | 0.880648 | 0.112011 | 0.833724 | 88.724658 |
| **Event-6** | -0.896275 | -1.345034 | 0.780241 | -0.829714 | 89.924865 |
| **Event-7** | -1.676698 | -0.201047 | 1.739627 | -1.047751 | 88.345632 |
| **Event-8** | -1.370245 | -0.959933 | 0.272871 | -1.164529 | -0.137281 |
| **Event-9** | -1.755111 | -0.136693 | -0.150392 | -0.001808 | 68.708643 |
| **Event-10** | 0.000000 | 57.077383 | 1.303720 | 76.542389 | 89.329409 |
| **Event-11** | 0.000000 | -0.343396 | 2.229397 | 0.119828 | -0.353262 |
| **Event-12** | 0.000000 | -0.211312 | 0.614674 | 76.743422 | 69.129827 |
| **Event-13** | 0.000000 | -0.723973 | 0.214320 | 1.138062 | 88.041569 |
| **Event-14** | 0.000000 | -1.255461 | 1.703323 | -1.805680 | 1.178772 |
| **Event-15** | 0.000000 | 0.255028 | 0.996913 | 0.764758 | -0.400422 |
| **Event-16** | 0.000000 | -0.298613 | 0.000000 | -0.835592 | 87.632063 |
| **Event-17** | 0.000000 | -0.934997 | 0.000000 | -0.669124 | -0.246226 |
| **Event-18** | 0.000000 | 0.000000 | 0.000000 | 0.464890 | -0.231902 |
| **Event-19** | 0.000000 | 0.000000 | 0.000000 | -1.119577 | 0.237968 |
| **Event-20** | 0.000000 | 0.000000 | 0.000000 | -0.468851 | -0.444446 |
| **Event-21** | 0.000000 | 0.000000 | 0.000000 | 55.189797 | -0.603047 |
| **Event-22** | 0.000000 | 0.000000 | 0.000000 | -1.489538 | 1.920526 |
| **Event-23** | 0.000000 | 0.000000 | 0.000000 | 0.000000 | 84.271705 |
| **Event-24** | 0.000000 | 0.000000 | 0.000000 | 0.000000 | 73.310413 |
| **Event-25** | 0.000000 | 0.000000 | 0.000000 | 0.000000 | 88.425699 |
| **Event-26** | 0.000000 | 0.000000 | 0.000000 | 0.000000 | -0.481938 |
| **Event-27** | 0.000000 | 0.000000 | 0.000000 | 0.000000 | 88.284444 |
| **Event-28** | 0.000000 | 0.000000 | 0.000000 | 0.000000 | 1.768907 |
| **Event-29** | 0.000000 | 0.000000 | 0.000000 | 0.000000 | 1.708772 |
| **Event-30** | 0.000000 | 0.000000 | 0.000000 | 0.000000 | 0.377320 |
| **Event-31** | 0.000000 | 0.000000 | 0.000000 | 0.000000 | 0.335972 |

**b**

| | Mean | Standard Deviation | SE of Mean |
|---|---|---|---|
| **Ex.I** | -1.2287 | 0.916710 | 0.264630 |
| **Ex.II** | 4.40787 | 16.595940 | 4.790840 |
| **Ex.III** | 6.11543 | 18.326220 | 5.290320 |
| **Ex.IV** | 19.06057 | 34.907220 | 10.076850 |
| **EX.V** | 48.90032 | 43.543650 | 12.569970 |

**Supplementary 15**: The raw data of real-time PCR to evaluate percentage of delta-delta $C_t$ mean from *BleoR* in the all experiments. **a**, Delta-delta $C_t$ mean of *BleoR* expression from grown buds. Each sample was investigated with three technical repeats. **b**, Descriptive statistic table of raw data calculated by ANOVA-One way.
