## Supplementary material for "Precise exogenous insertion and sequence replacements in poplar by simultaneous HDR overexpression and NHEJ suppression using CRISPR-Cas9": Suppl. 16 (from another version of this preprint: version 6, 2021-02-03)

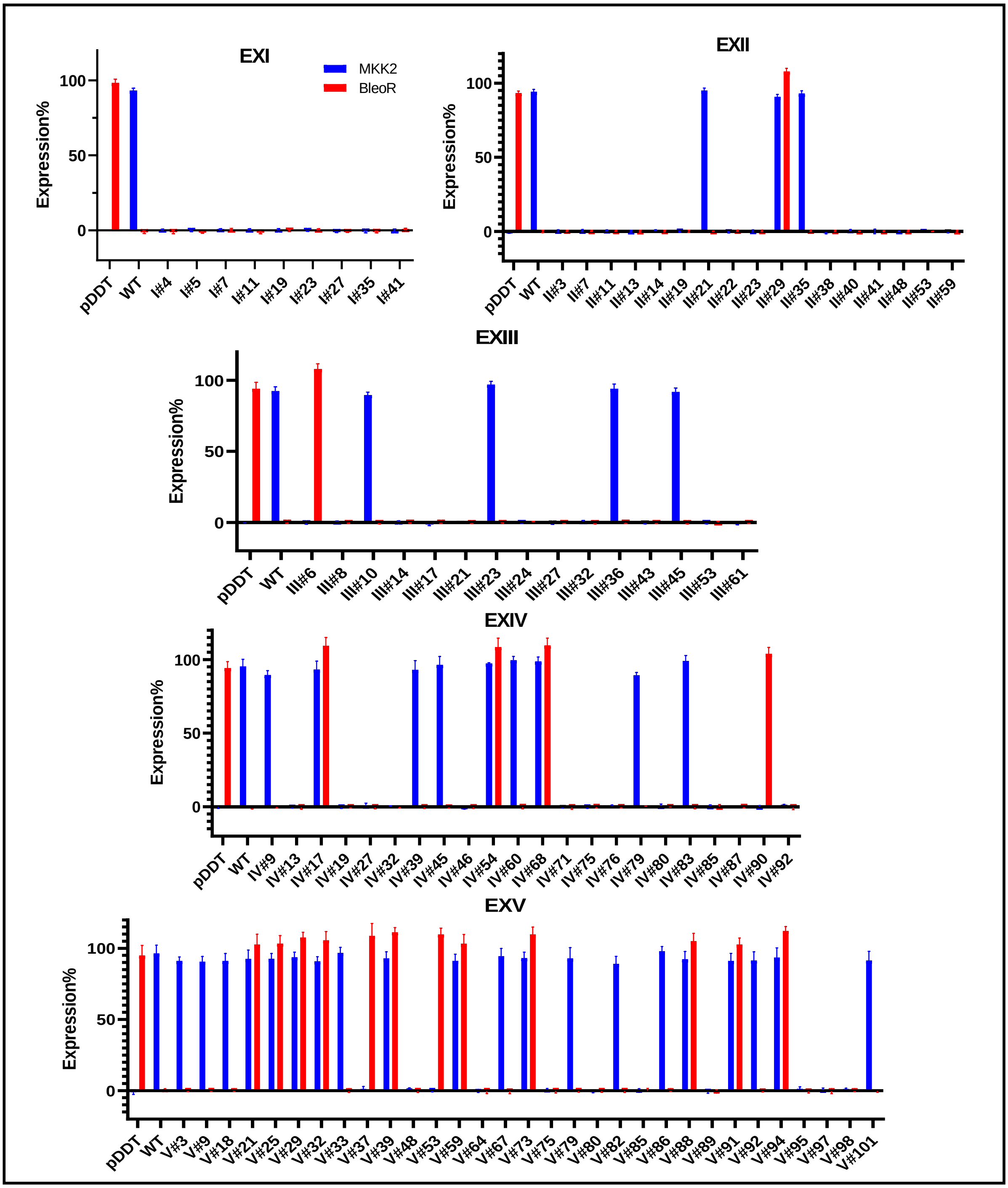


**Supplementary 16:** Column plots of the expression of integrated BleoR and new recombinant MKK2 genes via different designed experiments. Three technical repeats were used for each event in this assay; Error bars represent SD; WT and pDDT were used to control MKK2 and BleoR expression, respectively.
