## Supplementary material for "Precise exogenous insertion and sequence replacements in poplar by simultaneous HDR overexpression and NHEJ suppression using CRISPR-Cas9": Suppl. 17 (from another version of this preprint: version 6, 2021-02-03)

Supplementary 17: Chromatogram alignments of events included in experiment V.

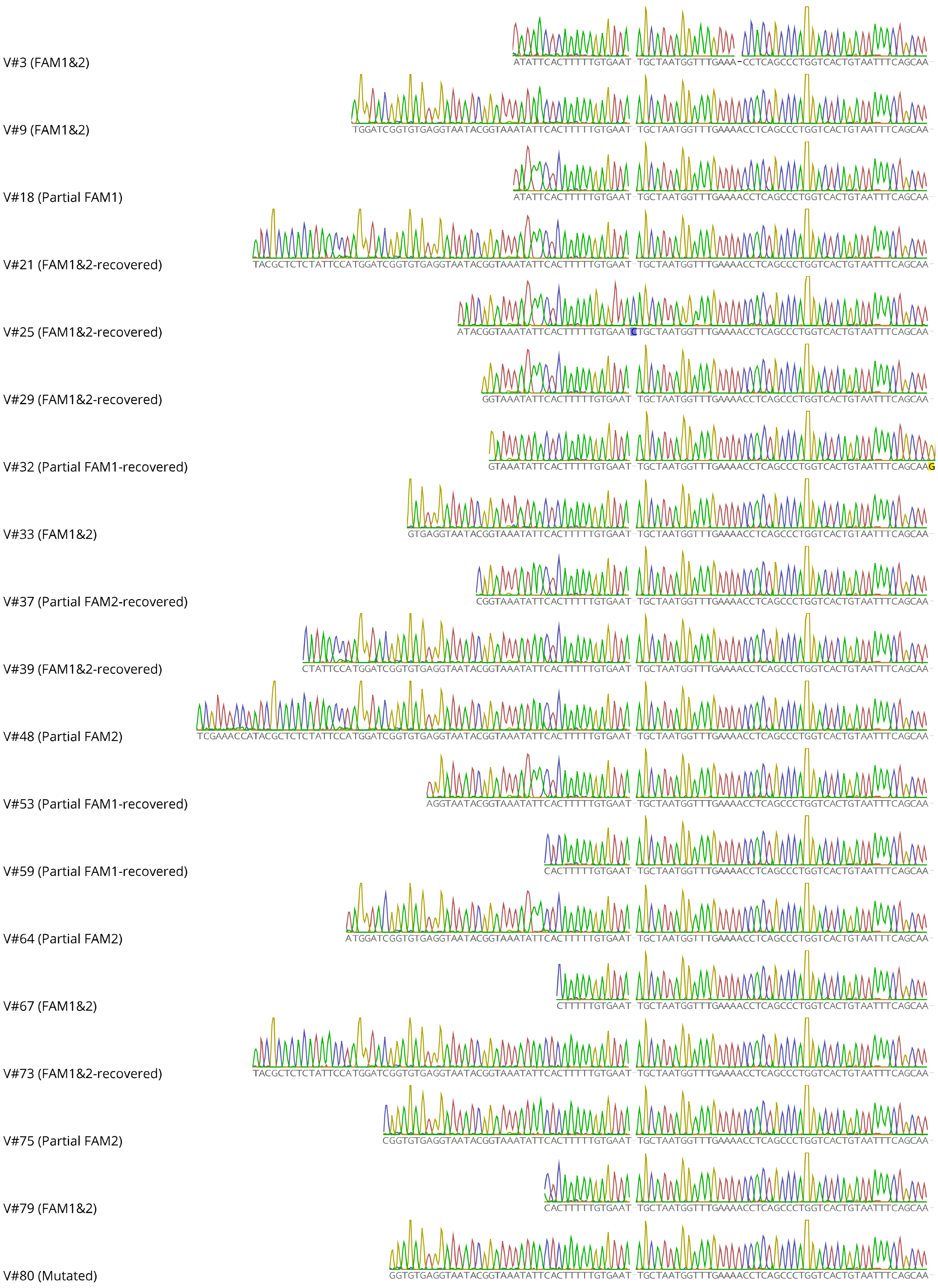

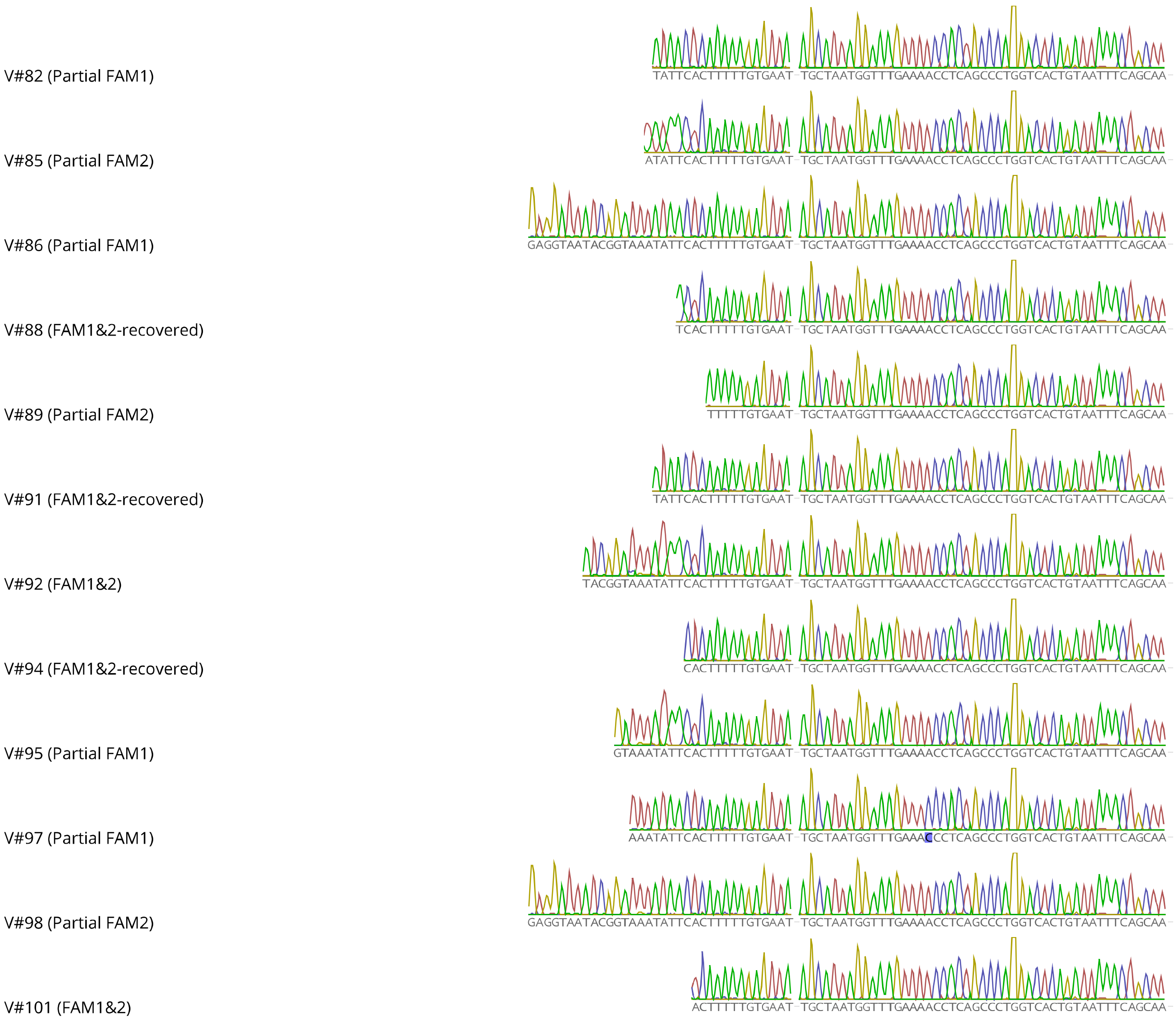
V#82 (Partial FAM1)
TATTCACTTTTTGTGAAT TGCTAATGGTTTGAAAACCTCAGCCCTGGTCACTGTAATTTCAGCAA
V#85 (Partial FAM2)
ATATTCACTTTTTGTGAAT TGCTAATGGTTTGAAAACCTCAGCCCTGGTCACTGTAATTTCAGCAA
V#86 (Partial FAM1)
GAGGTAATACGGTAAATATTCACTTTTTGTGAAT TGCTAATGGTTTGAAAACCTCAGCCCTGGTCACTGTAATTTCAGCAA
V#88 (FAM1&2-recovered)
TCACTTTTTGTGAAT TGCTAATGGTTTGAAAACCTCAGCCCTGGTCACTGTAATTTCAGCAA
V#89 (Partial FAM2)
TTTTTGTGAAT TGCTAATGGTTTGAAAACCTCAGCCCTGGTCACTGTAATTTCAGCAA
V#91 (FAM1&2-recovered)
TATTCACTTTTTGTGAAT TGCTAATGGTTTGAAAACCTCAGCCCTGGTCACTGTAATTTCAGCAA
V#92 (FAM1&2)
TACGGTAAATATTCACTTTTTGTGAAT TGCTAATGGTTTGAAAACCTCAGCCCTGGTCACTGTAATTTCAGCAA
V#94 (FAM1&2-recovered)
CACTTTTTGTGAAT TGCTAATGGTTTGAAAACCTCAGCCCTGGTCACTGTAATTTCAGCAA
V#95 (Partial FAM1)
GTAAATATTCACTTTTTGTGAAT TGCTAATGGTTTGAAAACCTCAGCCCTGGTCACTGTAATTTCAGCAA
V#97 (Partial FAM1)
AAATATTCACTTTTTGTGAAT TGCTAATGGTTTGAAACCCTCAGCCCTGGTCACTGTAATTTCAGCAA
V#98 (Partial FAM2)
GAGGTAATACGGTAAATATTCACTTTTTGTGAAT TGCTAATGGTTTGAAAACCTCAGCCCTGGTCACTGTAATTTCAGCAA
V#101 (FAM1&2)
ACTTTTTGTGAAT TGCTAATGGTTTGAAAACCTCAGCCCTGGTCACTGTAATTTCAGCAA

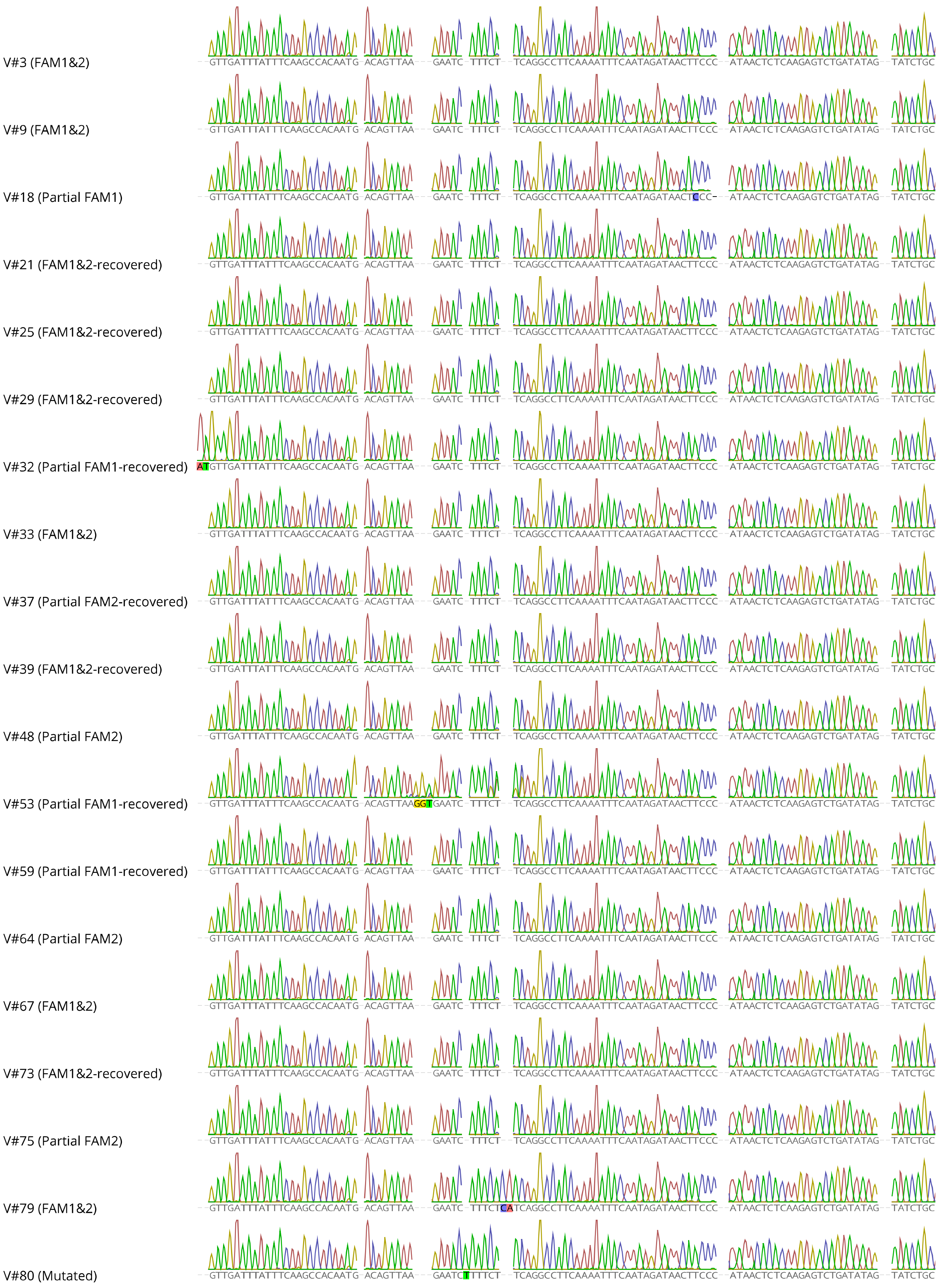
V#3 (FAM1&2)
GTTGATTTATTTCAAGCCACAATG ACAGTTAA GAATC TTTCT TCAGGCCTTCAAAATTTCAATAGATAACTTCCC ATAACTCTCAAGAGTCTGATATAG TATCTGC
V#9 (FAM1&2)
GTTGATTTATTTCAAGCCACAATG ACAGTTAA GAATC TTTCT TCAGGCCTTCAAAATTTCAATAGATAACTTCCC ATAACTCTCAAGAGTCTGATATAG TATCTGC
V#18 (Partial FAM1)
GTTGATTTATTTCAAGCCACAATG ACAGTTAA GAATC TTTCT TCAGGCCTTCAAAATTTCAATAGATAACTCCC- ATAACTCTCAAGAGTCTGATATAG TATCTGC
V#21 (FAM1&2-recovered)
GTTGATTTATTTCAAGCCACAATG ACAGTTAA GAATC TTTCT TCAGGCCTTCAAAATTTCAATAGATAACTTCCC ATAACTCTCAAGAGTCTGATATAG TATCTGC
V#25 (FAM1&2-recovered)
GTTGATTTATTTCAAGCCACAATG ACAGTTAA GAATC TTTCT TCAGGCCTTCAAAATTTCAATAGATAACTTCCC ATAACTCTCAAGAGTCTGATATAG TATCTGC
V#29 (FAM1&2-recovered)
GTTGATTTATTTCAAGCCACAATG ACAGTTAA GAATC TTTCT TCAGGCCTTCAAAATTTCAATAGATAACTTCCC ATAACTCTCAAGAGTCTGATATAG TATCTGC
V#32 (Partial FAM1-recovered)
ATGTTGATTTATTTCAAGCCACAATG ACAGTTAA GAATC TTTCT TCAGGCCTTCAAAATTTCAATAGATAACTTCCC ATAACTCTCAAGAGTCTGATATAG TATCTGC
V#33 (FAM1&2)
GTTGATTTATTTCAAGCCACAATG ACAGTTAA GAATC TTTCT TCAGGCCTTCAAAATTTCAATAGATAACTTCCC ATAACTCTCAAGAGTCTGATATAG TATCTGC
V#37 (Partial FAM2-recovered)
GTTGATTTATTTCAAGCCACAATG ACAGTTAA GAATC TTTCT TCAGGCCTTCAAAATTTCAATAGATAACTTCCC ATAACTCTCAAGAGTCTGATATAG TATCTGC
V#39 (FAM1&2-recovered)
GTTGATTTATTTCAAGCCACAATG ACAGTTAA GAATC TTTCT TCAGGCCTTCAAAATTTCAATAGATAACTTCCC ATAACTCTCAAGAGTCTGATATAG TATCTGC
V#48 (Partial FAM2)
GTTGATTTATTTCAAGCCACAATG ACAGTTAA GAATC TTTCT TCAGGCCTTCAAAATTTCAATAGATAACTTCCC ATAACTCTCAAGAGTCTGATATAG TATCTGC
V#53 (Partial FAM1-recovered)
GTTGATTTATTTCAAGCCACAATG ACAGTTAAGGTGAATC TTTCT TCAGGCCTTCAAAATTTCAATAGATAACTTCCC ATAACTCTCAAGAGTCTGATATAG TATCTGC
V#59 (Partial FAM1-recovered)
GTTGATTTATTTCAAGCCACAATG ACAGTTAA GAATC TTTCT TCAGGCCTTCAAAATTTCAATAGATAACTTCCC ATAACTCTCAAGAGTCTGATATAG TATCTGC
V#64 (Partial FAM2)
GTTGATTTATTTCAAGCCACAATG ACAGTTAA GAATC TTTCT TCAGGCCTTCAAAATTTCAATAGATAACTTCCC ATAACTCTCAAGAGTCTGATATAG TATCTGC
V#67 (FAM1&2)
GTTGATTTATTTCAAGCCACAATG ACAGTTAA GAATC TTTCT TCAGGCCTTCAAAATTTCAATAGATAACTTCCC ATAACTCTCAAGAGTCTGATATAG TATCTGC
V#73 (FAM1&2-recovered)
GTTGATTTATTTCAAGCCACAATG ACAGTTAA GAATC TTTCT TCAGGCCTTCAAAATTTCAATAGATAACTTCCC ATAACTCTCAAGAGTCTGATATAG TATCTGC
V#75 (Partial FAM2)
GTTGATTTATTTCAAGCCACAATG ACAGTTAA GAATC TTTCT TCAGGCCTTCAAAATTTCAATAGATAACTTCCC ATAACTCTCAAGAGTCTGATATAG TATCTGC
V#79 (FAM1&2)
GTTGATTTATTTCAAGCCACAATG ACAGTTAA GAATC TTTCTCATCAGGCCTTCAAAATTTCAATAGATAACTTCCC ATAACTCTCAAGAGTCTGATATAG TATCTGC
V#80 (Mutated)
GTTGATTTATTTCAAGCCACAATG ACAGTTAA GAATCTTTTCT TCAGGCCTTCAAAATTTCAATAGATAACTTCCC ATAACTCTCAAGAGTCTGATATAG TATCTGC

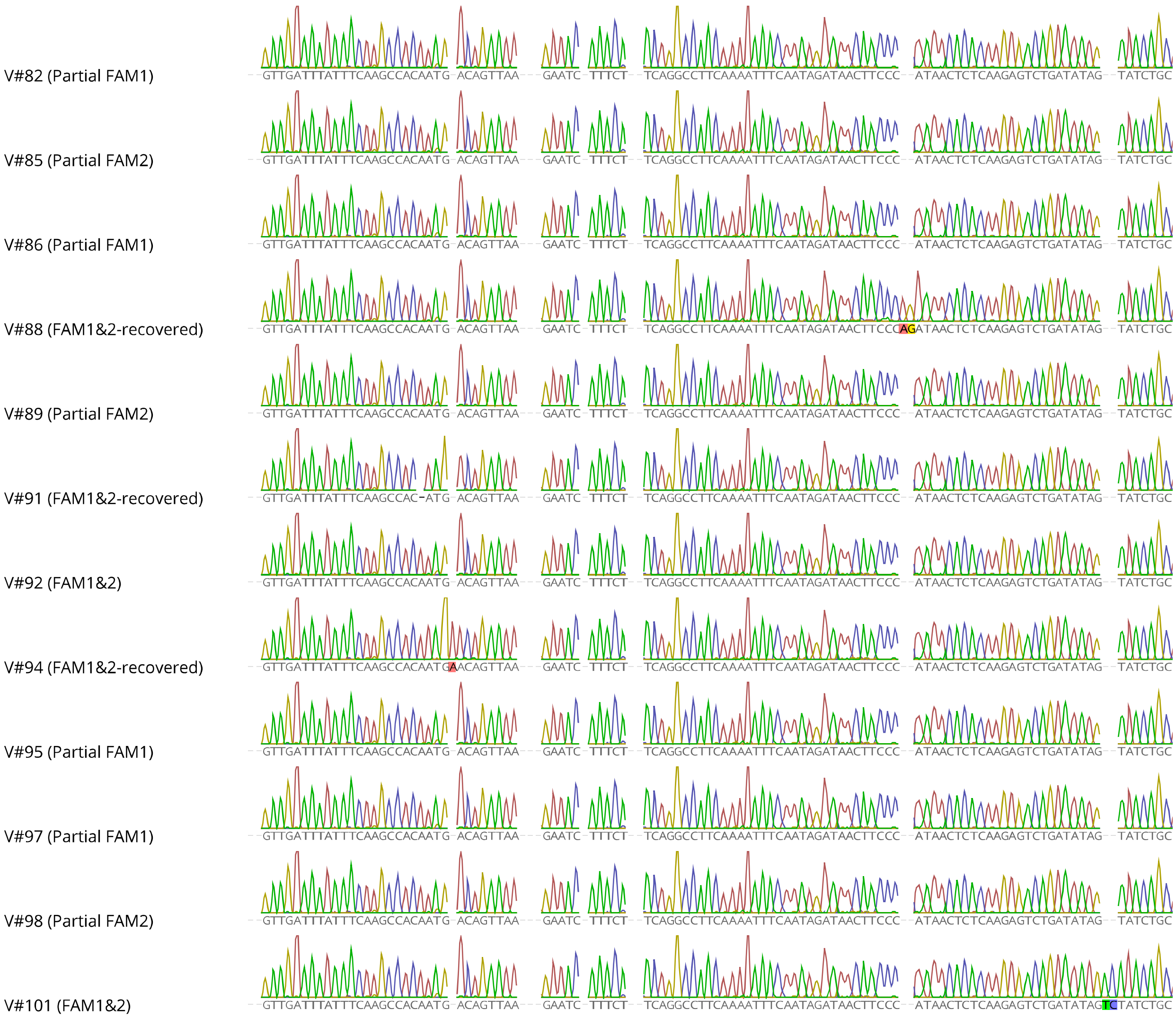
V#82 (Partial FAM1)
GTTGATTTATTTCAAGCCACAATG ACAGTTAA GAATC TTTCT TCAGGCCTTCAAAATTTCAATAGATAACTTCCC ATAACTCTCAAGAGTCTGATATAG TATCTGC
V#85 (Partial FAM2)
GTTGATTTATTTCAAGCCACAATG ACAGTTAA GAATC TTTCT TCAGGCCTTCAAAATTTCAATAGATAACTTCCC ATAACTCTCAAGAGTCTGATATAG TATCTGC
V#86 (Partial FAM1)
GTTGATTTATTTCAAGCCACAATG ACAGTTAA GAATC TTTCT TCAGGCCTTCAAAATTTCAATAGATAACTTCCC ATAACTCTCAAGAGTCTGATATAG TATCTGC
V#88 (FAM1&2-recovered)
GTTGATTTATTTCAAGCCACAATG ACAGTTAA GAATC TTTCT TCAGGCCTTCAAAATTTCAATAGATAACTTCCCAGATAACTCTCAAGAGTCTGATATAG TATCTGC
V#89 (Partial FAM2)
GTTGATTTATTTCAAGCCACAATG ACAGTTAA GAATC TTTCT TCAGGCCTTCAAAATTTCAATAGATAACTTCCC ATAACTCTCAAGAGTCTGATATAG TATCTGC
V#91 (FAM1&2-recovered)
GTTGATTTATTTCAAGCCAC-ATG ACAGTTAA GAATC TTTCT TCAGGCCTTCAAAATTTCAATAGATAACTTCCC ATAACTCTCAAGAGTCTGATATAG TATCTGC
V#92 (FAM1&2)
GTTGATTTATTTCAAGCCACAATG ACAGTTAA GAATC TTTCT TCAGGCCTTCAAAATTTCAATAGATAACTTCCC ATAACTCTCAAGAGTCTGATATAG TATCTGC
V#94 (FAM1&2-recovered)
GTTGATTTATTTCAAGCCACAATGAACAGTTAA GAATC TTTCT TCAGGCCTTCAAAATTTCAATAGATAACTTCCC ATAACTCTCAAGAGTCTGATATAG TATCTGC
V#95 (Partial FAM1)
GTTGATTTATTTCAAGCCACAATG ACAGTTAA GAATC TTTCT TCAGGCCTTCAAAATTTCAATAGATAACTTCCC ATAACTCTCAAGAGTCTGATATAG TATCTGC
V#97 (Partial FAM1)
GTTGATTTATTTCAAGCCACAATG ACAGTTAA GAATC TTTCT TCAGGCCTTCAAAATTTCAATAGATAACTTCCC ATAACTCTCAAGAGTCTGATATAG TATCTGC
V#98 (Partial FAM2)
GTTGATTTATTTCAAGCCACAATG ACAGTTAA GAATC TTTCT TCAGGCCTTCAAAATTTCAATAGATAACTTCCC ATAACTCTCAAGAGTCTGATATAG TATCTGC
V#101 (FAM1&2)
GTTGATTTATTTCAAGCCACAATG ACAGTTAA GAATC TTTCT TCAGGCCTTCAAAATTTCAATAGATAACTTCCC ATAACTCTCAAGAGTCTGATATAGTCTATCTGC

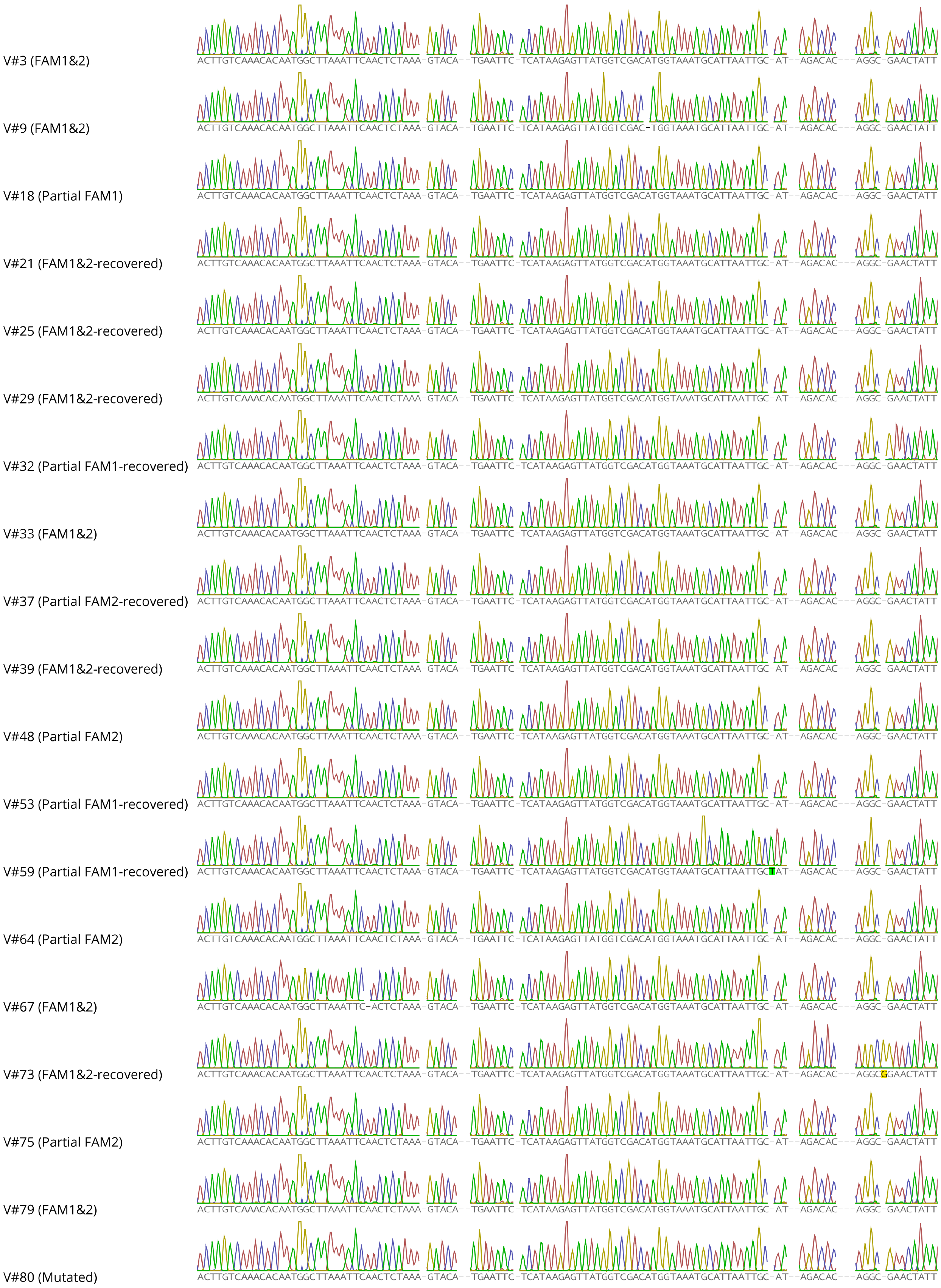
V#3 (FAM1&2)
V#9 (FAM1&2)
V#18 (Partial FAM1)
V#21 (FAM1&2-recovered)
V#25 (FAM1&2-recovered)
V#29 (FAM1&2-recovered)
V#32 (Partial FAM1-recovered)
V#33 (FAM1&2)
V#37 (Partial FAM2-recovered)
V#39 (FAM1&2-recovered)
V#48 (Partial FAM2)
V#53 (Partial FAM1-recovered)
V#59 (Partial FAM1-recovered)
V#64 (Partial FAM2)
V#67 (FAM1&2)
V#73 (FAM1&2-recovered)
V#75 (Partial FAM2)
V#79 (FAM1&2)
V#80 (Mutated)

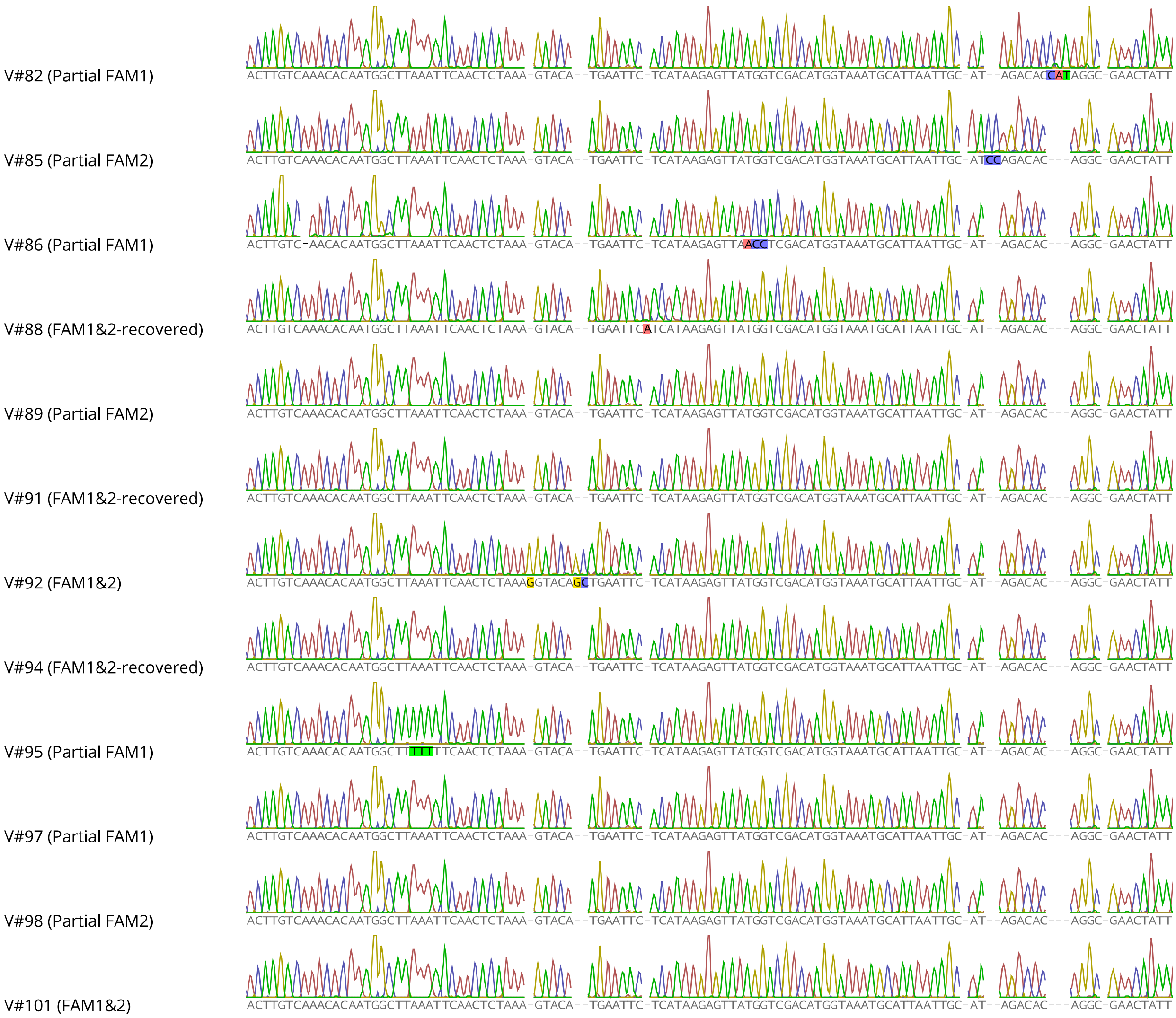
V#82 (Partial FAM1)
V#85 (Partial FAM2)
V#86 (Partial FAM1)
V#88 (FAM1&2-recovered)
V#89 (Partial FAM2)
V#91 (FAM1&2-recovered)
V#92 (FAM1&2)
V#94 (FAM1&2-recovered)
V#95 (Partial FAM1)
V#97 (Partial FAM1)
V#98 (Partial FAM2)
V#101 (FAM1&2)

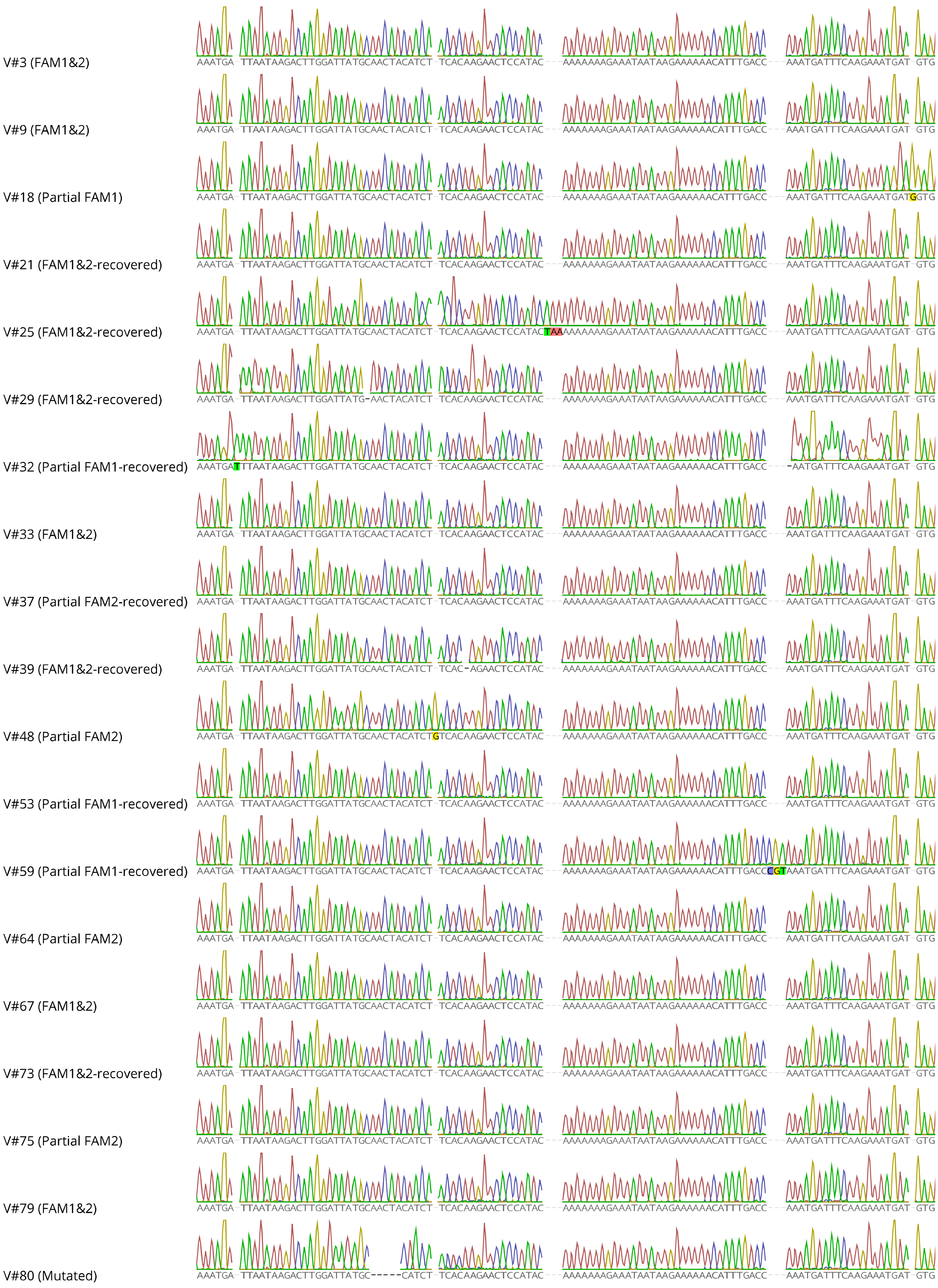

V#3 (FAM1&2)
AAATGA TTAATAAGACTTGGATTATGCAACTACATCT TCACAAGAACTCCATAC AAAAAAAGAAATAATAAGAAAAAACATTTGACC AAATGATTTCAAGAAATGAT GTG
V#9 (FAM1&2)
AAATGA TTAATAAGACTTGGATTATGCAACTACATCT TCACAAGAACTCCATAC AAAAAAAGAAATAATAAGAAAAAACATTTGACC AAATGATTTCAAGAAATGAT GTG
V#18 (Partial FAM1)
AAATGA TTAATAAGACTTGGATTATGCAACTACATCT TCACAAGAACTCCATAC AAAAAAAGAAATAATAAGAAAAAACATTTGACC AAATGATTTCAAGAAATGATGGTG
V#21 (FAM1&2-recovered)
AAATGA TTAATAAGACTTGGATTATGCAACTACATCT TCACAAGAACTCCATAC AAAAAAAGAAATAATAAGAAAAAACATTTGACC AAATGATTTCAAGAAATGAT GTG
V#25 (FAM1&2-recovered)
AAATGA TTAATAAGACTTGGATTATGCAACTACATCT TCACAAGAACTCCATACTAAAAAAAAGAAATAATAAGAAAAAACATTTGACC AAATGATTTCAAGAAATGAT GTG
V#29 (FAM1&2-recovered)
AAATGA TTAATAAGACTTGGATTATG-AACTACATCT TCACAAGAACTCCATAC AAAAAAAGAAATAATAAGAAAAAACATTTGACC AAATGATTTCAAGAAATGAT GTG
V#32 (Partial FAM1-recovered)
AAATGATTTAATAAGACTTGGATTATGCAACTACATCT TCACAAGAACTCCATAC AAAAAAAGAAATAATAAGAAAAAACATTTGACC -AATGATTTCAAGAAATGAT GTG
V#33 (FAM1&2)
AAATGA TTAATAAGACTTGGATTATGCAACTACATCT TCACAAGAACTCCATAC AAAAAAAGAAATAATAAGAAAAAACATTTGACC AAATGATTTCAAGAAATGAT GTG
V#37 (Partial FAM2-recovered)
AAATGA TTAATAAGACTTGGATTATGCAACTACATCT TCACAAGAACTCCATAC AAAAAAAGAAATAATAAGAAAAAACATTTGACC AAATGATTTCAAGAAATGAT GTG
V#39 (FAM1&2-recovered)
AAATGA TTAATAAGACTTGGATTATGCAACTACATCT TCAC-AGAACTCCATAC AAAAAAAGAAATAATAAGAAAAAACATTTGACC AAATGATTTCAAGAAATGAT GTG
V#48 (Partial FAM2)
AAATGA TTAATAAGACTTGGATTATGCAACTACATCTGTCACAAGAACTCCATAC AAAAAAAGAAATAATAAGAAAAAACATTTGACC AAATGATTTCAAGAAATGAT GTG
V#53 (Partial FAM1-recovered)
AAATGA TTAATAAGACTTGGATTATGCAACTACATCT TCACAAGAACTCCATAC AAAAAAAGAAATAATAAGAAAAAACATTTGACC AAATGATTTCAAGAAATGAT GTG
V#59 (Partial FAM1-recovered)
AAATGA TTAATAAGACTTGGATTATGCAACTACATCT TCACAAGAACTCCATAC AAAAAAAGAAATAATAAGAAAAAACATTTGACCCGTAAATGATTTCAAGAAATGAT GTG
V#64 (Partial FAM2)
AAATGA TTAATAAGACTTGGATTATGCAACTACATCT TCACAAGAACTCCATAC AAAAAAAGAAATAATAAGAAAAAACATTTGACC AAATGATTTCAAGAAATGAT GTG
V#67 (FAM1&2)
AAATGA TTAATAAGACTTGGATTATGCAACTACATCT TCACAAGAACTCCATAC AAAAAAAGAAATAATAAGAAAAAACATTTGACC AAATGATTTCAAGAAATGAT GTG
V#73 (FAM1&2-recovered)
AAATGA TTAATAAGACTTGGATTATGCAACTACATCT TCACAAGAACTCCATAC AAAAAAAGAAATAATAAGAAAAAACATTTGACC AAATGATTTCAAGAAATGAT GTG
V#75 (Partial FAM2)
AAATGA TTAATAAGACTTGGATTATGCAACTACATCT TCACAAGAACTCCATAC AAAAAAAGAAATAATAAGAAAAAACATTTGACC AAATGATTTCAAGAAATGAT GTG
V#79 (FAM1&2)
AAATGA TTAATAAGACTTGGATTATGCAACTACATCT TCACAAGAACTCCATAC AAAAAAAGAAATAATAAGAAAAAACATTTGACC AAATGATTTCAAGAAATGAT GTG
V#80 (Mutated)
AAATGA TTAATAAGACTTGGATTATGC-----CATCT TCACAAGAACTCCATAC AAAAAAAGAAATAATAAGAAAAAACATTTGACC AAATGATTTCAAGAAATGAT GTG

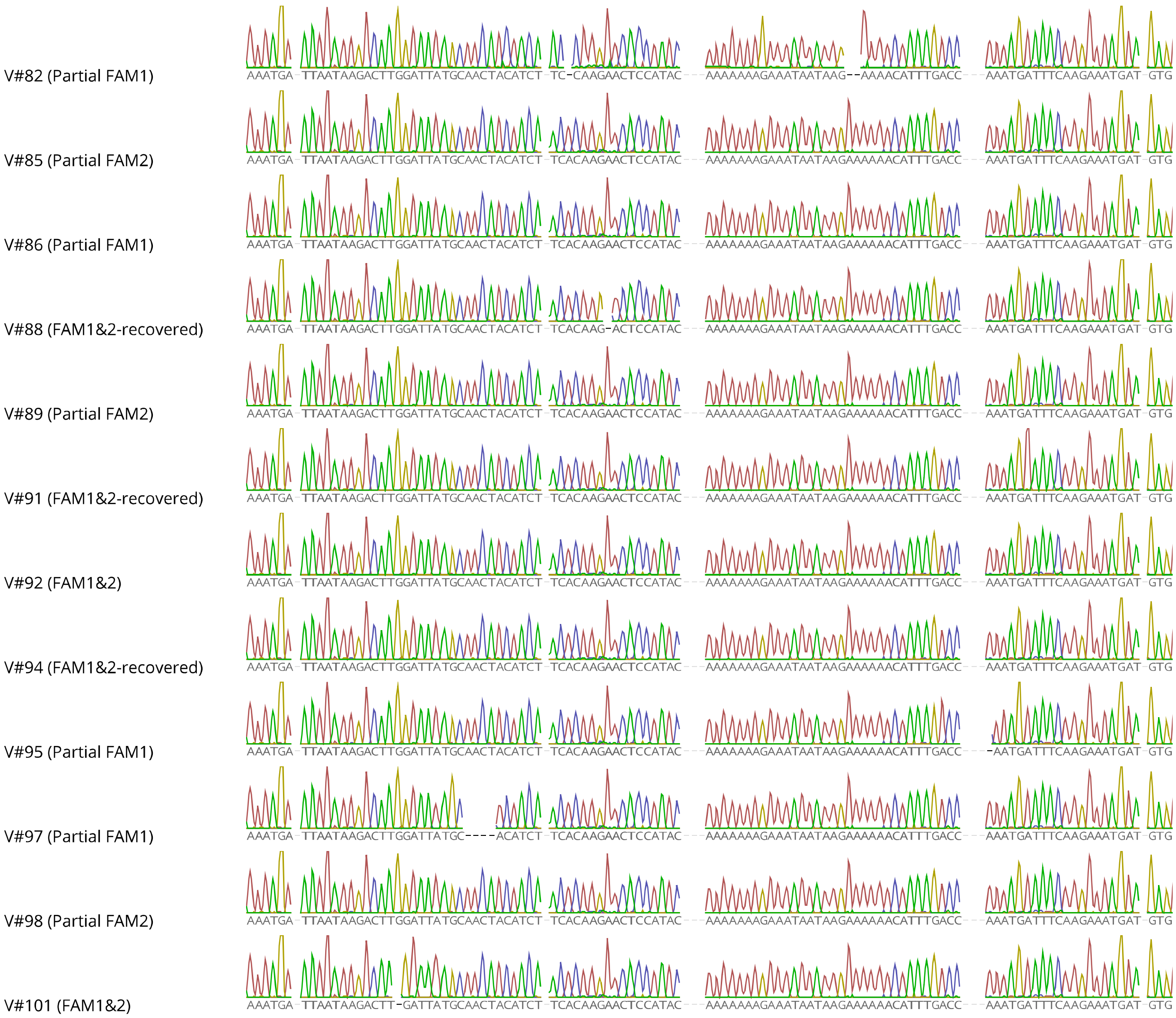
V#82 (Partial FAM1)
AAATGA TTAATAAGACTTGGATTATGCAACTACATCT TC-CAAGAACTCCATAC AAAAAAAGAAATAATAAG--AAAACATTTGACC AAATGATTTCAAGAAATGAT GTG
V#85 (Partial FAM2)
AAATGA TTAATAAGACTTGGATTATGCAACTACATCT TCACAAGAACTCCATAC AAAAAAAGAAATAATAAGAAAAAACATTTGACC AAATGATTTCAAGAAATGAT GTG
V#86 (Partial FAM1)
AAATGA TTAATAAGACTTGGATTATGCAACTACATCT TCACAAGAACTCCATAC AAAAAAAGAAATAATAAGAAAAAACATTTGACC AAATGATTTCAAGAAATGAT GTG
V#88 (FAM1&2-recovered)
AAATGA TTAATAAGACTTGGATTATGCAACTACATCT TCACAAG-ACTCCATAC AAAAAAAGAAATAATAAGAAAAAACATTTGACC AAATGATTTCAAGAAATGAT GTG
V#89 (Partial FAM2)
AAATGA TTAATAAGACTTGGATTATGCAACTACATCT TCACAAGAACTCCATAC AAAAAAAGAAATAATAAGAAAAAACATTTGACC AAATGATTTCAAGAAATGAT GTG
V#91 (FAM1&2-recovered)
AAATGA TTAATAAGACTTGGATTATGCAACTACATCT TCACAAGAACTCCATAC AAAAAAAGAAATAATAAGAAAAAACATTTGACC AAATGATTTCAAGAAATGAT GTG
V#92 (FAM1&2)
AAATGA TTAATAAGACTTGGATTATGCAACTACATCT TCACAAGAACTCCATAC AAAAAAAGAAATAATAAGAAAAAACATTTGACC AAATGATTTCAAGAAATGAT GTG
V#94 (FAM1&2-recovered)
AAATGA TTAATAAGACTTGGATTATGCAACTACATCT TCACAAGAACTCCATAC AAAAAAAGAAATAATAAGAAAAAACATTTGACC AAATGATTTCAAGAAATGAT GTG
V#95 (Partial FAM1)
AAATGA TTAATAAGACTTGGATTATGCAACTACATCT TCACAAGAACTCCATAC AAAAAAAGAAATAATAAGAAAAAACATTTGACC -AATGATTTCAAGAAATGAT GTG
V#97 (Partial FAM1)
AAATGA TTAATAAGACTTGGATTATGC----ACATCT TCACAAGAACTCCATAC AAAAAAAGAAATAATAAGAAAAAACATTTGACC AAATGATTTCAAGAAATGAT GTG
V#98 (Partial FAM2)
AAATGA TTAATAAGACTTGGATTATGCAACTACATCT TCACAAGAACTCCATAC AAAAAAAGAAATAATAAGAAAAAACATTTGACC AAATGATTTCAAGAAATGAT GTG
V#101 (FAM1&2)
AAATGA TTAATAAGACTT-GATTATGCAACTACATCT TCACAAGAACTCCATAC AAAAAAAGAAATAATAAGAAAAAACATTTGACC AAATGATTTCAAGAAATGAT GTG

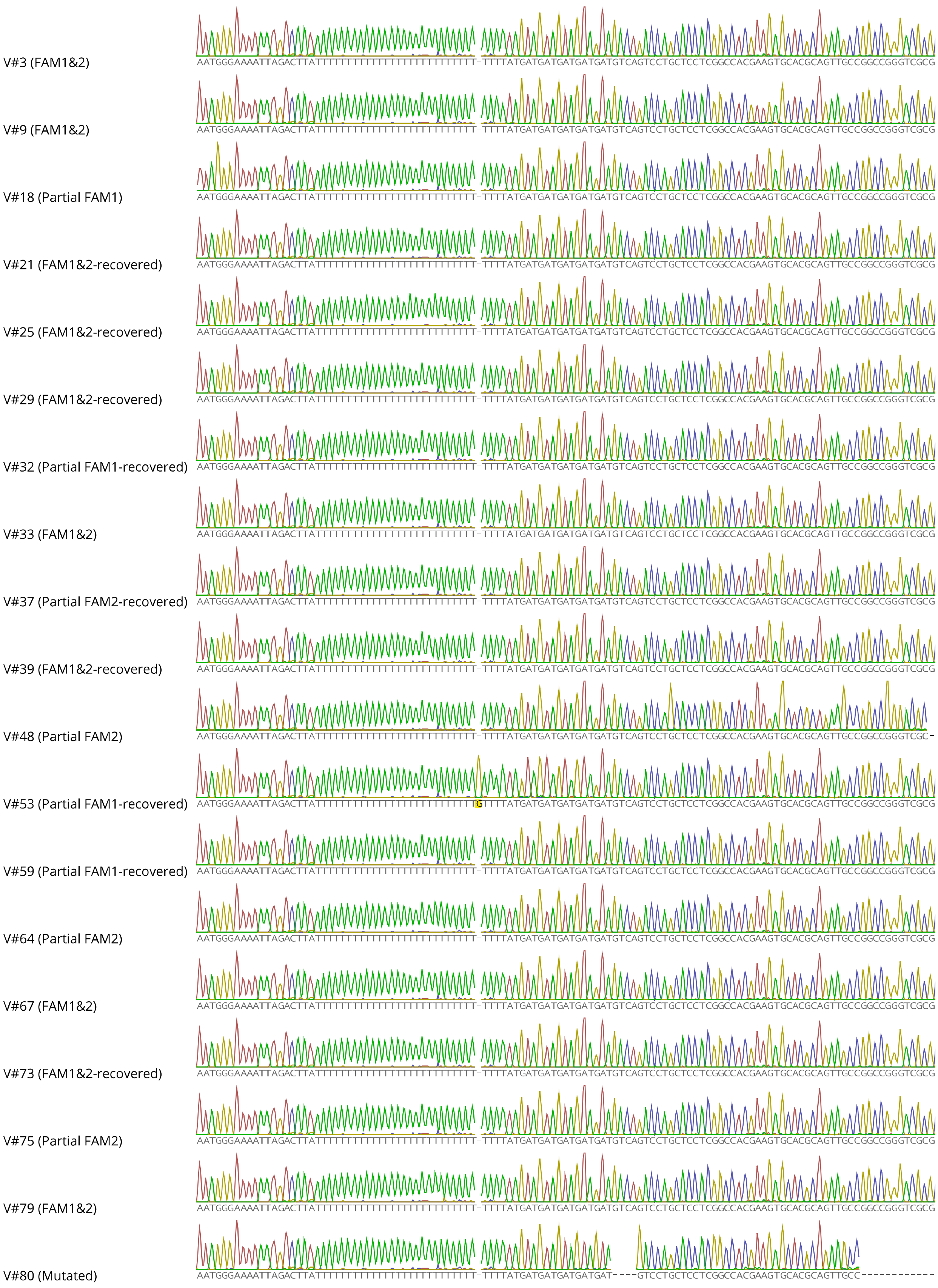

V#3 (FAM1&2)
V#9 (FAM1&2)
V#18 (Partial FAM1)
V#21 (FAM1&2-recovered)
V#25 (FAM1&2-recovered)
V#29 (FAM1&2-recovered)
V#32 (Partial FAM1-recovered)
V#33 (FAM1&2)
V#37 (Partial FAM2-recovered)
V#39 (FAM1&2-recovered)
V#48 (Partial FAM2)
V#53 (Partial FAM1-recovered)
V#59 (Partial FAM1-recovered)
V#64 (Partial FAM2)
V#67 (FAM1&2)
V#73 (FAM1&2-recovered)
V#75 (Partial FAM2)
V#79 (FAM1&2)
V#80 (Mutated)

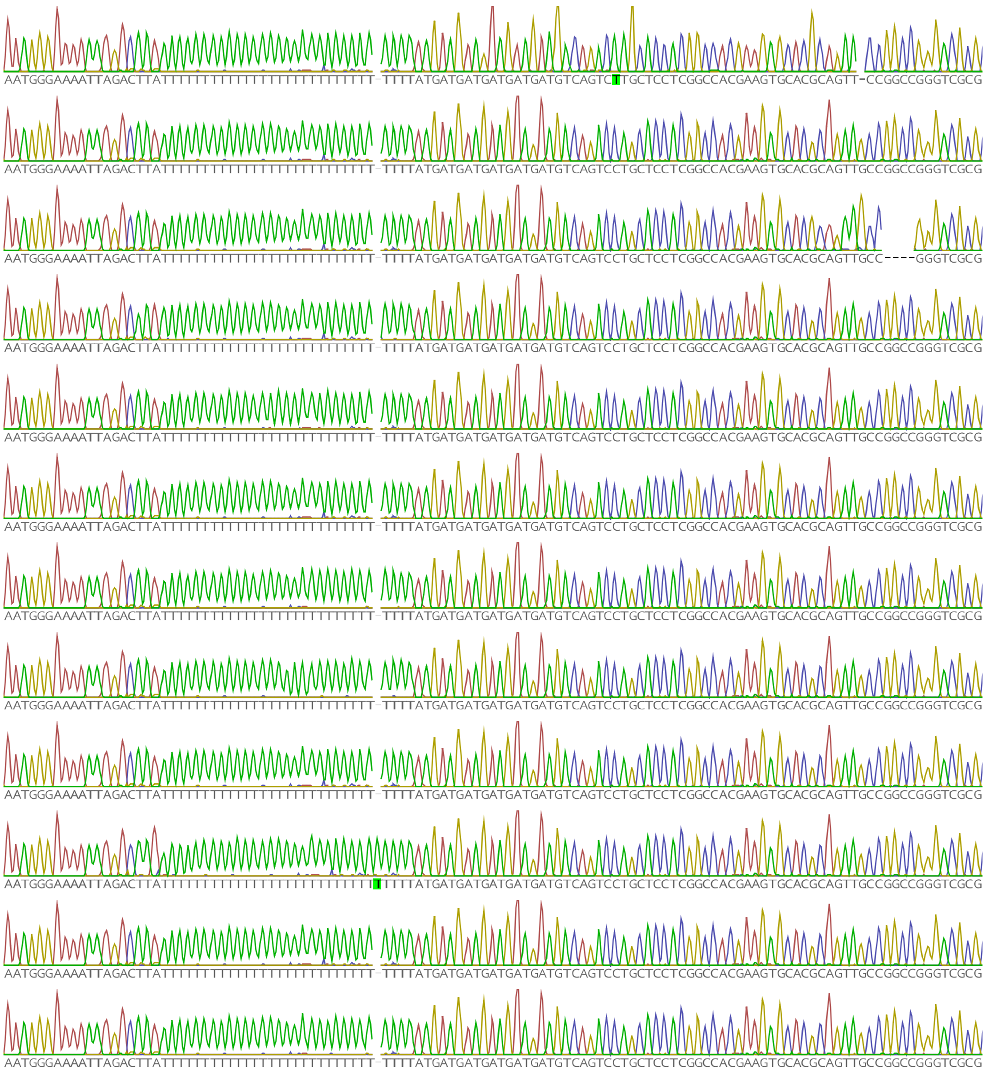

V#82 (Partial FAM1)
V#85 (Partial FAM2)
V#86 (Partial FAM1)
V#88 (FAM1&2-recovered)
V#89 (Partial FAM2)
V#91 (FAM1&2-recovered)
V#92 (FAM1&2)
V#94 (FAM1&2-recovered)
V#95 (Partial FAM1)
V#97 (Partial FAM1)
V#98 (Partial FAM2)
V#101 (FAM1&2)

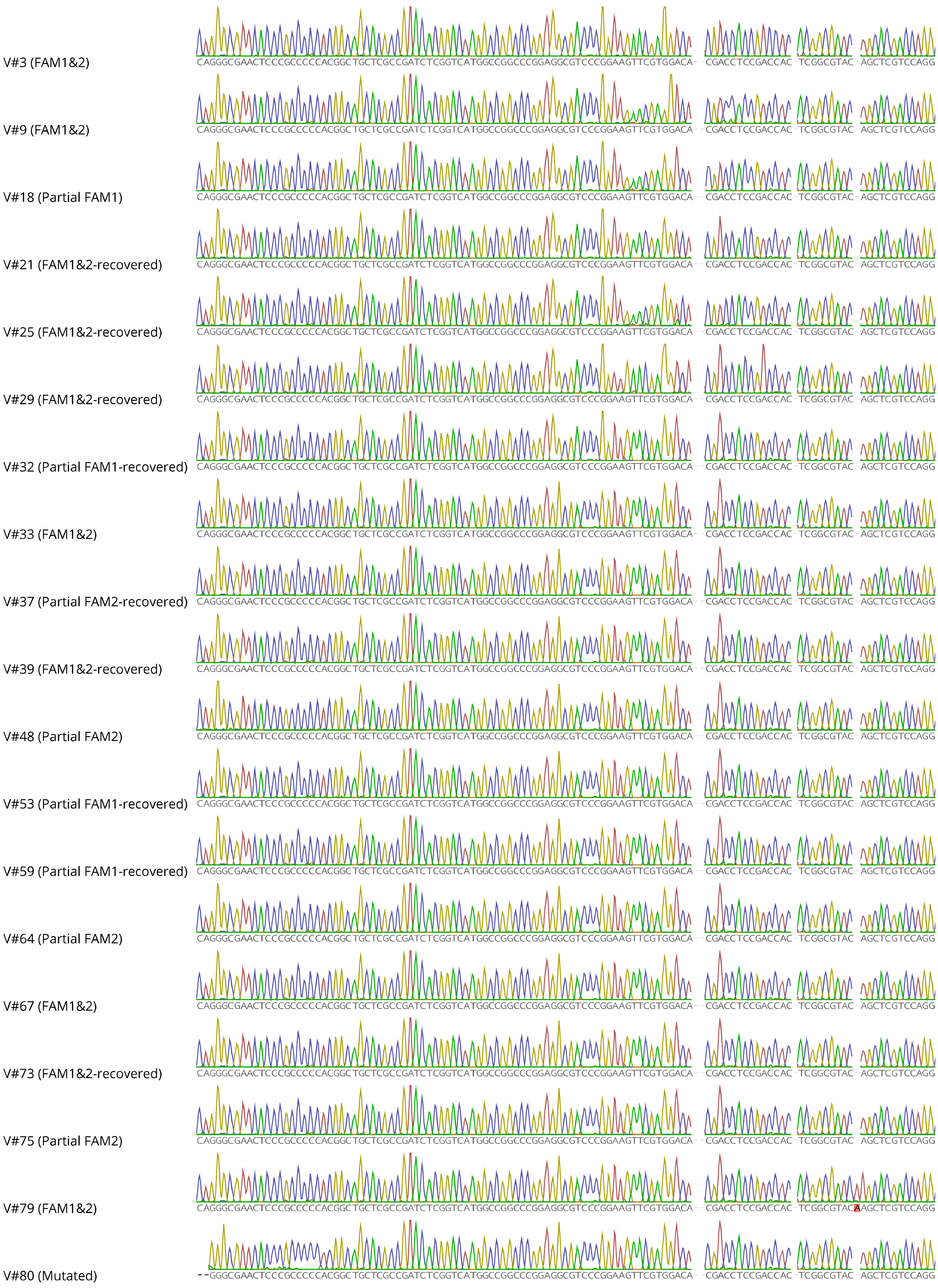
V#3 (FAM1&2)
CAGGGCGAACTCCCGCCCCCACGGCTGCTCGCCGATCTCGGTCATGGCCGGCCCGGAGGCGTCCCGGAAGTTCGTGGACA CGACCTCCGACCAC TCGGCGTAC AGCTCGTCCAGG
V#9 (FAM1&2)
CAGGGCGAACTCCCGCCCCCACGGCTGCTCGCCGATCTCGGTCATGGCCGGCCCGGAGGCGTCCCGGAAGTTCGTGGACA CGACCTCCGACCAC TCGGCGTAC AGCTCGTCCAGG
V#18 (Partial FAM1)
CAGGGCGAACTCCCGCCCCCACGGCTGCTCGCCGATCTCGGTCATGGCCGGCCCGGAGGCGTCCCGGAAGTTCGTGGACA CGACCTCCGACCAC TCGGCGTAC AGCTCGTCCAGG
V#21 (FAM1&2-recovered)
CAGGGCGAACTCCCGCCCCCACGGCTGCTCGCCGATCTCGGTCATGGCCGGCCCGGAGGCGTCCCGGAAGTTCGTGGACA CGACCTCCGACCAC TCGGCGTAC AGCTCGTCCAGG
V#25 (FAM1&2-recovered)
CAGGGCGAACTCCCGCCCCCACGGCTGCTCGCCGATCTCGGTCATGGCCGGCCCGGAGGCGTCCCGGAAGTTCGTGGACA CGACCTCCGACCAC TCGGCGTAC AGCTCGTCCAGG
V#29 (FAM1&2-recovered)
CAGGGCGAACTCCCGCCCCCACGGCTGCTCGCCGATCTCGGTCATGGCCGGCCCGGAGGCGTCCCGGAAGTTCGTGGACA CGACCTCCGACCAC TCGGCGTAC AGCTCGTCCAGG
V#32 (Partial FAM1-recovered)
CAGGGCGAACTCCCGCCCCCACGGCTGCTCGCCGATCTCGGTCATGGCCGGCCCGGAGGCGTCCCGGAAGTTCGTGGACA CGACCTCCGACCAC TCGGCGTAC AGCTCGTCCAGG
V#33 (FAM1&2)
CAGGGCGAACTCCCGCCCCCACGGCTGCTCGCCGATCTCGGTCATGGCCGGCCCGGAGGCGTCCCGGAAGTTCGTGGACA CGACCTCCGACCAC TCGGCGTAC AGCTCGTCCAGG
V#37 (Partial FAM2-recovered)
CAGGGCGAACTCCCGCCCCCACGGCTGCTCGCCGATCTCGGTCATGGCCGGCCCGGAGGCGTCCCGGAAGTTCGTGGACA CGACCTCCGACCAC TCGGCGTAC AGCTCGTCCAGG
V#39 (FAM1&2-recovered)
CAGGGCGAACTCCCGCCCCCACGGCTGCTCGCCGATCTCGGTCATGGCCGGCCCGGAGGCGTCCCGGAAGTTCGTGGACA CGACCTCCGACCAC TCGGCGTAC AGCTCGTCCAGG
V#48 (Partial FAM2)
CAGGGCGAACTCCCGCCCCCACGGCTGCTCGCCGATCTCGGTCATGGCCGGCCCGGAGGCGTCCCGGAAGTTCGTGGACA CGACCTCCGACCAC TCGGCGTAC AGCTCGTCCAGG
V#53 (Partial FAM1-recovered)
CAGGGCGAACTCCCGCCCCCACGGCTGCTCGCCGATCTCGGTCATGGCCGGCCCGGAGGCGTCCCGGAAGTTCGTGGACA CGACCTCCGACCAC TCGGCGTAC AGCTCGTCCAGG
V#59 (Partial FAM1-recovered)
CAGGGCGAACTCCCGCCCCCACGGCTGCTCGCCGATCTCGGTCATGGCCGGCCCGGAGGCGTCCCGGAAGTTCGTGGACA CGACCTCCGACCAC TCGGCGTAC AGCTCGTCCAGG
V#64 (Partial FAM2)
CAGGGCGAACTCCCGCCCCCACGGCTGCTCGCCGATCTCGGTCATGGCCGGCCCGGAGGCGTCCCGGAAGTTCGTGGACA CGACCTCCGACCAC TCGGCGTAC AGCTCGTCCAGG
V#67 (FAM1&2)
CAGGGCGAACTCCCGCCCCCACGGCTGCTCGCCGATCTCGGTCATGGCCGGCCCGGAGGCGTCCCGGAAGTTCGTGGACA CGACCTCCGACCAC TCGGCGTAC AGCTCGTCCAGG
V#73 (FAM1&2-recovered)
CAGGGCGAACTCCCGCCCCCACGGCTGCTCGCCGATCTCGGTCATGGCCGGCCCGGAGGCGTCCCGGAAGTTCGTGGACA CGACCTCCGACCAC TCGGCGTAC AGCTCGTCCAGG
V#75 (Partial FAM2)
CAGGGCGAACTCCCGCCCCCACGGCTGCTCGCCGATCTCGGTCATGGCCGGCCCGGAGGCGTCCCGGAAGTTCGTGGACA CGACCTCCGACCAC TCGGCGTAC AGCTCGTCCAGG
V#79 (FAM1&2)
CAGGGCGAACTCCCGCCCCCACGGCTGCTCGCCGATCTCGGTCATGGCCGGCCCGGAGGCGTCCCGGAAGTTCGTGGACA CGACCTCCGACCAC TCGGCGTACAAGCTCGTCCAGG
V#80 (Mutated)
--GGGCGAACTCCCGCCCCCACGGCTGCTCGCCGATCTCGGTCATGGCCGGCCCGGAGGCGTCCCGGAAGTTCGTGGACA CGACCTCCGACCAC TCGGCGTAC AGCTCGTCCAGG

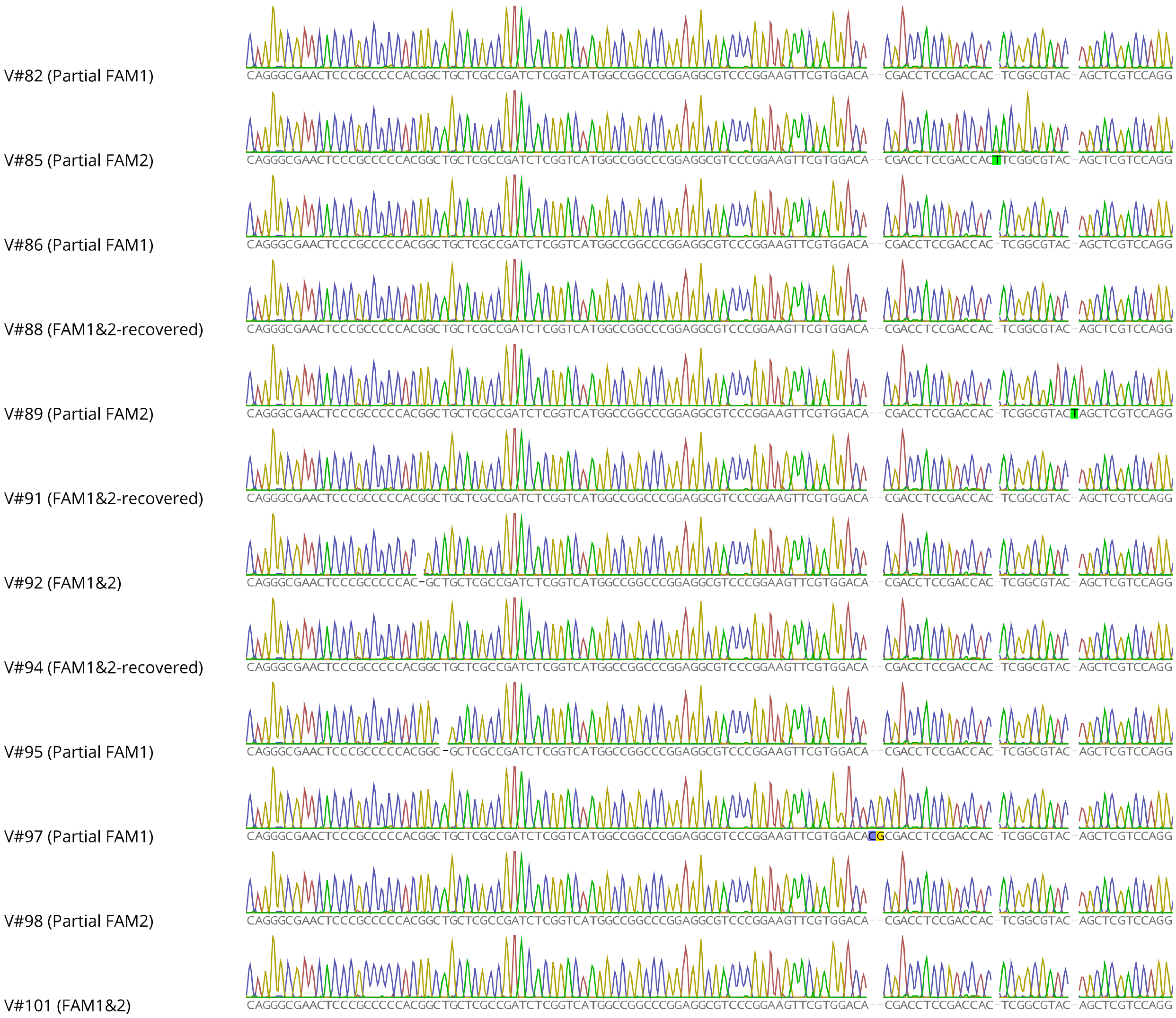
V#82 (Partial FAM1)
CAGGGCGAACTCCCGCCCCCACGGCTGCTCGCCGATCTCGGTCATGGCCGGCCCGGAGGCGTCCCGGAAGTTCGTGGACA CGACCTCCGACCAC TCGGCGTAC AGCTCGTCCAGG
V#85 (Partial FAM2)
CAGGGCGAACTCCCGCCCCCACGGCTGCTCGCCGATCTCGGTCATGGCCGGCCCGGAGGCGTCCCGGAAGTTCGTGGACA CGACCTCCGACCACTTCGGCGTAC AGCTCGTCCAGG
V#86 (Partial FAM1)
CAGGGCGAACTCCCGCCCCCACGGCTGCTCGCCGATCTCGGTCATGGCCGGCCCGGAGGCGTCCCGGAAGTTCGTGGACA CGACCTCCGACCAC TCGGCGTAC AGCTCGTCCAGG
V#88 (FAM1&2-recovered)
CAGGGCGAACTCCCGCCCCCACGGCTGCTCGCCGATCTCGGTCATGGCCGGCCCGGAGGCGTCCCGGAAGTTCGTGGACA CGACCTCCGACCAC TCGGCGTAC AGCTCGTCCAGG
V#89 (Partial FAM2)
CAGGGCGAACTCCCGCCCCCACGGCTGCTCGCCGATCTCGGTCATGGCCGGCCCGGAGGCGTCCCGGAAGTTCGTGGACA CGACCTCCGACCAC TCGGCGTACTAGCTCGTCCAGG
V#91 (FAM1&2-recovered)
CAGGGCGAACTCCCGCCCCCACGGCTGCTCGCCGATCTCGGTCATGGCCGGCCCGGAGGCGTCCCGGAAGTTCGTGGACA CGACCTCCGACCAC TCGGCGTAC AGCTCGTCCAGG
V#92 (FAM1&2)
CAGGGCGAACTCCCGCCCCCAC-GCTGCTCGCCGATCTCGGTCATGGCCGGCCCGGAGGCGTCCCGGAAGTTCGTGGACA CGACCTCCGACCAC TCGGCGTAC AGCTCGTCCAGG
V#94 (FAM1&2-recovered)
CAGGGCGAACTCCCGCCCCCACGGCTGCTCGCCGATCTCGGTCATGGCCGGCCCGGAGGCGTCCCGGAAGTTCGTGGACA CGACCTCCGACCAC TCGGCGTAC AGCTCGTCCAGG
V#95 (Partial FAM1)
CAGGGCGAACTCCCGCCCCCACGGC-GCTCGCCGATCTCGGTCATGGCCGGCCCGGAGGCGTCCCGGAAGTTCGTGGACA CGACCTCCGACCAC TCGGCGTAC AGCTCGTCCAGG
V#97 (Partial FAM1)
CAGGGCGAACTCCCGCCCCCACGGCTGCTCGCCGATCTCGGTCATGGCCGGCCCGGAGGCGTCCCGGAAGTTCGTGGACACGCGACCTCCGACCAC TCGGCGTAC AGCTCGTCCAGG
V#98 (Partial FAM2)
CAGGGCGAACTCCCGCCCCCACGGCTGCTCGCCGATCTCGGTCATGGCCGGCCCGGAGGCGTCCCGGAAGTTCGTGGACA CGACCTCCGACCAC TCGGCGTAC AGCTCGTCCAGG
V#101 (FAM1&2)
CAGGGCGAACTCCCGCCCCCACGGCTGCTCGCCGATCTCGGTCATGGCCGGCCCGGAGGCGTCCCGGAAGTTCGTGGACA CGACCTCCGACCAC TCGGCGTAC AGCTCGTCCAGG

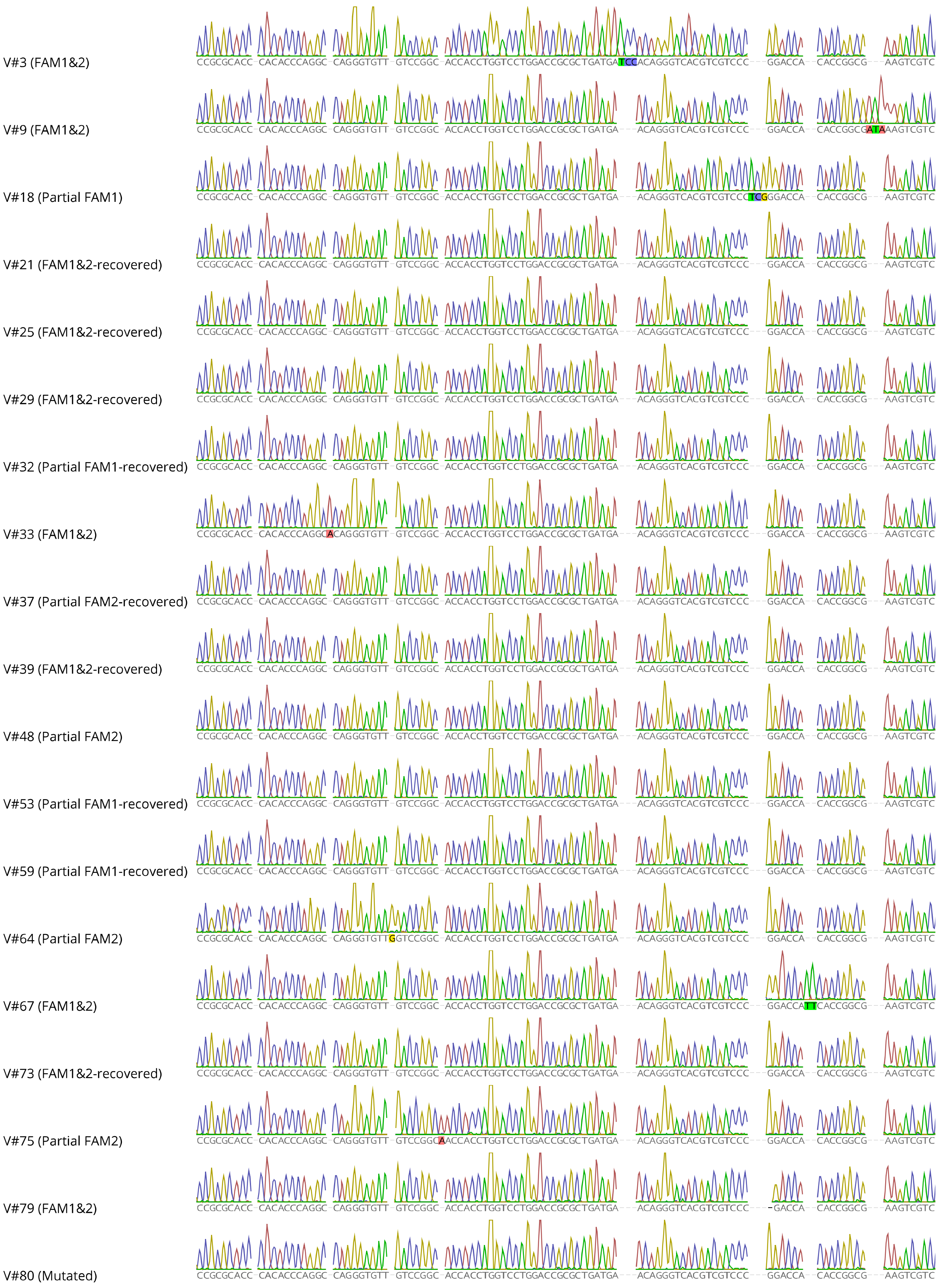

V#3 (FAM1&2)
V#9 (FAM1&2)
V#18 (Partial FAM1)
V#21 (FAM1&2-recovered)
V#25 (FAM1&2-recovered)
V#29 (FAM1&2-recovered)
V#32 (Partial FAM1-recovered)
V#33 (FAM1&2)
V#37 (Partial FAM2-recovered)
V#39 (FAM1&2-recovered)
V#48 (Partial FAM2)
V#53 (Partial FAM1-recovered)
V#59 (Partial FAM1-recovered)
V#64 (Partial FAM2)
V#67 (FAM1&2)
V#73 (FAM1&2-recovered)
V#75 (Partial FAM2)
V#79 (FAM1&2)
V#80 (Mutated)

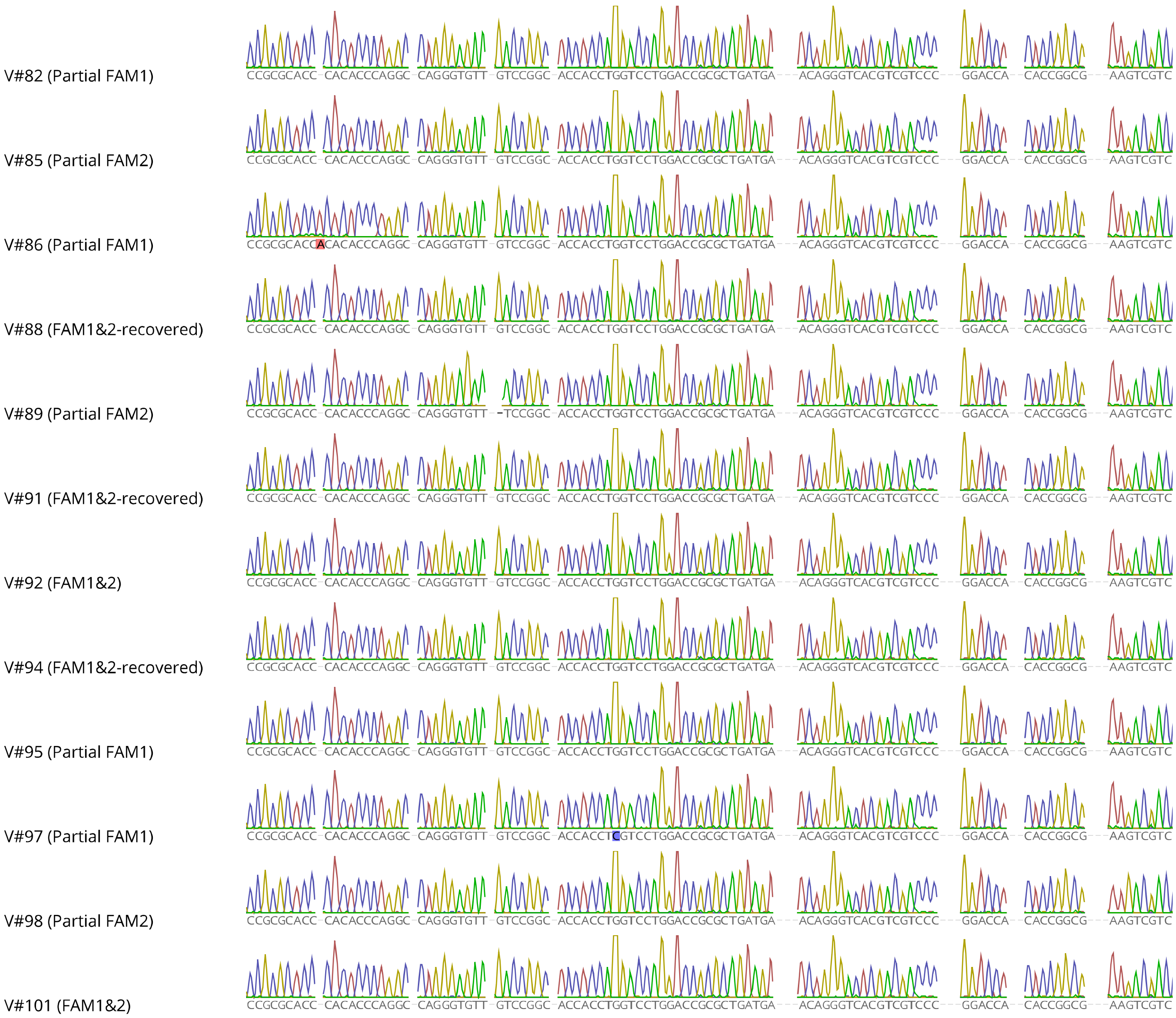
V#82 (Partial FAM1)
CCGCGCACC CACACCCAGGC CAGGGTGTT GTCCGGC ACCACCTGGTCCTGGACCGCGCTGATGA ACAGGGTCACGTCGTCCC GGACCA CACCGGCG AAGTCGTC
V#85 (Partial FAM2)
CCGCGCACC CACACCCAGGC CAGGGTGTT GTCCGGC ACCACCTGGTCCTGGACCGCGCTGATGA ACAGGGTCACGTCGTCCC GGACCA CACCGGCG AAGTCGTC
V#86 (Partial FAM1)
CCGCGCACCACACACCCAGGC CAGGGTGTT GTCCGGC ACCACCTGGTCCTGGACCGCGCTGATGA ACAGGGTCACGTCGTCCC GGACCA CACCGGCG AAGTCGTC
V#88 (FAM1&2-recovered)
CCGCGCACC CACACCCAGGC CAGGGTGTT GTCCGGC ACCACCTGGTCCTGGACCGCGCTGATGA ACAGGGTCACGTCGTCCC GGACCA CACCGGCG AAGTCGTC
V#89 (Partial FAM2)
CCGCGCACC CACACCCAGGC CAGGGTGTT -TCCGGC ACCACCTGGTCCTGGACCGCGCTGATGA ACAGGGTCACGTCGTCCC GGACCA CACCGGCG AAGTCGTC
V#91 (FAM1&2-recovered)
CCGCGCACC CACACCCAGGC CAGGGTGTT GTCCGGC ACCACCTGGTCCTGGACCGCGCTGATGA ACAGGGTCACGTCGTCCC GGACCA CACCGGCG AAGTCGTC
V#92 (FAM1&2)
CCGCGCACC CACACCCAGGC CAGGGTGTT GTCCGGC ACCACCTGGTCCTGGACCGCGCTGATGA ACAGGGTCACGTCGTCCC GGACCA CACCGGCG AAGTCGTC
V#94 (FAM1&2-recovered)
CCGCGCACC CACACCCAGGC CAGGGTGTT GTCCGGC ACCACCTGGTCCTGGACCGCGCTGATGA ACAGGGTCACGTCGTCCC GGACCA CACCGGCG AAGTCGTC
V#95 (Partial FAM1)
CCGCGCACC CACACCCAGGC CAGGGTGTT GTCCGGC ACCACCTGGTCCTGGACCGCGCTGATGA ACAGGGTCACGTCGTCCC GGACCA CACCGGCG AAGTCGTC
V#97 (Partial FAM1)
CCGCGCACC CACACCCAGGC CAGGGTGTT GTCCGGC ACCACCTCGTCCTGGACCGCGCTGATGA ACAGGGTCACGTCGTCCC GGACCA CACCGGCG AAGTCGTC
V#98 (Partial FAM2)
CCGCGCACC CACACCCAGGC CAGGGTGTT GTCCGGC ACCACCTGGTCCTGGACCGCGCTGATGA ACAGGGTCACGTCGTCCC GGACCA CACCGGCG AAGTCGTC
V#101 (FAM1&2)
CCGCGCACC CACACCCAGGC CAGGGTGTT GTCCGGC ACCACCTGGTCCTGGACCGCGCTGATGA ACAGGGTCACGTCGTCCC GGACCA CACCGGCG AAGTCGTC

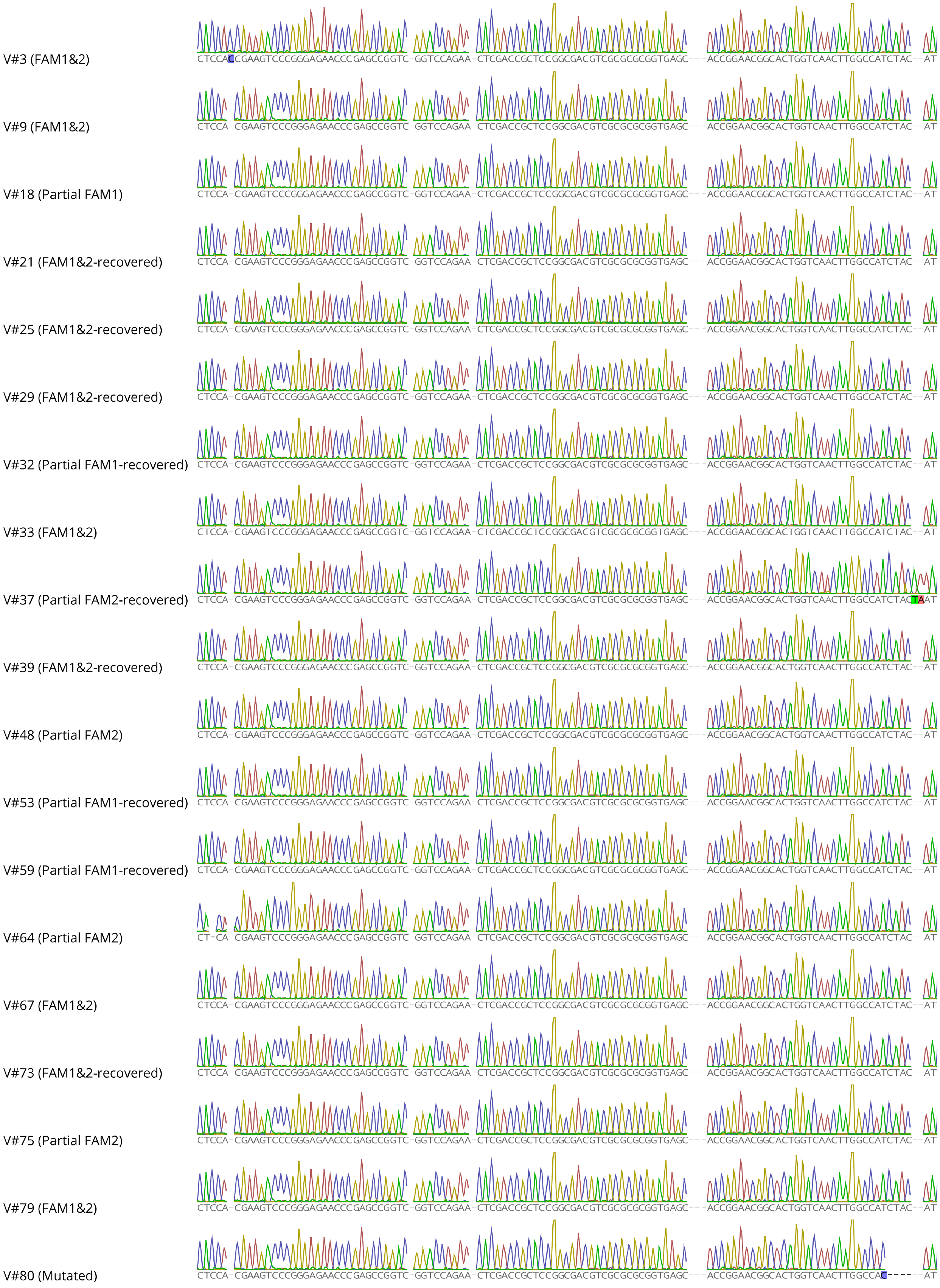

V#3 (FAM1&2)
V#9 (FAM1&2)
V#18 (Partial FAM1)
V#21 (FAM1&2-recovered)
V#25 (FAM1&2-recovered)
V#29 (FAM1&2-recovered)
V#32 (Partial FAM1-recovered)
V#33 (FAM1&2)
V#37 (Partial FAM2-recovered)
V#39 (FAM1&2-recovered)
V#48 (Partial FAM2)
V#53 (Partial FAM1-recovered)
V#59 (Partial FAM1-recovered)
V#64 (Partial FAM2)
V#67 (FAM1&2)
V#73 (FAM1&2-recovered)
V#75 (Partial FAM2)
V#79 (FAM1&2)
V#80 (Mutated)

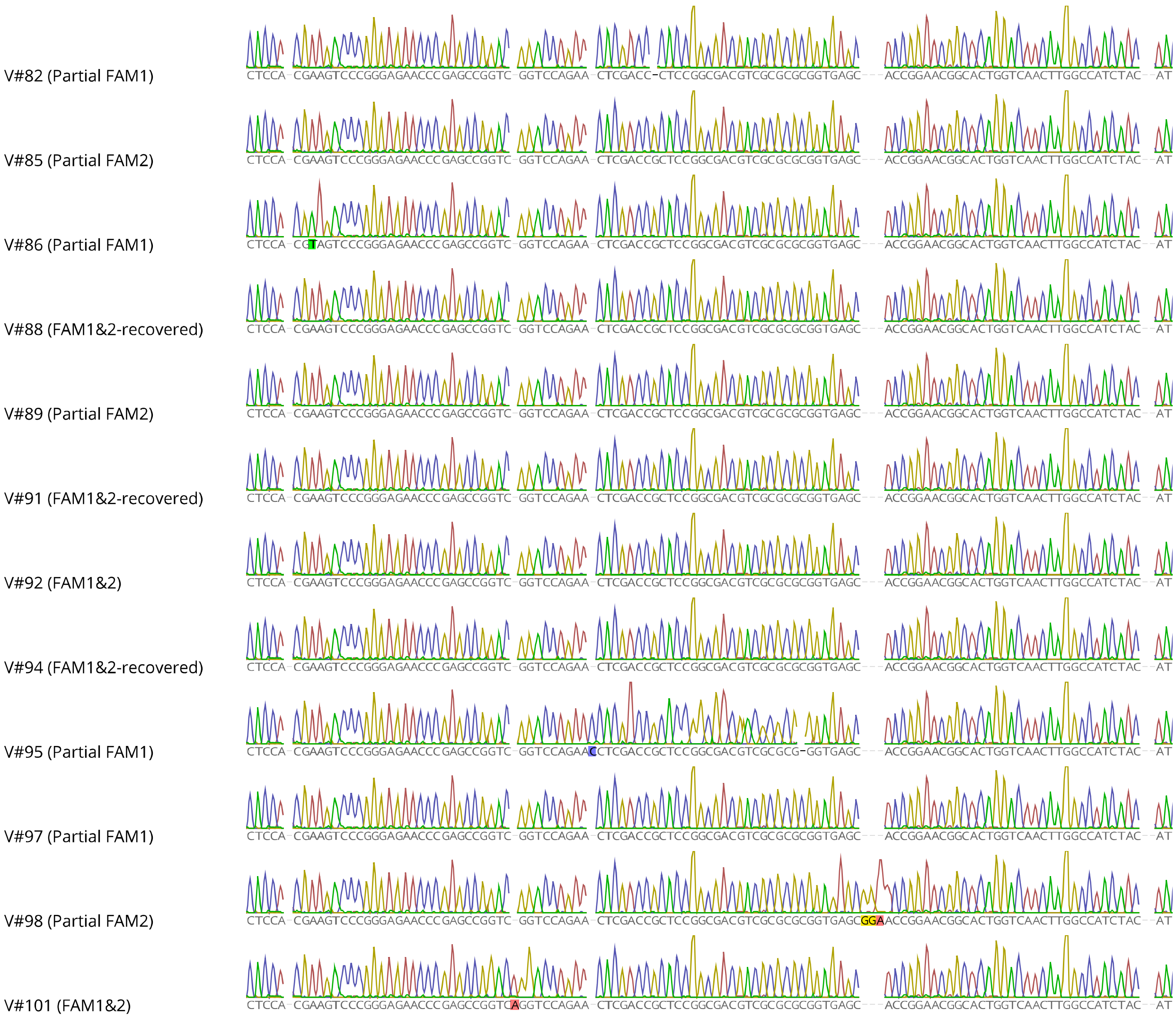
V#82 (Partial FAM1)
V#85 (Partial FAM2)
V#86 (Partial FAM1)
V#88 (FAM1&2-recovered)
V#89 (Partial FAM2)
V#91 (FAM1&2-recovered)
V#92 (FAM1&2)
V#94 (FAM1&2-recovered)
V#95 (Partial FAM1)
V#97 (Partial FAM1)
V#98 (Partial FAM2)
V#101 (FAM1&2)

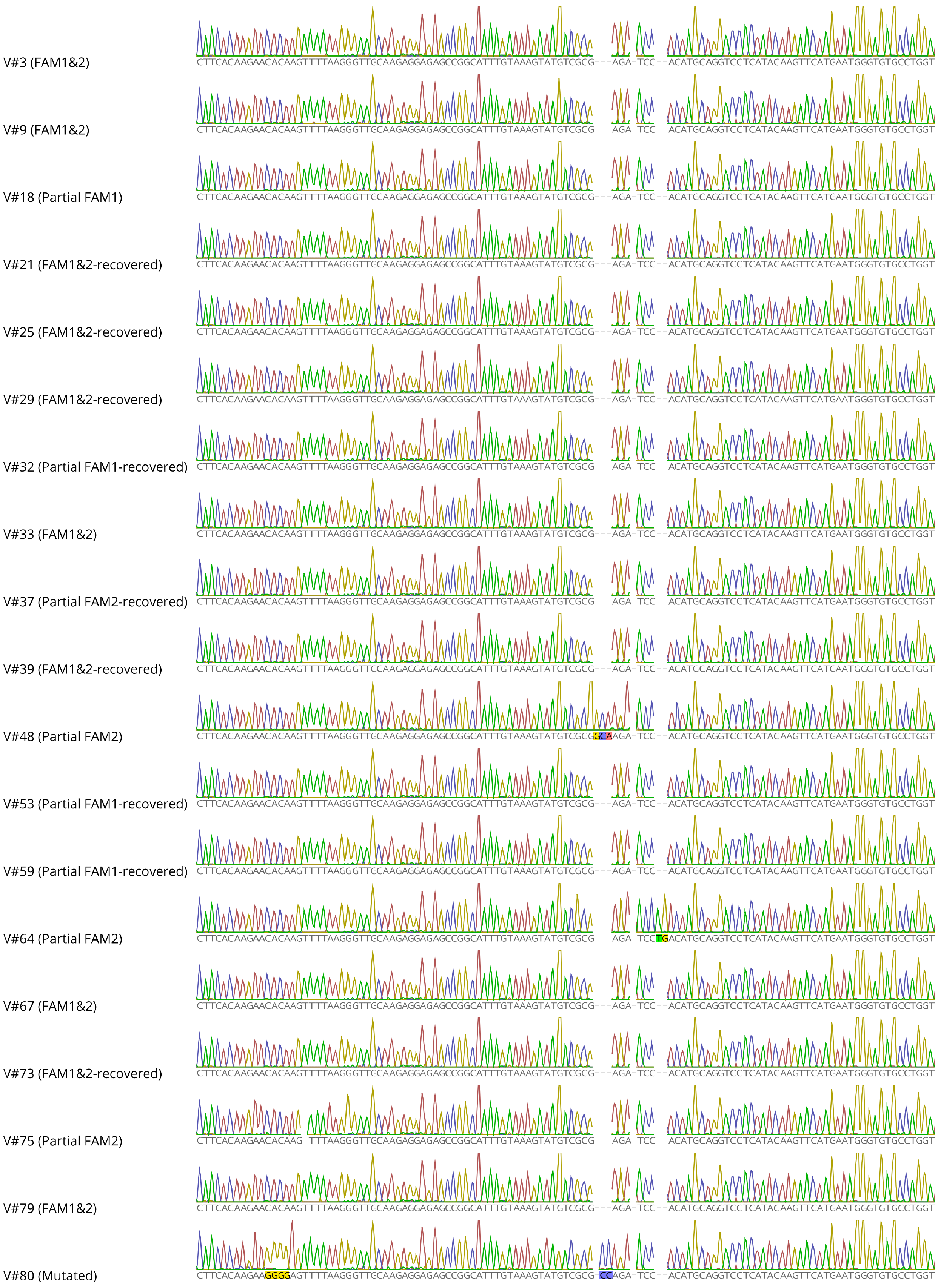
V#3 (FAM1&2)
V#9 (FAM1&2)
V#18 (Partial FAM1)
V#21 (FAM1&2-recovered)
V#25 (FAM1&2-recovered)
V#29 (FAM1&2-recovered)
V#32 (Partial FAM1-recovered)
V#33 (FAM1&2)
V#37 (Partial FAM2-recovered)
V#39 (FAM1&2-recovered)
V#48 (Partial FAM2)
V#53 (Partial FAM1-recovered)
V#59 (Partial FAM1-recovered)
V#64 (Partial FAM2)
V#67 (FAM1&2)
V#73 (FAM1&2-recovered)
V#75 (Partial FAM2)
V#79 (FAM1&2)
V#80 (Mutated)

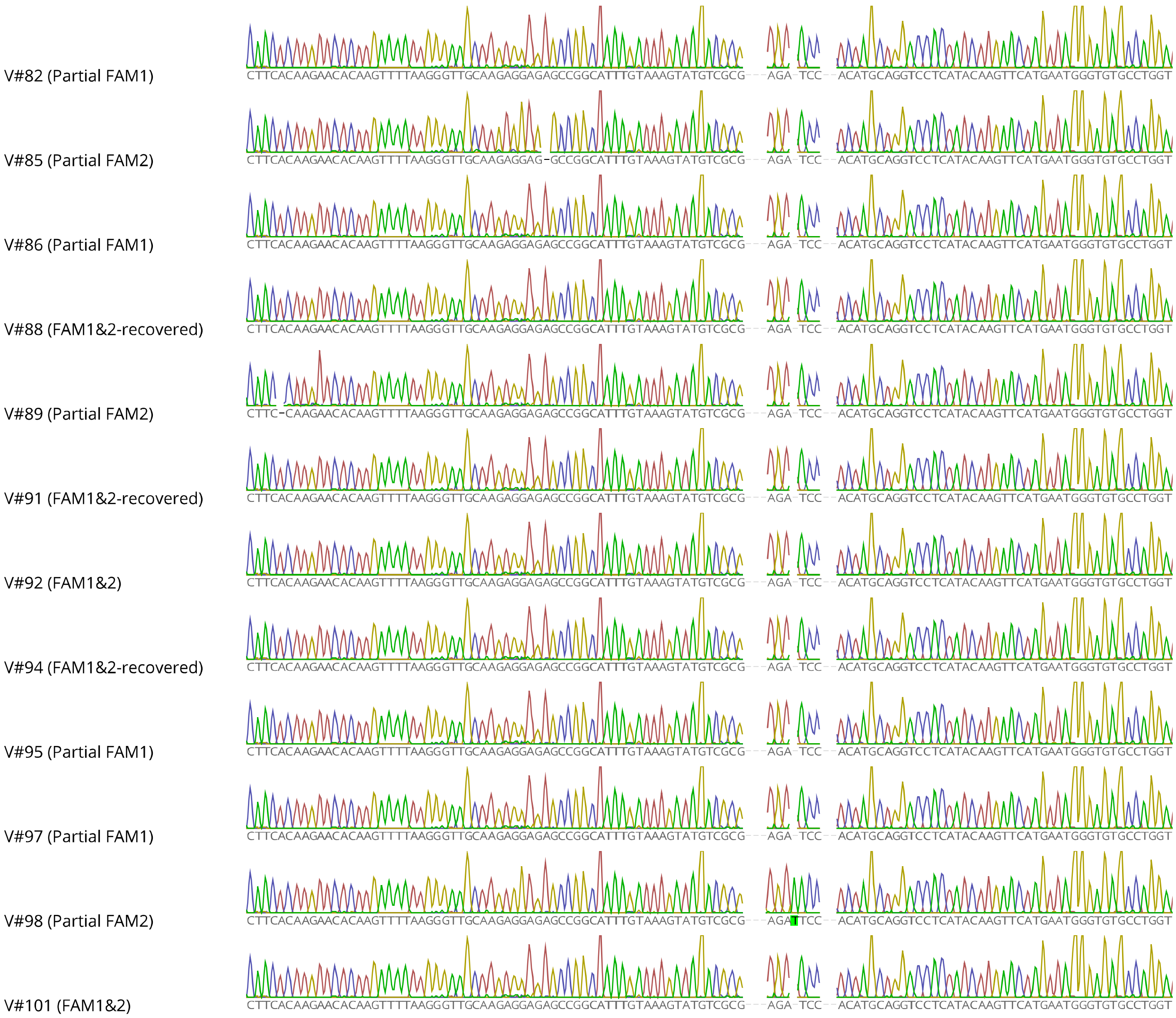
V#82 (Partial FAM1)
CTTCACAAGAACACAAGTTTTAAGGGTTGCAAGAGGAGAGCCGGCATTTGTAAAGTATGTCGCG AGA TCC ACATGCAGGTCCTCATACAAGTTCATGAATGGGTGTGCCTGGT
V#85 (Partial FAM2)
CTTCACAAGAACACAAGTTTTAAGGGTTGCAAGAGGAG-GCCGGCATTTGTAAAGTATGTCGCG AGA TCC ACATGCAGGTCCTCATACAAGTTCATGAATGGGTGTGCCTGGT
V#86 (Partial FAM1)
CTTCACAAGAACACAAGTTTTAAGGGTTGCAAGAGGAGAGCCGGCATTTGTAAAGTATGTCGCG AGA TCC ACATGCAGGTCCTCATACAAGTTCATGAATGGGTGTGCCTGGT
V#88 (FAM1&2-recovered)
CTTCACAAGAACACAAGTTTTAAGGGTTGCAAGAGGAGAGCCGGCATTTGTAAAGTATGTCGCG AGA TCC ACATGCAGGTCCTCATACAAGTTCATGAATGGGTGTGCCTGGT
V#89 (Partial FAM2)
CTTC-CAAGAACACAAGTTTTAAGGGTTGCAAGAGGAGAGCCGGCATTTGTAAAGTATGTCGCG AGA TCC ACATGCAGGTCCTCATACAAGTTCATGAATGGGTGTGCCTGGT
V#91 (FAM1&2-recovered)
CTTCACAAGAACACAAGTTTTAAGGGTTGCAAGAGGAGAGCCGGCATTTGTAAAGTATGTCGCG AGA TCC ACATGCAGGTCCTCATACAAGTTCATGAATGGGTGTGCCTGGT
V#92 (FAM1&2)
CTTCACAAGAACACAAGTTTTAAGGGTTGCAAGAGGAGAGCCGGCATTTGTAAAGTATGTCGCG AGA TCC ACATGCAGGTCCTCATACAAGTTCATGAATGGGTGTGCCTGGT
V#94 (FAM1&2-recovered)
CTTCACAAGAACACAAGTTTTAAGGGTTGCAAGAGGAGAGCCGGCATTTGTAAAGTATGTCGCG AGA TCC ACATGCAGGTCCTCATACAAGTTCATGAATGGGTGTGCCTGGT
V#95 (Partial FAM1)
CTTCACAAGAACACAAGTTTTAAGGGTTGCAAGAGGAGAGCCGGCATTTGTAAAGTATGTCGCG AGA TCC ACATGCAGGTCCTCATACAAGTTCATGAATGGGTGTGCCTGGT
V#97 (Partial FAM1)
CTTCACAAGAACACAAGTTTTAAGGGTTGCAAGAGGAGAGCCGGCATTTGTAAAGTATGTCGCG AGA TCC ACATGCAGGTCCTCATACAAGTTCATGAATGGGTGTGCCTGGT
V#98 (Partial FAM2)
CTTCACAAGAACACAAGTTTTAAGGGTTGCAAGAGGAGAGCCGGCATTTGTAAAGTATGTCGCG AGATTCC ACATGCAGGTCCTCATACAAGTTCATGAATGGGTGTGCCTGGT
V#101 (FAM1&2)
CTTCACAAGAACACAAGTTTTAAGGGTTGCAAGAGGAGAGCCGGCATTTGTAAAGTATGTCGCG AGA TCC ACATGCAGGTCCTCATACAAGTTCATGAATGGGTGTGCCTGGT

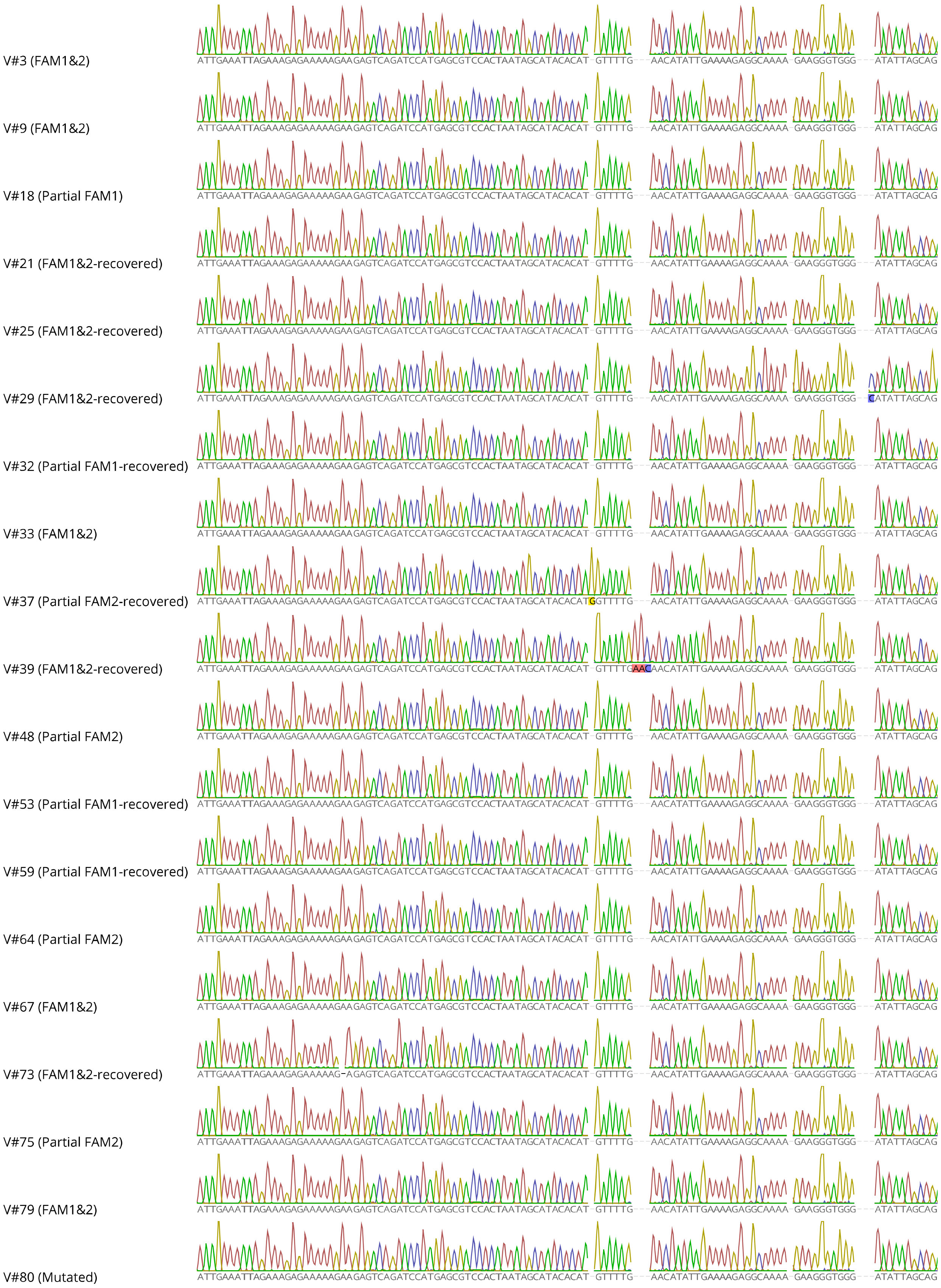
V#3 (FAM1&2)
ATTGAAATTAGAAAGAGAAAAAGAAGAGTCAGATCCATGAGCGTCCACTAATAGCATACACAT GTTTTG AACATATTGAAAAGAGGCAAAA GAAGGGTGGG ATATTAGCAG
V#9 (FAM1&2)
ATTGAAATTAGAAAGAGAAAAAGAAGAGTCAGATCCATGAGCGTCCACTAATAGCATACACAT GTTTTG AACATATTGAAAAGAGGCAAAA GAAGGGTGGG ATATTAGCAG
V#18 (Partial FAM1)
ATTGAAATTAGAAAGAGAAAAAGAAGAGTCAGATCCATGAGCGTCCACTAATAGCATACACAT GTTTTG AACATATTGAAAAGAGGCAAAA GAAGGGTGGG ATATTAGCAG
V#21 (FAM1&2-recovered)
ATTGAAATTAGAAAGAGAAAAAGAAGAGTCAGATCCATGAGCGTCCACTAATAGCATACACAT GTTTTG AACATATTGAAAAGAGGCAAAA GAAGGGTGGG ATATTAGCAG
V#25 (FAM1&2-recovered)
ATTGAAATTAGAAAGAGAAAAAGAAGAGTCAGATCCATGAGCGTCCACTAATAGCATACACAT GTTTTG AACATATTGAAAAGAGGCAAAA GAAGGGTGGG ATATTAGCAG
V#29 (FAM1&2-recovered)
ATTGAAATTAGAAAGAGAAAAAGAAGAGTCAGATCCATGAGCGTCCACTAATAGCATACACAT GTTTTG AACATATTGAAAAGAGGCAAAA GAAGGGTGGG CATATTAGCAG
V#32 (Partial FAM1-recovered)
ATTGAAATTAGAAAGAGAAAAAGAAGAGTCAGATCCATGAGCGTCCACTAATAGCATACACAT GTTTTG AACATATTGAAAAGAGGCAAAA GAAGGGTGGG ATATTAGCAG
V#33 (FAM1&2)
ATTGAAATTAGAAAGAGAAAAAGAAGAGTCAGATCCATGAGCGTCCACTAATAGCATACACAT GTTTTG AACATATTGAAAAGAGGCAAAA GAAGGGTGGG ATATTAGCAG
V#37 (Partial FAM2-recovered)
ATTGAAATTAGAAAGAGAAAAAGAAGAGTCAGATCCATGAGCGTCCACTAATAGCATACACATGGTTTTG AACATATTGAAAAGAGGCAAAA GAAGGGTGGG ATATTAGCAG
V#39 (FAM1&2-recovered)
ATTGAAATTAGAAAGAGAAAAAGAAGAGTCAGATCCATGAGCGTCCACTAATAGCATACACAT GTTTTGAACAACATATTGAAAAGAGGCAAAA GAAGGGTGGG ATATTAGCAG
V#48 (Partial FAM2)
ATTGAAATTAGAAAGAGAAAAAGAAGAGTCAGATCCATGAGCGTCCACTAATAGCATACACAT GTTTTG AACATATTGAAAAGAGGCAAAA GAAGGGTGGG ATATTAGCAG
V#53 (Partial FAM1-recovered)
ATTGAAATTAGAAAGAGAAAAAGAAGAGTCAGATCCATGAGCGTCCACTAATAGCATACACAT GTTTTG AACATATTGAAAAGAGGCAAAA GAAGGGTGGG ATATTAGCAG
V#59 (Partial FAM1-recovered)
ATTGAAATTAGAAAGAGAAAAAGAAGAGTCAGATCCATGAGCGTCCACTAATAGCATACACAT GTTTTG AACATATTGAAAAGAGGCAAAA GAAGGGTGGG ATATTAGCAG
V#64 (Partial FAM2)
ATTGAAATTAGAAAGAGAAAAAGAAGAGTCAGATCCATGAGCGTCCACTAATAGCATACACAT GTTTTG AACATATTGAAAAGAGGCAAAA GAAGGGTGGG ATATTAGCAG
V#67 (FAM1&2)
ATTGAAATTAGAAAGAGAAAAAGAAGAGTCAGATCCATGAGCGTCCACTAATAGCATACACAT GTTTTG AACATATTGAAAAGAGGCAAAA GAAGGGTGGG ATATTAGCAG
V#73 (FAM1&2-recovered)
ATTGAAATTAGAAAGAGAAAAAG-AGAGTCAGATCCATGAGCGTCCACTAATAGCATACACAT GTTTTG AACATATTGAAAAGAGGCAAAA GAAGGGTGGG ATATTAGCAG
V#75 (Partial FAM2)
ATTGAAATTAGAAAGAGAAAAAGAAGAGTCAGATCCATGAGCGTCCACTAATAGCATACACAT GTTTTG AACATATTGAAAAGAGGCAAAA GAAGGGTGGG ATATTAGCAG
V#79 (FAM1&2)
ATTGAAATTAGAAAGAGAAAAAGAAGAGTCAGATCCATGAGCGTCCACTAATAGCATACACAT GTTTTG AACATATTGAAAAGAGGCAAAA GAAGGGTGGG ATATTAGCAG
V#80 (Mutated)
ATTGAAATTAGAAAGAGAAAAAGAAGAGTCAGATCCATGAGCGTCCACTAATAGCATACACAT GTTTTG AACATATTGAAAAGAGGCAAAA GAAGGGTGGG ATATTAGCAG

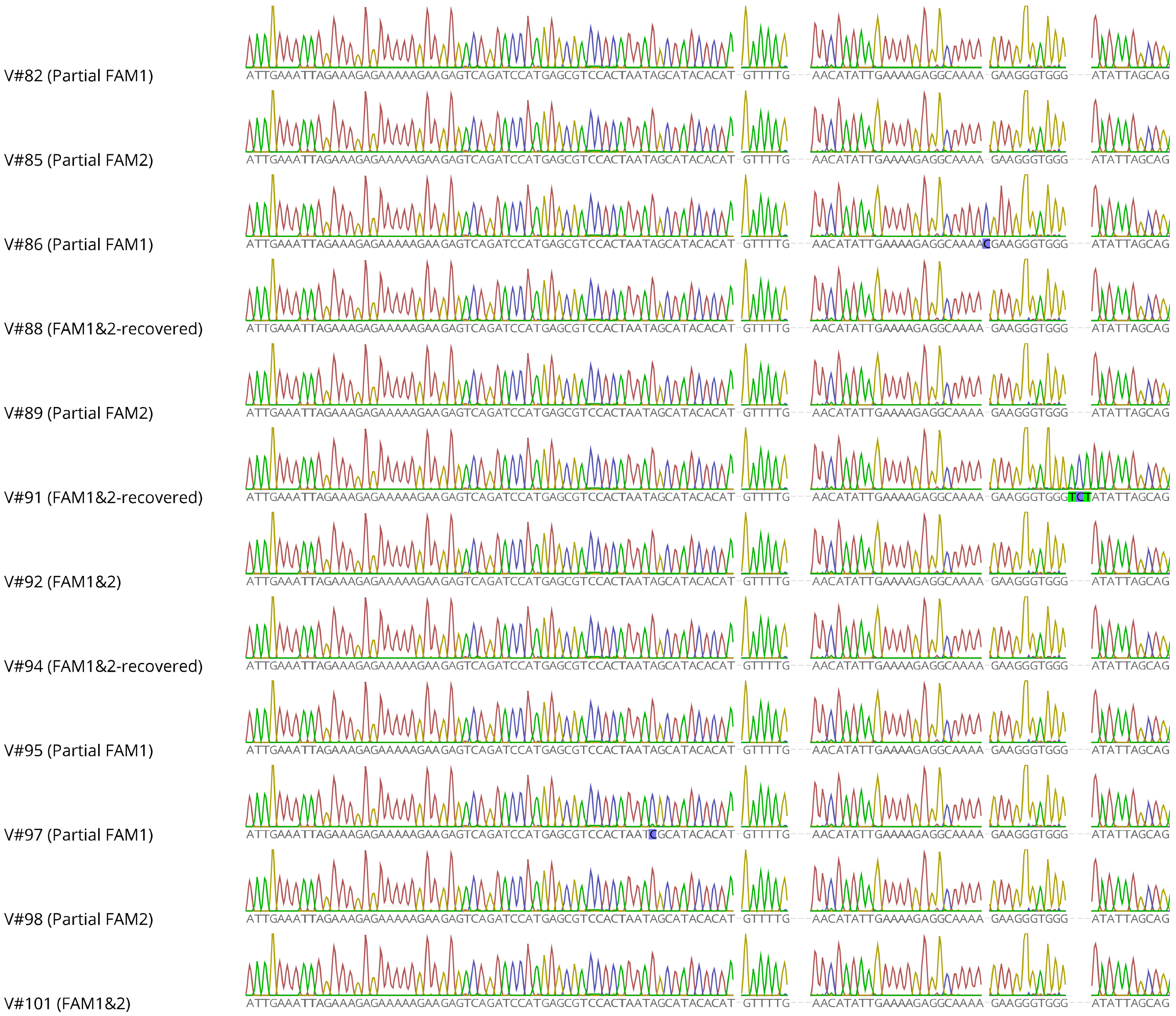
V#82 (Partial FAM1)
ATTGAAATTAGAAAGAGAAAAAGAAGAGTCAGATCCATGAGCGTCCACTAATAGCATACACAT GTTTTG AACATATTGAAAAGAGGCAAAA GAAGGGTGGG ATATTAGCAG
V#85 (Partial FAM2)
ATTGAAATTAGAAAGAGAAAAAGAAGAGTCAGATCCATGAGCGTCCACTAATAGCATACACAT GTTTTG AACATATTGAAAAGAGGCAAAA GAAGGGTGGG ATATTAGCAG
V#86 (Partial FAM1)
ATTGAAATTAGAAAGAGAAAAAGAAGAGTCAGATCCATGAGCGTCCACTAATAGCATACACAT GTTTTG AACATATTGAAAAGAGGCAAAACGAAGGGTGGG ATATTAGCAG
V#88 (FAM1&2-recovered)
ATTGAAATTAGAAAGAGAAAAAGAAGAGTCAGATCCATGAGCGTCCACTAATAGCATACACAT GTTTTG AACATATTGAAAAGAGGCAAAA GAAGGGTGGG ATATTAGCAG
V#89 (Partial FAM2)
ATTGAAATTAGAAAGAGAAAAAGAAGAGTCAGATCCATGAGCGTCCACTAATAGCATACACAT GTTTTG AACATATTGAAAAGAGGCAAAA GAAGGGTGGG ATATTAGCAG
V#91 (FAM1&2-recovered)
ATTGAAATTAGAAAGAGAAAAAGAAGAGTCAGATCCATGAGCGTCCACTAATAGCATACACAT GTTTTG AACATATTGAAAAGAGGCAAAA GAAGGGTGGGTCTATATTAGCAG
V#92 (FAM1&2)
ATTGAAATTAGAAAGAGAAAAAGAAGAGTCAGATCCATGAGCGTCCACTAATAGCATACACAT GTTTTG AACATATTGAAAAGAGGCAAAA GAAGGGTGGG ATATTAGCAG
V#94 (FAM1&2-recovered)
ATTGAAATTAGAAAGAGAAAAAGAAGAGTCAGATCCATGAGCGTCCACTAATAGCATACACAT GTTTTG AACATATTGAAAAGAGGCAAAA GAAGGGTGGG ATATTAGCAG
V#95 (Partial FAM1)
ATTGAAATTAGAAAGAGAAAAAGAAGAGTCAGATCCATGAGCGTCCACTAATAGCATACACAT GTTTTG AACATATTGAAAAGAGGCAAAA GAAGGGTGGG ATATTAGCAG
V#97 (Partial FAM1)
ATTGAAATTAGAAAGAGAAAAAGAAGAGTCAGATCCATGAGCGTCCACTAATCGCATACACAT GTTTTG AACATATTGAAAAGAGGCAAAA GAAGGGTGGG ATATTAGCAG
V#98 (Partial FAM2)
ATTGAAATTAGAAAGAGAAAAAGAAGAGTCAGATCCATGAGCGTCCACTAATAGCATACACAT GTTTTG AACATATTGAAAAGAGGCAAAA GAAGGGTGGG ATATTAGCAG
V#101 (FAM1&2)
ATTGAAATTAGAAAGAGAAAAAGAAGAGTCAGATCCATGAGCGTCCACTAATAGCATACACAT GTTTTG AACATATTGAAAAGAGGCAAAA GAAGGGTGGG ATATTAGCAG

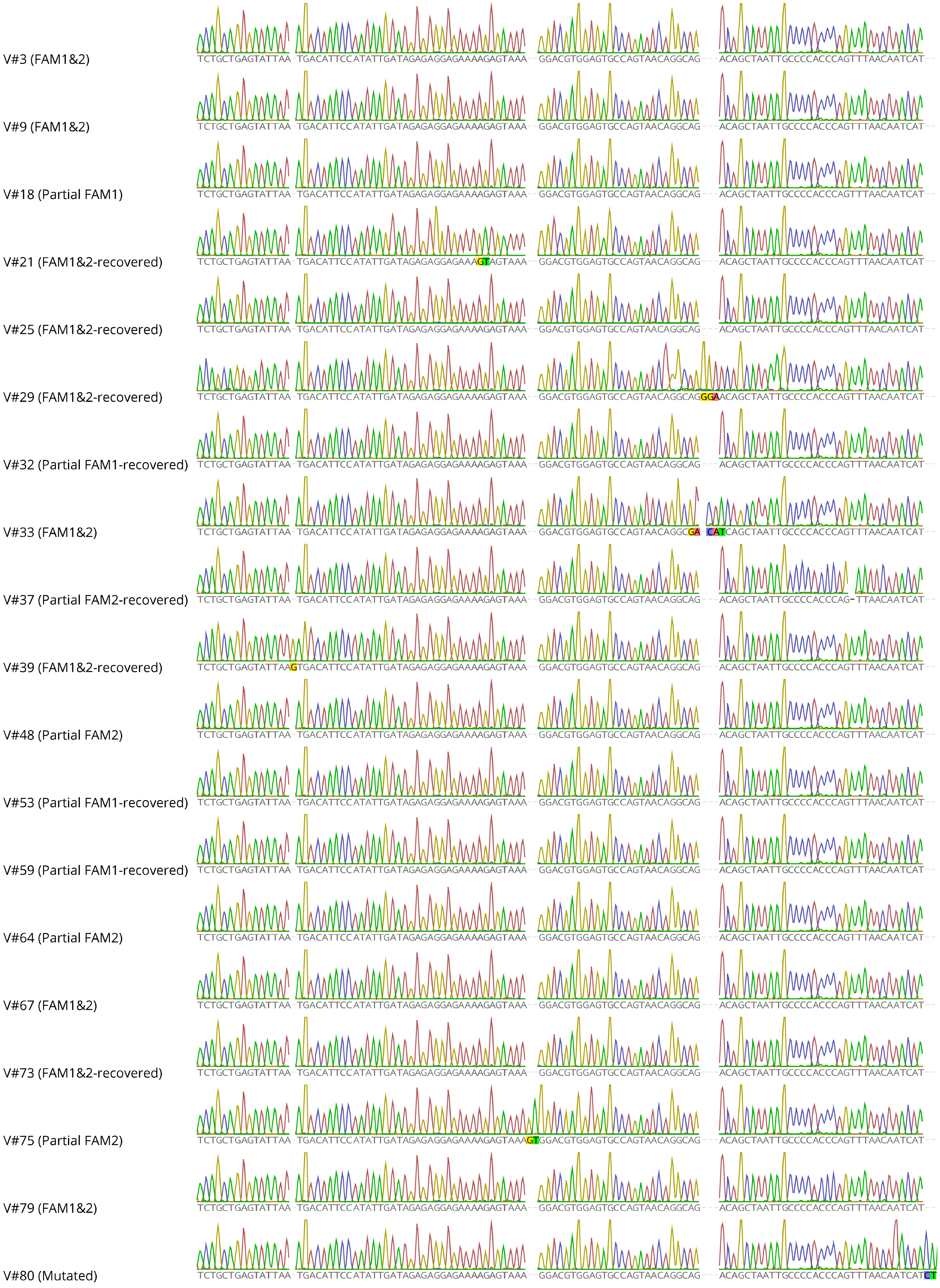
V#3 (FAM1&2)
V#9 (FAM1&2)
V#18 (Partial FAM1)
V#21 (FAM1&2-recovered)
V#25 (FAM1&2-recovered)
V#29 (FAM1&2-recovered)
V#32 (Partial FAM1-recovered)
V#33 (FAM1&2)
V#37 (Partial FAM2-recovered)
V#39 (FAM1&2-recovered)
V#48 (Partial FAM2)
V#53 (Partial FAM1-recovered)
V#59 (Partial FAM1-recovered)
V#64 (Partial FAM2)
V#67 (FAM1&2)
V#73 (FAM1&2-recovered)
V#75 (Partial FAM2)
V#79 (FAM1&2)
V#80 (Mutated)

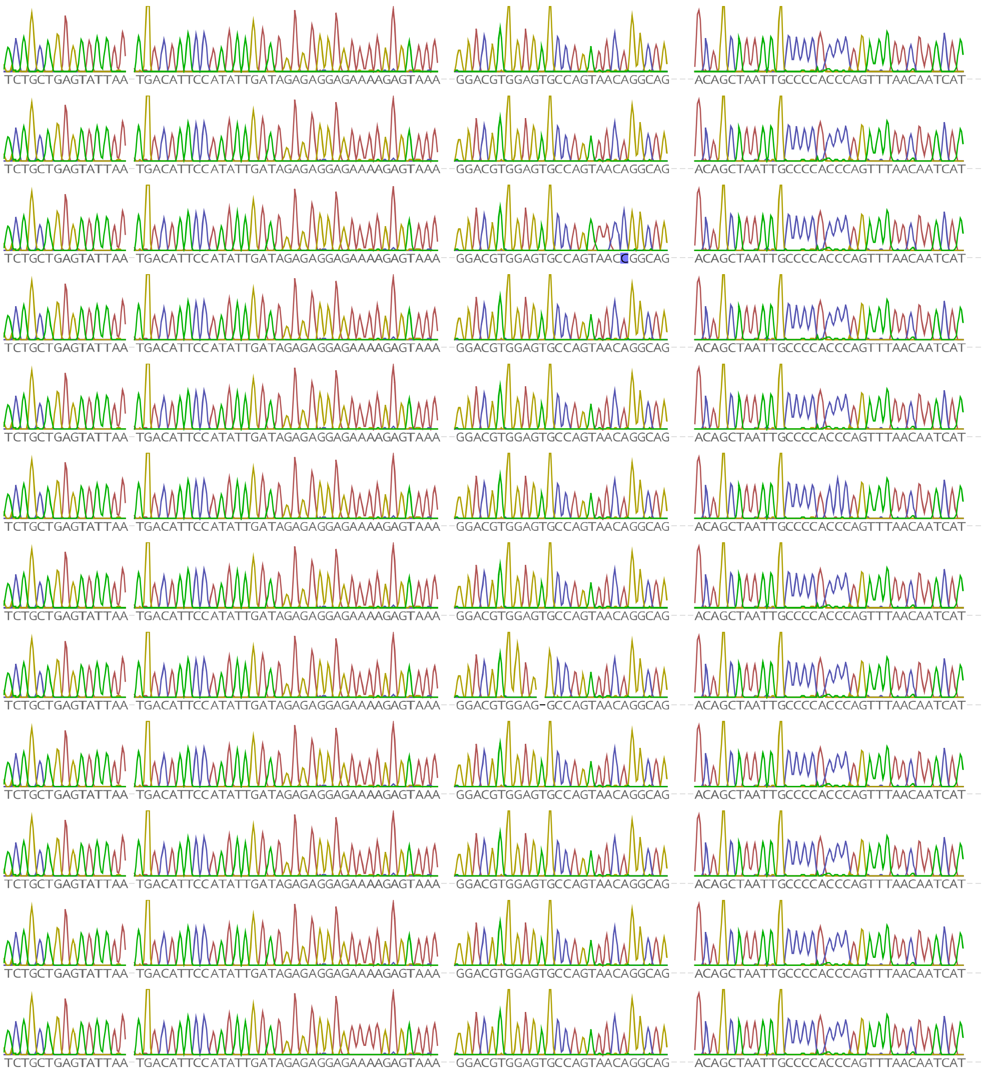


V#82 (Partial FAM1)
V#85 (Partial FAM2)
V#86 (Partial FAM1)
V#88 (FAM1&2-recovered)
V#89 (Partial FAM2)
V#91 (FAM1&2-recovered)
V#92 (FAM1&2)
V#94 (FAM1&2-recovered)
V#95 (Partial FAM1)
V#97 (Partial FAM1)
V#98 (Partial FAM2)
V#101 (FAM1&2)

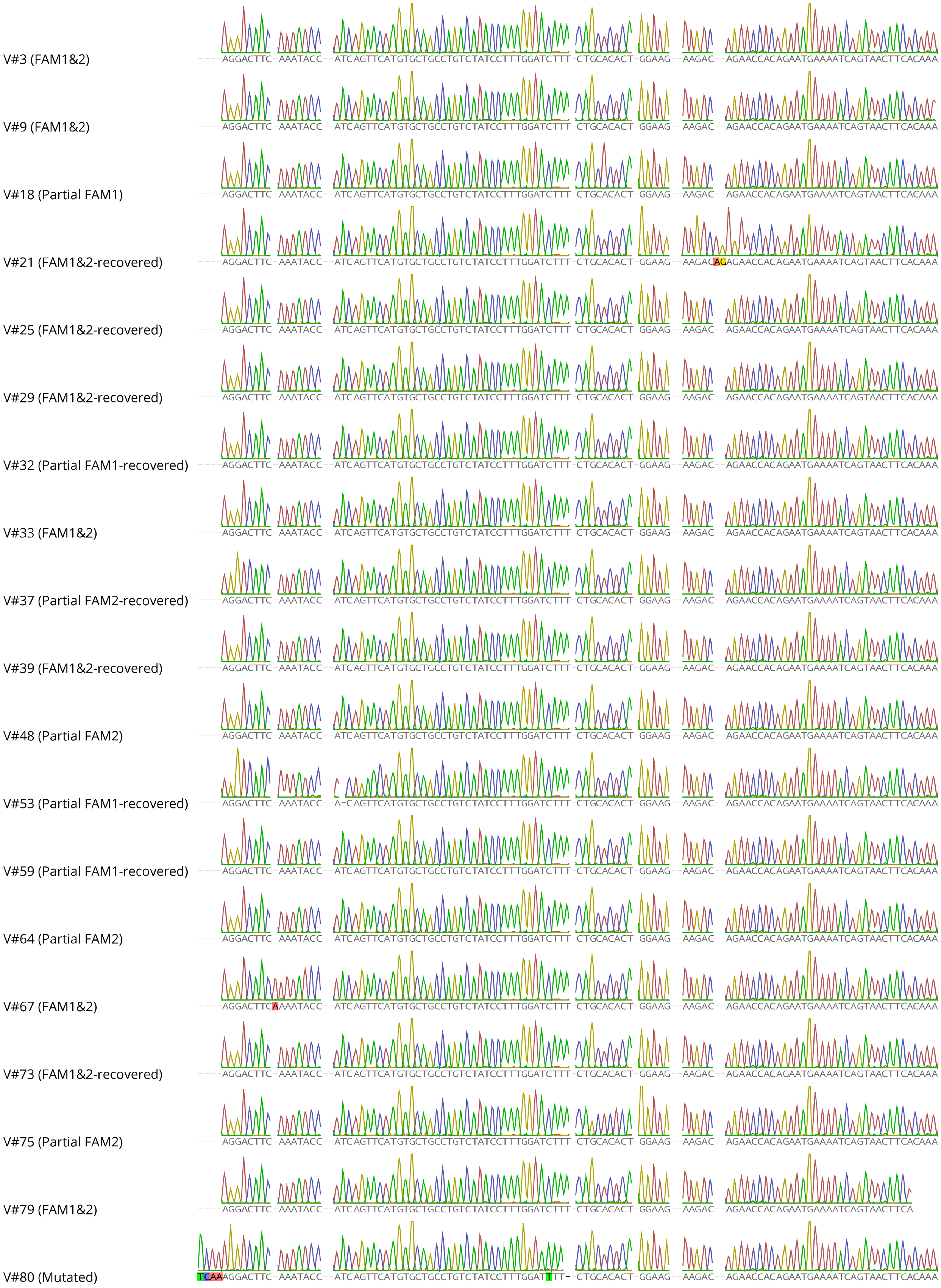
V#3 (FAM1&2)
V#9 (FAM1&2)
V#18 (Partial FAM1)
V#21 (FAM1&2-recovered)
V#25 (FAM1&2-recovered)
V#29 (FAM1&2-recovered)
V#32 (Partial FAM1-recovered)
V#33 (FAM1&2)
V#37 (Partial FAM2-recovered)
V#39 (FAM1&2-recovered)
V#48 (Partial FAM2)
V#53 (Partial FAM1-recovered)
V#59 (Partial FAM1-recovered)
V#64 (Partial FAM2)
V#67 (FAM1&2)
V#73 (FAM1&2-recovered)
V#75 (Partial FAM2)
V#79 (FAM1&2)
V#80 (Mutated)

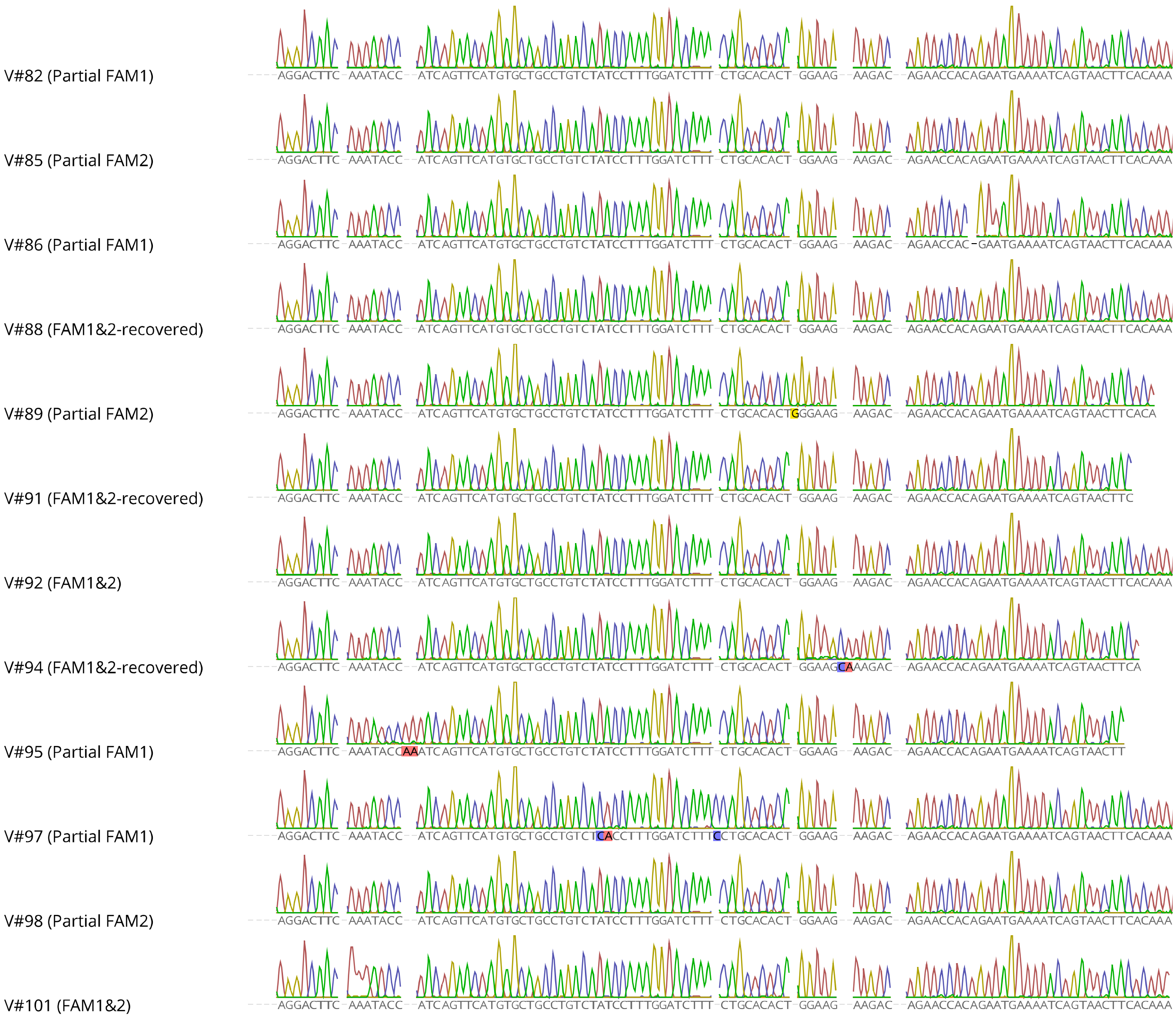
V#82 (Partial FAM1)
V#85 (Partial FAM2)
V#86 (Partial FAM1)
V#88 (FAM1&2-recovered)
V#89 (Partial FAM2)
V#91 (FAM1&2-recovered)
V#92 (FAM1&2)
V#94 (FAM1&2-recovered)
V#95 (Partial FAM1)
V#97 (Partial FAM1)
V#98 (Partial FAM2)
V#101 (FAM1&2)

V#3 (FAM1&2)

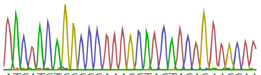


V#9 (FAM1&2)

V#18 (Partial FAM1)

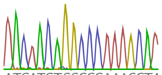


V#21 (FAM1&2-recovered)

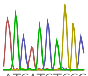


V#25 (FAM1&2-recovered)

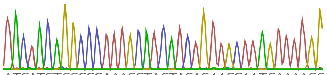


V#29 (FAM1&2-recovered)

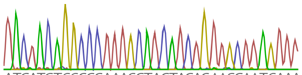


V#32 (Partial FAM1-recovered)

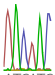


V#33 (FAM1&2)

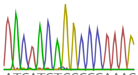


V#37 (Partial FAM2-recovered)

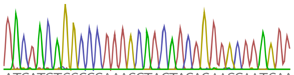


V#39 (FAM1&2-recovered)

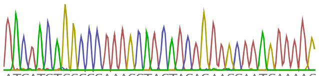


V#48 (Partial FAM2)

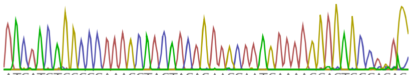


V#53 (Partial FAM1-recovered)

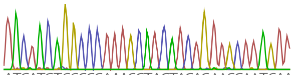


V#59 (Partial FAM1-recovered)

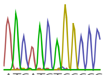


V#64 (Partial FAM2)

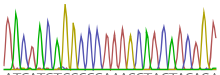


V#67 (FAM1&2)

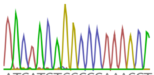


V#73 (FAM1&2-recovered)

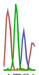


V#75 (Partial FAM2)




V#79 (FAM1&2)

V#80 (Mutated)

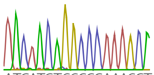

V#82 (Partial FAM1)

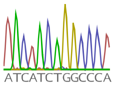


V#85 (Partial FAM2)

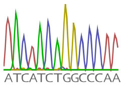


V#86 (Partial FAM1)

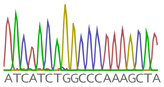


V#88 (FAM1&2-recovered)

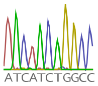


V#89 (Partial FAM2)

V#91 (FAM1&2-recovered)

V#92 (FAM1&2)

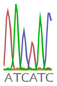


V#94 (FAM1&2-recovered)

V#95 (Partial FAM1)

V#97 (Partial FAM1)

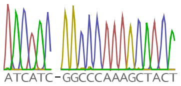


V#98 (Partial FAM2)

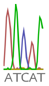


V#101 (FAM1&2)
