## Supplementary material for "Precise exogenous insertion and sequence replacements in poplar by simultaneous HDR overexpression and NHEJ suppression using CRISPR-Cas9": Suppl T.1 (from another version of this preprint: version 6, 2021-02-03)

| Name | Length | Direction | # Off-target Sites | Doench (2014) Activity Score | Protospacer Adjacent Motif (PAM) | Zhang (2013) Specificity Score | Target Sequence |
|---|---|---|---|---|---|---|---|
| CRISPR guide 3 | 23 | forward | 57 (1 in CDS) | 0.237 | AGG | 32.44% | AAGTTCCTCAAGAGTTTGAA |
| CRISPR guide 4 | 23 | reverse | 140 (1 in CDS) | 0.063 | TGG | 48.14% | TTATTATTTCTTTTTTTGTA |
| CRISPR guide 6 | 23 | reverse | 3 (0 in CDS) | 0.26 | AGG | 77.76% | ATAGAGAGCGTATGGTTTCG |
| CRISPR guide 7 | 23 | forward | 3 (0 in CDS) | 0.126 | TGG | 78.25% | TGGTTTGAAAACCTCAGCCC |
| CRISPR guide 11 | 23 | reverse | 7 (0 in CDS) | 0.067 | AGG | 81.71% | GTTATCTATTGAAATTTTGA |
| CRISPR guide 12 | 23 | reverse | 2 (0 in CDS) | 0.631 | TGG | 81.72% | TATTACCTCACACCGATCCA |
| CRISPR guide 13 | 23 | forward | 10 (0 in CDS) | 0.091 | CGG | 82.08% | ACAATAAAATCACTTTATTT |
| CRISPR guide 15 | 23 | reverse | 4 (1 in CDS) | 0.099 | AGG | 82.80% | GCATCAAACTCCTGTTTGAA |
| CRISPR guide 16 | 23 | forward | 3 (0 in CDS) | 0.217 | AGG | 82.94% | TATATTGACACCTTTCAAAC |
| CRISPR guide 20 | 23 | forward | 2 (0 in CDS) | 0.371 | CGG | 83.09% | TGGATCGGTGTGAGGTAATA |
| CRISPR guide 22 | 23 | forward | 21 (2 in CDS) | 0.112 | TGG | 83.15% | GAAAAGAGGCAAAAGAAGGG |
| CRISPR guide 25 | 23 | forward | 1 (0 in CDS) | 0.031 | AGG | 83.33% | ACAGTTAAGAATCTTTCTTC |
| CRISPR guide 27 | 23 | reverse | 1 (1 in CDS) | 0.122 | AGG | 83.33% | CTCTTTCTAATTTCAATACC |
| CRISPR guide 28 | 23 | forward | 1 (0 in CDS) | 0.289 | AGG | 83.33% | CTATTCCATGGATCGGTGTG |
| CRISPR guide 30 | 23 | reverse | 1 (0 in CDS) | 0.55 | AGG | 83.33% | AATTACAGTGACCAGGGCTG |
| CRISPR guide 31 | 23 | reverse | 1 (0 in CDS) | 0.107 | TGG | 83.33% | TCGAGGCTTTTGATTTGTGA |
| CRISPR guide 36 | 23 | forward | 2 (1 in CDS) | 0.137 | GGG | 84.30% | CCTCATACAAGTTCATGAAT |
| CRISPR guide 42 | 23 | forward | 1 (0 in CDS) | 0.139 | AGG | 89.21% | AAGAGACATCAGATCGTAAG |
| CRISPR guide 43 | 23 | forward | 14 (1 in CDS) | 0.449 | GGG | 89.43% | ATTGAAAAGAGGCAAAAGAA |
| CRISPR guide 44 | 23 | forward | 2 (2 in CDS) | 0.085 | TGG | 89.48% | TCCTCATACAAGTTCATGAA |
| CRISPR guide 48 | 23 | reverse | 4 (1 in CDS) | 0.052 | AGG | 91.10% | TTGATTTGTGATGGATGTCC |
| CRISPR guide 50 | 23 | reverse | 4 (0 in CDS) | 0.075 | AGG | 92.63% | TTTGACCTTTCAAACTCTTG |
| CRISPR guide 51 | 23 | forward | 8 (0 in CDS) | 0.081 | TGG | 92.73% | ATTTCAAGAAATGATGTGAA |
| CRISPR guide 52 | 23 | reverse | 3 (2 in CDS) | 0.104 | TGG | 92.76% | TCCATGGAATAGAGAGCGTA |
| CRISPR guide 54 | 23 | reverse | 8 (0 in CDS) | 0.215 | TGG | 93.33% | CATCATTTCTTGAAATCATT |
| CRISPR guide 55 | 23 | reverse | 2 (0 in CDS) | 0.027 | AGG | 93.40% | TTGCTGAAATTACAGTGACC |
| CRISPR guide 57 | 23 | forward | 9 (2 in CDS) | 0.057 | GGG | 93.66% | ATTCTTACTATTCTCCAATT |
| CRISPR guide 60 | 23 | forward | 4 (1 in CDS) | 0.515 | GGG | 94.19% | AGACTTACCACAAGTTTTAA |
| CRISPR guide 64 | 23 | reverse | 1 (1 in CDS) | 0.22 | CGG | 94.59% | GCGACATACTTTACAAATGC |
| CRISPR guide 65 | 23 | forward | 6 (0 in CDS) | 0.017 | TGG | 94.59% | TATACAGAAAACTATTTTCC |
| CRISPR guide 66 | 23 | forward | 6 (1 in CDS) | 0.363 | AGG | 94.63% | GTTTTGAACATATTGAAAAG |
| CRISPR guide 67 | 23 | forward | 3 (0 in CDS) | 0.13 | TGG | 94.67% | CACTTTTTGTGAATTGCTAA |
| CRISPR guide 72 | 23 | reverse | 6 (0 in CDS) | 0.006 | TGG | 95.21% | AAGTGATTTTATTGTTTGTC |
| CRISPR guide 77 | 23 | reverse | 2 (1 in CDS) | 0.279 | CGG | 95.85% | TGGAGAATAGTAAGAATTTG |
| CRISPR guide 78 | 23 | reverse | 3 (0 in CDS) | 0.111 | TGG | 95.98% | TTACGATCTGATGTCTCTTT |
| CRISPR guide 80 | 23 | forward | 1 (1 in CDS) | 0.268 | AGG | 96.24% | ATGTCGCGAGATCCACATGC |
| CRISPR guide 82 | 23 | reverse | 2 (1 in CDS) | 0.12 | GGG | 96.43% | ATCAGACTCTTGAGAGTTAT |
| CRISPR guide 87 | 23 | forward | 1 (1 in CDS) | 0.158 | TGG | 96.67% | AGTTCATGAATGGGTGTGCC |
| CRISPR guide 89 | 23 | reverse | 3 (1 in CDS) | 0.065 | TGG | 96.81% | TCTTTTGGGTGGCCACGAGG |
| CRISPR guide 90 | 23 | forward | 2 (0 in CDS) | 0.087 | TGG | 96.84% | ATGAATTCTCATAAGAGTTA |
| CRISPR guide 93 | 23 | reverse | 1 (0 in CDS) | 0.447 | GGG | 96.94% | TGCTGAAATTACAGTGACCA |
| CRISPR guide 94 | 23 | reverse | 3 (1 in CDS) | 0.223 | AGG | 96.99% | CCCATTCATGAACTTGTATG |
| CRISPR guide 95 | 23 | reverse | 3 (1 in CDS) | 0.133 | AGG | 97.09% | AGAATTTGCGGATCAAGAAA |
| CRISPR guide 96 | 23 | reverse | 1 (1 in CDS) | 0.15 | TGG | 97.12% | CTTGTATGAGGACCTGCATG |
| CRISPR guide 97 | 23 | forward | 9 (0 in CDS) | 0.196 | GGG | 97.19% | AAAAGAGGCAAAAGAAGGGT |
| CRISPR guide 100 | 23 | forward | 2 (2 in CDS) | 0.19 | TGG | 97.48% | ACCATACGCTCTCTATTCCA |
| CRISPR guide 102 | 23 | reverse | 1 (0 in CDS) | 0.374 | TGG | 97.55% | GATCTGATGTCTCTTTTGGG |
| CRISPR guide 104 | 23 | forward | 6 (0 in CDS) | 0.139 | TGG | 97.70% | ATTAAATGATTAATAAGACT |
| CRISPR guide 108 | 23 | forward | 5 (0 in CDS) | 0.278 | AGG | 98.08% | TATTGAAAAGAGGCAAAAGA |
| CRISPR guide 111 | 23 | reverse | 3 (2 in CDS) | 0.107 | TGG | 98.34% | AAACATGTGTATGCTATTAG |
| CRISPR guide 116 | 23 | forward | 3 (1 in CDS) | 0.113 | GGG | 98.59% | TTTCAAGAAATGATGTGAAT |
| CRISPR guide 117 | 23 | reverse | 1 (0 in CDS) | 0.224 | AGG | 98.73% | GTCTCTTTTGGGTGGCCACG |
| CRISPR guide 118 | 23 | reverse | 2 (0 in CDS) | 0.075 | GGG | 98.79% | TACGATCTGATGTCTCTTTT |
| CRISPR guide 120 | 23 | forward | 2 (0 in CDS) | 0.127 | TGG | 98.80% | AATTCTTACTATTCTCCAAT |
| CRISPR guide 123 | 23 | forward | 3 (0 in CDS) | 0.159 | CGG | 98.98% | AAGGGTTGCAAGAGGAGAGC |
| CRISPR guide 125 | 23 | forward | 1 (0 in CDS) | 0.26 | CGG | 99.06% | ACGCTCTCTATTCCATGGAT |
| CRISPR guide 126 | 23 | forward | 1 (1 in CDS) | 0.103 | AGG | 99.13% | TGCATTAATTGCATAGACAC |
| CRISPR guide 137 | 23 | forward | 2 (0 in CDS) | 0.308 | TGG | 99.41% | TCTGCACTTGTCAAACACAA |
| CRISPR guide 138 | 23 | forward | 1 (0 in CDS) | 0.401 | TGG | 99.44% | GTCAGGGAAAAACCACCTCG |
| CRISPR guide 139 | 23 | forward | 1 (0 in CDS) | 0.403 | AGG | 99.52% | CAAGTTTTAAGGGTTGCAAG |
| CRISPR guide 140 | 23 | reverse | 2 (0 in CDS) | 0.02 | TGG | 99.53% | AAGATTCTTAACTGTCATTG |
| CRISPR guide 148 | 23 | forward | 1 (0 in CDS) | 0.238 | TGG | 99.78% | CATAAGAGTTATGGTCGACA |
| CRISPR guide 162 | 23 | forward | 0 (0 in CDS) | 0.072 | AGG | 100.00% | TCTCCAATTGGGAATCAGTC |
| CRISPR guide 163 | 23 | forward | 0 (0 in CDS) | 0.175 | GGG | 100.00% | CTCCAATTGGGAATCAGTCA |
| CRISPR guide 164 | 23 | forward | 0 (0 in CDS) | 0.014 | AGG | 100.00% | TAGACTTACCACAAGTTTTA |
| CRISPR guide 179 | 23 | reverse | 0 (0 in CDS) | 0.084 | TGG | 100.00% | ATGCTATTAGTGGACGCTCA |
| CRISPR guide 180 | 23 | reverse | 0 (0 in CDS) | 0.243 | TGG | 100.00% | CTTGCAACCCTTAAAACTTG |
| CRISPR guide 181 | 23 | reverse | 0 (0 in CDS) | 0.069 | TGG | 100.00% | TATCAGACTCTTGAGAGTTA |
| CRISPR guide 182 | 23 | reverse | 0 (0 in CDS) | 0.241 | TGG | 100.00% | TTCCCTGACTGATTCCCAAT |

**Supplementary Table 1:** CRISPR sites located on 3' region of *MKK2*. Yellow highlight reveals the selected CRISPR target in this study.
