## Supplementary material for "Precise exogenous insertion and sequence replacements in poplar by simultaneous HDR overexpression and NHEJ suppression using CRISPR-Cas9": Suppl T.2 (from another version of this preprint: version 6, 2021-02-03)

| Product name | Primers | Oligos | Sequence with the extension (5' to 3') | Tm |
|---|---|---|---|---|
| DDT | Os2-R | - | CCTGCAGGAAGGAATCTTTAAACATACGAACAGATC | 58.4 |
| | Os2-F | - | CCATGGAAAAAAGCACCGACTCGGTGC | 62.3 |
| | Os1-R | - | GGATCCAAAAAAGCACCGACTCGGTG | 59.3 |
| | Os1-F | - | AAGCTTAAGGAATCTTTAAACATACGAACAG | 55.9 |
| | BleoR-overlap3-R | - | TTTTTTTTTTTTTTTTTTTTTTTTTTTTTTATG | 53.4 |
| | BleoR-overlap3-F | - | CTTGCAACCCTTAAAACTTG | 53.2 |
| | BleoR-overlap2-R | - | TTTTTTTTTTTTTTTTTTATGATGATGA | 52.5 |
| | BleoR-overlap2-F | - | CCCTTAAAACTTGTGTTCTTGTGAAGA | 60.4 |
| | BleoR-overlap1-R | - | ATGATGATGATGTCAGTCCTGC | 58.3 |
| | BleoR-overlap1-F | - | CTTGTGAAGATGTAGATGGCCAAGT | 61.1 |
| | BleoR - 2,276 R | - | TCAGTCCTGCTCCTCGGC | 61.1 |
| | BleoR - 1,902 F | - | ATGGCCAAGTTGACCAGTGC | 61.2 |
| | 5' Ho-R-2 | - | CTACATCTTCACAAGAACACAAGTTTTAAGGGTTGCAAGAGGAGAGCCGGCATTTGTAA | 59.4 |
| | 5' Ho-R-1 | - | AGGAGAGCCGGCATTTGTAA | 59.4 |
| | 5' Ho-F-2 | - | GGATCCGCACCCGAGGAGGTGGGTTACTGATTTTCATTCTGTGGTTC | 56.9 |
| | 5' Ho-F-1 | - | GTTACTGATTTTCATTCTGTGGTTC | 56.9 |
| | 3' Ho-R-2 | - | CCATGGGCACCCGAGGAGGTGGTTTTGTGAATTGCTAATGGTTTGAA | 56.9 |
| | 3' Ho-R-1 | - | TTTTGTGAATTGCTAATGGTTTGAA | 56.9 |
| | 3' Ho-F-2 | - | TTTTTTTTTTTTTTTTTTTTTTTTTTTTTTTAAGTCTAATTTTCCCATTCACATC | 55.4 |
| | 3' Ho-F-1 | - | TAAGTCTAATTTTCCCATTCACATC | 55.4 |
| | - | Special gRNA oligo2-R | AAACCCACCTCCTCGGGTGC | |
| | - | Special gRNA oligo2-F | GGCAGCACCCGAGGAGGTGG | |
| | - | Special gRNA oligo1-R | AAACCCACCTCCTCGGGTGC | |
| | - | Special gRNA oligo1-F | GGCAGCACCCGAGGAGGTGG | |
| pgRNA | - | MKK2 Oligo-F | GGCACTTGCAACCCTTAAAACTTGTGG | |
| | - | MKK2 Oligo-R | AAACCCACAAGTTTTAAGGGTTGCAAG | |
| pgCtIP | Cam-F | - | TGAGACTTTTCAACAAAGGGTAATA | 56.6 |
| | Cam-R | - | TGTTCTCTCCAAATGAAATGAACTTC | 58.3 |
| | Cas-F | - | GACAAGAAGTACAGCATCGGC | 59.3 |
| | Cas-R | - | GTCGCCTCCCAGCTGAGA | 61.1 |
| | CtIP-F | - | ATGAAGGGCGACCTGGAG | 58.7 |
| | CtIP-R | - | TCACATTTCAGATTCAAATCCAATA | 55.3 |
| | Flag-F | - | ATGGACTATAAGGACCACGAC | 56.6 |
| | Flag-R | - | CTTATCGTCATCGTCTTTGTAATCA | 57.3 |
| | PolyA-F | - | TATGGGCACCAAAGAACCTG | 57.8 |
| | PolyA-R | - | CTACCTCGAGTTTTTTTTTTTT | 51.8 |
| pgMR | Cam-F | - | CCTGCAGGTGAGACTTTTCAACAAAGGGTAATA | 56.6 |
| | Cam-R | - | CTCGAGTGTTCTCTCCAAATGAAATGAACTTC | 58.3 |
| | Cas-F | - | CATATGGACAAGAAGTACAGCATCGGC | 59.3 |
| | Cas-R | - | GGATCCGTCGCCTCCCAGCTGAGA | 61.1 |
| | Flag-F | - | GGATCCATGAAGGGCGACCTGGAG | 56.6 |
| | Flag-R2 | - | AGGCCTTCACATTTCAGATTCAAATCCAATA | 57.3 |
| | MRE-F | - | CTCGAGATGGACTATAAGGACCACGAC | 56.1 |
| | MRE-R | - | GCGGCCGCCTTATCGTCATCGTCTTTGTAATCA | 58 |
| | PolyA-R | - | GCGGCCGCATGGAGAAGGATGAGGTACG | 51.8 |
| | PolyA2-F | - | CATATGTCAGCAATTCTTCAAACAAGGATC | 57.8 |
| pgCtMR | Cam-F | - | CCTGCAGGTGAGACTTTTCAACAAAGGGTAATA | 56.6 |
| | Cam-R | - | CTCGAGTGTTCTCTCCAAATGAAATGAACTTC | 58.3 |
| | Cas-F2 | - | CATATGGACAAGAAGTACAGCATCGGC | 59.3 |
| | Cas-R | - | GGATCCGTCGCCTCCCAGCTGAGA | 61.1 |
| | CtIP-F | - | GGATCCATGAAGGGCGACCTGGAG | 58.7 |
| | CtIP-R | - | AGGCCTTCACATTTCAGATTCAAATCCAATA | 55.3 |
| | Flag-F | - | CTCGAGATGGACTATAAGGACCACGAC | 56.6 |
| | Flag-R2 | - | GCGGCCGCCTTATCGTCATCGTCTTTGTAATCA | 57.3 |
| | MRE-F | - | GCGGCCGCATGGAGAAGGATGAGGTACG | 56.1 |
| | MRE-R | - | CATATGTCAGCAATTCTTCAAACAAGGATC | 58 |
| | PolyA-F | - | AGGCCTTATGGGCACCAAAGAACCTG | 57.8 |
| | PolyA-R | - | GTTTAAACCTACCTCGAGTTTTTTTTTTTT | 51.8 |
| XXRC4-cassette | XR-Cass1-F | - | AAGGAATCTTTAAACATACGAACAG | 55.9 |
| | XR-Cass1-R | - | AAAAAAGCACCGACTCGGTG | 59.3 |
| | XR-Cass2-F | - | AAGGAATCTTTAAACATACGAACAG | 55.9 |
| | XR-Cass2-R | - | AAAAAAGCACCGACTCGGTG | 59.3 |
| | - | XRCC4-Oligo1 | TTCAATCTCTCCATCACCGACGG | |
| | - | XRCC4-Oligo2 | CCGTCGGTGATGGAGAGATTGAA | |
| pggCtMR | Cam-F | - | CCTGCAGGTTTTTTAAAAAATGATCATGAGACTTTTCAACAAAGGGTAATA | 56.6 |
| | Cam-R | - | CTCGAGTGTTCTCTCCAAATGAAATGAACTTC | 58.3 |
| | Cas-F2 | - | CATATGGACAAGAAGTACAGCATCGGC | 59.3 |
| | Cas-R | - | GGATCCGTCGCCTCCCAGCTGAGA | 61.1 |
| | CtIP-F | - | GGATCCATGAAGGGCGACCTGGAG | 58.7 |
| | CtIP-R | - | AGGCCTTCACATTTCAGATTCAAATCCAATA | 55.3 |
| | Flag-F | - | CTCGAGATGGACTATAAGGACCACGAC | 56.6 |
| | Flag-R2 | - | GCGGCCGCCTTATCGTCATCGTCTTTGTAATCA | 57.3 |
| | MRE-F | - | GCGGCCGCATGGAGAAGGATGAGGTACG | 56.1 |
| | MRE-R | - | CATATGTCAGCAATTCTTCAAACAAGGATC | 58 |
| | PolyA-F | - | AGGCCTTATGGGCACCAAAGAACCTG | 57.8 |
| | PolyA-R | - | GTTTAAACCTACCTCGAGTTTTTTTTTTTT | 51.8 |
| TaqMan real-time PCR | FAM1-F | - | GCACACCCATTCATGAACTTGTAT | 59.8 |
| | FAM1-R | - | GGTGAGCACCGGAACGGC | 63.7 |
| | FAM2-F | - | CCGGCCGGCAACTGCGTG | 67.3 |
| | FAM2-R | - | AAAAAACATTTGACCAAATGATTTCA | 56 |
| | VIC F | - | ATTTCCCGTTCAGCAGTGGT | 59.9 |
| | VIC R | - | ACGCCATCCTTCGTTTGGAT | 60 |
| HDR efficiency | BleoR - 52 F | - | GGAGCGGTCGAGTTCTGG | 59.8 |
| | BleoR - 151 R | - | TGATGAACAGGGTCACGTCG | 60 |
| RT-qPCR | 2,817 F-MKK2 | - | TCTGCTAGAGATCCACCATCCA | 60.1 |
| | 2,968 R-MKK2 | - | ATTGAAGAGCCTGCACGCAA | 60.9 |
| | 3,260 F-XRCC4 | - | GGGATCCCAACCAAGTTCGT | 60 |
| | 3,438 R-XRCC4 | - | TGCGACCTGGGACTCATTTC | 60 |
| | 7 F-BleoR | - | AAGTTGACCAGTGCCGTTCC | 60.8 |
| | 7,831 F-MRE11 | - | AGCCGTTCATCCCGAATGTT | 60 |
| | 8,025 R-MRE11 | - | ATGGCACTTGAGGGTTCTGG | 60 |
| | 11,007 F-Lig4 | - | GCTCCTCAATCGTCAACCCA | 60 |
| | 194 R-BleoR | - | CAGGCCAGGGTGTTGTCC | 60.3 |
| | 242 F-Rad50 | - | CTTACTCTCATCGTCGGCCC | 60 |
| | 244 F-BRCA1 | - | GGTCAGGCACCAGGTACTTC | 60 |
| | 410 R-Rad50 | - | ACCTAGCATTCGGAGGCAAC | 60.1 |
| | 422 R-BRCA1 | - | CTCGCCGCATAGGACATGAA | 60.2 |
| | 557 F-Rad51 | - | GTGGTGGCCTAGGAGGAAAG | 59.7 |
| | 643 F-CtIP | - | GGGCGTGGAAAGAGAAGGAA | 60 |
| | 723 R-Rad51 | - | TTTCTTGAGCACGCTCTCCC | 60.3 |
| | 828 R-CtIP | - | TCACAACCTCCTGTTCCAGC | 59.9 |
| | 9860 R-Lig4 | - | GCAATTTCTCTCTCGTTGCTGC | 60.7 |
| Sequencing | MKK2-S-7F | - | GCATTTGCAACACCAGACAA | 57.8 |
| | MKK2-S-1,139 R | - | TGACTGAGAACTCAATCTGCACT | 59.7 |
| Southern blotting | S-F | - | CCAGGGTGTTGTCCGGCA | 60.1 |
| | S-R | - | CTCACCGCGCGCGACGTC | 62.3 |
| RT-PCR | RT-F | - | TTCACAAGAACACAAGTTTTAAGGG | 58.1 |
| | RT-R | - | CCACCTCCAATAACGCCTTC | 59.2 |
| | RT-F-107 | - | GAGGCGTCCCGGAAGTTC | 60.5 |
| | Rt-R-519 | - | AAGGATAGACAGGCAGCACA | 57.3 |
| | Actin-F | - | ATTTCCCGTTCAGCAGTGGT | 57.3 |
| | Actin-R | - | ACGCCATCCTTCGTTTGGAT | 57.3 |

**Supplementary Table 2:** Oligos and primers used in this study.
