## Supplementary material for "Precise exogenous insertion and sequence replacements in poplar by simultaneous HDR overexpression and NHEJ suppression using CRISPR-Cas9": Suppl T.3 (from another version of this preprint: version 6, 2021-02-03)

| | | Experiment | | | | |
|---|---|---|---|---|---|---|
| | | I | II | III | IV | V |
| **5' homology arm** | **Deletion** | 105 | 19 | 17 | 19 | 5 |
| | **Del (tandem repeat)** | 2 | 2 | 1 | 4 | 2 |
| | **Insertion** | 53 | 34 | 29 | 10 | 11 |
| | **Ins (tandem repeat)** | 7 | 2 | 0 | 4 | 2 |
| | **SNP (transition)** | 4 | 1 | 4 | 5 | 1 |
| | **SNP (transversion)** | 3 | 7 | 0 | 10 | 3 |
| | **Substitution** | 8 | 0 | 2 | 2 | 3 |
| **3' homology arm** | **Deletion** | 32 | 77 | 57 | 17 | 10 |
| | **Del (tandem repeat)** | 3 | 1 | 1 | 4 | 5 |
| | **Insertion** | 16 | 22 | 24 | 13 | 18 |
| | **Ins (tandem repeat)** | 11 | 1 | 5 | 5 | 2 |
| | **SNP (transition)** | 1 | 2 | 7 | 4 | 1 |
| | **SNP (transversion)** | 6 | 6 | 1 | 14 | 1 |
| | **Substitution** | 0 | 1 | 2 | 0 | 2 |

Supplementary Table 3: All polymorphisms detected in homology arms happened by HDR through experiments.
