## Supplementary material for "Precise exogenous insertion and sequence replacements in poplar by simultaneous HDR overexpression and NHEJ suppression using CRISPR-Cas9": Suppl T.4 (from another version of this preprint: version 6, 2021-02-03)

### Supplemntary Table 4 : Variant nucleotides happened from experiment I

| Variant events | Polymorphism type | Variant genotyping | Protein effect | Nucleotide genotyping |
|---|---|---|---|---|
| I#11 | T | Deletion | Frame Shift | CCG |
| I#11 | T | Deletion | Frame Shift | A--------CAAGAAC |
| I#11 | T | Insertion (tandem repeat) | | - |
| I#11 | T | SNP (transition) | | C |
| I#11 | T | Insertion (tandem repeat) | | - |
| I#11 | G | Insertion | | - |
| I#11 | G | SNP | None | --------G |
| I#11 | G | Insertion | | - |
| I#11 | C | Deletion | Frame Shift | GCCGATCTCGGTCATGGCCGGCCGGAGGCGTCCCG GAAGTTCGTGGACACGACCTCCGACCACTC |
| I#11 | C | Deletion | Frame Shift | TC |
| I#11 | C | Deletion | Frame Shift | TC |
| I#11 | C | Insertion | | - |
| I#11 | C | Insertion | | - |
| I#11 | C | SNP (transversion) | | A |
| I#11 | C | Insertion | | - |
| I#11 | A | Deletion | Truncation | GTCCTCCCGAAGTCCCGG-GA |
| I#11 | A | Insertion | | - |
| I#11 | A | Insertion | | - |
| I#11 | A | SNP (transversion) | | T |
| I#11 | A | Insertion | | - |
| I#11 | A | Insertion | | - |
| I#11 | | Deletion | | A |
| I#11 | | Deletion | | C |
| I#11 | | Deletion | Frame Shift | CCCA |
| I#11 | | Deletion | Frame Shift | C |
| I#11 |  | Deletion | Frame Shift | GTACAGCTCG---------T- CCAGGCCGCGCACCCACACCCAGGCCAGGGTGTTG TCCGGCACCACCTGGTCCTGGACCGCGCTGATGAA CA |
| I#11 | | Deletion | Frame Shift | AT |
| I#11 | | Deletion | Frame Shift | AT |
| I#11 | | Deletion | Frame Shift | C |
| I#11 | | Deletion | Frame Shift | C |
| I#11, I#19 | A | Insertion | Frame Shift | - |
| I#11, I#19, I#23 | | Deletion | Frame Shift | A |
| I#11, I#19, I#23, I#27, I#35, I#4, I#41, I#5, I#7 | C | Insertion | | - |
| I#11, I#19, I#23, I#27, I#35, I#4, I#5, I#7 | | Deletion | | C |
| I#11, I#19, I#23, I#27, I#4, I#5 | A | Insertion | | - |
| I#11, I#19, I#23, I#27, I#5 | | Deletion | | T |
| I#11, I#19, I#23, I#35, I#4, I#41, I#5, I#7 | | Deletion | | A |
| I#11, I#19, I#27 | | Deletion | Deletion | ACAAGTTTTAAGGGTTGCAAG |
| I#11, I#19, I#27 | | Deletion | Deletion | GCC |
| I#11, I#19, I#27 | | Deletion | Deletion | GCC |
| I#11, I#19, I#27 | | Deletion | Deletion | GGC |
| I#11, I#19, I#27 | | Deletion | Deletion | GGC |
| I#11, I#19, I#27 | | Deletion | Deletion | CAC |
| I#11, I#19, I#27 | | Deletion | Deletion | CAC |
| I#11, I#19, I#27 | | Deletion | Deletion | ATG |
| I#11, I#19, I#27 | | Deletion | Deletion | ATG |
| I#11, I#19, I#27 | | Deletion | Deletion | CAG |
| I#11, I#19, I#27 | | Deletion | Deletion | CAG |
| I#11, I#19, I#27 | | Deletion | Deletion | GTC |
| I#11, I#19, I#27 | | Deletion | Deletion | GTC |
| I#11, I#19, I#27, I#35 | | Deletion | Deletion | AGG |
| I#11, I#19, I#27, I#35 | | Deletion | Deletion | AGG |
| I#11, I#19, I#27, I#35 | | Deletion | Deletion | AGA |
| I#11, I#19, I#27, I#35 | | Deletion | Deletion | AGA |
| I#11, I#19, I#35 | | Deletion (tandem repeat) | | A |
| I#11, I#19, I#35, I#41 | T | SNP | None | -------T |
| I#11, I#23 | TA | Insertion (tandem repeat) | | -- |
| I#11, I#23 | T | Insertion (tandem repeat) | | - |
| I#11, I#23 | C | Insertion | | - |
| I#11, I#23 | A | Insertion | | - |
| I#11, I#23, I#35 | | Deletion | | AATAAGACTTGGATT |
| I#11, I#23, I#35, I#4, I#41, I#5 | | Deletion | | C |
| I#11, I#27 | | Deletion | Deletion | GGTCACGTCGTCCCGGACCACACCGGCGAAGTC |
| I#11, I#27, I#35 | | Deletion | Deletion | CGGCTGCTC |
| I#11, I#27, I#35, I#5 | | Deletion (tandem repeat) | | TTTTTT |
| I#11, I#27, I#35, I#5 | | Deletion | | TTTTTTTTTATGATGATGATGATGATG |
| I#11, I#27, I#35, I#5 | | Deletion | Extension | TCAGTCCTGCTCCTCGGCCACGAAGTGCACGCAGT TGCCGGCCGGGTCGCGCAGGGCGAA |
| I#11, I#35 | | Deletion | | A |
| I#11, I#35 | | Deletion | | T |

| | | | | |
|---|---|---|---|---|
| I#11, I#35 | | Deletion | | GCAACTACATCTTCACAAGAACTCCATACAAAAAAGAAATAATAA |
| I#11, I#35 | | Deletion (tandem repeat) | | A |
| I#11, I#35 | | Deletion | | AAAA |
| I#11, I#35 | | Deletion | | A |
| I#11, I#35 | | Deletion | | TTGA |
| I#11, I#35 | | Deletion | | AA |
| I#11, I#35 | | Deletion | | ATGATTTCAAGAAATGATGTGAATGGGAAAATTAGACTTA |
| I#11, I#35, I#4 | | Deletion (tandem repeat) | | T |
| I#11, I#35, I#5 | | Deletion | | C |
| I#11, I#35, I#5 | | Deletion (tandem repeat) | | TTTTTTTTTTTTTTT |
| I#11, I#35, I#5 | | Deletion | Deletion | CTC |
| I#11, I#4, I#5 | C | Insertion | | - |
| I#11, I#41 | A | Insertion | | - |
| I#19 | T | SNP | None | -T |
| I#19 | T | SNP (transversion) | None | G |
| I#19 | T | SNP (transversion) | None | G |
| I#19 | T | Deletion | Frame Shift | AA |
| I#19 | T | Deletion | Frame Shift | AA |
| I#19 | G | Deletion | Extension | CTACATCTTCA--------CAAGAAC |
| I#19 | G | Insertion | | - |
| I#19 | A | Insertion (tandem repeat) | | - |
| I#19 | A | Insertion | | - |
| I#19 | | Deletion | | TAAATTC |
| I#19 | | Deletion | Frame Shift | GTCACGTCGTCCCGGACCACACCGGCGAAGTC |
| I#19 | | Deletion | Frame Shift | CG |
| I#19 | | Deletion | Frame Shift | CG |
| I#19 | | Deletion | | ATAGC |
| I#19 | | Deletion | | C |
| I#19 | | Deletion | | A |
| I#19 | | Deletion | | CAT |
| I#19 | | Deletion | | TGA |
| I#19 | | Deletion | | A |
| I#19 | | Deletion (tandem repeat) | | A |
| I#19, I#23, I#27, I#35, I#4, I#41, I#5, I#7 | | Deletion | | C |
| I#19, I#23, I#27, I#35, I#41 | C | Insertion | | - |
| I#19, I#27 | | Deletion | Deletion | ATT |
| I#19, I#27 | | Deletion | Deletion | ATT |
| I#19, I#27 | | Deletion | Deletion | TGT |
| I#19, I#27 | | Deletion | Deletion | TGT |
| I#19, I#27 | | Deletion | Deletion | AAA |
| I#19, I#27 | | Deletion | Deletion | AAA |
| I#19, I#27 | | Deletion | Deletion | GTA |
| I#19, I#27 | | Deletion | Deletion | GTA |
| I#19, I#27 | | Deletion | Deletion | TGT |
| I#19, I#27 | | Deletion | Deletion | TGT |
| I#19, I#27 | | Deletion | Deletion | ATC |
| I#19, I#27 | | Deletion | Deletion | ATC |
| I#19, I#27 | | Deletion | Deletion | CTC |
| I#19, I#27 | | Deletion | Deletion | CTC |
| I#19, I#27 | | Deletion | | AAATTAGAA |
| I#19, I#27 | | Deletion | | AGAAAAA |
| I#19, I#27 | | Deletion | | AGAGTCAGATCCATG |
| I#19, I#27 | | Deletion | | TCCACTA |
| I#19, I#27 | | Deletion | | ATA |
| I#19, I#27 | | Deletion | | TTT |
| I#19, I#27 | | Deletion | | CATAT |
| I#19, I#27 | | Deletion | | GA |
| I#19, I#27, I#4 | | Deletion | Deletion | GAA |
| I#19, I#27, I#4 | | Deletion | Deletion | GAA |
| I#19, I#27, I#4, I#41 | | Deletion | | A |
| I#19, I#27, I#4, I#5 | | Deletion | Deletion | TGG |
| I#19, I#27, I#4, I#5 | | Deletion | Deletion | TGG |
| I#19, I#27, I#4, I#5 | | Deletion | Deletion | GTG |
| I#19, I#27, I#4, I#5 | | Deletion | Deletion | GTG |
| I#19, I#27, I#4, I#5 | | Deletion | Deletion | TGC |
| I#19, I#27, I#4, I#5 | | Deletion | Deletion | TGC |
| I#19, I#27, I#4, I#5 | | Deletion | | C |
| I#19, I#27, I#5 | | Deletion | Deletion | GTCCTCCACGAAGTCCCGG-GA |
| I#19, I#27, I#5 |  | Deletion | Deletion | GAACCCG------AGCCGGTCGGTCCAGAACTCGACC---------GCTCCGGCGACGTCGCGCGCGGTGAG |
| I#19, I#27, I#5 | | Deletion | Start Codon Loss | CACCGGAACGGCACTGGTCAACTTGGCCAT |
| I#19, I#27, I#5 | | Deletion | Deletion | CAA |
| I#19, I#27, I#5 | | Deletion | Deletion | CAA |
| I#19, I#27, I#5 | | Deletion | Deletion | GTT |
| I#19, I#27, I#5 | | Deletion | Deletion | GTT |
| I#19, I#27, I#5 | | Deletion | Deletion | CAT |

| | | | | |
|---|---|---|---|---|
| I#19, I#27, I#5 | | Deletion | Deletion | CAT |
| I#19, I#27, I#5 | | Deletion | | TATTG |
| I#19, I#35 | | Deletion | | AACTCTAAAG |
| I#19, I#4, I#5 | | Deletion | | TG |
| I#19, I#5 | | Deletion | | G |
| I#23 | T | Deletion | Extension | CTACATCTTCA--------CAAGAAC |
| I#23 | T | Insertion | | - |
| I#23 | T | Insertion (tandem repeat) | | - |
| I#23 | GTCCTG | Insertion |  | ------ |
| I#23 | G | SNP | None | ---------G |
| I#23 | G | Insertion | | - |
| I#23 | G | Insertion | | - |
| I#23 | G | Insertion | | - |
| I#23 | CCGG | Insertion | Frame Shift | ---- |
| I#23 | CCGG | Insertion | Frame Shift | ---- |
| I#23 | C | SNP | None | ------C |
| I#23 | C | Insertion | | - |
| I#23 | ACATTCCCCCT | Insertion | Frame Shift | ---------T- |
| I#23 | A | Insertion | | - |
| I#23 | A | Insertion (tandem repeat) | | - |
| I#23 | A | SNP (transversion) | | T |
| I#23 | A | Insertion | | - |
| I#23 | A | Insertion | Frame Shift | - |
| I#23 | | Deletion | | G |
| I#23 | | Deletion | Frame Shift | ACAAGTTTTAAGGGTT |
| I#23, I#27 | T | SNP | None | ------T |
| I#23, I#5 | T | Insertion (tandem repeat) | | - |
| I#23, I#7 | C | Insertion (tandem repeat) | | - |
| I#27 | TCCG | Insertion |  | ---- |
| I#27 | GGGCGAA | Insertion |  | ------- |
| I#27 | G | Insertion | | - |
| I#27 | G | Insertion | | - |
| I#27 | G | SNP (transition) | | A |
| I#27 | C | Insertion (tandem repeat) | | - |
| I#27 | C | Insertion (tandem repeat) | | - |
| I#27 | C | SNP (transversion) | | A |
| I#27 | C | SNP (transversion) | | A |
| I#27 | C | Insertion | | - |
| I#27 | C | Insertion | Frame Shift | - |
| I#27 | AT | Insertion | Frame Shift | --------- |
| I#27 | A | Insertion (tandem repeat) | | - |
| I#27 | A | Insertion | | - |
| I#27 | A | Insertion | | - |
| I#27 | A | Insertion | | - |
| I#27 | A | Insertion | | - |
| I#27 | A | SNP (transversion) | | C |
| I#27 | A | Deletion | Extension | CTACATCTTCCAAGAAC |
| I#27 | | Deletion | Frame Shift | T-<br>CCAGGCCGCGCACCCACACCCAGGCCAGGGTGTTG<br>TCCGGCACCACCTGGTCCTGGACCGCGC |
| I#27 | | Deletion | Frame Shift | TGAACAG |
| I#27 | | Deletion | Deletion | CGC |
| I#27 | | Deletion | Deletion | CGC |
| I#27 | | Deletion | Deletion | GAG |
| I#27 | | Deletion | Deletion | GAG |
| I#27 | | Deletion | | AG |
| I#27 | | Deletion | | G |
| I#27 | | Deletion | | AGCG |
| I#27 | | Deletion (tandem repeat) | | A |
| I#27, I#35, I#41 | | Deletion | Frame Shift | A--T |
| I#27, I#35, I#41, I#7 | A | Insertion | | - |
| I#27, I#35, I#41, I#7 | | Deletion | | A |
| I#27, I#41, I#7 | C | Insertion (tandem repeat) | | - |
| I#27, I#5 | | Deletion | Deletion | CCG |
| I#27, I#5 | | Deletion | Deletion | CCCCCA |
| I#27, I#5 | | Deletion | Deletion | ATA |
| I#27, I#5 | | Deletion | Deletion | ATA |
| I#27, I#7 | A | Insertion | | - |
| I#35 | TTTGTAA | Insertion |  | ------- |
| I#35 | T | Insertion | Frame Shift | - |
| I#35 | GAAA | Insertion | Frame Shift | ---- |
| I#35 | AAAA | Insertion | Frame Shift | ---- |
| I#35 | AAAA | Insertion | Frame Shift | ---- |
| I#35 | A | Deletion | Frame Shift | GTACAGCTCG---------T-CCAGGCCGCGCA |
| I#35 | A | SNP (transition) | None | G |
| I#35 | | Deletion | | TACA |
| I#35 | | Deletion | | T |
| I#35 | | Deletion | | G |
| I#35 | | Deletion | | AATTCTCATAAGAGT |

| | | | | |
|---|---|---|---|---|
| I#35 | | Deletion | | T |
| I#35 | | Deletion | | ATGGTCGACATGGTAAATGCATTA |
| I#35 | | Deletion | | ATTG |
| I#35 | | Deletion | | C |
| I#35 | | Deletion | | ATAGACA |
| I#35 | | Deletion | | GAACTAT |
| I#35 | | Deletion | | T |
| I#35 | | Deletion | | AAATGATT |
| I#35 | | Deletion | | G |
| I#35 | | Deletion | Frame Shift | CC |
| I#35 | | Deletion | Frame Shift | CCCCA |
| I#35 | | Deletion | Frame Shift | G |
| I#35 |  | Deletion | Frame Shift | TCCAGAACTCGACC---------GCTCCGGCGACGTCGCGCGCGGTGAG |
| I#35 | | Deletion | Deletion | CCGGAACGGCACTGG |
| I#35 | | Deletion | Start Codon Loss | AACTTGGCCAT |
| I#35 | | Deletion | Frame Shift | G |
| I#35 | | Deletion | Frame Shift | GC |
| I#35 | | Deletion | Frame Shift | GC |
| I#35, I#4, I#41 | A | Insertion (tandem repeat) | | - |
| I#35, I#41 | AT | Insertion | Frame Shift | -------- |
| I#35, I#41 | A | Insertion | | - |
| I#35, I#41, I#5, I#7 | T | Insertion | Frame Shift | - |
| I#35, I#41, I#5, I#7 | T | Insertion | Frame Shift | - |
| I#35, I#41, I#5, I#7 | | Deletion | Frame Shift | T |
| I#35, I#41, I#5, I#7 | | Deletion | Frame Shift | T |
| I#35, I#5 |  | Deletion | Deletion | GCCGATCTCGGTCATGGCCGGC------CCGGAGGCGTCCCGGAAGTTCGTGGACACGACCTCCGACCACTC |
| I#4 | TTTATCGCA | Insertion | Truncation | A-------- |
| I#4 | TCCTAA | Insertion | Truncation | ------ |
| I#4 | T | SNP | None | T- |
| I#4 | G | Insertion (tandem repeat) | Frame Shift | - |
| I#4 | G | Insertion | | - |
| I#4 | G | Insertion | | - |
| I#4 | CCACAC | Insertion |  | ------ |
| I#4 | A | Insertion | | - |
| I#4 | A | SNP | None | A- |
| I#4 | | Deletion | | AACAGGCAG |
| I#4 | | Deletion | | A |
| I#4 | | Deletion | | CAGCTA |
| I#4, I#7 | T | SNP | None | T- |
| I#4, I#7 | C | Insertion (tandem repeat) | | - |
| I#41 | T | SNP | None | --------T |
| I#41 | GG | Insertion | Frame Shift | -- |
| I#41 | CTGTTCGAC | Insertion | Insertion | --------- |
| I#41 | C | Insertion | Frame Shift | - |
| I#41 | C | Insertion | Frame Shift | - |
| I#41 | AAGAACACA | Insertion |  | --------- |
| I#41 | A | Insertion | | - |
| I#41 | | Deletion | | A |
| I#41 | | Deletion | Frame Shift | G |
| I#41 | | Deletion | Frame Shift | C |
| I#41 | | Deletion | Frame Shift | C |
| I#41 | | Deletion | Deletion | TGC |
| I#5 | T | SNP (transversion) | | G |
| I#5 | T | SNP (transversion) | | A |
| I#5 | T | SNP (transition) | | C |
| I#5 | G | SNP (transversion) | | T |
| I#5 | G | SNP (transversion) | | T |
| I#5 | G | SNP (transversion) | | C |
| I#5 | CT | Insertion | | -- |
| I#5 | C | SNP (transition) | | T |
| I#5 | C | SNP (transversion) | | A |
| I#5 | A | Deletion | Frame Shift | GGTCCGTCGTCCCGGACCACACCGGCGAAGTC |
| I#5 | | Deletion | | CCC |
| I#5 | | Deletion | Frame Shift | GGCTGCTC |
| I#5 | | Deletion | Deletion | GGC |
| I#5 |  | Deletion | Frame Shift | GTACAGCTCG---------T-CCAGGCCGCGCACC |
| I#5 | | Deletion | Frame Shift | CCCAGGCCAGGGTGTTGTCCGGCACCACCTGGTCCTGGACCGCGCTGATGAACAG |
| I#5 |  | Deletion | Extension | CTACATCTTCA--------CAAGAAC |
| I#5 | | Deletion | Frame Shift | TC |
| I#5 | | Deletion | Frame Shift | TC |
| I#5 | | Deletion | Frame Shift | AA |
| I#5 | | Deletion | Frame Shift | AA |
| I#5 | | Deletion | Deletion | AC-A |
| I#5 | | Deletion | | CTGG |
| I#7 | T | Insertion (tandem repeat) | | - |

| | | | | |
|---|---|---|---|---|
| I#7 | T | Insertion | | - |
| I#7 | T | Insertion | | - |
| I#7 | T | Insertion | | - |
| I#7 | GA | Insertion | Frame Shift | -- |
| I#7 | G | Insertion | | - |
| I#7 | C | Insertion (tandem repeat) | | - |
| I#7 | C | Insertion (tandem repeat) | | - |
| I#7 | C | Insertion | | - |
| I#7 | AGTTCGAGTATGC | Insertion | Frame Shift | ------------- |
| I#7 | AATAAT | Insertion | Insertion | ------ |
| I#7 | AAACCTCAGCCCTGGTT | Insertion | Frame Shift | ---------------- |
| I#7 | A | Insertion | | - |
| I#7 | | Deletion | Frame Shift | GGCACCACCT |
| I#7 | | Deletion | | G |

### Variant nucleotides happened from experiment II

| Variant events | Polymorphism type | Variant genotyping | Protein effect | Nucleotide genotyping |
|---|---|---|---|---|
| II#11 | G | SNP (transversion) | Substitution | T |
| II#11 | G | SNP (transversion) | Substitution | T |
| II#11 | C | SNP (transversion) | Substitution | A |
| II#11 | C | Insertion | | - |
| II#11 | AC | Insertion | | -- |
| II#11 | A | Insertion | Frame Shift | - |
| II#11, II#23 | | Deletion | | ATACACA |
| II#13 | T | Insertion | Frame Shift | - |
| II#13 | | Deletion | | A |
| II#13 | | Deletion | | TGCAAC |
| II#13 | | Deletion | | A |
| II#13 | | Deletion | | C |
| II#13 | | Deletion | | TTA |
| II#13 | | Deletion | Deletion | CCA |
| II#13 | | Deletion | Frame Shift | C |
| II#13 | | Deletion (tandem repeat) | Frame Shift | T |
| II#13, II#40, II#48 | | Deletion | Deletion | CTCCGA |
| II#13, II#41, II#48 | | Deletion | | TGAT |
| II#13, II#48 | | Deletion | | TTTTTTTTTTTTTTTTTTTTTTTTTTTTTTTATGATGA |
| II#13, II#48 | | Deletion | | GATGATG |
| II#13, II#48 | | Deletion | Extension | TCAGTCCT-<br>GCTCCTCGGCCACGAAGTGCACGCAGTTGCCGGCCGGGTCGC<br>GCAGGGCGAACTCCCGCCCCCACGGCTGCTCGCCGATCT-<br>[illegible] |
| II#13, II#48 | | Deletion | Deletion | CTCGGCGTACAGCTC |
| II#13, II#48 | | Deletion | Deletion | GTC |
| II#14 | TTA | Insertion | | --- |
| II#14 | T | Insertion (tandem repeat) | Frame Shift | - |
| II#14 | A | Insertion | Frame Shift | - |
| II#14 | | Deletion (tandem repeat) | | A |
| II#19 | T | Insertion | | - |
| II#19 | C | SNP (transition) | Substitution | T |
| II#19 | AAGAC | Insertion |  | ----- |
| II#19 | A | Insertion (tandem repeat) | | - |
| II#19 | A | SNP (transversion) | Substitution | C |
| II#19 | | Deletion | | C |
| II#19 | | Deletion | Deletion | CAGGCC |
| II#21 | T | SNP (transversion) | | G |
| II#21 | T | Insertion | | - |
| II#21 | G | Insertion | | - |
| II#21 | CGGGGA | Insertion | Insertion | ------ |
| II#21 | | Deletion | Frame Shift | T |
| II#22 | G | Insertion | Frame Shift | - |
| II#22 | G | SNP (transition) | | A |
| II#22 | C | SNP (transversion) | | A |
| II#22 | C | SNP (transversion) | | A |
| II#22 | ACC | Insertion | Insertion | --- |
| II#22 | ACC | Insertion | Insertion | --- |
| II#22 | A | SNP (transversion) | | C |
| II#22 | | Deletion | Frame Shift | ATCT |
| II#23 | T | SNP (transition) | Substitution | C |
| II#23 | C | Insertion | Frame Shift | - |
| II#23 | AAT | Insertion | Insertion | --- |
| II#23 | AAT | Insertion | Insertion | --- |
| II#23 | A | Insertion | Frame Shift | - |
| II#23 | | Deletion | Frame Shift | GTCGTCCCGGACCACACCGGCGAAGT |
| II#23 | | Deletion | Frame Shift | C |
| II#23 | | Deletion | Frame Shift | GT |
| II#23 | | Deletion | Frame Shift | GT |
| II#23 | | Deletion | Deletion | CGC |
| II#23 | | Deletion | Deletion | CGC |
| II#23 | | Deletion | Deletion | GAG |
| II#23 | | Deletion | Deletion | GAG |
| II#23 | | Deletion | Deletion | ATC |
| II#23 | | Deletion | Deletion | ATC |
| II#23 | | Deletion | Deletion | CAC |
| II#23 | | Deletion | Deletion | CAC |
| II#23 | | Deletion | Deletion | ATG |
| II#23 | | Deletion | Deletion | ATG |
| II#23 | | Deletion | Deletion | CAG |
| II#23 | | Deletion | Deletion | CAG |

| | | | | |
|---|---|---|---|---|
| II#23 | | Deletion | Deletion | GTC |
| II#23 | | Deletion | Deletion | GTC |
| II#23 | | Deletion | Deletion | CTC |
| II#23 | | Deletion | Deletion | CTC |
| II#23 | | Deletion | | AAAA |
| II#23 | | Deletion | | GAAGA |
| II#23 | | Deletion | | CATGAGCG |
| II#23 | | Deletion | | CC |
| II#23 | | Deletion | | ATAGC |
| II#23 | | Deletion | | TGT |
| II#23, II#38, II#48 | | Deletion | Deletion | GGTCAC |
| II#23, II#38, II#48, II#53 | | Deletion | Deletion | CACCTGGTCCTGGACCGCGCTGATGAACAG |
| II#23, II#40 | | Deletion | Deletion | ATA |
| II#23, II#40 | | Deletion | Deletion | ATA |
| II#23, II#40 | | Deletion | Deletion | CAA |
| II#23, II#40 | | Deletion | Deletion | CAA |
| II#23, II#40 | | Deletion | Deletion | GTT |
| II#23, II#40 | | Deletion | Deletion | GTT |
| II#23, II#40 | | Deletion | Deletion | CAT |
| II#23, II#40 | | Deletion | Deletion | CAT |
| II#23, II#40 | | Deletion | Deletion | GAA |
| II#23, II#40 | | Deletion | Deletion | GAA |
| II#23, II#40 | | Deletion | Deletion | TGG |
| II#23, II#40 | | Deletion | Deletion | TGG |
| II#23, II#40 | | Deletion | Deletion | GTG |
| II#23, II#40 | | Deletion | Deletion | GTG |
| II#23, II#40 | | Deletion | Deletion | TGC |
| II#23, II#40 | | Deletion | Deletion | TGC |
| II#23, II#40 | | Deletion | | CTGGTATTGAAA |
| II#23, II#40 | | Deletion | | TTAGAAAGAGA |
| II#23, II#48 | | Deletion | | ACTA |
| II#23, II#53 | | Deletion | | GTCAGA |
| II#23, II#53 | | Deletion | | TC |
| II#29 | T | Insertion | | - |
| II#29 | T | Insertion | | - |
| II#29 | T | Insertion | | - |
| II#29 | C | Insertion | | - |
| II#29 | C | Insertion | | - |
| II#29 | C | Insertion | | - |
| II#29 | A | Insertion | | - |
| II#29 | A | Insertion | | - |
| II#29 | | Deletion | | A |
| II#29 | | Deletion | | GTCT |
| II#29, II#59 | AT | Insertion | | -- |
| II#3 | TGAA | Insertion |  | ---- |
| II#3 | T | Insertion | | - |
| II#3 | A | SNP (transversion) | | T |
| II#3 | | Deletion | | A |
| II#3 | | Deletion | | CAC |
| II#3 | | Deletion | Start Codon Loss | GAACGGCACTGGTCAACTTGGCCAT |
| II#3 | | Deletion | Deletion | A-CAAGTTTTAAGGGTTGC |
| II#3 | | Deletion | Deletion | GCC |
| II#3 | | Deletion | Deletion | GCC |
| II#3 | | Deletion | Deletion | G---GC |
| II#3 | | Deletion | Deletion | G---GC |
| II#3 | | Deletion | Deletion | ATT |
| II#3 | | Deletion | Deletion | ATT |
| II#3 | | Deletion | Deletion | TGT |
| II#3 | | Deletion | Deletion | TGT |
| II#3 | | Deletion | Deletion | AAA |
| II#3 | | Deletion | Deletion | AAA |
| II#3 | | Deletion | Deletion | GTA |
| II#3 | | Deletion | Deletion | GTA |
| II#3 | | Deletion | Frame Shift | T---G |
| II#3 | | Deletion | Frame Shift | T---G |
| II#3, II#48 | | Deletion | Extension | CTACATC-TTCACAAGAAC |
| II#3, II#48 | | Deletion | Deletion | AAG |
| II#3, II#48 | | Deletion | Deletion | AGG |
| II#3, II#48 | | Deletion | Deletion | AGG |
| II#3, II#48 | | Deletion | Deletion | AGA |

| | | | | |
|---|---|---|---|---|
| II#3, II#48 | | Deletion | Deletion | AGA |
| II#35 | TCC | Insertion | Insertion | --- |
| II#35 | T | SNP (transition) | None | C |
| II#35 | G | Insertion | Frame Shift | - |
| II#35 | A | Insertion (tandem repeat) | | - |
| II#35 | A | Deletion | | AC |
| II#35 | | Deletion | | G |
| II#35 | | Deletion | | C |
| II#35 | | Deletion | | A |
| II#35 | | Deletion | | TTT |
| II#35 | | Deletion | | G |
| II#35, II#41 | | Deletion (tandem repeat) | | A |
| II#38 | TAA | Insertion | | --- |
| II#38 | C | Insertion | | - |
| II#38 | A | SNP (transition) | Substitution | G |
| II#38 | A | Insertion | Frame Shift | - |
| II#38 | A | Insertion | Frame Shift | - |
| II#38 | | Deletion | | CAGCA |
| II#38 | | Deletion | Frame Shift | AGGGTGTTGTCCGGCAC |
| II#38 | | Deletion | | A |
| II#40 | T | Insertion | Truncation | - |
| II#40 | G | SNP (transversion) | | C |
| II#40 | | Deletion | | A |
| II#40 | | Deletion | Frame Shift | CCCACGGCTGCTCGCCGATCT-CGGTCATGGCCGGCCCGGAGGCGTCCCGGAAGTTCGTGGACACCAC |
| II#40 | | Deletion | Frame Shift | TC |
| II#40 | | Deletion | Frame Shift | TC |
| II#41 | T | SNP (transversion) | None | G |
| II#41 | T | SNP (transition) | Substitution | C |
| II#41 | G | SNP (transversion) | Substitution | T |
| II#41 | CGG | Insertion | | --- |
| II#41 | C | Insertion | | - |
| II#41 | A | SNP (transversion) | Substitution | C |
| II#41 | | Deletion | Frame Shift | A |
| II#48 | T | Insertion | | - |
| II#48 | T | Insertion | Frame Shift | - |
| II#48 | CCGC | Insertion |  | ---- |
| II#48 | C | SNP (transversion) | | A |
| II#48 | C | SNP (transversion) | | G |
| II#48 | AT | Insertion | | -- |
| II#48 | A | SNP (transversion) | | T |
| II#48 | | Deletion | Deletion | C-AGGCC |
| II#48 | | Deletion | Deletion | GCGCACCCACACCCAGGCCAGGGTGTTGTCCGGCAC |
| II#48 | | Deletion | Deletion | GTCGTCCCGGACCACACCGGCGAAGTCGTC |
| II#48 | | Deletion | Deletion | CTCCAC |
| II#48 | | Deletion | Deletion | GAAGTCCCG |
| II#48 | | Deletion | Deletion | GGAGAACCCGAGCCGGTCGGTCCA |
| II#48 | | Deletion | Start Codon Loss | GAACTCGACCGCTCCGGCGACGTCGCGCGCGGTGAGCACCGGAACGGCACTGGTCAACTTGGCCAT |
| II#48 | | Deletion | Frame Shift | A |
| II#48 | | Deletion | | A |
| II#48 | | Deletion | Frame Shift | TT |
| II#48 | | Deletion | Frame Shift | CTGCAC-A |
| II#48 | | Deletion | | CT |
| II#48, II#59 | | Deletion | | GG |
| II#53 | T | Insertion | | - |
| II#53 | G | SNP (transversion) | Substitution | C |
| II#53 | CCCACG | Insertion | Insertion | ------ |
| II#53 | C | Insertion | | - |
| II#53 | | Deletion | Frame Shift | TTGTCCGGCAC |
| II#53 | | Deletion | | A |
| II#59 | T | Insertion (tandem repeat) | | - |
| II#59 | G | Insertion | | - |
| II#59 | G | Insertion | | - |
| II#59 | C | SNP (transition) | | T |
| II#59 | C | Insertion | | - |
| II#59 | C | Insertion | | - |
| II#59 | C | Insertion | | - |
| II#59 | C | Insertion | | - |
| II#59 | C | SNP (transversion) | Substitution | G |
| II#59 | AA | Substitution | Substitution | TC |

| | | | | |
|---|---|---|---|---|
| II#59 | A | Insertion | | - |
| II#59 | | Deletion | | A |
| II#59 | | Deletion | | AA |
| II#7 | C | Insertion | | - |
| II#7 | C | Insertion | | - |
| II#7 | C | Deletion | | AGGCAAA |
| II#7 | C | SNP (transversion) | | G |
| II#7 | C | SNP (transversion) | Substitution | A |
| II#7 | AC | Substitution | Substitution | CA |
| II#7 | | Deletion | | G |
| II#7 | | Deletion | Deletion | CCG |

### Variant nucleotides happened from experiment III

| Variant events | Polymorphism type | Variant genotyping | Protein effect | Nucleotide genotyping |
|---|---|---|---|---|
| III#6 | ----- | Insertion |  | AGCTC |
| III#6 | TCC | Deletion | Deletion | |
| III#6 | A | SNP (transversion) | Substitution | C |
| III#6 | AAA | Deletion | | |
| III#6 | A | Deletion (tandem repeat) | | |
| III#8 | A | SNP (transversion) | | T |
| III#8 | GA | Substitution | | AG |
| III#8 | AG | Deletion | | |
| III#8 | TTA | Deletion | | |
| III#8 | T | Deletion (tandem repeat) | | |
| III#8 | - | Insertion | Frame Shift | C |
| III#8 | C | Deletion | Frame Shift | |
| III#8 | TCATACAAGTT | Deletion | Frame Shift | |
| III#8 | CATGAATGGGTGTGC | Deletion | Deletion | |
| III#8 | CTGG | Deletion | | |
| III#8 | CGT | Deletion | | C |
| III#8 | -- | Insertion | Frame Shift | TA |
| III#10 | TT | Deletion | | |
| III#10 | GCCG | Deletion | Frame Shift | |
| III#10 | TTGTCCGGC | Deletion | Deletion | |
| III#10 | - | Insertion | | C |
| III#10 | -- | Insertion | | TT |
| III#14 | ACTATT | Deletion | | |
| III#14 | A | Deletion | | |
| III#14 | - | Insertion | Frame Shift | C |
| III#14 | ------- | Insertion | Frame Shift | TTAGGTA |
| III#14, III#17, III#43 | GGCGACGTCGCGCGCGGTGAG | Deletion | Deletion | |
| III#14, III#17, III#43 | CACCGG | Deletion | Deletion | |
| III#14, III#17, III#43 | AACGGCACTGGTCAACTTGGCCAT | Deletion | Start Codon Loss | |
| III#14, III#43 | T | SNP (transition) | | C |
| III#14, III#43 | GAACCCGAGCCGGTCGGTCC----- AGAACTCGAC | Deletion | Deletion |  |
| III#14, III#43 | CGCTCC | Deletion | Deletion | |
| III#14, III#43 | CTACATCTTCACAAGAA | Deletion | Extension | |
| III#14, III#43, III#8 | TTTTTTTTTTTTTTTTTTTTTTTTTTTTTTA TGATGATGATGATGATG | Deletion | | |
| III#14, III#43, III#8 | TCAGTCCTGCTCCTCGGCCACGAAGTG | Deletion | Extension | |
| III#14, III#43, III#8 | CAC | Deletion | Deletion | |
| III#14, III#43, III#8 | GCA | Deletion | Deletion | |
| III#14, III#43, III#8 | GTT | Deletion | Deletion | |
| III#14, III#43, III#8 | GCCGGCCGGGTC | Deletion | Deletion | |
| III#14, III#43, III#8 | GCG | Deletion | Deletion | |
| III#17 | - | Insertion | | C |
| III#17 | ATT | Deletion | | |
| III#17 | CTACATCTTCACAAGAAC | Deletion | Extension | |
| III#17 | ACAAGTTTTAAGGGTT | Deletion | Frame Shift | |
| III#17 | A | Deletion | Frame Shift | |
| III#17 | --------- | Insertion |  | CGCAGCCAG |
| III#21 | - | Insertion | | C |
| III#21 | ----- | Insertion | Frame Shift | ACAAT |
| III#21 | G | Deletion | | |
| III#21 | GC | Substitution | Substitution | CG |
| III#21, III#43 | ACTCCAT | Deletion | | |
| III#23 | CTTTTTGT | Deletion | | |
| III#23 | -- | Insertion | | AA |
| III#23 | A | Deletion | Frame Shift | |
| III#23 | - | Insertion | | T |
| III#24 | - | Insertion | Frame Shift | T |

| | | | | |
|---|---|---|---|---|
| III#24 | -------- | Insertion | Truncation | ATGGTACT |
| III#24 | A | Deletion | Frame Shift | |
| III#24 | T | SNP (transversion) | Substitution | G |
| III#24 | TT | Substitution | Substitution | AC |
| III#24, III#8 | AC | Deletion | | |
| III#27 | ------- | Insertion |  | CCAATTC |
| III#27 | CCGA | Deletion | Frame Shift | |
| III#27 | C | Deletion | | |
| III#27 | - | Insertion | Frame Shift | A |
| III#32 | A | Deletion | | |
| III#32 | G | SNP (transition) | Substitution | A |
| III#32 | CAC | Substitution | Substitution | GTA |
| III#32 | GCA | Substitution | Substitution | AAC |
| III#32 | GCACCA | Deletion | Deletion | |
| III#32 | ---- | Insertion | Frame Shift | CTCG |
| III#32 | - | Insertion | Frame Shift | T |
| III#36 | - | Insertion | | A |
| III#36 | ------- | Insertion |  | TGGGTTC |
| III#36 | C | Deletion | Frame Shift | |
| III#36 | GAAC | Deletion | | |
| III#43 | --------- | Insertion |  | TCCGTAAGG |
| III#43 | A | Deletion | | |
| III#43 | - | Insertion | | G |
| III#43 | T | SNP (transversion) | | A |
| III#43 | CTTCACAAAATCATCTGGC | Deletion | | |
| III#43, III#61 | A | Deletion | | |
| III#45 | T | Deletion | | |
| III#45 | CTC | Substitution | | TCA |
| III#45 | - | Insertion | Frame Shift | T |
| III#45 | --------- | Insertion | Insertion | TTGGAGCTT |
| III#45 | G | Deletion | | |
| III#53 | TTAAAT | Deletion | | |
| III#53 | --------- | Insertion |  | TGGGGTTTA |
| III#53 | G | Deletion | Frame Shift | |
| III#53 | - | Insertion | | C |
| III#53 | CTGC-ACA | Deletion | Frame Shift | |
| III#61 | --------- | Insertion |  | GGCGCTTTA |
| III#61 | - | Insertion | Frame Shift | T |
| III#61 | GTTC | Deletion | Frame Shift | |

### Variant nucleotides happened from experiment IV

| Variant events | Polymorphism type | Variant genotyping | Protein effect | Nucleotide genotyping |
|---|---|---|---|---|
| IV#13 | TT | Insertion | | -- |
| IV#13 | T | Insertion (tandem repeat) | | - |
| IV#13 | T | SNP (transition) | Substitution | C |
| IV#13 | C | SNP (transition) | | T |
| IV#13 | C | SNP (transversion) | Substitution | A |
| IV#13 | C | SNP (transversion) | Substitution | G |
| IV#13 | C | SNP (transition) | | T |
| IV#13 | A | Insertion | | - |
| IV#13 | | Deletion | Frame Shift | G |
| IV#13 | | Deletion | Frame Shift | GC |
| IV#13, IV#80 | | Deletion | Frame Shift | TG |
| IV#17 | T | SNP (transversion) | | A |
| IV#17 | GC | Insertion | | -- |
| IV#17 | C | Insertion | | - |
| IV#17 | C | SNP (transversion) | | G |
| IV#17 | | Deletion | | A |
| IV#17 | | Deletion | | AGGC |
| IV#17, IV#9 | A | Insertion (tandem repeat) | | - |
| IV#19 | T | SNP (transversion) | | G |
| IV#19 | G | Insertion | | - |
| IV#19 | AA | Insertion | Frame Shift | -- |
| IV#19 | A | Insertion (tandem repeat) | | - |
| IV#19 | | Deletion (tandem repeat) | Frame Shift | C |
| IV#19 | | Deletion | | GAG |
| IV#27 | G | Insertion | | - |
| IV#27 | G | Insertion (tandem repeat) | | - |
| IV#27 | CC | Insertion | | -- |
| IV#27 | C | Insertion | Frame Shift | - |
| IV#27 | AG | Insertion | | -- |
| IV#27 | A | SNP (transition) | None | G |
| IV#27 | | Deletion | Frame Shift | T |
| IV#32 | G | Insertion | | - |
| IV#32 | G | Insertion | Frame Shift | - |
| IV#32 | C | SNP (transition) | | T |
| IV#32 | C | Insertion | | - |
| IV#32 | A | Insertion | | - |
| IV#32 | | Deletion | Frame Shift | TCCCGGGAGAACCCGAGCC--<br>GGTCGGTCCAGAACTCGACCGCTCCGGC<br>GACGTCGCGCGCGGTGAG |
| IV#32 | | Deletion | Frame Shift | C |
| IV#32 | | Deletion | Frame Shift | C |
| IV#32 | | Deletion | | T |
| IV#32 | | Deletion | | ACA |
| IV#32 | | Deletion | | CAT |
| IV#32 | | Deletion (tandem repeat) | | T |
| IV#32 | | Deletion | | TTT |
| IV#32 | | Deletion | | AACAT |
| IV#32 | | Deletion | | ATT |
| IV#32, IV#39 | | Deletion | | G |
| IV#32, IV#80 | | Deletion | | G |
| IV#39 | T | Insertion | | - |
| IV#39 | G | SNP (transversion) | | T |
| IV#39 | G | Insertion | | - |
| IV#39 | G | Deletion | Frame Shift | CG |
| IV#39 | C | Insertion | | - |
| IV#39 | A | Insertion | | - |
| IV#45 | T | Insertion | Frame Shift | - |
| IV#45 | G | SNP (transversion) | | T |
| IV#45 | AA | Insertion | | -- |
| IV#45 | A | Insertion | | - |
| IV#45 | | Deletion | | A |
| IV#46 | T | SNP (transversion) | Substitution | A |
| IV#46 | G | Insertion | | - |
| IV#46 | C | Insertion | Frame Shift | - |
| IV#46 | C | Insertion | Frame Shift | - |
| IV#46 | A | Insertion | | - |
| IV#46 | | Deletion | Frame Shift | G |
| IV#46 | | Deletion | | CC |
| IV#46, IV#71 | T | Insertion (tandem repeat) | | - |
| IV#54 | T | Insertion | | - |

| | | | | |
|---|---|---|---|---|
| IV#54 | G | Insertion (tandem repeat) | | - |
| IV#54 | G | SNP (transversion) | | C |
| IV#54 | G | Insertion | | - |
| IV#54 | C | SNP (transversion) | | G |
| IV#54 | C | Insertion | | - |
| IV#54 | C | SNP (transition) | | T |
| IV#54 | | Deletion | | G |
| IV#54 | | Deletion (tandem repeat) | | T |
| IV#54, IV#80 | A | SNP (transition) | | G |
| IV#60 | T | SNP (transversion) | | A |
| IV#60 | T | SNP (transversion) | | A |
| IV#60 | GA | Insertion | Frame Shift | -- |
| IV#60 | C | Insertion | Frame Shift | - |
| IV#60 | A | Insertion | | - |
| IV#60 | | Deletion | | T |
| IV#60, IV#83 | | Deletion | | GA |
| IV#60, IV#87 | | Deletion (tandem repeat) | | T |
| IV#68 | CA | Insertion | | -- |
| IV#68 | CA | Insertion | | -- |
| IV#68 | C | Insertion | | - |
| IV#71 | TG | Insertion | Frame Shift | -- |
| IV#71 | T | SNP (transition) | None | C |
| IV#71 | GC | Substitution | Substitution | AA |
| IV#71 | GA | Deletion | Deletion | ACAAG |
| IV#71 | C | SNP (transition) | | T |
| IV#71 | C | SNP (transversion) | | A |
| IV#71 | A | SNP (transversion) | Substitution | C |
| IV#71 | A | Insertion | Frame Shift | - |
| IV#71 | | Deletion (tandem repeat) | | C |
| IV#71 | | Deletion | | TTATTTTTTTTTTTTTTTTTTTTTTTTTTTT TTATGATGATGATGATGATG |
| IV#71 | | Deletion | Extension | TCAGTCCT-GC--TCCTCGGCCACGAAGT-- GCACGCAGTTGCC |
| IV#71 | | Deletion | Frame Shift | GGCCGGGTCG- CGCAGGGCGAACTCCCGCCCCCACGGCT GCTCGCCGA |
| IV#71 | | Deletion | Frame Shift | CGGTCATGGC |
| IV#71 | | Deletion | Deletion | CGGCCC |
| IV#71 | | Deletion | Deletion | GGAGGCGTC |
| IV#71 | | Deletion | Deletion | CCGGAAGTTCGTGGA |
| IV#71 | | Deletion | Deletion | CACGACCTCCGACCACTCGGCGTA |
| IV#71 | | Deletion | Deletion | CAGCTCGTC |
| IV#71 | | Deletion | Deletion | CAGGCCGC-G |
| IV#71 | | Deletion | Deletion | CACCCACACCCAGGCC-AG |
| IV#71 | | Deletion | Deletion | GGTGTTGTC |
| IV#71 | | Deletion | Deletion | CG-GCACCA-CCTGGTCCTGGACCGC |
| IV#71 | | Deletion | Deletion | GCTGAT |
| IV#71 | | Deletion | Deletion | GAACAGGGTCACGTCGTCCCGGACCAC ACCGGCGAAGTCGTCCTCCACGAAGTCC CGGGAGAACCCGAGCC-- GGTCGGTCCAGAACTCGACCGCTCCGGC GACGTCGCGCGCGGTGAG |
| IV#71 | | Deletion | Deletion | CACCGGAACGGC |
| IV#71 | | Deletion | Frame Shift | ACTGGTC |
| IV#71 | | Deletion | Start Codon Loss | GGCCAT |
| IV#71 | | Deletion | Extension | CTACATCTTCAC |
| IV#71 | | Deletion | Deletion | A---AG |
| IV#71 | | Deletion | | T |
| IV#71 | | Deletion | | ACC |
| IV#71, IV#87 | | Deletion | Deletion | AAC |
| IV#75 | T | Insertion | Frame Shift | - |
| IV#75 | T | Insertion | | - |
| IV#75 | GA | Insertion | | -- |
| IV#75 | G | SNP (transversion) | Substitution | C |
| IV#75 | A | SNP (transversion) | None | C |
| IV#75 | | Deletion | | TCAAGA |
| IV#75 | | Deletion | | T |
| IV#75 | | Deletion | Frame Shift | G |
| IV#76 | T | SNP (transversion) | | A |
| IV#76 | G | Insertion | Frame Shift | - |
| IV#76 | G | SNP (transversion) | | C |

| | | | | |
|---|---|---|---|---|
| IV#76 | CA | Insertion | Frame Shift | A- |
| IV#76 | C | SNP (transversion) | | A |
| IV#76 | C | SNP (transversion) | Substitution | A |
| IV#76 | A | SNP (transversion) | | T |
| IV#76 | A | SNP (transversion) | | T |
| IV#76 | A | SNP (transversion) | | T |
| IV#76 | A | SNP (transversion) | Substitution | C |
| IV#76 | | Deletion | | TA |
| IV#76 | | Deletion | Frame Shift | A |
| IV#76 | | Deletion | Deletion | AAG |
| IV#76 | | Deletion | Deletion | AGG |
| IV#76, IV#87 | | Deletion | | G |
| IV#79 | GA | Insertion | Frame Shift | -- |
| IV#79 | G | Insertion | | - |
| IV#79 | G | Insertion | | - |
| IV#79 | A | SNP (transversion) | | T |
| IV#79 | | Deletion (tandem repeat) | | C |
| IV#80 | G | Insertion (tandem repeat) | Frame Shift | - |
| IV#80 | G | SNP (transversion) | | T |
| IV#80 | G | Insertion | | - |
| IV#80 | CC | Insertion | | -- |
| IV#80 | C | SNP (transition) | | T |
| IV#80 | C | SNP (transition) | | T |
| IV#80 | C | SNP (transition) | | T |
| IV#80 | C | Insertion | | - |
| IV#80 | C | Insertion | Frame Shift | - |
| IV#80 | C | Insertion | | - |
| IV#80 | C | Insertion | | - |
| IV#80 | | Deletion | Frame Shift | G |
| IV#80 | | Deletion | Deletion | TGC |
| IV#80 | | Deletion | | C |
| IV#83 | G | Insertion | | - |
| IV#83 | C | Insertion | Frame Shift | - |
| IV#83 | AG | Insertion | | -- |
| IV#83 | A | SNP (transversion) | Substitution | C |
| IV#83 | | Deletion | | C |
| IV#85 | TT | Insertion | | -- |
| IV#85 | TC | Substitution | Substitution | CT |
| IV#85 | T | Insertion | | - |
| IV#85 | G | Insertion (tandem repeat) | | - |
| IV#85 | G | SNP (transition) | Substitution | A |
| IV#85 | G | Deletion | | GGAA |
| IV#85 | | Deletion | | GAG |
| IV#85 | | Deletion | | T |
| IV#85 | | Deletion | | T |
| IV#85 | | Deletion | | A |
| IV#85 | | Deletion | | TGGT |
| IV#85 | | Deletion (tandem repeat) | | A |
| IV#85 | | Deletion (tandem repeat) | Frame Shift | T |
| IV#87 | G | Insertion | Frame Shift | - |
| IV#87 | G | SNP (transition) | None | A |
| IV#87 | CCGAC | Substitution | Substitution | ATCCA |
| IV#87 | AGC | Insertion | Frame Shift | #NAME? |
| IV#87 | AA | Insertion | Frame Shift | -- |
| IV#87 | A | SNP (transition) | Substitution | G |
| IV#87 | | Deletion | | TTGCTAAT |
| IV#87 | | Deletion | | G |
| IV#87 | | Deletion | Frame Shift | AG |
| IV#87 | | Deletion | Frame Shift | AC |
| IV#87 | | Deletion | Frame Shift | G |
| IV#87, IV#92 | | Deletion | | A |
| IV#9 | G | Insertion | | - |
| IV#9 | C | Insertion | | - |
| IV#9 | C | Insertion (tandem repeat) | Frame Shift | - |
| IV#9 | | Deletion (tandem repeat) | | A |
| IV#9 | | Deletion | | GC |
| IV#90 | T | Insertion | | - |
| IV#90 | G | Insertion | | - |
| IV#90 | G | Insertion | | - |
| IV#90 | C | SNP (transversion) | | G |
| IV#90 | C | SNP (transversion) | | G |

| | | | | |
|---|---|---|---|---|
| IV#90 | A | SNP (transversion) | | T |
| IV#90 | A | Insertion | | - |
| IV#90 | | Deletion | | A |
| IV#90 | | Deletion | Frame Shift | T |
| IV#92 | T | SNP (transversion) | | G |
| IV#92 | T | Insertion | | - |
| IV#92 | G | Insertion (tandem repeat) | Frame Shift | - |
| IV#92 | CCG | Insertion | Insertion | --- |
| IV#92 | C | SNP (transversion) | | A |
| IV#92 | C | SNP (transition) | | T |
| IV#92 | AA | Insertion | Frame Shift | -- |
| IV#92 | A | Insertion | | - |
| IV#92 | A | Insertion (tandem repeat) | | - |
| IV#92 | A | Insertion (tandem repeat) | | - |
| IV#92 | | Deletion | Frame Shift | A |
| IV#92 | | Deletion | Deletion | TAC |
| IV#92 | | Deletion | | CAGTAAC |

#### Variant nucleotides happened from experiment V

| Variant events | Polymorphism type | Variant genotyping | Protein effect | Nucleotide genotyping |
|---|---|---|---|---|
| V#101 | TC | Insertion | | -- |
| V#101 | A | Insertion | Truncation | - |
| V#101 | | Deletion | | G |
| V#18 | TCG | Insertion | Insertion | --- |
| V#18 | G | Insertion | | - |
| V#18 | C | SNP (transition) | | T |
| V#18 | | Deletion (tandem repeat) | | C |
| V#21 | GT | Substitution | | AG |
| V#21 | AG | Insertion | | -- |
| V#25 | TAA | Insertion | | --- |
| V#25 | C | Insertion | | - |
| V#25 | | Deletion (tandem repeat) | | T |
| V#29 | G | Insertion | | - |
| V#29 | G | Insertion | | - |
| V#29 | C | Insertion | | - |
| V#29 | | Deletion | | C |
| V#29, V#33 | A | Insertion | | - |
| V#3 | TCC | Insertion | Insertion | --- |
| V#3 | C | Insertion | Frame Shift | - |
| V#3 | | Deletion (tandem repeat) | | A |
| V#32 | T | Insertion (tandem repeat) | | - |
| V#32 | GAT | Insertion | | --- |
| V#32, V#95 | | Deletion (tandem repeat) | | A |
| V#33 | T | SNP (transversion) | | A |
| V#33 | GA | Substitution | | AG |
| V#33 | C | Insertion | | - |
| V#33 | A | Insertion | Frame Shift | - |
| V#37 | TA | Insertion | Frame Shift | -- |
| V#37 | G | Insertion | | - |
| V#37 | | Deletion (tandem repeat) | | T |
| V#39 | G | Insertion | | - |
| V#39 | AAC | Insertion | | --- |
| V#39 | | Deletion | | A |
| V#48 | GCA | Insertion | Insertion | --- |
| V#48 | G | Insertion | | - |
| V#48 | | Deletion | Frame Shift | G |
| V#53 | GGT | Insertion | | --- |
| V#53 | G | Insertion | | - |
| V#53 | | Deletion | Frame Shift | T |
| V#59 | T | Insertion | | - |
| V#59 | CGT | Insertion | | --- |
| V#59 | | Deletion | | G |
| V#64 | TG | Insertion | Frame Shift | -- |
| V#64 | G | Insertion | Frame Shift | - |
| V#64 | | Deletion | Frame Shift | C |
| V#67 | TT | Insertion | Frame Shift | -- |
| V#67 | A | Insertion (tandem repeat) | | - |
| V#67 | | Deletion | | A |
| V#73 | G | Insertion | | - |
| V#73 | | Deletion | | A |
| V#75 | GT | Insertion | | -- |
| V#75 | A | Insertion | Frame Shift | - |

| | | | | |
|---|---|---|---|---|
| V#75 | | Deletion (tandem repeat) | Frame Shift | T |
| V#79 | CA | Insertion | | -- |
| V#79 | A | Insertion | Frame Shift | - |
| V#79 | | Deletion | Frame Shift | G |
| V#80 | T | Insertion (tandem repeat) | | - |
| V#80 | T | SNP (transition) | Substitution | C |
| V#80 | GGG | Substitution | Substitution | ACA |
| V#80 | G | SNP (transversion) | Substitution | C |
| V#80 | CTTCAA | Insertion |  | ------ |
| V#80 | CC | Insertion | Frame Shift | -- |
| V#80 | C | SNP (transition) | Start Codon Loss | T |
| V#80 | | Deletion | | A |
| V#80 | | Deletion | | G |
| V#80 | | Deletion | Extension | TCA |
| V#80 | | Deletion | Deletion | GGCCGGGTCGCG |
| V#80 | | Deletion | Frame Shift | CA |
| V#80 | | Deletion | Extension | CTAC |
| V#80 | | Deletion (tandem repeat) | Frame Shift | T |
| V#80, V#97 | | Deletion | | AACT |
| V#82 | T | SNP (transition) | None | C |
| V#82 | CAT | Insertion | | --- |
| V#82 | | Deletion | | A |
| V#82 | | Deletion (tandem repeat) | | AA |
| V#82 | | Deletion | Frame Shift | G |
| V#82 | | Deletion | Frame Shift | G |
| V#85 | T | Insertion | Frame Shift | - |
| V#85 | CC | Insertion | | -- |
| V#85 | | Deletion | Frame Shift | A |
| V#86 | T | SNP (transversion) | Substitution | A |
| V#86 | C | Insertion | | - |
| V#86 | C | SNP (transversion) | | A |
| V#86 | ACC | Substitution | | TGG |
| V#86 | A | Insertion | Frame Shift | - |
| V#86 | | Deletion (tandem repeat) | | A |
| V#86 | | Deletion | Deletion | GCC |
| V#86 | | Deletion | Frame Shift | G |
| V#86 | | Deletion | | A |
| V#88 | AG | Insertion | | -- |
| V#88 | A | Insertion | | - |
| V#88 | | Deletion | | A |
| V#89 | T | Insertion | Frame Shift | - |
| V#89 | G | Insertion (tandem repeat) | | - |
| V#89 | G | Insertion | | - |
| V#89 | | Deletion | Frame Shift | G |
| V#89 | | Deletion | Frame Shift | A |
| V#89, V#9, V#91 | A | Insertion (tandem repeat) | | - |
| V#9 | T | SNP (transversion) | | A |
| V#9 | ATA | Insertion | Insertion | --- |
| V#9 | | Deletion | | A |
| V#91 | TC | Insertion | | -- |
| V#91 | T | Insertion | | - |
| V#91 | T | SNP (transversion) | | A |

| | | | | |
|---|---|---|---|---|
| V#91 | | Deletion | | A |
| V#92 | GC | Insertion | | -- |
| V#92 | G | Insertion | | - |
| V#92 | | Deletion | Frame Shift | G |
| V#94 | CA | Insertion | | -- |
| V#94 | A | Insertion | | - |
| V#94 | | Deletion | | T |
| V#95 | TTT | Substitution | | AAA |
| V#95 | C | Insertion | Frame Shift | - |
| V#95 | AA | Insertion | Frame Shift | -- |
| V#95 | | Deletion | Frame Shift | T |
| V#95 | | Deletion | Frame Shift | C |
| V#97 | T | Insertion (tandem repeat) | | - |
| V#97 | CG | Insertion | Frame Shift | -- |
| V#97 | CA | Substitution | Substitution | AT |
| V#97 | C | SNP (transversion) | | A |
| V#97 | C | SNP (transversion) | Substitution | G |
| V#97 | C | SNP (transversion) | | A |
| V#97 | C | Insertion | Frame Shift | - |
| V#97 | | Deletion | Frame Shift | A |
| V#97 | | Deletion | | T |
| V#98 | T | Insertion | Frame Shift | - |
| V#98 | GGA | Insertion | Insertion | --- |
