## Supplementary material for "Precise exogenous insertion and sequence replacements in poplar by simultaneous HDR overexpression and NHEJ suppression using CRISPR-Cas9": Suppl T.5 (from another version of this preprint: version 6, 2021-02-03)

| | | MKK2 expression (%) | |
|---|---|---|---|
| | | Before stress | After stress |
| | WT | 94.092 | 169.343 |
| II | II#29 | 99.746 | 169.281 |
| IV | IV#17 | 100.934 | 171.940 |
| | IV#54 | 98.417 | 172.380 |
| | IV#68 | 101.447 | 170.223 |
| V | V#21 | 99.839 | 170.455 |
| | V#25 | 97.520 | 171.290 |
| | V#29 | 99.312 | 171.915 |
| | V#32 | 99.154 | 170.925 |
| | V#39 | 98.009 | 168.850 |
| | V#59 | 97.821 | 172.377 |
| | V#73 | 99.962 | 169.636 |
| | V#88 | 97.916 | 170.665 |
| | V#91 | 100.696 | 172.734 |
| | V#94 | 99.267 | 170.267 |

| | | Stem length (mm) | |
|---|---|---|---|
| | | Before stress | After stress |
| | WT | 93 | 95 |
| II | II#29 | 96 | 98 |
| IV | IV#17 | 90 | 93 |
| | IV#54 | 95 | 97 |
| | IV#68 | 93 | 96 |
| V | V#21 | 93 | 95 |
| | V#25 | 98 | 100 |
| | V#29 | 95 | 95 |
| | V#32 | 99 | 102 |
| | V#39 | 98 | 99 |
| | V#59 | 96 | 97 |
| | V#73 | 91 | 92 |
| | V#88 | 91 | 93 |
| | V#91 | 94 | 96 |
| | V#94 | 96 | 97 |

| | | Stem diameter (mm) | |
|---|---|---|---|
| | | Before stress | After stress |
| | WT | 6 | 6 |
| II | II#29 | 6 | 6 |
| IV | IV#17 | 6 | 7 |
| | IV#54 | 6 | 6 |
| | IV#68 | 6 | 7 |
| V | V#21 | 6 | 6 |
| | V#25 | 5 | 6 |
| | V#29 | 6 | 7 |
| | V#32 | 6 | 6 |
| | V#39 | 6 | 6 |
| | V#59 | 6 | 6 |
| | V#73 | 6 | 6 |
| | V#88 | 6 | 6 |
| | V#91 | 6 | 7 |
| | V#94 | 6 | 7 |

**Supplementary Table 5**: The raw data of RT-qPCR of *MKK2* expressions and phenotypic analyzes in WT and survived recovered events before and after salt stress. All events were analyzed in triplicates for RT-qPCR.
